# Rational screening for cooperativity in small-molecule inducers of protein–protein associations

**DOI:** 10.1101/2023.05.22.541439

**Authors:** Shuang Liu, Bingqi Tong, Jeremy W. Mason, Jonathan M. Ostrem, Antonin Tutter, Bruce K. Hua, Sunny A. Tang, Simone Bonazzi, Karin Briner, Frédéric Berst, Frédéric J. Zécri, Stuart L. Schreiber

## Abstract

The hallmark of a molecular glue is its ability to induce cooperative protein–protein interactions, leading to the formation of a ternary complex, despite weaker binding towards one or both individual proteins. Notably, the extent of cooperativity distinguishes molecular glues from bifunctional compounds, a second class of inducers of protein–protein interactions. However, apart from serendipitous discovery, there have been limited rational screening strategies for the high cooperativity exhibited by molecular glues. Here, we propose a binding-based screen of DNA-barcoded compounds on a target protein in the presence and absence of a presenter protein, using the “presenter ratio”, the ratio of ternary enrichment to binary enrichment, as a predictive measure of cooperativity. Through this approach, we identified a range of cooperative, noncooperative, and uncooperative compounds in a single DNA-encoded library screen with bromodomain (BRD)9 and the VHL–elongin C–elongin B (VCB) complex. Our most cooperative hit compound, **13-7**, exhibits micromolar binding affinity to BRD9 but nanomolar affinity for the ternary complex with BRD9 and VCB, with cooperativity comparable to classical molecular glues. This approach may enable the discovery of molecular glues for pre-selected proteins and thus facilitate the transition to a new paradigm of molecular therapeutics.

## Introduction

Small-molecule drugs that induce protein–protein associations possess novel properties with substantial current medical impact and tremendous potential for future therapeutic applications. These drugs have been demonstrated to stabilize, degrade, translocate, inhibit, or activate protein targets, to rewire cellular circuitry, and to restrict drug actions to targeted tissues co-expressing their two protein partners^1^. In some cases, the induced protein–protein associations exist within a single target protein (intramolecular), which underlies the ability to modulate otherwise challenging therapeutic targets such as the NLRP3 inflammasome^2^ and dysfunctional CFTR channel proteins in the genetic disorder cystic fibrosis^3^.

These “chemical inducers of proximity” (CIPs) comprise two major categories, namely molecular glues and bifunctional compounds. Molecular glues exhibit high cooperativity, as quantitatively defined by cooperativity factor α >>1, where α = *K*Dbinary/*K*Dternary (**Figure 1a**). In contrast, bifunctional compounds bind their two protein targets largely noncooperatively (α ∽ 1) and therefore exhibit the hook effect – a loss of ternary complex at high compound concentrations due to the formation of 1:1 compound/protein binary complexes. Bifunctional compounds such as protein-degrading PROTACs are appealing due to the conceptual simplicity of their design – link two binders with a connector – but their associated hook effect creates dosing challenges during clinical development and limits their therapeutic potential. Molecular glues overcome this limitation through their cooperative binding, which results in a wider therapeutic window.

**Figure 1.**
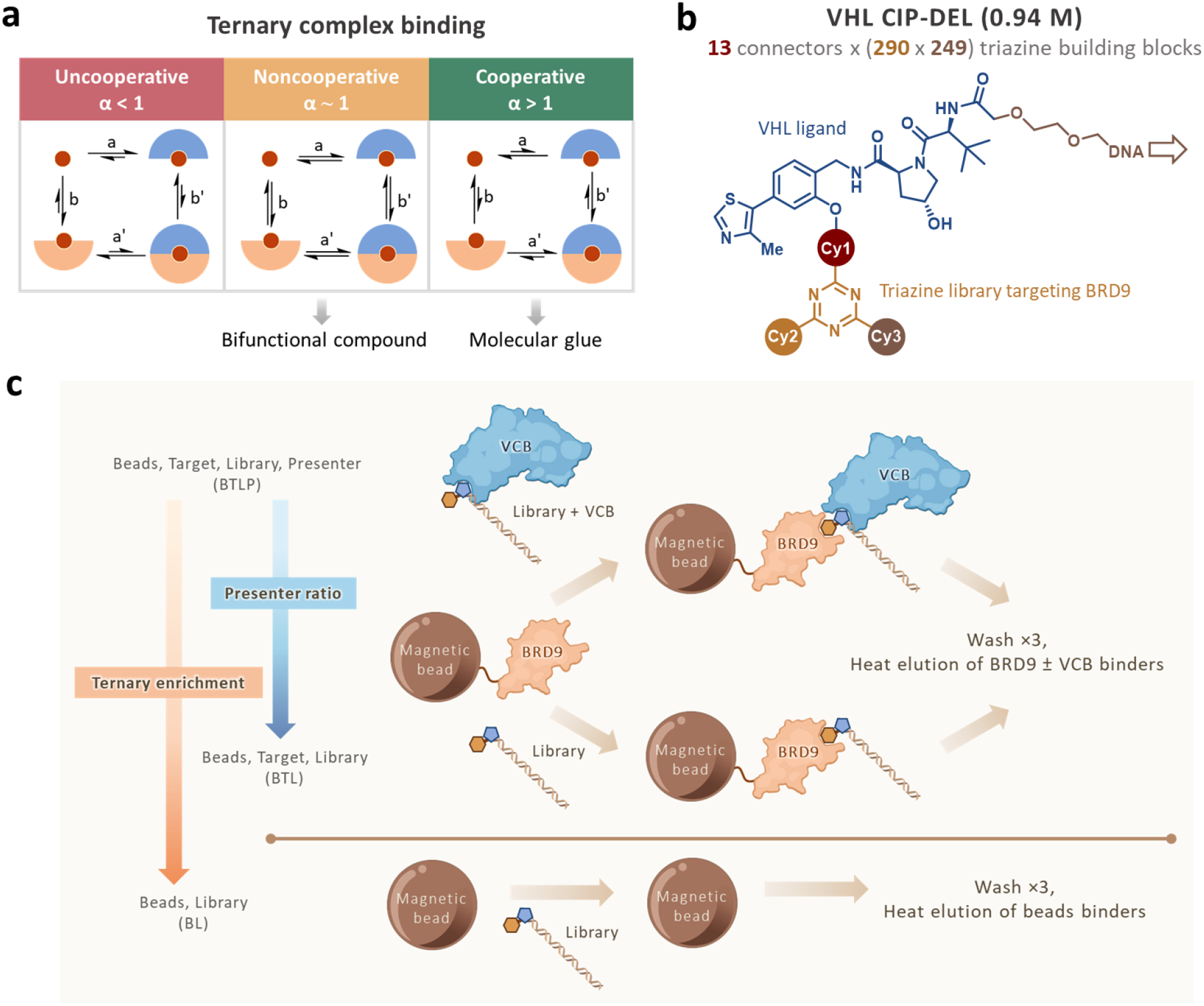
Screening of BRD9 using a VHL CIP-DEL in the presence and absence of VCB led to the discovery of cooperative, noncooperative and uncooperative compounds. **a**. A quantitative way to distinguish molecular glues and bifunctional compounds is by measuring the cooperativity factor, α, in their induced ternary complex binding. α is calculated by *K*_D_^binary^/*K*_D_^ternary^, i.e. either a/a’ or b/b’ as labeled. Bifunctional compounds usually have α ∽1 (noncooperative) and molecular glues have high cooperativity with α >1 (cooperative). **b**. A 0.94 million-member VHL CIP-DEL library used in the CIP-DEL screen. A VHL ligand was connected through 13 connectors of various lengths and structures (cycle 1) to a diversified triazine library with two appendage building blocks (cycles 2 and 3). **c**. The CIP-DEL screening workflow with the target protein, BRD9 and the presenter protein, VCB. BRD9 was immobilized on magnetic beads and incubated with the library [Beads, Target, Library (BTL)] or library preincubated with VCB [(Beads, Target, Library, Presenter (BTLP)]. Buffer instead of BRD9 was added to the magnetic beads in the beads only control [(Beads, Library (BL)]. Ternary enrichment was calculated by comparing BTLP with BL; presenter ratio was calculated by comparing BTLP with BTL.

CIPs induce the associations of two proteins, a target protein and a presenter protein. Typically, the target protein is subject to modulating effects, while the presenter protein serves as a tool protein that exerts functional effects on the target protein. The presence of tissue-specific or cell-specific presenter proteins enables high selectivity of molecular glues, as productive binding only occurs in the presence of both the target and presenter proteins. ‘Undruggable’ proteins can also be targeted by molecular glues, as demonstrated by the binding of WDB002 to an otherwise featureless coiled coil domain of CEP250, mediated by the presenter protein FKBP12^4^. Despite their therapeutic advantages, molecular glues have in general been challenging to design and have been most frequently identified serendipitously – by uncovering their ability to induce protein associations during mechanism-of-action studies of compounds having novel functional consequences on cells and in animals; for example, tacrolimus (FK506)^5^, sirolimus (rapamycin)^6,7^ and lenalidomide/thalidomide^8^.

There has been a tendency to differentiate bifunctional compounds from molecular glues based on their ‘binder-linker-binder’ structure, which is less recognizable in many molecular glues. However, the hallmark of molecular glues is their high cooperativity in inducing protein–protein associations to form ternary complexes. In this study, we aim to develop a rational approach to discover compounds with high cooperativity, regardless of their chemical structures. We recently described a screening technique using pre-selected target and presenter proteins to identify bifunctional compounds from a pool of DNA-barcoded compounds prepared using the split–pool synthesis strategy of DNA-encoded libraries (DELs). We used the VHL–elongin C–elongin B (VCB) E3 ligase complex as the presenter protein to assess the functional outcomes of induced associations in cells. The resulting noncooperative PROTACs were effective in forming ternary complexes in cells and degrading their pre-selected targets^9^. Here, we adapt this system to identify rationally compounds from DELs that induce protein associations of pre-selected targets and presenters cooperatively. The resulting functional compounds exhibit cooperativity *in vitro* and in cells comparable to classical molecular glues.

## Results

### Library design and screening workflow

We used a VHL-targeted CIP-DEL to screen for compounds that induce associations between BRD9 and VHL. The library was constructed analogously to the one previously described^9^, but used different vectors from the VHL ligand to attach to the DNA barcodes and connectors. In this CIP-DEL iteration, the diversified triazine-based library was connected to the VHL-targeting ligand via a phenolic oxygen (**Figure 1b**), in accordance with the reported BRD7/BRD9 degrader, VZ185^10^. The DNA headpiece and barcode were attached to the distal amide of the VHL-targeting ligand, which points towards the solvent exposed region in the crystal structure of a similar compound co-crystallized with the bromodomain of SMARCA2 and VCB (PDB 6HAX)^11^. The split-and-pool synthesis consisted of three cycles, where cycle 1 (cy1) included 13 connectors between the triazine library and the VHL-targeting ligand (**Supplementary Figure S1**), and cycle 2 (cy2) and cycle 3 (cy3) included 290 and 249 appendage building blocks on the triazine core, respectively, as previously described^9^. In total, the VHL-targeted CIP-DEL contains 938,730 library members, with DNA-encoded VZ185 (**Supplementary Figure S2**) added as a positive control at an equimolar amount to a single library member in the screen.

In the CIP-DEL screen, His-tagged BRD9 (target protein) was first immobilized to Co^2+^-coated magnetic beads. Immobilized BRD9 was added to either the CIP-DEL to form binary complexes [Beads, Target, Library (BTL)], or the CIP-DEL pre-incubated with VCB (presenter protein) to form ternary complexes [Beads, Target, Library, Presenter (BTLP)]. To control for nonspecific bead binders, [Beads, Library (BL)] was included where no protein was immobilized on beads or present in the buffer. The beads were then subjected to three washes before the proteins were heat-denatured to release the binders. The DNA barcodes of the binders were PCR-amplified and sequenced to obtain the counts of each library member in BL, BTL and BTLP. We modeled the count data as a Poisson sampling of barcodes, which normalizes the counts against the total sample counts while also taking into account the sequencing data uncertainty^12^. The binary enrichment was calculated by dividing BTL over BL, the ternary enrichment by dividing BTLP over BL, and the presenter ratio by dividing BTLP over BTL, essentially giving the ratio of ternary enrichment over binary enrichment (**Figure 1c**). *A higher presenter ratio suggests that the binding of the CIP-DEL compound to BRD9 is more dependent on the presence of VCB, thus implying higher cooperativity of the compound*. We hypothesized that we could use the presenter ratio to select small molecules with various degrees of cooperativity in their ternary complex binding to BRD9 and VCB.

### Screening results

The CIP-DEL screening results of BRD9 with and without VCB were displayed as a plot of ternary enrichment (BTLP/BL) versus presenter ratio (BTLP/BTL), whereby each datum point represents a CIP-DEL library member (**Figure 2a**). The enrichment and presenter ratio values were reported as the lower bound of the 95% confidence interval^9,12^. Interestingly, the enriched compounds for BRD9 showed a distribution that could be categorized into three groups: 1) compounds with high ternary enrichment and low presenter ratio, which were dominated by those with connectors 9 (9-series) and 12 (12-series); 2) compounds with medium ternary enrichment and medium presenter ratio, which were dominated by those with connector 11 (11-series); and 3) compounds with low ternary enrichment and high presenter ratio, which were dominated by those with connector 13 (13-series). Notably, all enriched connectors were relatively short and cyclic among the 13 connectors in the library (**Supplementary Figure S1**). We selected 16 library members (**9-1** to **9-4, 11-1, 11-2, 12-1, 13-1** to **13-9**; **Figure 2a** for plot, **Table 1** for summarized results, **Supplementary Figure S3** for compound structures) for off-DNA synthesis and validation using biophysical and cellular assays. The synthesis of compounds with connector 12 was unique, as it was attached to the VHL ligand via the carbon *para* to the anticipated phenolic oxygen, due to the distinct reactivity of the 4-(piperazin-1-yl)-benzyl tosylate (**Supplementary Figure S4**). Our subsequent discussion focuses on compounds with connectors 9, 11 and 13, which share a common scaffold.

**Figure 2.**
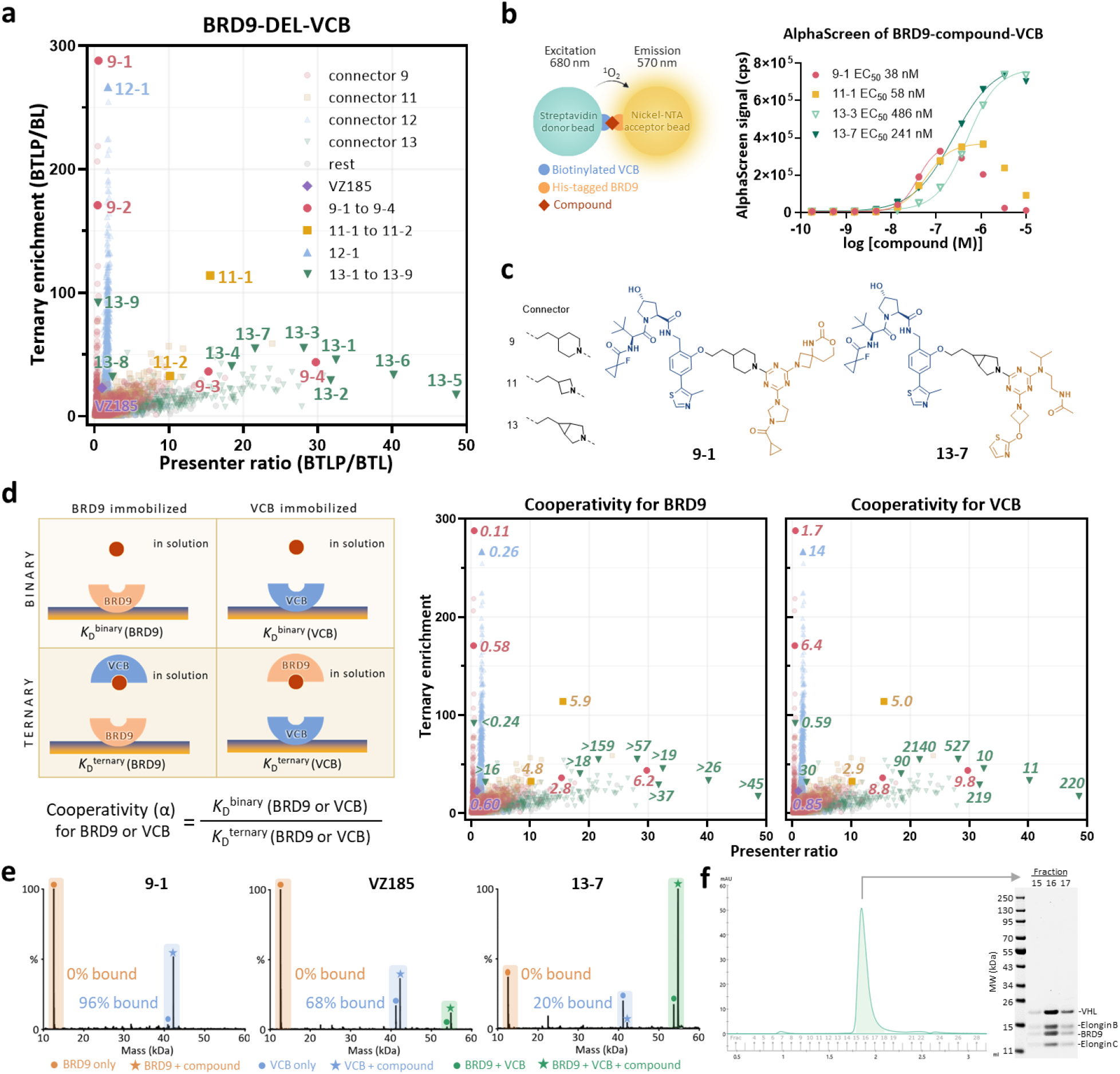
Biophysical characterization of uncooperative, noncooperative and cooperative compounds with BRD9 and VCB. **a**. A plot of ternary complex enrichment against presenter ratio. Enriched compounds of BRD9 in the presence of VCB can be divided into three categories: 1. High ternary enrichment and low presenter ratio, dominated by compounds with connectors 9 (coral) and 12 (blue); 2. Medium ternary enrichment and medium presenter ratio, dominated by compounds with connector 11 (yellow); 3. Low ternary enrichment and high presenter ratio, dominated by compounds with connector 13 (green). **b**. Compounds with relatively high presenter ratio, **13-3** and **13-7**, have higher maximum AlphaScreen signals than the compounds with relatively low presenter ratio, **9-1** and **11-1**, due to the onset of the hook effect at higher compound concentrations for the less cooperative **9-1** and **11-1**. Data are mean ± SD, n = 2 technical replicates. Buffer: 25 mM HEPES, pH 7.4, 150 mM NaCl, 0.05% Tween-20 (w/v). See ‘Methods’ for conditions. **c**. Chemical structures of connectors 9, 11 and 13, compounds **9-1** and **13-7. d**. Binary and ternary *K*_D_s can be measured by SPR with BRD9 immobilized in the absence or presence of 2 μM VCB in the analyte solution, or with VCB immobilized in the absence or presence of 20 μM BRD9 in the analyte solution. Cooperativity factor, α, is calculated by binary *K*_D_ over ternary *K*_D_. The measured cooperativity has a general correlation with the presenter ratio. Buffer: 20 mM HEPES, pH 7.5, 100 mM NaCl, 0.005% Tween-20 (w/v) and 0.2 mM TCEP. See ‘Methods’ for conditions. **e**. Deconvoluted non-denaturing MS spectra showing the formation of ternary complexes of BRD9 + compound + VCB (5 μM for all) for the highly cooperative compound, **13-7**. The noncooperative VZ185 and the uncooperative/noncooperative **9-1** showed reduced or no ternary complex despite potent binary binding to VCB. Buffer: 200 mM ammonium acetate, pH 7.5. See ‘Methods’ for conditions. **f**. Size exclusion chromatography analysis revealed a single peak corresponding to the ternary complex of BRD9, **13-7** and VCB (separated into VHL, elongin B and elongin C under the denaturing SDS-PAGE gel condition) when mixed at an equimolar amount. Buffer: 20 mM HEPES, 100 mM NaCl, pH 7.4.

**Table 1.** Profiling of compounds selected for off-DNA synthesis and validation. Cycle IDs are (connectors, triazine appendage building block 1, triazine appendage building block 2). See **Supplementary Figure S3** for compound structures. Ternary enrichment and presenter ratio values are presented as the most probable values with the lower and upper bounds of 95% confidence interval of a Poisson distribution. Cooperativity values for BRD9 and VCB were calculated based on binary and ternary *K*_D_s with immobilized BRD9 and VCB, respectively, as measured by SPR. See **Supplementary Figure S6** for SPR graphs. See **Supplementary Figure S9** for NanoBiT graphs and **Supplementary Figure S11** for HiBiT (± MLN4924) and CTG graphs.

| Compound | Cycle ID | Ternary enrichment | Presenter ratio | Cooperativity for BRD9 | Cooperativity for VCB | NanoBiT EC<sub>50</sub> (nM) | NanoBiT E<sub>max</sub> (%) | HiBiT DC<sub>50</sub> (nM) | HiBiT D<sub>max</sub> (%) |
| --- | --- | --- | --- | --- | --- | --- | --- | --- | --- |
| **9-1** | (9, 275, 131) | 404 (288-608) | 0.56 (0.54-0.57) | 0.11 | 1.7 | 904 | 363 | 4470 | 49 |
| **9-2** | (9, 275, 235) | 262 (171-448) | 0.46 (0.44-0.49) | 0.58 | 6.4 | 29 | 319 | 1590 | 80 |
| **9-3** | (9, 87, 174) | 58 (36-107) | 21 (15-32) | 2.8 | 8.8 | 51 | 404 | 1880 | 73 |
| **9-4** | (9, 126, 158) | 90 (44-273) | 56 (30-139) | 6.2 | 9.8 | 759 | 306 | 3650 | 78 |
| **11-1** | (11, 87, 174) | 248 (114-879) | 21 (16-29) | 5.9 | 5 | 517 | 285 | 4110 | 60 |
| **11-2** | (11, 197, 174) | 52 (33-93) | 14 (10-19) | 4.8 | 2.9 | 2980 | 421 | >30000 | 12 |
| **12-1** | (12, 275, 235) | 437 (267-841) | 1.9 (1.8-2.1) | 0.26 | 14 | 158 | 291 | 1120 | 47 |
| **13-1** | (13, 163, 109) | 77 (46-154) | 52 (32-96) | >19 | 10 | 726 | 878 | 3650 | 67 |
| **13-2** | (13, 253, 109) | 42 (29-64) | 48 (32-78) | >37 | 219 | 54 | 1065 | 582 | 77 |
| **13-3** | (13, 117, 161) | 91 (55-176) | 41 (28-65) | >57 | 527 | 33 | 1299 | 323 | 80 |
| **13-4** | (13, 117, 12) | 60 (40-97) | 25 (18-34) | >18 | 90 | 191 | 824 | 3960 | 59 |
| **13-5** | (13, 130, 109) | 25 (17-38) | 95 (49-255) | >45 | 220 | 195 | 1141 | 539 | 76 |
| **13-6** | (13, 18, 109) | 52 (34-90) | 68 (40-136) | >26 | 11 | 201 | 918 | 2760 | 61 |
| **13-7** | (13, 117, 156) | 107 (54-288) | 34 (22-58) | >159 | 2140 | 17 | 1452 | 108 | 82 |
| **13-8** | (13, 117, 185) | 44 (32-64) | 2.8 (2.4-3.1) | >16 | 30 | 464 | 841 | 3540 | 58 |
| **13-9** | (13, 52, 185) | 137 (92-224) | 0.54 (0.50-0.57) | <0.24 | 0.59 | >30000 | 205 | >30000 | 10 |
| **VZ185** | (14, 291, 250) | 166 (23-453) | 1.8 (1.0-3.1) | 0.6 | 0.85 | 5.5 | 847 | 1.9 | 77 |

### Compounds with high presenter ratios showed no hook effect

We first tested the ternary complex formation of the off-DNA compounds with purified recombinant BRD9 and VCB using AlphaScreen assays (**Figure 2b**, see **Supplementary Figure S5** for complete AlphaScreen data of all compounds). Compounds that induced ternary complexes with BRD9 immobilized on the donor beads and VCB on the acceptor beads exhibited luminescent AlphaScreen signals. Among the compounds tested, **9-1**, a compound with low presenter ratio (0.54), showed increasing AlphaScreen signals up to a concentration of 120 nM, but the signals decreased at higher concentrations due to the hook effect (**Figure 2b**). Similarly, **11-1**, a compound with medium presenter ratio (16), showed AlphaScreen signals over a broader concentration range but still exhibited the hook effect at concentrations above 1 μM (**Figure 2b**). However, **13-3** and **13-7**, both of which had a high presenter ratio (28 and 22 respectively) and high ternary enrichment within the 13-series, showed increasing AlphaScreen signals without any significant hook effect up to the highest tested concentration of 10 μM (**Figure 2b**). These results demonstrated that compounds with higher presenter ratios are more cooperative. Given that connectors 9, 11 and 13 are all short and cyclic (**Figure 2c**), it is difficult to predict the cooperativity of the tested compounds based on their chemical structures alone, underscoring the value of CIP-DEL screening for discovering cooperative compounds.

### Cooperativity of 13-7 is comparable to that of rapamycin

To quantify the cooperativity of the off-DNA compounds observed in AlphaScreen assays, we measured the cooperativity factor, α, using surface plasmon resonance (SPR) (**Figure 2d**, see **Supplementary Figure S6** for complete SPR data of all compounds). α of the compounds for BRD9 is the ratio of the binary *K*D for BRD9 to the ternary *K*D for BRD9. The binary dissociation constant (*K*D) for BRD9 was determined by immobilizing biotinylated BRD9 on a streptavidin sensor chip with off-DNA compounds in solution, and the ternary *K*D for BRD9 was determined by immobilizing biotinylated BRD9 on the chip with off-DNA compounds preincubated with tagless VCB in solution. For most of the 13-series compounds, binary *K*D values could not be accurately determined by SPR when titrating the compounds up to 20 μM (**Supplementary Figure S6b**), indicating weak binding to BRD9 alone and resulting in α values greater than a lower limit (**Figure 2d**). Among the compounds tested, **13-7** exhibited the highest cooperativity for BRD9 (α >159) (**Figure 2d**). The α values for VCB were similarly calculated by immobilizing biotinylated VCB on the chip with compound alone, or compound preincubated with tagless BRD9 in solution. In theory, α of the compounds for BRD9 should equate to that for VCB; in practice, different *K*Ds of the compounds to the protein in solution, concentration of the protein in solution, and immobilization of the protein to a surface resulted in imperfect experimental conditions and in some cases, different α values obtained. Importantly, the relative trend remained consistent where the 9-series with low presenter ratios showed low α values, whereas the 13-series with high presenter ratios showed α as high as 2140 for **13-7**, which is comparable to the cooperativity of the classical molecular glue, rapamycin (α = 2170)^13^ binding to FKBP12 and the FRB domain of mTOR. We focus our following discussion on **13-7**, which has the highest cooperativity for both BRD9 and VCB.

### 13-7 induced less binary but more ternary complexes

To measure directly the proportion of binary and ternary complexes induced by various compounds, we used non-denaturing mass spectrometry (MS) to detect any non-covalent interactions in a mixture of equimolar off-DNA compound, BRD9 and VCB. The deconvoluted non-denaturing mass spectra for the uncooperative/noncooperative **9-1**, noncooperative **VZ185** (the reported bifunctional PROTAC of BRD9 and VCB^10^), and the cooperative **13-7** were shown in **Figure 2e** (see **Supplementary Figure S7** for complete MS data of all compounds). No binary binding to BRD9 was observed for any of the three compounds at the MS conditions. Binary binding to VCB was the strongest for **9-1** (96% bound), followed by **VZ185** (68% bound) and **13-7** (20% bound). However, despite potent binary binding of **9-1** to VCB, no MS peak corresponding to the ternary complex of **9-1** with BRD9 and VCB was observed. In contrast, a small population (12% relative to the mass intensity of unbound BRD9) of ternary complex with BRD9 and VCB was observed for VZ185, while a large population (270% relative to the mass intensity of unbound BRD9) of ternary complex with BRD9 and VCB was observed for the cooperative **13-7**, despite its weak binary binding to VCB. The gas-phase non-denaturing MS results were further validated by a solution-phase experiment where a mixture of equimolar **13-7**, BRD9 and VCB were injected into a size exclusion column, yielding a single elution peak containing BRD9 and VCB, with no significant binary complexes or unbound proteins observed (**Figure 2f**). The results suggest that **13-7** cooperatively induces the formation of ternary complexes with BRD9 and VCB, resulting in their co-elution (see **Supplementary Figure S8** for control experiments).

### 13-7 induced ternary complexes without hook effect in cells

To evaluate the cooperativity of the off-DNA compounds in cells, we conducted dose-dependent NanoBiT assays^14^ using SmBiT-BRD9 and VHL-LgBiT fusion proteins in HEK293T cells (see **Supplementary Figure S9** for complete BRD9 NanoBiT data of all compounds). Based on the measured luminescence signals, the uncooperative/noncooperative **9-1** induced a relatively low amount of ternary complexes with BRD9 and VHL in cells, with a half-maximal effective concentration (EC50) of 904 nM (**Figure 3a**). The noncooperative **VZ185**, on the other hand, induced more ternary complexes with a low EC50 of 5.5 nM, but with a hook effect observed above 1 μM. In contrast, the cooperative **13-7**, which showed cooperative behavior with purified BRD9 and VCB proteins in the AlphaScreen assays (**Figure 2b**), induced dose-dependent formation of ternary complexes in cells with EC50 of 17 nM without any significant hook effect up to the highest tested concentration of 30 μM (**Figure 3a**).

**Figure 3.**
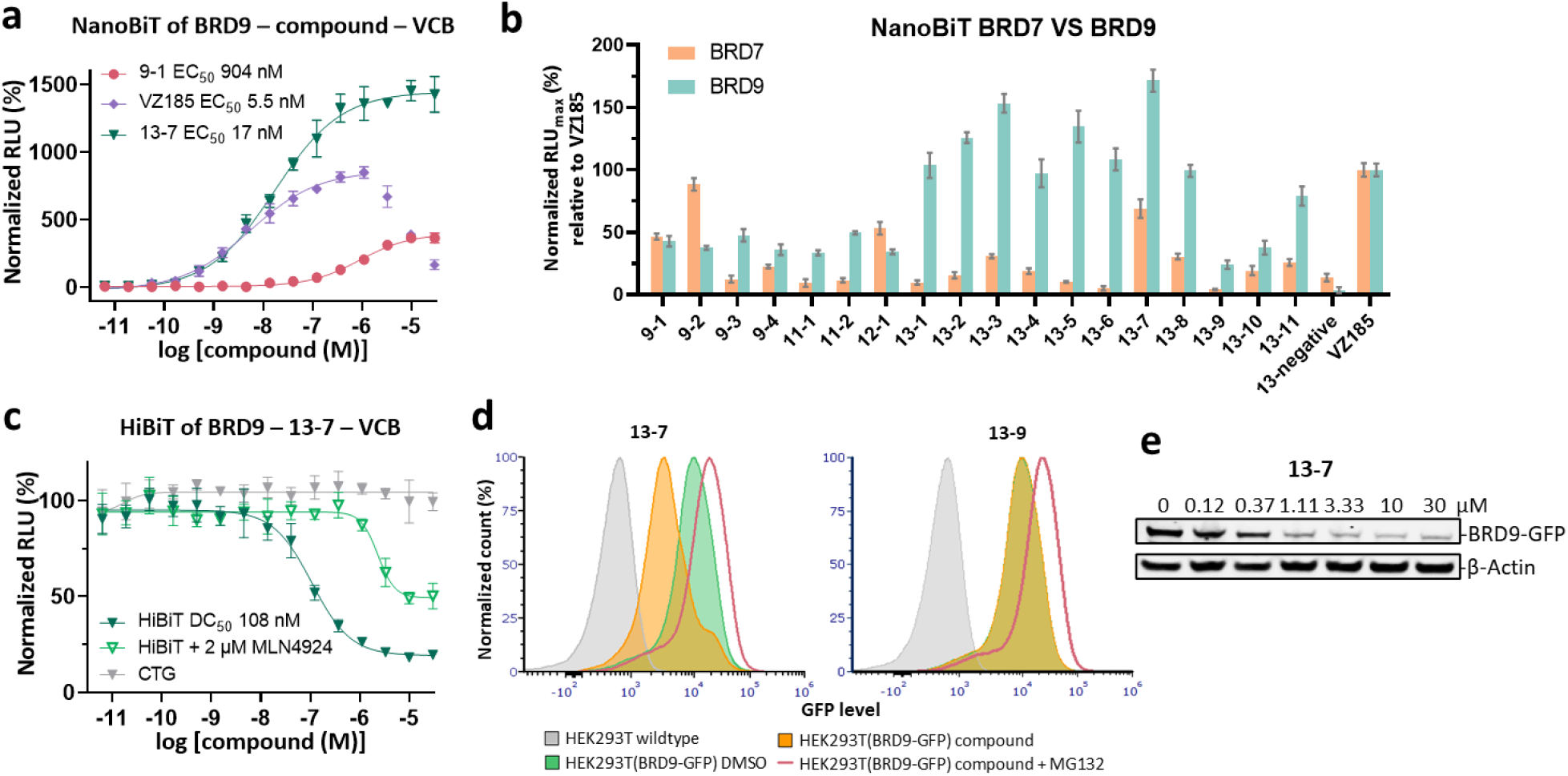
Cellular evaluation of the ternary complex formation of 13-7 with BRD9 and VCB using NanoBiT assays, and BRD9 degradation induced by 13-7 using HiBiT assays, FACS and western blot analysis. **a**. Cooperative **13-7** induced more ternary complexes with BRD9 and VCB in HEK293T cells than their noncooperative and uncooperative counterparts, VZ185 and **9-1**, as measured by NanoBiT assays. Data are mean ± SD, n = 4 technical replicates. No hook effect was observed for **13-7** at the highest tested concentration of 30 μM. **b**. Comparison of maximum ternary complex induced by compounds with BRD9/VCB and with BRD7/VCB, normalized with 100% being the highest ternary complex levels observed with VZ185, as measured by NanoBiT assays. Data are mean ± SD, n = 4 technical replicates. The cooperative compounds generally showed a greater selectivity towards a ternary complex with BRD9 over BRD7. **c. 13-7** induced degradation of BRD9 in HEK293T cells and the degradation was partially reversed by neddylation inhibitor MLN4924 (2 μM), as measured by HiBiT assays. Data are mean ± SD, n = 4 technical replicates. **d**. FACS analysis in HEK293T cells showed that **13-7**, but not **13-9**, induced degradation of BRD9 and this effect was reversed by the addition of proteasomal inhibitor MG132. **e**. Western blot analysis of BRD9 degradation in HEK293T cells without a hook effect being observed up to the highest tested concentration of 30 μM.

### Cooperative compounds showed better selectivity for close homologs

**VZ185** has been shown to be a dual degrader of BRD9 and its close homolog, BRD7^10^. While BRD9 is a promising therapeutic target for synovial sarcoma^15^, non-selective degradation of BRD7 could have undesirable consequences. To investigate whether cooperativity could improve selectivity of the off-DNA compounds in inducing ternary complexes with BRD9/VCB over BRD7/VCB, we conducted NanoBiT assays with SmBiT-BRD7 in place of SmBiT-BRD9 (see **Supplementary Figure S10** for complete BRD7 NanoBiT data of all compounds). The maximum effect (Emax) values were normalized relative to VZ185, which was set to 100% for both BRD9 and BRD7. The uncooperative/noncooperative **9-1, 9-2** and **12-1** showed no selectivity for BRD9 over BRD7 (**Figure 3b**). In contrast, the cooperative 13-series demonstrated high ternary complex formation with BRD9/VCB and varying degrees of selectivity for BRD9/VCB over BRD7/VCB (**Figure 3b**). Notably, compounds **13-5** and **13-6** exhibited the highest selectivity for BRD9/VCB over BRD7/VCB, with ratios of 13-fold and 21-fold, respectively, compared to VZ185 (**Figure 3b**). **13-5** and **13-6** also had the highest presenter ratios based on the screening results (**Figure 2a, Table 1**).

### 13-7 degrades BRD9 without hook effect in cells

As the presenter protein in our CIP-DEL screen was VCB, BRD9 degradation was assessed using HiBiT assays^16^ as a functional consequence of compound-induced ternary complex formation with BRD9/VCB (see **Supplementary Figure S11** for complete HiBiT data of all compounds). Although less potent than VZ185 (half-maximal degradation concentration (DC50) 1.9 nM), **13-7** degraded BRD9-HiBiT fusion proteins in HEK293T cells without the presence of a hook effect, with a DC50 of 108 nM (**Figure 3c**). The BRD9 degradation induced by **13-7** was at least partly VHL dependent, as demonstrated by the shift of the DC50 to 2.3 μM in the presence of the neddylation inhibitor, MLN4924^17^ (**Figure 3c**). Cell viability was not compromised at all tested concentrations of **13-7**, as confirmed by the CellTiter-Glo assay (**Figure 3c**), indicating that the decrease in HiBiT signals was mainly due to BRD9 degradation and not cellular toxicity. The calculation of degradation selectivity of the off-DNA compounds using HiBiT assays requires caution, as we could not solely attribute the maximal degradation (Dmax) to VHL dependence for some compounds (**Supplementary Figure S11**). Additional HiBiT data including VHL ligand competition and ubiquitin dependency (MLN7243) are provided in **Supplementary Figures S12 and S13**.

To verify our HiBiT results, we used fluorescence-activated cell sorting (FACS) to measure the degradation of BRD9-GFP fusion proteins in response to selected compounds in HEK293T cells (**Figure 3d**). BRD9-GFP was cloned in-frame with internal ribosomal entry site (IRES)-mCherry for simultaneous expression of mCherry, allowing us to exclude expression level changes as a source of any reduction in GFP signal. Addition of **13-7** (10 μM) to cells overexpressing BRD9-GFP resulted in reduced GFP signals intensity without effect on mCherry signal, consistent with loss of BRD9-GFP protein (**Figure 3d**). Co-treatment with the proteasome inhibitor MG132 prevented this effect, demonstrating that **13-7** induces proteasomal degradation of BRD9-GFP. We performed western blot analysis to confirm the absence of hook effect in the degradation of BRD9 induced by **13-7**. We detected the level of BRD9-GFP at varying concentrations of **13-7** and did not observe a significant hook effect up to the highest tested concentration of 30 μM (**Figure 3e**).

We also investigated **13-9** which showed high ternary enrichment but low presenter ratio and was an outlier of the 13-series based on the screening results (**Figure 2a, Table 1**). **13-9** was uncooperative (α <1) by SPR (**Figure 2d**), suggesting that its ternary complex formation with BRD9 and VCB is disfavored. As expected, we observed no degradation of BRD9-HiBiT with up to 30 μM **13-9** by HiBiT assays (**Supplementary Figure S11**) and no degradation of BRD9-GFP at 10 μM **13-9** by FACS (**Figure 3d**).

## Discussion

Rational screening for cooperative molecular glues in a high-throughput manner remains a challenging task in drug discovery. In this study, we have demonstrated the feasibility of using a binding-based CIP-DEL approach to identify compounds that induce ternary complex formation and target degradation cooperatively. By screening a library of approximately a million DNA-barcoded compounds against BRD9 with and without the VCB protein complex, we identified compounds that fall under opposite extremes: the 9-series with high ternary enrichment but low presenter ratio, and the 13-series with low ternary enrichment but high presenter ratio.

Our subsequent analysis of these compounds, which have similar chemical structures but varying cooperativity for BRD9 and VCB, reveals that the choice of connectors (e.g. connector 9 versus 13) between the BRD9 targeting ligand and the VHL ligand plays a dominant role in determining cooperativity of the compounds (**Figure 2a**). While the choice of appendage building blocks in the BRD9 targeting ligand plays a less dominant role in determining cooperativity, it can occasionally have a drastic effect on improving (**9-3** and **9-4**, cooperative despite being the 9-series) or reducing (**13-9**, uncooperative despite being the 13-series) cooperativity.

Interestingly, the uncooperative/noncooperative **9-1** behaves differently from the cooperative **13-7** for both binding measured by biophysical methods and functional effect in cells. Although **9-1** has high binary affinity to BRD9, it is less capable of inducing ternary complex formation with BRD9/VCB (**Figures 2b, 2e, 3a**) and degrading BRD9 (**Supplementary Figure S11**). By contrast, **13-7** has low binding affinity to BRD9 alone but demonstrated the highest degree of ternary complex formation (**Figures 2b, 2e, 3a**) and BRD9 degradation in cells (**Supplementary Figure S11**). These findings emphasize that binary affinity to the target protein is less consequential for ternary complex formation and target degradation compared to cooperativity and ternary *K*D.

Furthermore, our AlphaScreen and NanoBiT assay results demonstrated that cooperative compounds identified in this study are less susceptible to the hook effect (**Figures 2b, 3a**), similar to classical molecular glues. Minimizing the hook effect could widen the therapeutic window and mitigate dosing challenges, highlighting the potential advantages of cooperative compounds in therapeutic applications.

The presenter ratios based on the screening results were observed to have a positive correlation with cooperativity measured by SPR (Spearman’s correlation= 0.84, Pearson’s correlation= 0.40, calculated based on lower bound of presenter ratios and cooperativity for BRD9 in **Table 1**). Notably, compounds with high presenter ratios exhibited greater cooperativity (**13-7** and **13-3**) and greater selectivity towards the target BRD9 over its close homolog BRD7 (**13-5** and **13-6, Figure 3b**). Among them, the most cooperative compounds, **13-7** and **13-3**, also have the highest ternary enrichment. Thus, it is advisable to consider both high presenter ratios and ternary enrichment values in identifying cooperative compounds.

As we strive to extend the scope of our findings, ongoing investigations are in progress to validate the generality of analyzing data in this manner for systems other than BRD9/VCB and to explore the potential benefits of utilizing monovalent molecules in the CIP-DEL for identifying molecular glues with improved drug properties. These endeavors aim to facilitate the rational discovery of molecular glues for next-generation therapeutics.

## Methods

### Recombinant protein preparation

#### i) His-tagged BRD9

The plasmid pET-22b(+) containing the recombinant BRD9 gene (E135-A242) fused with an N-terminal His6 tag, via Ndel and Xhol restriction sites, was expressed in *Escherichia coli* (*E. coli*) BL21(DE3) cells. A starter culture was grown in LB media supplemented with ampicillin (50 μg/mL) at 37 °C overnight. The starter culture (1:100 v/v) was used to inoculate 3 L of LB media supplemented with ampicillin (50 μg/mL) and grown at 37 °C to an OD600 nm= 0.6. Expression was induced with 0.5 mM isopropyl-β-D-1-thiogalactopyranoside (IPTG) at 20 °C for 16 hr. The cells were collected by centrifugation (Thermo Scientific™ Sorvall™ LYNX 6000 centrifuge) at 8,000 ×g for 10 min. The pellets were suspended in lysis buffer containing 20 mM HEPES, 500 mM NaCl, pH 7.4, 1% Triton X-100, lysozyme (1 mg/mL), benzonase, 1 mM TCEP and 1 mM phenylmethylsulfonyl fluoride (PMSF). The cells were lysed on ice by sonication (50% amplitude, 3 s pulse ‘on’ and 3 s pulse ‘off’, for total 5 min ‘on’ time). The cell debris was precipitated by centrifugation (Thermo Scientific™ Sorvall™ LYNX 6000 centrifuge) at 30,000 ×g for 60 min. The cell lysates were filtered through a 0.45 μm syringe filter and purified by affinity chromatography using a 5 mL HisTrap HP column (GE Healthcare Life Sciences) with 5 CV of wash buffer (20 mM HEPES, 500 mM NaCl, pH 7.4), 10 CV of 4% elution buffer (20 mM HEPES, 500 mM NaCl, 500 mM imidazole, pH 7.4), and a gradient of 4-100% elution buffer over 10 CV. The fractions containing BRD9 were pooled, concentrated and further purified by size exclusion chromatography (SEC) using a Superdex 16/600 75 pg column in 20 mM HEPES, 100 mM NaCl, pH 7.4. Pure fractions of BRD9 were pooled, concentrated, aliquoted and flash frozen for storage at -80 °C.

#### ii) Tagless BRD9

Tagless BRD9 was produced in a similar manner as His-tagged BRD9, but with a 3C cleavage site between the N-terminal His6 tag and the BRD9 gene. After the HisTrap purification, BRD9 protein was buffer exchanged to 20 mM HEPES, 500 mM NaCl, pH 7.4 and the His6 tag was cleaved off by gently shaking the protein (10 mL) with HRV 3C protease (Pierce™ 88946, 90 μL) at 4 °C for 14 hr. Complete cleavage was checked by MS before the tagless BRD9 undergoing a second HisTrap purification where the protein was washed with 5 CV of wash buffer, 5 CV of 4% elution buffer, and a gradient of 4-50% elution buffer over 10 CV. The fractions containing tagless BRD9 were pooled, concentrated and further purified by SEC as described for His-tagged BRD9.

#### iii) Biotinylated BRD9

Biotinylated BRD9 was produced in a similar manner as tagless BRD9, but with a 3C cleavage site followed by an Avi tag between the N-terminal His6 tag and the BRD9 gene. After the HisTrap purification, BRD9 protein was buffer exchanged to 20 mM HEPES, 100 mM NaCl, pH 7.4, supplemented with 5 mM MgCl2, 2 mM ATP, 300 μM biotin, BirA (R&D systems 10464-BA, 90 μL) and HRV 3C protease (Pierce 88946, 90 μL) for concurrent His tag cleavage and biotinylation at 4 °C for 14 hr. Complete cleavage and biotinylation was checked by MS before the second HisTrap and SEC purification as described for tagless BRD9.

#### iv) His-tagged VCB

The plasmid pETite (Lucigen) containing the recombinant VHL (M54 – D213, fused with an His8 tag and an HRV 3C protease site at the N-terminus), elongin C (M17 – C112) and elongin B (M1 – K104) (VCB complex) genes was co-expressed in the *E. coli* BL21 Star™ (DE3) strain supplemented with pRARE. The plasmid and glycerol stocks were generous gifts from Dustin Dovala (NIBR, Emeryville, USA) and Michael Romanowski (NIBR, Cambridge, USA) respectively. A starter culture was grown in TB media supplemented with chloramphenicol (25 μg/mL) and kanamycin (50 μg/mL) at 37 °C overnight. The starter culture (1:100 v/v) was used to inoculate 3 L of TB media supplemented with chloramphenicol (25 μg/mL) and kanamycin (50 μg/mL) and grown at 37 °C to an OD600 nm= 1.0. Expression was induced with 0.5 mM IPTG at 20 °C for 15 hr. The cells were collected by centrifugation (Thermo Scientific™ Sorvall™ LYNX 6000 centrifuge) at 8,000 ×g for 10 min. The pellets were suspended in lysis buffer containing 20 mM HEPES, 500 mM NaCl, pH 7.4, 1% Triton X-100, lysozyme (1 mg/mL), benzonase, 1 mM TCEP and 1 mM PMSF. The cells were lysed on ice by sonication (50% amplitude, 3 s pulse ‘on’ and 3 s pulse ‘off’, for total 5 min ‘on’ time). The cell debris was precipitated by centrifugation (Thermo Scientific™ Sorvall™ LYNX 6000 centrifuge) at 30,000 ×g for 60 min. The cell lysates were filtered through a 0.45 μm syringe filter and purified by affinity chromatography using a 5 mL HisTrap HP column (GE Healthcare Life Sciences) with 5 CV of wash buffer (20 mM HEPES, 500 mM NaCl, pH 7.4), 10 CV of 4% elution buffer (20 mM HEPES, 500 mM NaCl, 500 mM imidazole, pH 7.4), and a gradient of 4-100% elution buffer over 10 CV. The fractions containing VCB were pooled, concentrated and further purified by size exclusion chromatography (SEC) using a Superdex 26/600 200 pg column in 20 mM HEPES, 100 mM NaCl, pH 7.4. Pure fractions of VCB were pooled, concentrated, aliquoted and flash frozen for storage at -80 °C.

#### v) Tagless VCB

Tagless VCB was produced in a similar manner as His-tagged VCB with the following exceptions. The same plasmid for His-tagged VCB was co-expressed in *E. coli* BL21 (DE3) cells and the TB media was supplemented with kanamycin (50 μg/mL). After the HisTrap purification, VCB protein was buffer exchanged to 20 mM HEPES, 500 mM NaCl, pH 7.4 and the His8 tag was cleaved off by gently shaking the protein (10 mL) with HRV 3C protease (Pierce™ 88946, 90 μL) at 4 °C for 14 hr. Complete cleavage was checked by MS before the tagless VCB undergoing a second HisTrap purification where the protein was washed with 5 CV of wash buffer, 5 CV of 4% elution buffer, and a gradient of 4-50% elution buffer over 10 CV. The fractions containing tagless VCB were pooled, concentrated and further purified by SEC as described for His-tagged VCB.

#### vi) Biotinylated VCB

Biotinylated VCB was produced in a similar manner as tagless VCB with the following exceptions. The plasmid pETite (Lucigen) containing the recombinant VHL (M54 – D213, fused with a His8 tag, an HRV 3C protease site and an Avi tag at the N terminus), elongin C (M17 – C112) and elongin B (M1 – Q118) (VCB complex) genes was co-expressed in *E. coli* BL21 (DE3) cells. The plasmid was a generous gift from Dustin Dovala (NIBR, Emeryville, USA). After the HisTrap purification, VCB protein was buffer exchanged to 20 mM HEPES, 100 mM NaCl, pH 7.4, supplemented with 5 mM MgCl2, 2 mM ATP, 100 μM biotin, BirA (R&D systems 10464-BA, 100 μg) and HRV 3C protease (Pierce 88946, 100 μL) for concurrent His tag cleavage and biotinylation at 4 °C for 18 hr. Complete cleavage and biotinylation was checked by MS before the second HisTrap and SEC purification as described for tagless VCB.

### DNA-encoded library screening

The DEL screening was performed with 3 different selection conditions in parallel: 1) beads-only control (BL: beads, library), 2) BRD9^BD^ -VCB (BTL: beads, target protein, library), and 3) BRD9^BD^ + VCB (BTLP: beads, target protein, library, presenter protein). Proteins from commercial sources were used for the DEL screening while the subsequent validation was performed using proteins prepared in-house following the method described above. His-tagged BRD9 was purchased from BPS Bioscience (31090). Tagless VCB complex was purchased from R&D Systems (E3-600) excluding DTT in the supplied buffer to ensure compatibility with the His-Tag Dynabeads. The screening buffers used are ‘B (binding) Buffer’ containing 25 mM HEPES pH 7.4, 150 mM NaCl, 0.05% Tween-20 (w/v), and ‘S (selection) Buffer’ containing 25 mM HEPES, pH 7.4, 150 mM NaCl, 0.05% Tween-20 (w/v) and 0.3 mg/mL Ultrapure Salmon Sperm DNA (ThermoFisher Scientific 15632011).

His-Tag Dynabeads (Invitrogen 10103D, 10 μL per sample) were manually washed three times with B buffer and resuspended in B buffer (100 μL per sample). Using a KingFisher Duo Prime (Thermo Scientific 5400110) in a 96-well deepwell plate (Thermo Scientific 95040452) at r.t., the washed beads were transferred to His-tagged BRD9 (10 μM in B buffer, 100 μL per sample) for BRD9 immobilization (1 hr, medium mix). The beads were washed once with B buffer (200 μL) and twice with S buffer (200 μL) (3 min each, medium mix). The beads were then incubated (1 hr, medium mix) with the library (10 million copies per library member, 156 nM, 100 μL in S buffer) only, or the library pre-incubated with tagless VCB (2 μM in S buffer). The positive control compound, DNA-encoded VZ185, was spiked into the library at an equimolar amount to a single library member in the screen i.e. 10 million copies of this single compound per selection. The beads were then washed once with S buffer (200 μL) and twice with B buffer (200 μL) (3 min each, medium mix). Finally, the beads were transferred to B buffer (100 μL) and heated (95 °C, 5 min) to elute bound DEL members into the supernatant. 20 μL of the supernatant was restriction digested by 0.1 μL StuI (NEB R0187) in 56.5 μL 1× SmartCutter buffer (NEB B7204S) per sample (37 °C, 1 hr) and cleaned up using the ChargeSwitch PCR Clean-Up Kit (Thermo Scientific CS12000). Successful immobilization of the BRD9 protein was checked by resuspending the beads in 133.3 μL of NuPAGE LDS Sample Buffer (1×) supplemented with 3.33 μL of 0.5 M TCEP, heating at 95°C for 5 min, and loading the supernatant onto an SDS-PAGE gel. The barcodes of the eluted DEL were PCR amplified using 3 μL i5 index primer (10 μM stock in water), 3 μL i7 index primer (10 μM stock in water), 19 μL cleaned up elution samples, and 25 μL Invitrogen Platinum™ Hot Start PCR Master Mix (2×) (Invitrogen 13000012). The PCR method is as follows: 95 °C for 2 min; 19 cycles of 95 °C (15 s), 55 °C (15 s), 72 °C (30 s); 72 °C for 7 min; hold at 4 °C. The PCR products were cleaned up using the ChargeSwitch PCR Clean-Up Kit, pooled in equimolar amounts, and the 187 bp amplicon was gel purified using a 2% E-Gel™ EX Agarose Gels (ThermoFisher Scientific G401002) and QIAquick Gel Extraction Kit (Qiagen 28704). The DNA concentration was measured using the Qubit dsDNA BR assay kit and sequenced using a HiSeq SBS v4 50 cycle kit (Illumina FC-401-4002) and HiSeq SR Cluster Kit v4 (Illumina GD-401-4001) on a HiSeq 2500 instrument (Illumina) in a single 50-base read with custom primer CTTAGCTCCCAGCGACCTGCTTCAATGTCGGATAGTG and 8-base index read 32 using custom primer CTGATGGAGGTAGAAGCCGCAGTGAGCATGGT. The sequencing data were analyzed as previously described^9^ and reported as the lower bound of 95% confidence interval unless stated otherwise.

### AlphaScreen assays

AlphaScreen assays were carried out in AlphaPlate-384 plates (PerkinElmer 6008350) with a total volume of 20 μL sample mixture in 25 mM HEPES, pH 7.4, 150 mM NaCl, 0.05% Tween-20 (w/v) in each well. All experiments were conducted in duplicates. His-tagged BRD9 (5 μL, final 50 nM), biotinylated VCB (5 μL, final 20 nM), compounds (5 μL, 3-fold serial dilution from a top concentration of final 10 μM, 1% DMSO), and a mixture of streptavidin donor beads (PerkinElmer 6760619C, 2.5 μL, final 0.02 mg/mL) and nickel chelate (Ni-NTA) acceptor beads (PerkinElmer 6760619C, 2.5 μL, final 0.02 mg/mL) were sequentially added to the wells of the plates. The mixture was incubated for 1 hr at r.t. in the dark. Fluorescence readings at excitation 680 nm and emission 570 nm were measured on an EnVision 2105 plate reader (PerkinElmer). EC50 values were calculated from nonlinear regression curve fits (excluding any data points above the hook effect concentration, if any) using GraphPad Prism v9.5.

### Surface Plasmon Resonance

All SPR experiments were performed on a Biacore T200 or 8K instrument (GE Healthcare) at 25 °C using a running buffer containing 20 mM HEPES, pH 7.5, 100 mM NaCl, 0.005% tween-20, 0.2 mM TCEP, 1% or 2% DMSO. The streptavidin sensor chip (Cytiva BR100531) was preconditioned with 1M NaCl and 40 mM NaOH (60 s ×3 cycles, 100 μL/min) and running buffer (60 s, 100 μL/min). The run has a startup run of 12 cycles using the running buffer, flow rate of 50 μL/min, contact time of 70 s and dissociation time of 140– 200 s, with two negative controls using the running buffer between each compound. The syringe was washed with 50% DMSO in water between injections. *K*D values were determined using the steady state affinity mode of the Biacore Evaluation Software. Cooperativity (α) was calculated by dividing the binary *K*D by the ternary *K*D, i.e. (i)/(ii) or (iii)/(iv) below.

1. Binary *K*D for BRD9 Biotinylated BRD9 (RU ∽500) was immobilized to the streptavidin chip. The compounds were 3-fold serial diluted with the running buffer (2% DMSO) from a top concentration of 20 μM (4 μL of 1 mM compound in DMSO added to 196 μL of running buffer without DMSO).
2. Ternary *K*D for BRD9 Biotinylated BRD9 (RU ∽55) was immobilized to the streptavidin chip. The compounds were 3-fold serial diluted with the running buffer (1% DMSO) containing 2 μM tagless VCB from a top concentration of 2 μM (2 μL of 0.2 mM compound in DMSO added to 198 μL of running buffer without DMSO and with 2.02 μM tagless VCB).
3. Binary *K*D for VCB Biotinylated VCB (RU ∽1700) was immobilized to the streptavidin chip. The compounds were 3-fold serial diluted with the running buffer (1% DMSO) from a top concentration of 2 μM (2 μL of 0.2 mM compound in DMSO added to 198 μL of running buffer without DMSO).
4. Ternary *K*D for VCB Biotinylated VCB (RU ∽140) was immobilized to the streptavidin chip. The compounds were 3-fold serial diluted with the running buffer (1% DMSO) containing 20 μM tagless BRD9 from a top concentration of 2 μM (2 μL of 0.2 mM compound in DMSO added to 198 μL of running buffer without DMSO and with 20.2 μM tagless BRD9). The sample compartment temperature was lowered to 15°C for better BRD9 stability.

### Non-denaturing Mass Spectrometry

Native mass spectrometry experiments were carried out using a Thermo Q-Exactive UHMR Mass Spectrometer (Thermo Scientific). Tagless BRD9 and tagless VCB stock solutions were buffer exchanged into 200 mM ammonium acetate, pH 7.5 using a Micro Bio-Spin™ 6 column (Bio-Rad 732-6221). The compound of interest (5 μM, 1% DMSO) was mixed with tagless BRD9 (5 μM) and/or tagless VCB (5 μM) in 200 mM ammonium acetate, pH 7.5. The samples (20 μL) were then pipetted into wells of a 384-well plate and introduced into the mass spectrometer using an automated chip-based electrospray ionization (ESI) source (TriVersa Nanomate, Advion, Ithaka, NY, USA) with ESI voltage of 1.8 kV and spray backing gas pressure of 1.1 psi. Spectra were obtained in positive ion mode with a resolving power of 6,250 (at m/z 400) and a mass range of 1000-7000 m/z. The capillary temperature was set to 200 °C. The in-source trapping voltage was set to -5 V to help desolvate the ions while maintaining the compound binding to proteins. The detector was set to low m/z mode while the ion transfer region was manually adjusted to a setting between low and high m/z mode with a trapping gas pressure of 7.0. Spectra were deconvoluted and relevant peaks were integrated to determine percent binary and ternary complexes formed using UniDec software^18^.

### Size exclusion chromatography for ternary complex measurement

The Superdex 200 Increase 5/150 GL column was equilibrated with 20 mM HEPES, 100 mM NaCl, pH 7.4, 1% DMSO (v/v) on a ÄKTA™ pure micro instrument. A mixture (40 μL) of tagless BRD9 (30 μM), tagless VCB (30 μM) and compound 13-7 (30 μM, 2% DMSO) was then injected into the column and the sample was eluted at a flow rate of 0.2 mL/min. UV was detected at 280 nm.

### NanoBiT assays

15 mL of HEK293 cells were plated at 0.625 × 10^6^ cells / mL in DMEM (Gibco 11995065) supplemented with 10% fetal bovine serum (FBS) in a T75 flask. 15 μg CMV-driven SmBiT-BRD9 (E135-A252) or SmBiT-BRD7 (E129-L250) plasmid, 15 μg HSVtk-driven VHL-LgBiT plasmid, 1.5 mL OptiMEM I (Gibco 31985070) and 90 μL Fμgene HD (Promega E2312) were incubated for 10 min at r.t. before being added dropwise to the cells. After 24 hr of incubation at 37 °C, 5% CO2, the transfected cells were trypsinized and normalized to 0.50 × 10^6^ cells / mL in DMEM supplemented with 10% FBS. MLN4924 (final 2 μM, 0.1% DMSO) was added to the cells. The cells (5 μL per well) were plated into a 1536-well TC-treated white plate (Greiner 789173-A) using a single tip dispenser (GNF Systems One Tip Dispenser) and incubated for 16 hr at 37 °C, 5% CO2. 15 nL of compound (3-fold serially diluted with DMSO, final top concentration 30 μM, 0.3% DMSO) or DMSO was dispensed into the target wells using an Echo 655 acoustic liquid handler. For NanoBiT assays with VHL ligand competition, 25 nL of 10 mM VH101 (final 50 μM) or the matching % DMSO was dispensed to the wells at the same time as the compound treatment using the Echo 655 acoustic liquid handler. The plates were incubated for 2 hr at 37 °C, 5% CO2, before being removed from the incubator and cooled to r.t. for 15 min. 5 μL of NanoBiT Nano-Glo reagent (Promega N2012, Nano-Glo substrate diluted 1:25 in a mixture of 50% Nano-Glo LCS buffer and 50% PBS) was dispensed into each well using the GNF single tip dispenser. After incubation at r.t. for 15 min, luminescence signals were read on a PHERAstar FS plate reader (BMG Labtech). Luminescence for each compound treated sample was normalized using the equation (well value-NC)/NC × 100%, where NC is the average neutral control (DMSO) well value. EC50 values were calculated from nonlinear regression curve fits (excluding any data points above the hook effect concentration, if any) using GraphPad Prism v9.5.

### HiBiT and CTG assays

15 mL of HEK293 cells were plated at 0.625 × 10^6^ cells / mL in DMEM (Gibco 11995065) supplemented with 10% FBS in a T75 flask. 15 μg HSVtk-driven BRD9 (M1-T597)-HiBiT or BRD7 (M1-S651)-HiBiT plasmid, 15 μg CMV-driven VHL plasmid, 1.5 mL OptiMEM I (Gibco 31985070) and 90 μL Fμgene HD (Promega E2312) were incubated for 10 min at r.t. before being added dropwise to the cells. After 19 hr of incubation at 37 °C, 5% CO2, the transfected cells were trypsinized and normalized to 0.50 × 10^6^ cells / mL in DMEM supplemented with 10% FBS. MLN4924 (final 2 μM, 0.1% DMSO) or DMSO was added to the cells. The cells (5 μL per well) were plated into a 1536-well TC-treated white plate (Greiner 789173-A) using a single tip dispenser (GNF Systems One Tip Dispenser) and incubated for 8 hr at 37 °C, 5% CO2. 15 nL of compound (3-fold serially diluted with DMSO, final top concentration 30 μM, 0.3% DMSO) or DMSO was dispensed into the target wells using an Echo 655 acoustic liquid handler. For HiBiT assays with VHL ligand competition or MLN7243, 25 nL of 10 mM VH101 (final 50 μM), 5 nL of 100 μM MLN7243 (final 0.1 μM), or the matching % DMSO was dispensed to the wells at the same time as the compound treatment using the Echo 655 acoustic liquid handler. The plates were incubated for 15 hr at 37 °C, 5% CO2, before being removed from the incubator and cooled to r.t. for 15 min. 5 μL of HiBiT lytic reagent (Promega N3040, LgBiT protein diluted 1:100 and lytic substrate diluted 1:50 in Nano-Glo HiBiT lytic buffer) was dispensed into each well using the GNF dispenser. For CellTiter-Glo (CTG) assays, instead of the HiBiT lytic reagent, 5 μL of CTG reagent (Promega G7571, CTG substrate dissolved in CTG buffer) was dispensed into each well. After incubation at r.t. for 15 min, luminescence signals were read on a PHERAstar FS plate reader (BMG Labtech). DCmax values were the maximum values after normalizing the luminescence for each compound-treated sample using the equation (well value-NC)/(AC-NC) × 100%, where NC is the average neutral control (DMSO) well value and AC is the average active control (mock-transfected with no DNA) well value. DC50 values were calculated from nonlinear regression curve fits (excluding any data points above the hook effect concentration, if any) using GraphPad Prism v9.5.

### Fluorescence-activated cell sorting

The Artichoke vector was a gift from Benjamin Ebert. Full length BRD9 was cloned into the Artichoke vector using gateway cloning (BRD9 (M1-T597)-eGFP-IRES-mCherry) and lentivirus was produced in HEK293T cells using standard methods. Resulting reporter cells were cultured in DMEM (Gibco 10564011) supplemented with 10% FBS and 1% penicillin/streptomycin at 37 °C, 5% CO2. Prior to the FACS analysis, cells were sorted on a Sony MA900 cell sorter gating for dual eGFP and mCherry positive cells. Sorted cells were allowed to grow to 90% confluency and plated at 0.7 × 10^6^ cells in FluroBrite DMEM (Gibco A1896701) supplemented with 10% FBS per well in a 6-well TC plate. 19 hr after plating the cells, compounds (final 10 μM, 0.5% DMSO) were added to the wells. After 24 hr incubation at 37 °C, 5% CO2, the cells were detached with TrypLE (200 μL) for 1 min and resuspended in chilled 10% FBS in PBS (800 μL). Cells were collected by centrifugation and resuspended in 300 μL of 1% FBS in PBS supplemented with 2 mM EDTA. The cells were strained through 35 μm nylon mesh (Falcon 352235) prior to FACS on a CytoFLEX S Flow cytometer analyzing 100k mCherry-positive cells per sample. A plot of normalized cell count against eGFP level was generated using FCS Express.

### Western blot

Degradation of BRD9-eGFP in HEK293T cells was validated by western blot. The same batch of cells sorted for the FACS experiment was plated at 35k cells in FluroBrite DMEM (Gibco A1896701) supplemented with 10% FBS per well in a 96-well TC plate. 19 hr after plating the cells, compounds (3-fold serially diluted from a top concentration of final 30 μM, 0.5% DMSO) were added to the wells. After 24 hr incubation at 37 °C, 5% CO2, the media was removed and the cells were lysed with 11 μL RIPA buffer with cOmplete™ EDTA-free Protease Inhibitor Cocktail on ice for 30 min. 4 μL of 8 M urea was added, and the supernatant was mixed with NuPAGE™ LDS Sample Buffer (4×) and heated for loading onto an SDS-PAGE gel. Proteins were transferred to PVDF membranes (Invitrogen IB24002) using the iBlot 2 Gel Transfer Device. The membranes were blocked with Intercept (PBS) Blocking Buffer (LI-COR 927-70001) at 4 °C for 1 hr. The blots were probed with an anti-BRD9 rabbit monoclonal antibody (Cell signaling technology 48306, 1:500 in the blocking buffer) and an anti-β actin mouse monoclonal antibody (Santa Cruz Biotechnology sc-47778, 1:500 in the blocking buffer) at 4 °C with gentle shaking overnight. The membranes were washed with TBST for three times, stained with IRDye 800CW goat anti-rabbit IgG (LI-COR 926-32211) and IRDye 680RD goat anti-mouse IgG (LI-COR 926-68070) secondary antibodies (1:10,000 in the blocking buffer), washed with TBST for three times, and imaged using a LI-COR Odyssey infrared imaging system.

## Acknowledgements

We sincerely acknowledge the following individuals for their technical and analytical expertise: Rishi Arora (SPR and microSEC), Andrew Reidenbach (DEL screening), Daniel Fuller (non-denaturing MS), Alicia Lindeman (next-generation sequencing), and Xiyu Jiao (statistical analysis). We are grateful to Dustin Dovala and Michael Romanowski for providing the VCB protein constructs, and Dennis Buckley for the compound VH101. Additionally, we express our appreciation to Raviraj Kulathila, Michael Romanowski, Wenzhi Tian, Liam Hudson, and Patricia Horton for their valuable support and contribution to this project. We are also thankful to Trevor Zandi, Zher Yin Tan, Zhihan Nan and Claire Harmange Magnani for fruitful discussions that enhanced the clarity of this manuscript. Special thanks to Cindy Hon for her excellent operational support throughout this project. S.L. thanks the Agency for Science, Technology and Research (A*STAR, Singapore) for a National Science Scholarship. The research was supported by the National Institute of General Medical Sciences (R35GM127045 awarded to S.L.S.).

## Author contributions

S.L. conceived the project, designed and performed the CIP-DEL screening, AlphaScreen assays, SPR, non-denaturing MS, size exclusion chromatography, and western blot experiments. B.T. synthesized and characterized screening hits off-DNA. J.W.M. designed, synthesized and characterized the CIP-DEL library. J.M.O. and S.L. designed and performed the FACS experiments. A.T. and S.L. designed and performed the NanoBiT, HiBiT and CTG experiments. B.K.H. analyzed the screening data. S.A.T. helped with the off-DNA synthesis. S.L.S., F.J.Z., F.B., K.B., and S.B. provided context for the framing of the original goals, supervision, guidance, operational support, and assisted in the interpretation of experimental outcomes. S.L. prepared the figures. S.L. and S.L.S. wrote the manuscript. All authors read and edited the paper.

## Supplementary Information

Supplementary information is available online at https://doi.org/xxx

