## Supplementary Information for "Rational screening for cooperativity in small-molecule inducers of protein–protein associations"

### DNA-ENCODED LIBRARY SYNTHESIS

#### Synthesis of modified VHL ligand-connector pairs for CIP-DEL library production

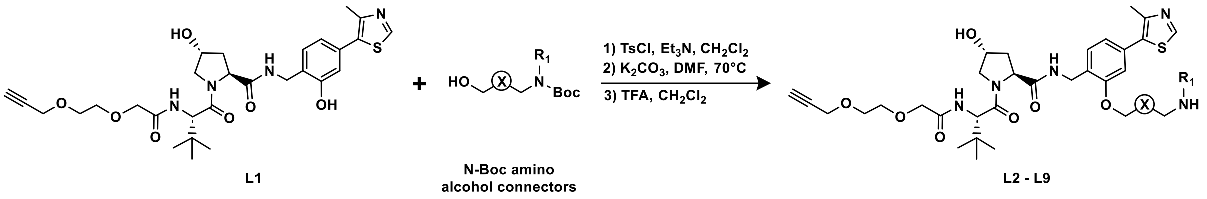

**Scheme L1.** Functionalization of phenolic VHL ligand hydroxyl with N-Boc amino alcohols.

***Synthesis of L2 – L9:*** N-Boc amino alcohols (2.0 equiv) were converted to tosylates with the addition of *p*-toluenesulfonyl chloride (2.0 equiv) and $Et_3N$ (3.0 equiv) in $CH_2Cl_2$ (2 mL). The reactions were stirred at room temperature overnight, then diluted with $CH_2Cl_2$ and concentrated. Crude material was purified by silica gel chromatography (heptanes – EtOAc gradients) and used directly in the next step. Purified tosylates were dissolved in DMF (2 mL) and **L1** (100 mg; 1 equiv) and $K_2CO_3$ (3.0 equiv) were added. The resulting solutions were heated to 70 °C in an oil-bath overnight. Reactions were then cooled to room temperature, diluted with $CH_2Cl_2$ and saturated aqueous $NH_4Cl$, and the organic phases were isolated. The aqueous phases were extracted twice more with $CH_2Cl_2$ and the combined organics were dried with $MgSO_4$, filtered, and concentrated under reduced pressure. Alkylated phenols were purified by silica gel chromatography ($CH_2Cl_2$ – MeOH gradients). Finally, the alkylated phenols were dissolved in $CH_2Cl_2$ (2.0 mL) and TFA (1.0 mL) and stirred at room temperature until UPLC-MS analysis showed complete removal of the Boc protecting groups. Once complete, reactions were diluted with $CH_2Cl_2$, concentrated, and purified by preparative HPLC (water – acetonitrile gradients both modified with 0.1% formic acid).

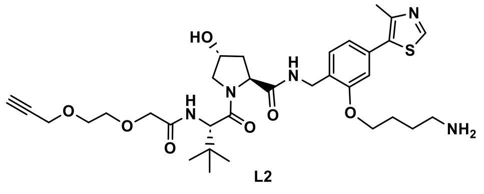

L2

***CIP-DEL presenter ligand with connector 3:***

**$^1$H NMR** (400 MHz, MeOD) δ = 8.89 (s, 1H), 8.50 (t, *J* = 5.8 Hz, 1H), 7.72 (d, *J* = 9.2 Hz, 1H), 7.49 (d, *J* = 7.7 Hz, 1H), 7.08 – 6.97 (m, 2H), 4.72 – 4.65 (m, 1H), 4.61 (dd, *J* = 9.1, 7.5 Hz, 1H), 4.53 – 4.48 (m, 1H), 4.47 – 4.37 (m, 2H), 4.31 – 4.18 (m, 2H), 4.17 – 4.09 (m, 2H), 4.05 (s, 2H), 3.89 (dt, *J* = 11.2, 1.7 Hz, 1H), 3.80 (dd, *J* = 11.0, 3.8 Hz, 1H), 3.77 – 3.69 (m, 4H), 3.12 – 3.00 (m, 2H), 2.85 (t, *J* = 2.4 Hz, 1H), 2.49 (s, 3H), 2.24 (ddt, *J* = 13.1, 7.6, 1.9 Hz, 1H), 2.10 (ddd, *J* = 13.3, 9.1, 4.5 Hz, 1H), 2.00 – 1.86 (m, 4H), 1.03 (s, 9H). **$^{13}$C NMR** (101 MHz, MeOD) δ = 174.4, 174.3, 172.2, 172.2, 171.9, 171.8, 171.6, 157.9, 157.7, 153.0, 152.9, 149.2, 149.1, 133.5, 133.1, 132.9, 129.8, 129.7, 127.8, 127.8, 122.8, 122.7, 113.4, 113.0, 80.5, 80.5, 76.3, 76.2, 72.1, 72.0, 71.0, 71.0, 69.8, 69.0, 68.8, 68.5, 60.8, 60.8, 59.1, 58.4, 58.3, 58.1, 56.4, 41.4, 40.7, 40.6, 39.5, 39.3, 38.9, 37.7, 36.9, 27.2, 27.1, 26.9, 25.9, 25.7, 15.9 (*note:* clear peak doubling present due to rotamers). **HRMS** (ESI+) req. for $C_{33}H_{48}N_5O_7S$ $[M+H]^+$ 658.3269, found 658.3329.

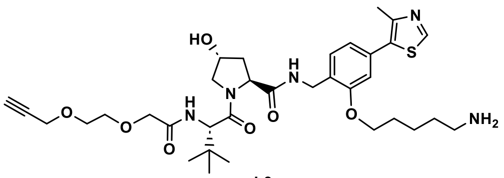

L3

***CIP-DEL presenter ligand with connector 4:***

**$^1$H NMR** (400 MHz, MeOD) δ = 8.87 (s, 1H), 7.49 (d, *J* = 7.7 Hz, 1H), 7.04 – 6.97 (m, 2H), 4.68 (s, 1H), 4.64 – 4.57 (m, 1H), 4.53 – 4.49 (m, 1H), 4.49 – 4.37 (m, 2H), 4.32 – 4.17 (m, 2H), 4.10 (t, *J* = 6.1 Hz, 2H), 4.05 (s, 2H), 3.89 (d, *J* = 11.6 Hz, 1H), 3.80 (dd, *J* = 11.0, 3.8 Hz, 1H), 3.77 – 3.69 (m, 4H), 2.98 (t, *J* = 7.4 Hz, 2H), 2.84 (t, *J* = 2.4 Hz, 1H), 2.48 (s, 3H), 2.24 (ddt, *J* = 13.3, 7.7, 1.9 Hz, 1H), 2.12 (dtd, *J* = 13.2, 8.8, 4.7 Hz, 1H), 1.97 – 1.85 (m, 2H), 1.83 – 1.71 (m, 2H), 1.70 – 1.58 (m, 2H), 1.03 (s, 9H). **$^{13}$C NMR** (101 MHz, MeOD) δ = 174.4, 173.7, 172.2, 172.1, 171.8, 171.4, 158.1, 157.8, 152.9, 152.8, 149.2, 149.1, 133.6, 133.4, 133.2, 132.8, 129.9, 129.6, 127.9, 127.7, 122.5, 113.3, 113.0, 80.5, 76.2, 76.2, 72.2, 72.0, 71.0, 71.0, 69.8, 69.0, 68.8, 61.2, 60.8, 59.1, 58.3, 58.1, 56.3, 41.4, 40.7, 39.5, 39.2, 38.9, 37.6, 37.0, 29.7, 29.7, 28.4, 28.3, 26.9, 24.4, 24.3, 15.9 (*note:* clear peak doubling present due to rotamers). **HRMS** (ESI+) req. for $C_{34}H_{50}N_5O_7S$ $[M+H]^+$ 672.3425, found 672.3471.

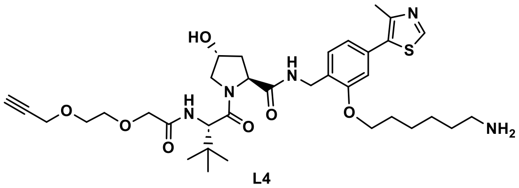

L4

***CIP-DEL presenter ligand with connector 5:***

**$^1$H NMR** (400 MHz, MeOD) δ = 8.87 (s, 1H), 7.49 (d, *J* = 7.7 Hz, 1H), 7.03 – 6.96 (m, 2H), 4.68 (s, 1H), 4.65 – 4.56 (m, 1H), 4.53 – 4.50 (m, 1H), 4.50 – 4.35 (m, 2H), 4.32 – 4.17 (m, 2H), 4.08 (t, *J* = 6.2 Hz, 2H), 4.05 (d, *J* = 1.0 Hz, 2H), 3.88 (dt, *J* = 11.3, 1.7 Hz, 1H), 3.80 (dd, *J* = 11.0, 3.8 Hz, 1H), 3.77 – 3.70 (m, 4H), 2.93 (t, *J* = 7.5 Hz, 2H), 2.84 (t, *J* = 2.4 Hz, 1H), 2.48 (s, 3H), 2.24 (ddt, *J* = 13.2, 7.8, 1.9 Hz, 1H), 2.12 (ddd, *J* = 13.2, 9.0, 4.5 Hz, 1H), 1.92 – 1.83 (m, 2H), 1.70 (p, *J* = 7.6 Hz, 2H), 1.65 – 1.55 (m, 2H), 1.55 – 1.44 (m, 2H), 1.03 (s, 9H). **$^{13}$C NMR** (101 MHz, MeOD) δ = 174.4, 173.6, 172.2, 172.1, 171.7, 171.3, 170.3, 158.2, 157.9, 152.9, 152.8, 149.2, 149.0, 133.6, 133.5, 133.2, 132.7, 130.2, 129.5, 127.9, 127.6, 122.4, 119.7, 116.8, 113.3, 113.0, 80.5, 76.2, 76.2, 72.2, 72.1, 71.0, 71.0, 69.8, 69.2, 69.1, 61.1, 60.7, 59.1, 58.3, 58.0, 56.3, 41.4, 40.8, 39.5, 39.2, 38.9, 37.6, 37.0, 30.1, 30.0, 29.0, 28.8, 27.2, 27.1, 26.9, 26.8, 26.8, 15.9 (*note:* clear peak doubling present due to rotamers). **HRMS** (ESI+) req. for $C_{35}H_{52}N_5O_7S$ $[M+H]^+$ 686.3582, found 686.3621.

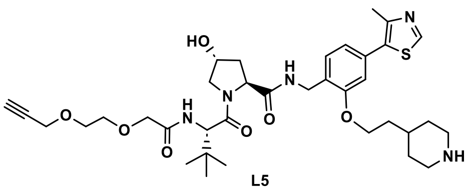

L5

***CIP-DEL presenter ligand with connector 9:***

**$^1$H NMR** (400 MHz, MeOD) δ = 8.88 (s, 1H), 7.49 (d, *J* = 8.1 Hz, 1H), 7.04 – 6.99 (m, 2H), 4.68 (s, 1H), 4.64 – 4.56 (m, 1H), 4.53 – 4.49 (m, 1H), 4.49 – 4.36 (m, 2H), 4.30 – 4.18 (m, 2H), 4.15 (t, *J* = 6.0 Hz, 2H), 4.05 (s, 2H), 3.88 (dt, *J* = 11.3, 1.7 Hz, 1H), 3.79 (dd, *J* = 11.0, 3.9 Hz, 1H), 3.77 – 3.70 (m, 4H), 3.37 (dt, *J* = 13.2, 3.3 Hz, 2H), 2.99 (tt, *J* = 12.9, 2.5 Hz, 2H), 2.85 (t, *J* = 2.4 Hz, 1H), 2.49 (s, 3H), 2.23 (ddt, *J* = 13.2, 7.6, 1.9 Hz, 1H), 2.12 (ddd, *J* = 13.2, 9.0, 4.5 Hz, 1H), 2.07 – 1.93 (m, 3H), 1.86 (q, *J* = 6.2 Hz, 2H), 1.47 (q, *J* = 11.1 Hz, 2H), 1.02 (s, 9H). **$^{13}$C NMR** (101 MHz, MeOD) δ = 174.3, 172.2, 171.8, 157.8, 152.8, 149.1, 133.5, 132.8, 129.7, 127.9, 122.6, 112.9, 80.5, 76.2, 72.2, 71.0, 71.0, 69.8, 66.5, 60.7, 59.1, 58.2, 58.0, 45.4, 39.3, 38.9, 37.0, 36.3, 32.4, 30.3, 30.2, 26.9, 15.9. **HRMS** (ESI+) req. for $C_{36}H_{52}N_5O_7S$ $[M+H]^+$ 698.3582, found 698.3651.

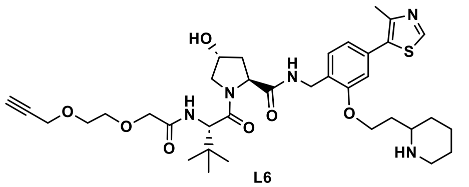

L6

***CIP-DEL presenter ligand with connector 10:***

**$^{1}$H NMR** (400 MHz, MeOD) δ = 8.88 (s, 1H), 8.55 (s, 1H), 7.48 (dd, *J* = 8.0, 1.8 Hz, 1H), 7.04 (d, *J* = 7.4 Hz, 2H), 4.67 (d, *J* = 1.2 Hz, 1H), 4.62 – 4.54 (m, 1H), 4.49 (d, *J* = 9.5 Hz, 1H), 4.45 – 4.36 (m, 2H), 4.30 – 4.17 (m, 4H), 4.04 (s, 2H), 3.88 (d, *J* = 11.1 Hz, 1H), 3.79 (dt, *J* = 10.7, 3.4 Hz, 1H), 3.77 – 3.69 (m, 4H), 3.22 – 3.12 (m, 1H), 2.95 – 2.82 (m, 2H), 2.49 (s, 3H), 2.27 – 2.18 (m, 1H), 2.09 (ddd, *J* = 13.4, 9.2, 4.6 Hz, 3H), 1.98 (d, *J* = 13.7 Hz, 1H), 1.88 (d, *J* = 11.1 Hz, 1H), 1.79 (d, *J* = 11.3 Hz, 1H), 1.58 (q, *J* = 12.2 Hz, 2H), 1.44 (t, *J* = 12.1 Hz, 1H), 1.01 (d, *J* = 1.9 Hz, 9H). **HRMS** (ESI+) req. for $C_{36}H_{52}N_5O_7S$ $[M+H]^+$ 698.3582, found 698.3633.

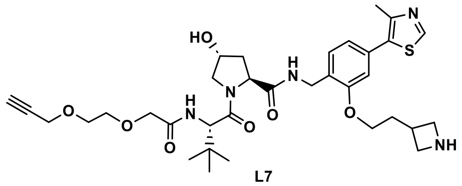

L7

***CIP-DEL presenter ligand with connector 11:***

**$^{1}$H NMR** (400 MHz, MeOD) δ = 8.88 (s, 1H), 7.71 (d, *J* = 9.3 Hz, 1H), 7.49 (d, *J* = 7.7 Hz, 1H), 7.04 (dd, *J* = 7.7, 1.6 Hz, 1H), 7.01 (d, *J* = 1.6 Hz, 1H), 4.72 – 4.67 (m, 1H), 4.61 (dd, *J* = 9.2, 7.6 Hz, 1H), 4.53 – 4.48 (m, 1H), 4.48 – 4.31 (m, 2H), 4.31 – 4.16 (m, 5H), 4.15 – 4.09 (m, 2H), 4.05 (s, 2H), 4.04 – 3.96 (m, 2H), 3.89 (dt, *J* = 11.2, 1.7 Hz, 1H), 3.80 (dd, *J* = 11.0, 3.8 Hz, 1H), 3.77 – 3.71 (m, 4H), 3.25 (q, *J* = 7.9 Hz, 1H), 2.86 (t, *J* = 2.4 Hz, 1H), 2.48 (s, 3H), 2.28 – 2.18 (m, 3H), 2.10 (ddd, *J* = 13.3, 9.1, 4.4 Hz, 1H), 1.03 (s, 9H). **$^{13}$C NMR** (101 MHz, MeOD) δ = 174.4, 174.4, 172.2, 172.2, 171.9, 171.8, 157.7, 152.9, 149.2, 133.4, 133.0, 129.9, 127.5, 122.9, 113.0, 80.5, 76.3, 76.2, 72.1, 72.0, 71.0, 71.0, 69.8, 69.1, 67.6, 60.8, 59.1, 58.4, 58.3, 58.1, 53.3, 53.1, 39.6, 39.5, 38.9, 37.7, 36.9, 33.7, 32.2, 26.9, 15.9, 15.9 (*note:* clear peak doubling present due to rotamers). **HRMS** (ESI+) req. for $C_{34}H_{48}N_5O_7S$ $[M+H]^+$ 670.3269, found 670.3328.

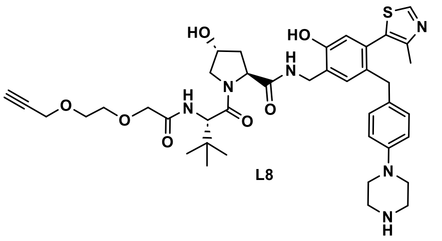

L8

***CIP-DEL presenter ligand with connector 12:***

****Note:*** for **L8**, alkylation occurred at the para-position of phenolic phenyl ring (see **Supplementary Figure S4** for additional details). Also, after preparative HPLC purification of the **L8** presenter ligand, UPLC-MS analysis indicated sample purity of ~80%. Since additional purification would be performed after conjugation to the DNA headpiece, the resulting material was progressed for library production.

**$^{1}$H NMR** (400 MHz, MeOD) δ = 8.86 (s, 1H), 8.53 (s, 1H), 7.30 (s, 1H), 7.03 – 6.96 (m, 1H), 6.78 (dd, *J* = 14.8, 9.0 Hz, 4H), 6.64 (s, 1H), 4.66 (s, 1H), 4.57 (t, *J* = 8.3 Hz, 1H), 4.51 – 4.46 (m, 1H), 4.41 (s, 1H), 4.14 (dd, *J* = 4.7, 2.4 Hz, 1H), 4.02 (d, *J* = 3.3 Hz, 2H), 3.86 (d, *J* = 11.2 Hz, 1H), 3.79 (dd, *J* = 11.1, 4.0 Hz, 1H), 3.75 – 3.71 (m, 2H), 3.71 – 3.62 (m, 4H), 3.27 (s, 6H), 2.83 (t, *J* = 2.4 Hz, 1H), 2.48 (s, 1H), 2.37 (s, 1H), 2.22 – 2.13 (m, 1H), 2.05 (dt, *J* = 8.7, 4.4 Hz, 1H), 1.95 (s, 2H), 1.02 (s, 9H). **$^{13}$C NMR** (101 MHz, MeOD) δ = 174.5, 172.1, 171.7, 164.0, 155.3, 154.7, 153.4, 150.5, 150.0, 135.5, 133.5, 132.1, 131.3, 130.4, 127.0, 119.1, 118.0, 80.5, 76.2, 72.1, 71.0, 71.0, 69.8, 60.7, 59.1, 59.1, 58.2, 58.0, 45.1, 39.4, 39.2, 38.9, 37.0, 27.0, 27.0, 15.0. **HRMS** (ESI+) req. for $C_{40}H_{53}N_6O_7S$ $[M+H]^+$ 761.3691, found 761.3742.

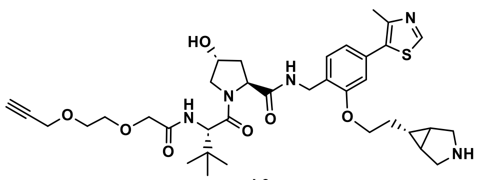

L9

***CIP-DEL presenter ligand with connector 13:***

**$^1$H NMR** (400 MHz, MeOD) δ = 8.90 (s, 1H), 7.50 (d, *J* = 7.7 Hz, 1H), 7.02 (dt, *J* = 10.7, 2.0 Hz, 2H), 4.68 (s, 1H), 4.61 (dd, *J* = 9.0, 7.6 Hz, 1H), 4.54 – 4.50 (m, 1H), 4.50 – 4.39 (m, 2H), 4.28 (dd, *J* = 16.0, 2.4 Hz, 1H), 4.21 (dd, *J* = 16.1, 2.4 Hz, 1H), 4.14 (t, *J* = 5.8 Hz, 2H), 4.05 (s, 2H), 3.89 (d, *J* = 10.7 Hz, 1H), 3.80 (dd, *J* = 11.0, 3.8 Hz, 1H), 3.77 – 3.70 (m, 4H), 3.49 – 3.37 (m, 4H), 2.85 (t, *J* = 2.4 Hz, 1H), 2.49 (s, 3H), 2.24 (ddt, *J* = 11.6, 7.9, 1.9 Hz, 1H), 2.11 (ddd, *J* = 13.3, 9.0, 4.4 Hz, 1H), 1.84 (dq, *J* = 9.7, 5.7 Hz, 2H), 1.77 (t, *J* = 3.2 Hz, 2H), 1.12 (tt, *J* = 7.1, 3.7 Hz, 1H), 1.03 (s, 9H). **$^{13}$C NMR** (101 MHz, MeOD) δ = 174.4, 172.2, 171.8, 157.8, 152.9, 149.0, 133.6, 132.8, 129.7, 127.9, 122.6, 113.0, 80.5, 76.3, 72.1, 71.0, 70.9, 69.8, 68.5, 60.8, 59.1, 58.3, 58.1, 39.3, 38.9, 36.9, 32.0, 26.9, 23.4, 23.2, 19.2, 15.9. **HRMS** (ESI+) req. for $C_{36}H_{50}N_5O_7S$ $[M+H]^+$ 696.3425, found 696.3474.

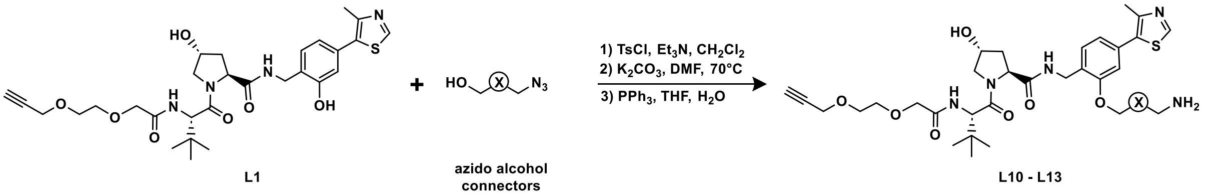

L1 azido alcohol connectors L10 - L13

**Scheme L2.** Functionalization of phenolic VHL ligand hydroxyl with azido amino alcohols.

***Synthesis of L10 – L13:*** Azido amino alcohols (2.0 equiv) were combined with *p*-toluenesulfonyl chloride (2.0 equiv) and $Et_3N$ (3.0 equiv) in $CH_2Cl_2$ (2 mL) and the reactions were stirred at room temperature overnight. The next day, reactions were diluted with $CH_2Cl_2$, concentrated, and purified by silica gel chromatography (heptanes – EtOAc gradients). Purified tosylates were dissolved in DMF (2 mL) and **L1** (125 mg; 1 equiv) and $K_2CO_3$ (3.0 equiv) were added. The reaction mixtures were heated to 70 °C in an oil-bath overnight, then cooled to room temperature and diluted with $CH_2Cl_2$ and saturated aqueous $NH_4Cl$. The organic phases were isolated and the aqueous phases were extracted twice more with $CH_2Cl_2$. The combined organics were dried with $MgSO_4$, filtered, and concentrated under reduced pressure. Alkylated phenols were purified by silica gel chromatography ($CH_2Cl_2$ – MeOH gradients). Lastly, the purified azido products were reduced with the addition of triphenylphosphine (3.0 equiv) in THF (3 mL) and water (1 mL) at room temperature overnight. The next day, methanol was added to the reactions which induced formation of a white precipitate. The precipitate was removed using a 0.45 µM filter and the filtrate was concentrated under reduced pressure. Upon concentration, additional precipitate formed and the slurries were loaded on to small silica gel columns and the desired products were isolated as clear colorless oils. Each product was further purified by preparative HPLC (water – acetonitrile gradients both modified with 0.1% formic acid) to obtain products ready for DNA conjugation.

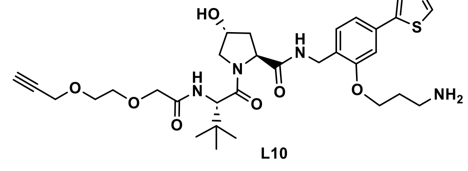

L10

***CIP-DEL presenter ligand with connector 1:***

[1]**H NMR** (400 MHz, MeOD) δ = 8.88 (s, 1H), 8.54 (s, 1H), 7.49 (d, *J* = 7.6 Hz, 1H), 7.11 – 6.96 (m, 2H), 4.68 (s, 1H), 4.62 – 4.54 (m, 1H), 4.50 (s, 1H), 4.43 (d, *J* = 5.8 Hz, 2H), 4.33 – 4.14 (m, 4H), 4.04 (s, 2H), 3.89 (d, *J* = 11.2 Hz, 1H), 3.80 (dd, *J* = 11.1, 4.0 Hz, 1H), 3.73 (d, *J* = 2.6 Hz, 4H), 3.19 (t, *J* = 7.1 Hz, 2H), 2.87 (t, *J* = 2.6 Hz, 1H), 2.49 (s, 3H), 2.30 – 2.14 (m, 3H), 2.09 (ddd, *J* = 13.6, 9.4, 4.5 Hz, 1H), 1.02 (s, 9H). [13]**C NMR** (101 MHz, MeOD) δ = 174.3, 172.2, 171.8, 170.3, 157.6, 152.9, 149.2, 133.4, 133.1, 130.2, 127.8, 123.0, 113.2, 80.5, 76.3, 72.1, 71.0, 71.0, 69.8, 66.6, 60.8, 59.1, 58.3, 58.1, 39.4, 38.9, 38.8, 36.9, 28.7, 26.9, 15.9. **HRMS** (ESI+) req. for $C_{32}H_{46}N_5O_7S$ $[M+H]^+$ 644.3112, found 644.3255.

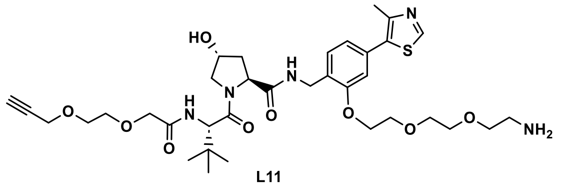

L11

***CIP-DEL presenter ligand with connector 2:***

[1]**H NMR** (400 MHz, MeOD) δ = 8.88 (s, 1H), 8.54 (s, 1H), 7.48 (d, *J* = 7.6 Hz, 1H), 7.10 – 7.00 (m, 2H), 4.67 (s, 1H), 4.63 – 4.54 (m, 2H), 4.53 – 4.41 (m, 3H), 4.31 – 4.17 (m, 4H), 4.05 (s, 2H), 3.97 – 3.86 (m, 3H), 3.83 – 3.62 (m, 10H), 3.10 (s, 2H), 2.85 (t, *J* = 2.7 Hz, 1H), 2.49 (s, 3H), 2.30 – 2.15 (m, 2H), 2.13 – 1.98 (m, 2H), 1.29 (s, 2H), 1.04 (s, 9H). **HRMS** (ESI+) req. for $C_{35}H_{52}N_5O_9S$ $[M+H]^+$ 718.3480, found 718.3570.

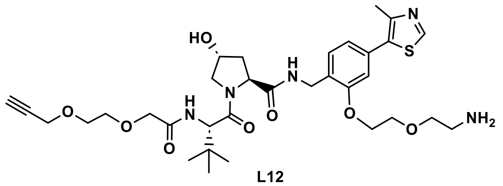

L12

***CIP-DEL presenter ligand with connector 6:***

[1]**H NMR** (400 MHz, MeOD) δ = 8.88 (s, 1H), 8.54 (s, 1H), 7.50 (d, *J* = 8.1 Hz, 1H), 7.09 – 7.00 (m, 2H), 4.68 (s, 1H), 4.63 – 4.56 (m, 1H), 4.53 – 4.49 (m, 1H), 4.49 – 4.38 (m, 2H), 4.31 – 4.18 (m, 4H), 4.04 (s, 2H), 3.95 (dd, *J* = 5.4, 3.2 Hz, 2H), 3.89 (dt, *J* = 11.1, 1.7 Hz, 1H), 3.83 – 3.76 (m, 3H), 3.77 – 3.70 (m, 4H), 3.12 (t, *J* = 5.1 Hz, 2H), 2.85 (t, *J* = 2.4 Hz, 1H), 2.49 (s, 3H), 2.23 (ddt, *J* = 11.6, 7.7, 1.9 Hz, 1H), 2.11 (ddd, *J* = 13.2, 9.1, 4.4 Hz, 1H), 1.02 (s, 9H). [13]**C NMR** (101 MHz, MeOD) δ = 174.3, 172.2, 171.8, 170.3, 157.8, 152.9, 149.1, 133.4, 132.9, 129.9, 128.2, 122.9, 113.4, 80.5, 76.3, 72.2, 71.0, 71.0, 70.8, 69.8, 69.2, 69.0, 60.8, 59.1, 58.3, 58.1, 40.9, 39.3, 38.9, 36.9, 26.9, 15.9. **HRMS** (ESI+) req. for $C_{33}H_{48}N_5O_8S$ $[M+H]^+$ 674.3218, found 674.3261.

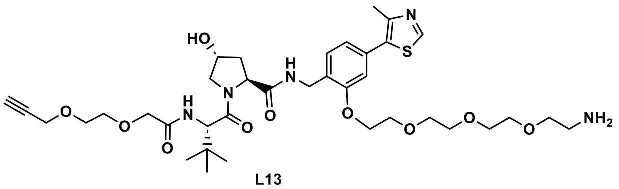

L13

***CIP-DEL presenter ligand with connector 7:***

[1]**H NMR** (400 MHz, MeOD) δ = 8.88 (s, 1H), 8.55 (s, 1H), 7.48 (d, *J* = 7.7 Hz, 1H), 7.08 – 7.00 (m, 2H), 4.68 (s, 1H), 4.60 (t, *J* = 8.3 Hz, 1H), 4.52 – 4.49 (m, 1H), 4.49 – 4.37 (m, 2H), 4.32 – 4.17 (m, 4H), 4.04 (s, 2H), 3.95 – 3.84 (m, 3H), 3.83 – 3.60 (m, 15H), 3.01 (t, *J* = 5.1 Hz, 2H), 2.85 (t, *J* = 2.4 Hz, 1H), 2.49 (s, 3H), 2.23 (ddt, *J* = 11.7, 7.7, 2.0 Hz, 1H), 2.09 (ddt, *J* = 13.3, 9.1, 4.9 Hz, 1H), 1.03 (s, 9H). [13]**C NMR** (101 MHz, MeOD) δ = 174.4, 172.2, 171.7, 170.3, 158.0, 152.9, 149.2, 133.5, 132.9, 129.9, 128.3, 122.9, 113.7, 80.5, 76.2, 72.2, 71.7, 71.5, 71.5, 71.2, 71.0, 71.0, 70.8, 69.8, 69.3, 69.3, 60.8, 59.1, 58.3, 58.1, 41.0, 39.4, 38.9, 37.0, 27.0, 15.9. **HRMS** (ESI+) req. for $C_{37}H_{56}N_5O_{10}S$ $[M+H]^+$ 762.3742, found 762.3821.

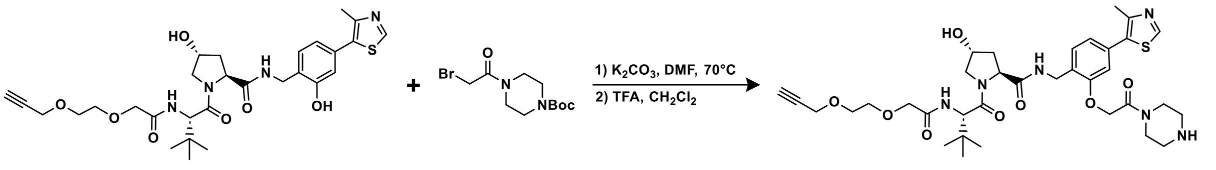

**Scheme L3.** Functionalization of phenolic VHL ligand hydroxyl with N-Boc amino alpha-bromo ketones.

***Synthesis of L15:*** alpha-bromo ketone **L14** (1.1 equiv) was combined with **L1** (125 mg; 1.0 equiv) and $K_2CO_3$ (3.0 equiv) in DMF (2 mL) and the reaction mixture was heated to 70 °C in an oil-bath. After 3 hr, UPLC-MS analysis showed complete consumption of starting material and the reaction was cooled to room temperature, diluted with $CH_2Cl_2$ and saturated aqueous $NH_4Cl$, and the organic phase was isolated. The aqueous phase was further extracted with $CH_2Cl_2$ (2x) and the combined organics were dried with $MgSO_4$, filtered, and concentrated under reduced pressure. The crude product was purified by silica gel chromatography ($CH_2Cl_2$ – MeOH gradients). Finally, the N-Boc protecting group was removed by dissolving the above product in $CH_2Cl_2$ (2 mL) followed by TFA (1 mL) and stirring at room temperature. After 3.5 hr, the reaction mixture was diluted with $CH_2Cl_2$ and concentrated. The crude product was purified by preparative HPLC (water – acetonitrile gradients both modified with 0.1% formic acid) to obtain pure product **L15**.

***<u>CIP-DEL presenter ligand with connector 8:</u>***
**$^1$H NMR** (400 MHz, MeOD) δ = 8.88 (s, 1H), 7.49 (d, *J* = 8.1 Hz, 1H), 7.10 – 7.01 (m, 2H), 5.05 (d, *J* = 14.6 Hz, 1H), 5.00 (d, *J* = 14.6 Hz, 1H), 4.66 (s, 1H), 4.58 (dd, *J* = 9.5, 7.2 Hz, 1H), 4.54 – 4.37 (m, 3H), 4.25 (dd, *J* = 16.1, 2.4 Hz, 1H), 4.19 (dd, *J* = 16.0, 2.4 Hz, 1H), 4.04 (s, 2H), 3.91 – 3.76 (m, 6H), 3.76 – 3.68 (m, 4H), 3.29 – 3.11 (m, 4H), 2.86 (t, *J* = 2.4 Hz, 1H), 2.47 (s, 3H), 2.22 (ddt, *J* = 13.1, 7.7, 2.0 Hz, 1H), 2.07 (ddt, *J* = 13.4, 9.1, 5.1 Hz, 1H), 1.01 (s, 9H). **$^{13}$C NMR** (101 MHz, MeOD) δ = 174.2, 172.1, 171.6, 169.0, 169.0, 157.4, 153.0, 149.3, 133.2, 133.1, 130.5, 128.4, 123.6, 114.0, 80.5, 76.3, 72.1, 71.0, 70.9, 69.8, 67.8, 60.8, 59.1, 58.3, 58.1, 44.8, 44.6, 43.5, 40.3, 39.7, 38.9, 37.7, 36.9, 26.9, 16.0. **HRMS** (ESI+) req. for $C_{35}H_{49}N_6O_8S$ $[M+H]^+$ 713.3327, found 713.3387.

#### CIP-DEL library production

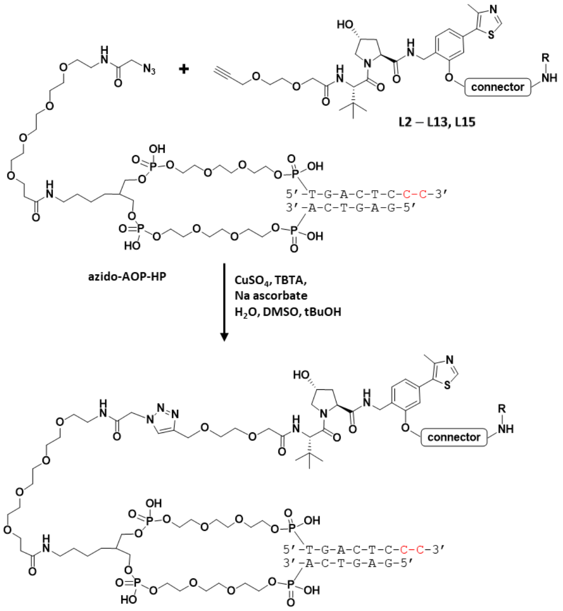

**Scheme L4.** *Library production step 1:* coupling of presenter ligands with connectors to the azido-AOP DNA headpiece.

**Synthesis of VHL-connector conjugates:** the VHL presenter ligands, conjugated to the 13 connectors included in the library, were appended on to the azido-AOP headpiece as described previously[1]. Following CuAAC coupling and EtOH precipitation, each DNA conjugate was purified by preparative HPLC and product fractions lyophilized. The purified conjugates were dissolved in 50 µL of 100 mM phosphate buffer pH 8.0 and quantified by NanoDrop. For the subsequent ligation step, each conjugate was normalized to 32 nmol of purified material in 37 µL of buffer.

**Conjugate of L2:** MS (ESI-) expected 5922 Da, found 5925 Da.

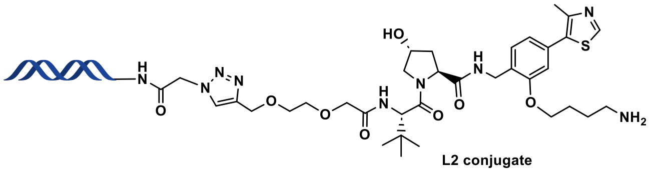

***UV:***

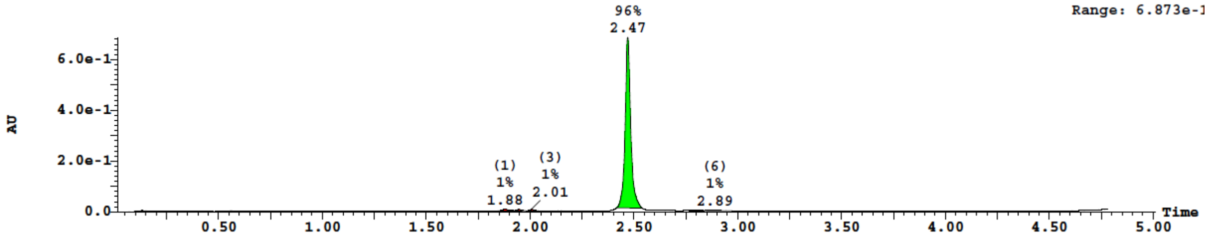

***TIC:***

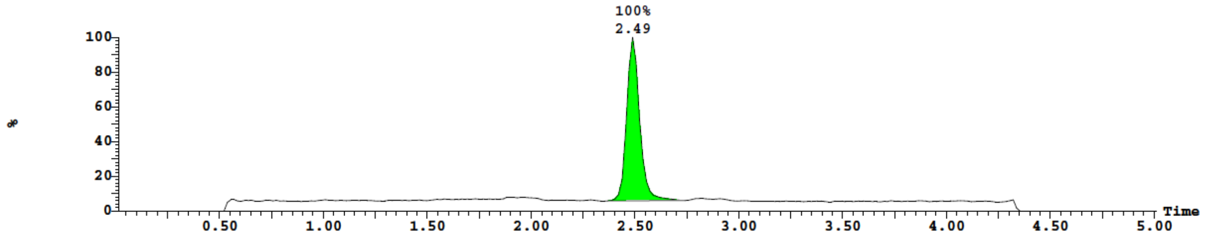

**Conjugate of L3:** MS (ESI-) expected 5936 Da, found 5940 Da.

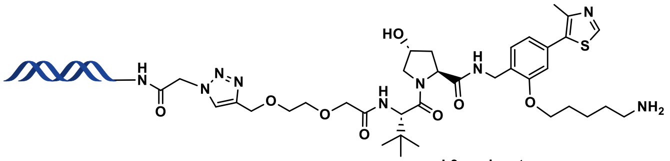

L3 conjugate

*UV:*

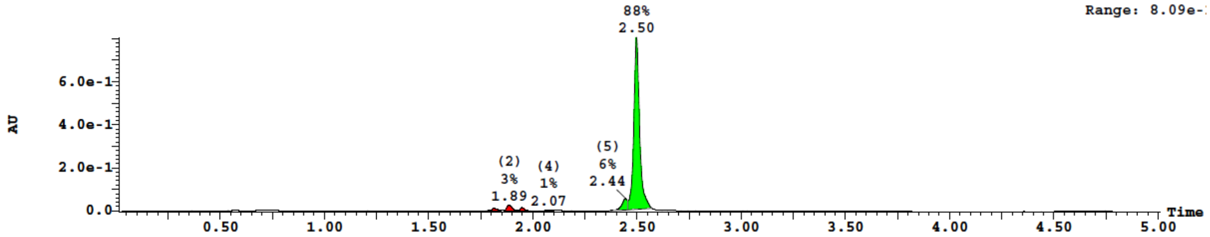

*TIC:*

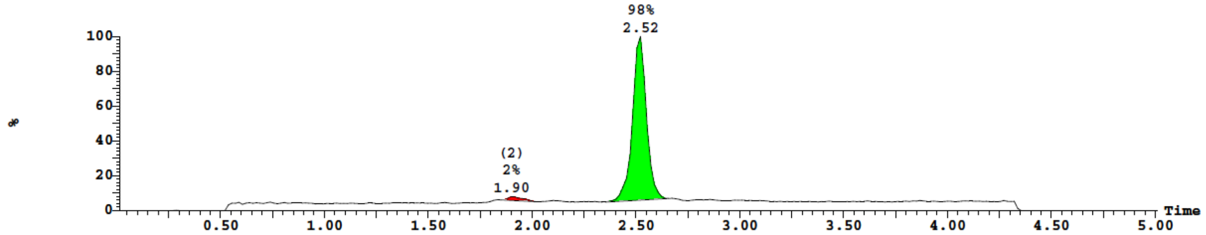

**Conjugate of L4:** MS (ESI-) expected 5950 Da, found 5953 Da.

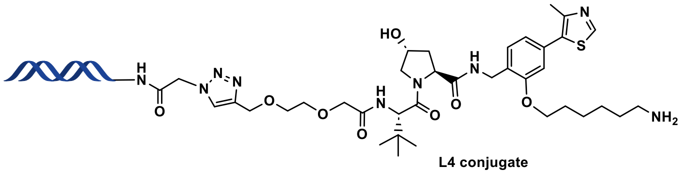

L4 conjugate

*UV:*

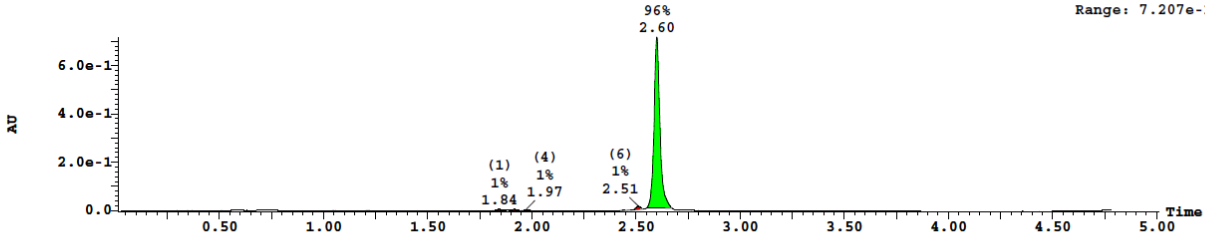

*TIC:*

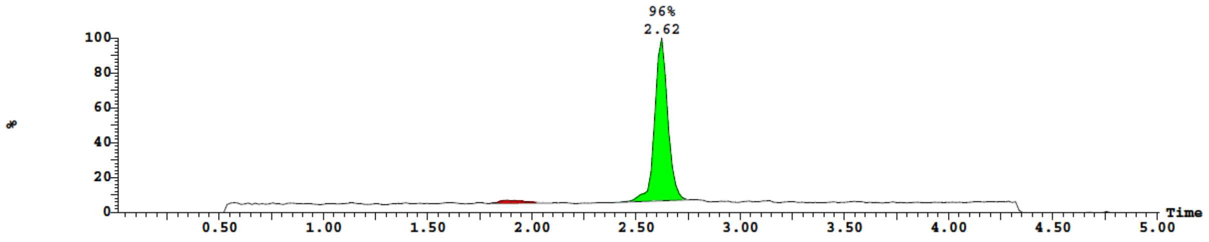

**Conjugate of L5:** MS (ESI-) expected 5962 Da, found 5965 Da.

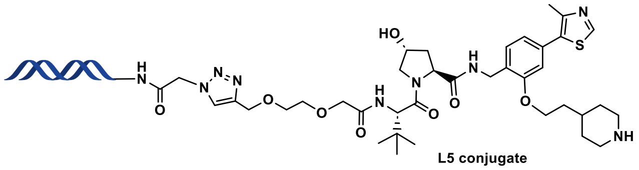

L5 conjugate

*UV:*

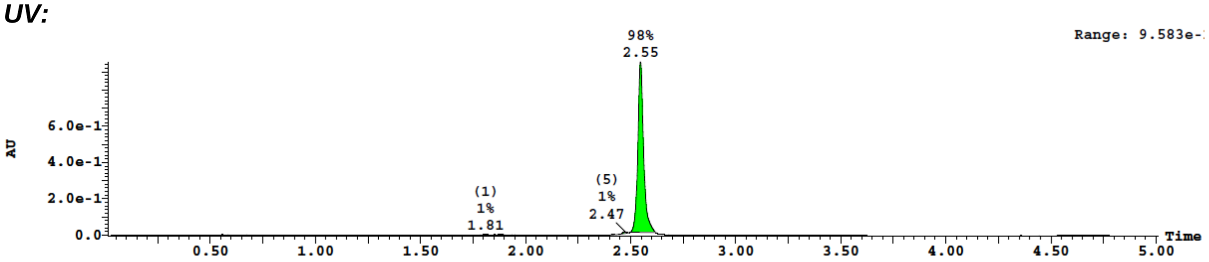

*TIC:*

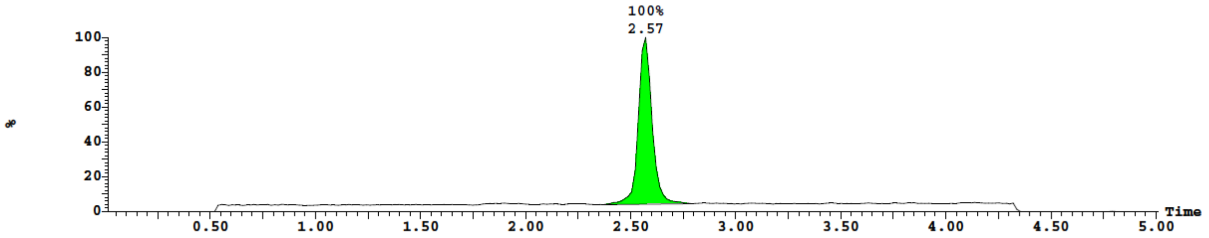

**Conjugate of L6:** MS (ESI-) expected 5962 Da, found 5965 Da.

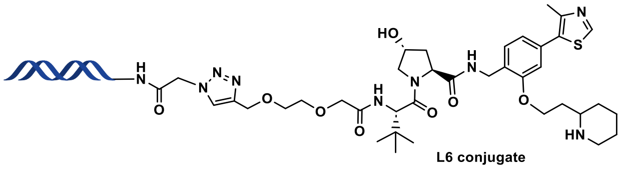

L6 conjugate

*UV:*

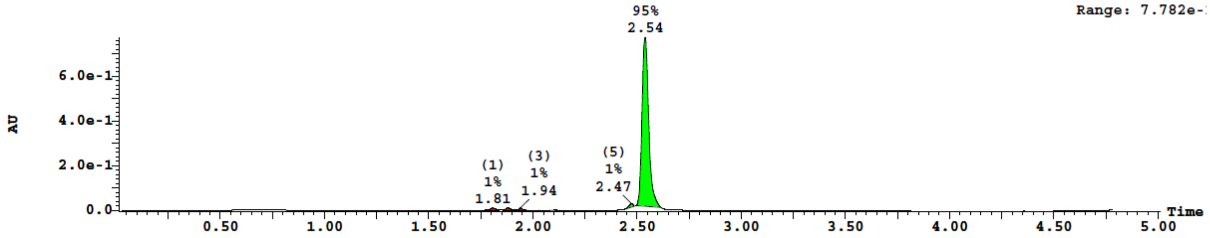

*TIC:*

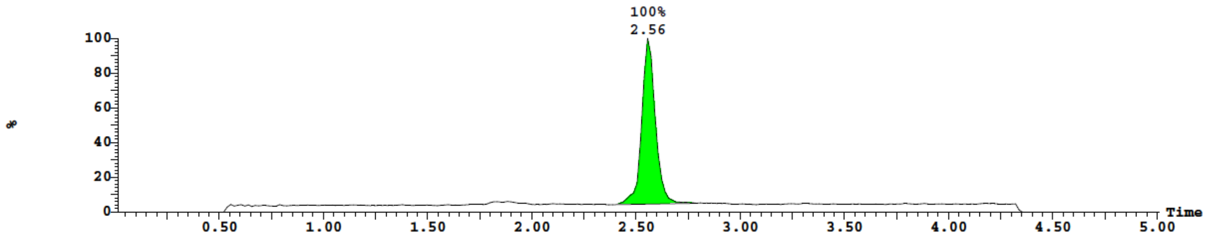

**Conjugate of L7:** MS (ESI-) expected 5933 Da, found 5937 Da.

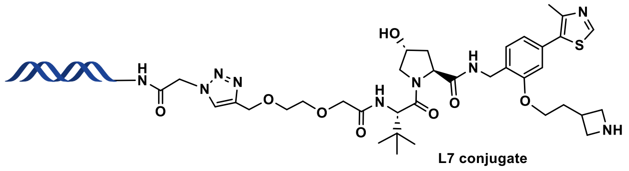

***UV:***

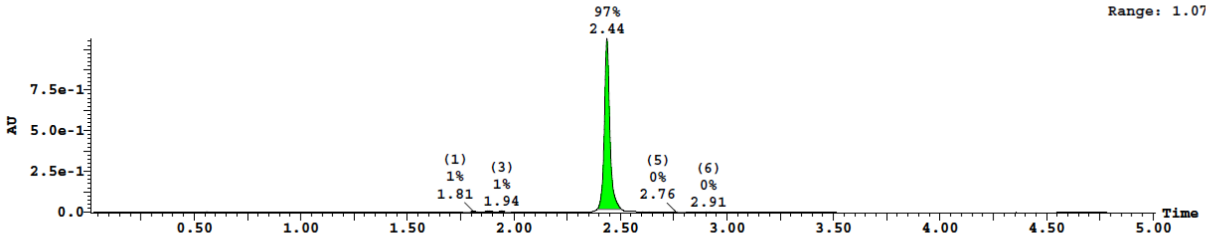

***TIC:***

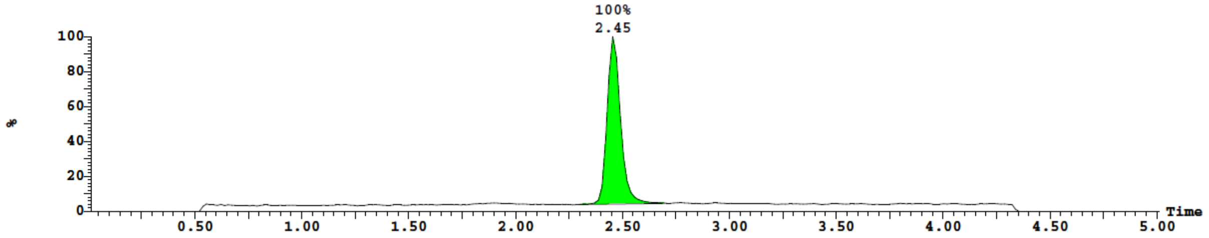

**Conjugate of L8:** MS (ESI-) expected 6025 Da, found 6029 Da.

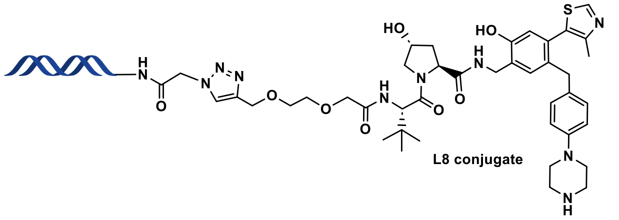

***UV:***

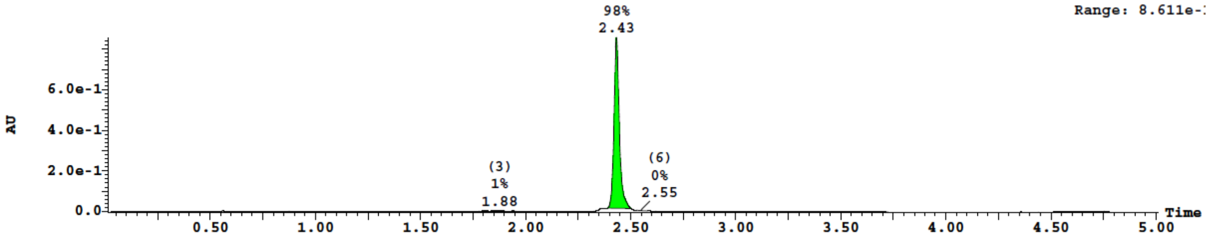

***TIC:***

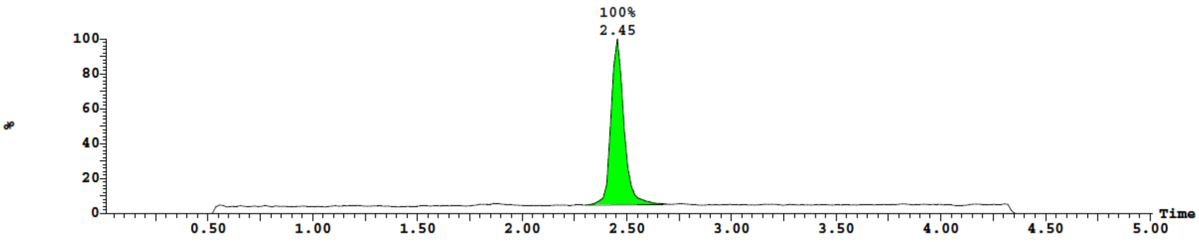

**Conjugate of L9:** MS (ESI-) expected 5960 Da, found 5963 Da.

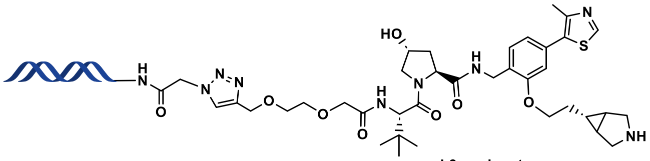

L9 conjugate

*UV:*

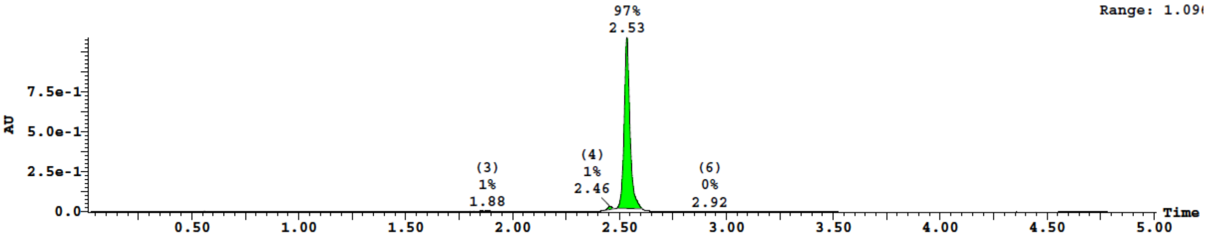

*TIC:*

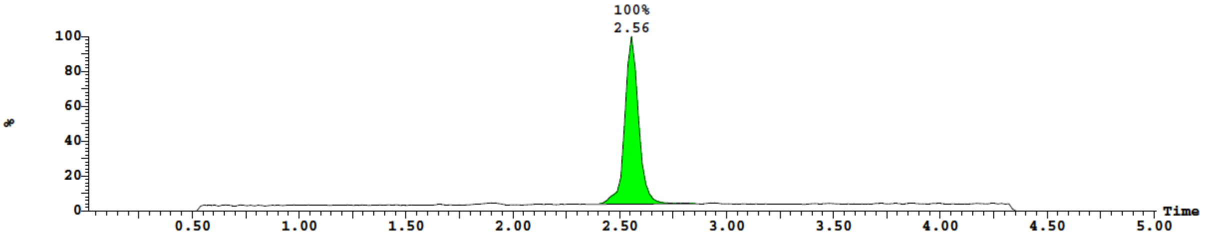

**Conjugate of L10:** MS (ESI-) expected 5908 Da, found 5912 Da.

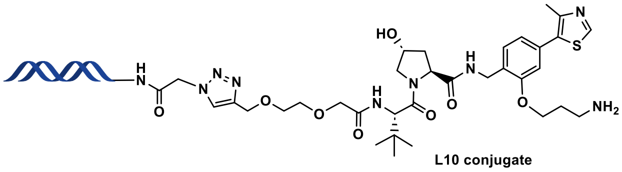

L10 conjugate

*UV:*

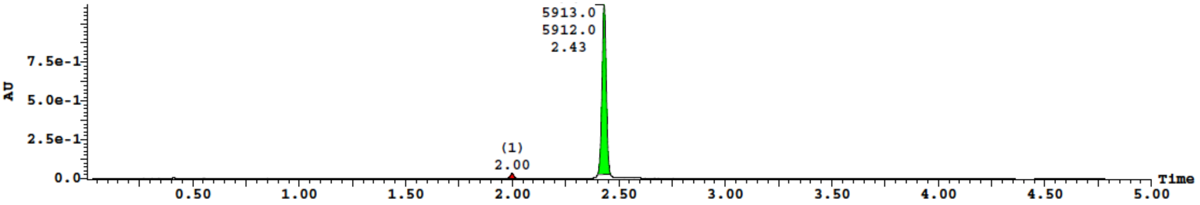

*TIC:*

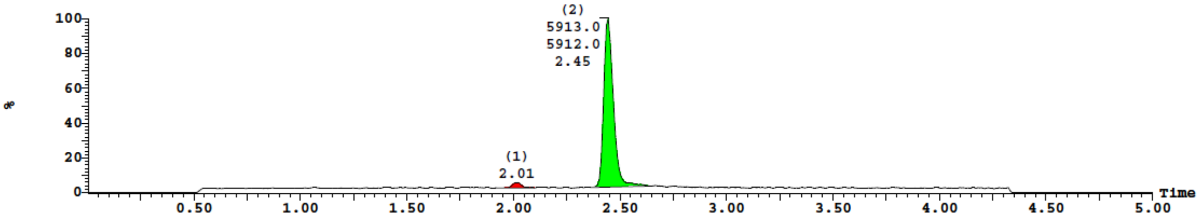

**Conjugate of L11:** MS (ESI-) expected 5982 Da, found 5985 Da.

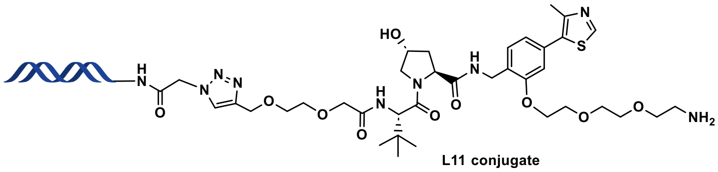

L11 conjugate

***UV:***

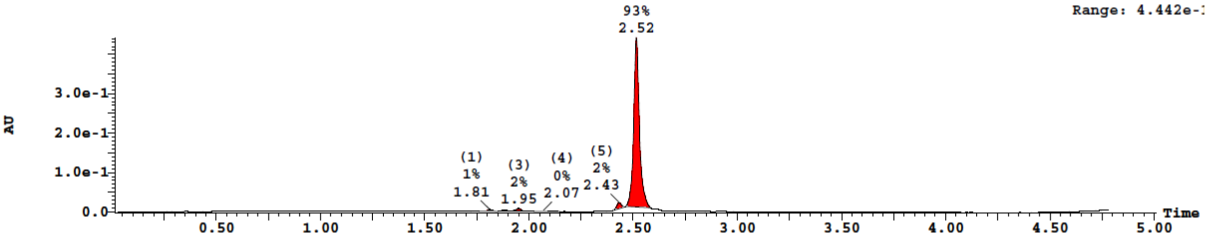

***TIC:***

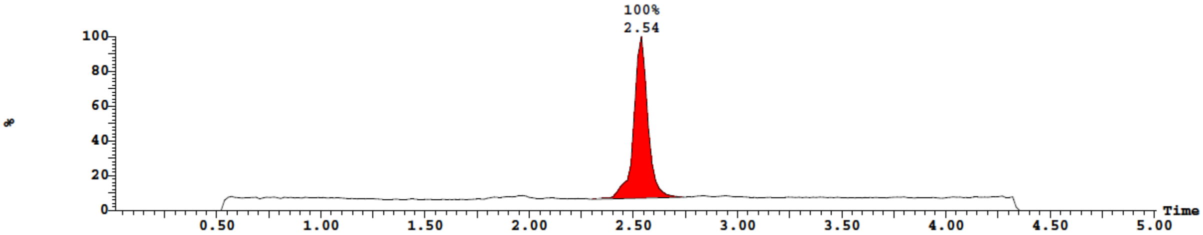

**Conjugate of L12:** MS (ESI-) expected 5938 Da, found 5942 Da.

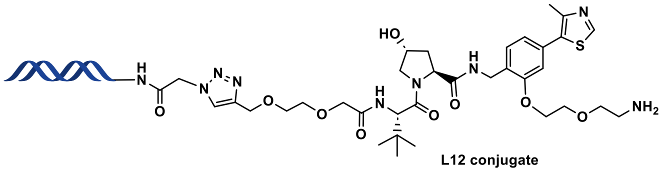

L12 conjugate

***UV:***

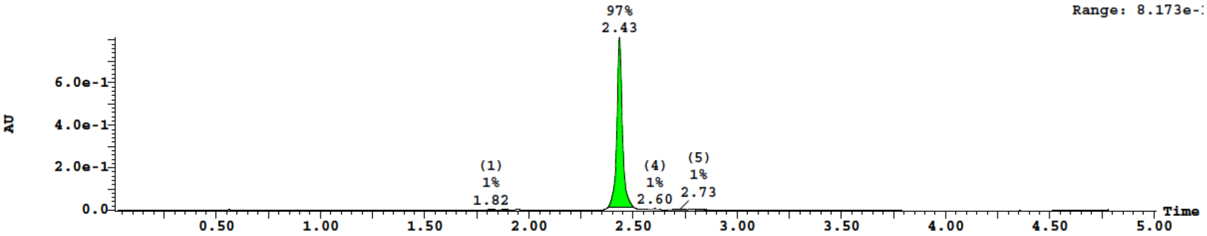

***TIC:***

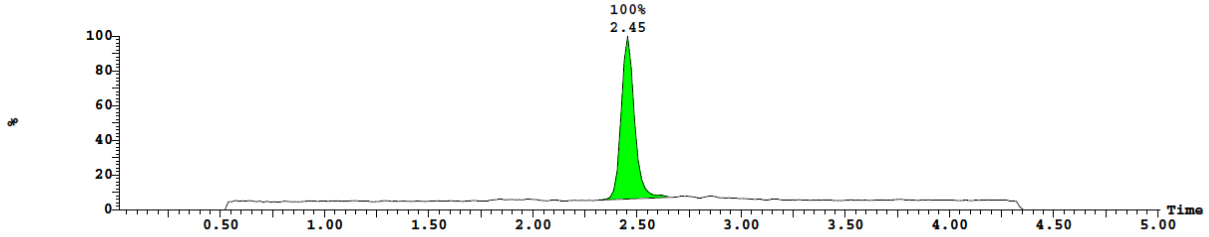

**Conjugate of L13:** MS (ESI-) expected 6026 Da, found 6029 Da.

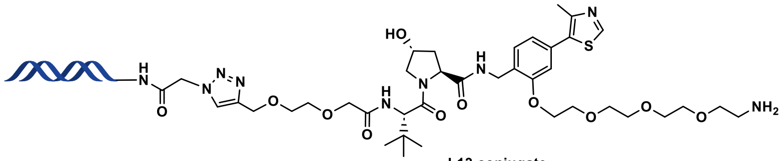

L13 conjugate

***UV:***

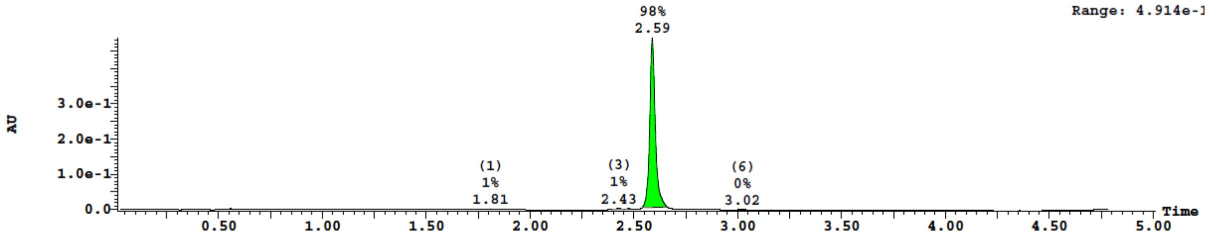

***TIC:***

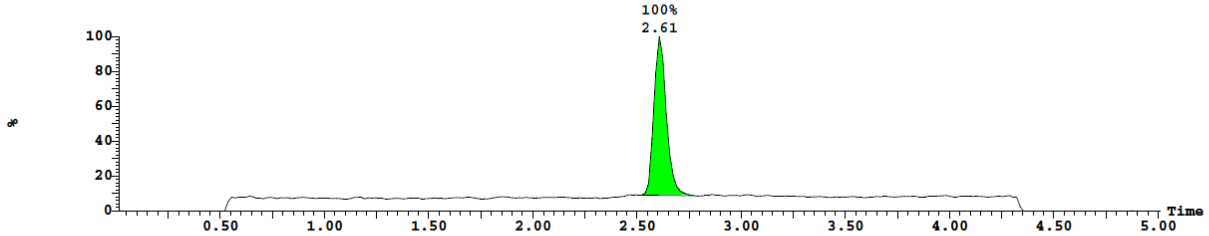

**Conjugate of L15:** MS (ESI-) expected 5977 Da, found 5980 Da.

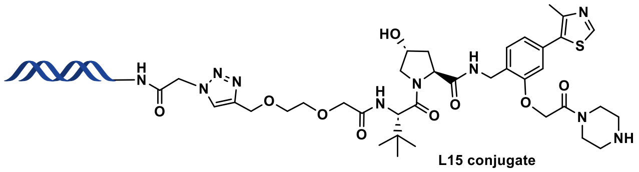

L15 conjugate

***UV:***

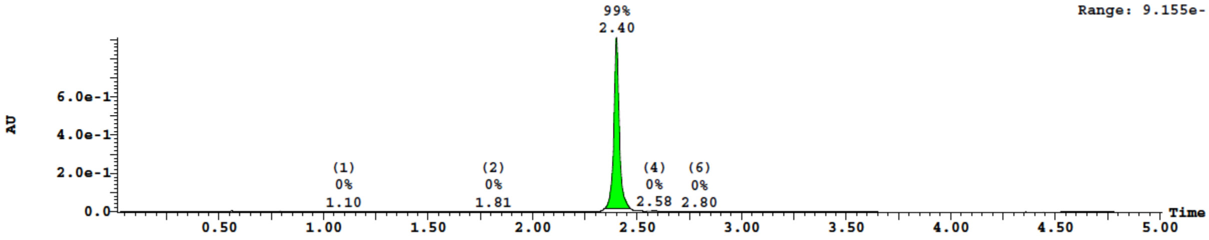

***TIC:***

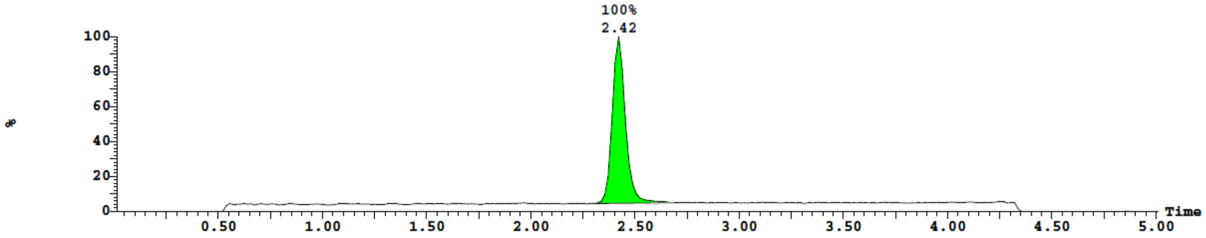

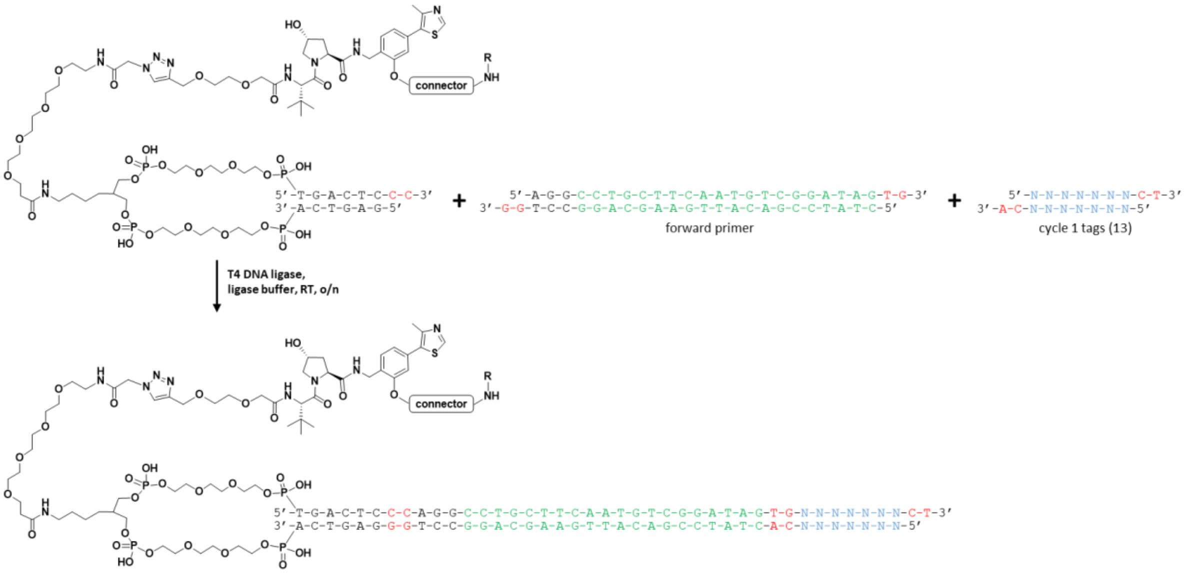

**Scheme L5.** *Library production step 2:* double ligation of forward primer binding sequence and cycle 1 barcodes encoding for connectors.

**Addition of forward primer binding sequence and cycle 1 tags:** Each conjugate (32 nmol, 37 µL, 1.6 equiv) was ligated to the forward primer binding oligo (20 nmol, 1 mM in $dH_2O$, 1 equiv) and a unique cycle 1 tag (44 nmol, 1 mM in $dH_2O$, 2.2 equiv) with the addition of 10× T4 ligase buffer (Lucigen, 20 µL) and T4 DNA ligase (Lucigen, 30 Weiss U / µL, 0.9 µL). Ligation mixtures were gently shaken at room temperature overnight, then analyzed by agarose gel (4% E-gel EX) to confirm reaction completion (**Figure L1**). Each ligation was then EtOH precipitated and the pellets were pooled in to an Amicon Ultra 3 kDa centrifugal filter. The pooled material was buffer exchanged with ~10 volumes of $dH_2O$ and the final concentrated material was analyzed UPLC-MS (**Figure L2**) and quantified by NanoDrop (~quantitative yield, ~260 nmol).

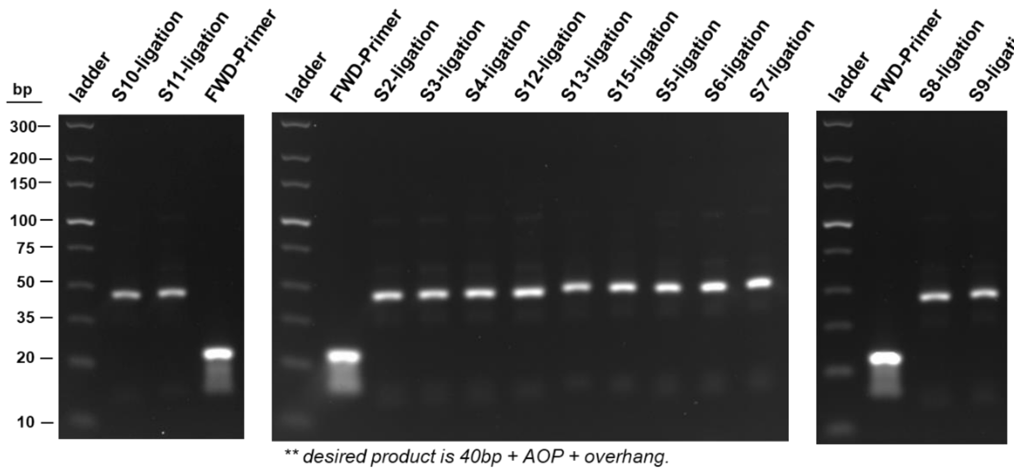

**Figure L1.** Agarose gel analysis of double ligations for addition of forward primer binding oligo and cycle 1 tags for connector encoding.

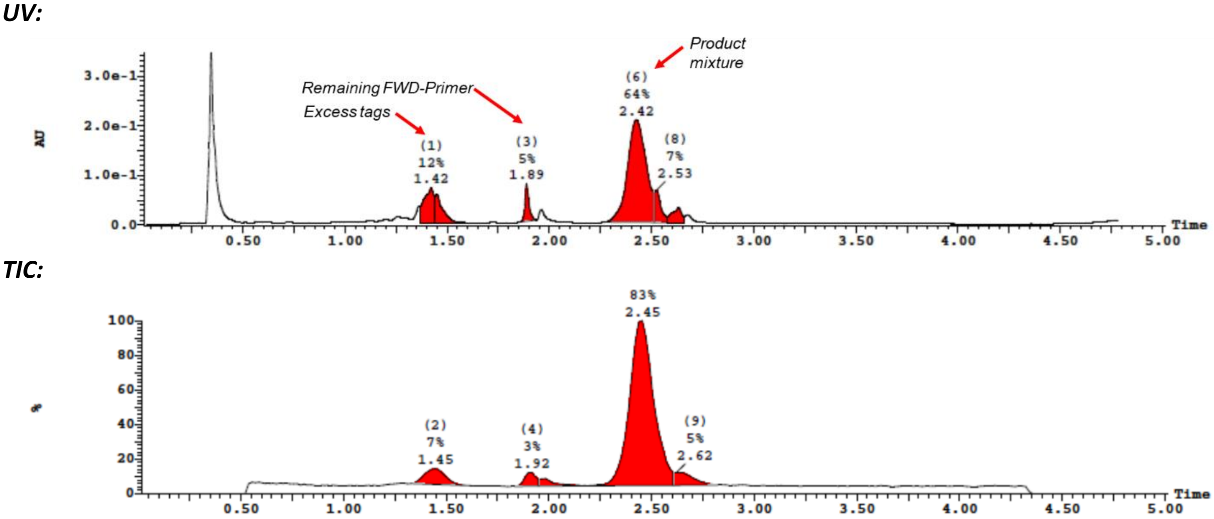

**Figure L2.** UPLC-MS analysis of pooled library after double ligation and buffer exchange.

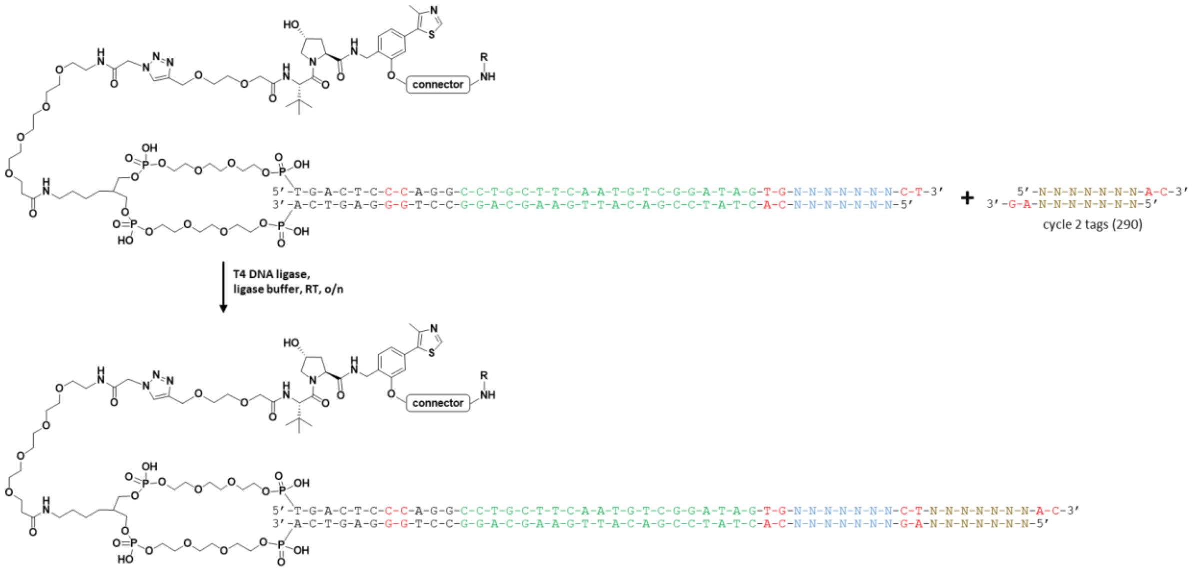

**Scheme L6.** *Library production step 3:* ligation of cycle 2 barcodes encoding for cycle 2 amine building blocks.

**Ligation of cycle 2 tags for amine building block encoding:** The pooled library from above (~260 nmol) was split across 290 wells using 96-well plates (~0.9 nmol per well, 1 equiv). To each well was added a unique cycle 2 tag (2.0 nmol, 2.2 equiv) followed by a mixture of ligase buffer and T4 DNA ligase (1.2 Weiss U / well) for a final volume of 10 µL per ligation. The plates were incubated at room temperature overnight and then analyzed by 4% agarose gel to confirm ligations were complete (**Figure L3**). Each well was EtOH precipitated and pellets were dissolved in 10 µL of 150 mM borate buffer pH 9.4 and used directly for the subsequent reactions.

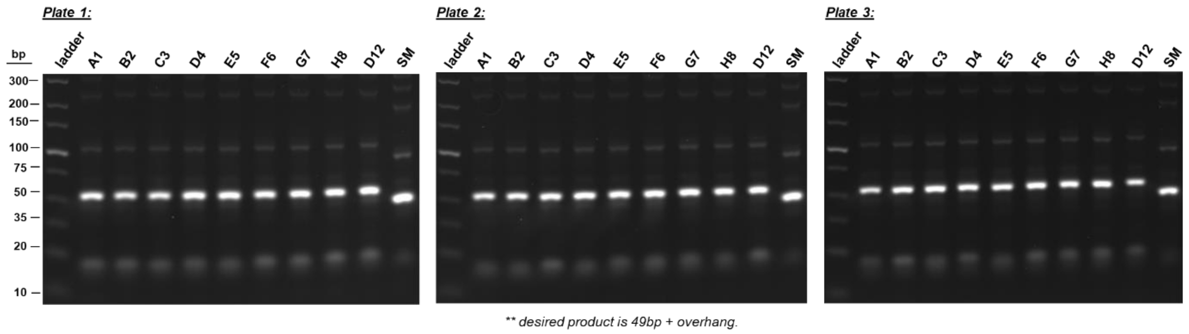

**Figure L3.** Agarose gel analysis of cycle 2 tag ligations across the diagonal wells of each 96-well plates.

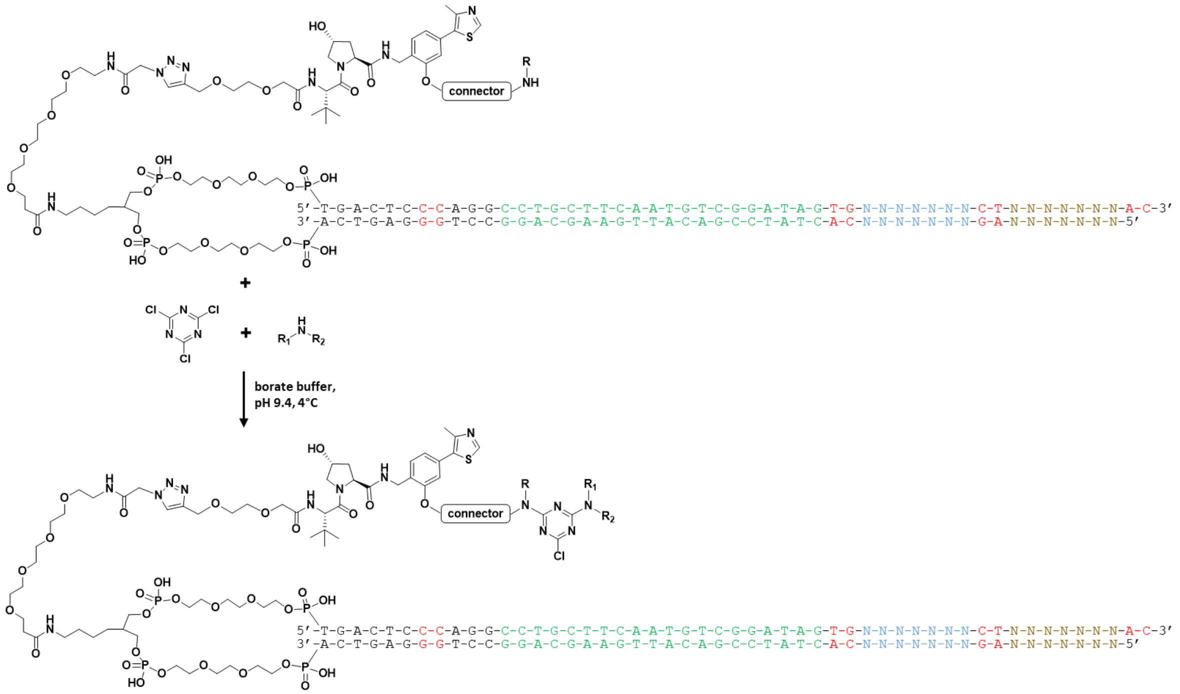

**Scheme L7.** *Library production step 4:* triazine addition to connectors and nucleophilic substitution of triazines with cycle 2 amine building blocks.

**Substitution of connector amines with cyanuric chloride followed by SNAr with 290 cycle 2 amines:** The plates from above containing 290 ligated products (~0.9 nmol in 10 µL of 150 mM borate buffer pH 9.4, 1 equiv) were cooled to 4 °C and cyanuric chloride (5 µL of 3 mM solution in $CH_3CN$, 16 equiv) was added to each well. After standing at 4 °C for 1 hr, each well received one of 290 unique amines (200 mM solutions in 1:1 $CH_3CN$:$dH_2O$, 1 µL, 222 equiv) and the plates were kept at 4 °C overnight. Each well was EtOH precipitated, dissolved in $dH_2O$, and pooled in to an Amicon Ultra 10 kDa spin column. The pooled library was buffer exchanged with ~10 volumes of $dH_2O$ and concentrated.

***UV:***

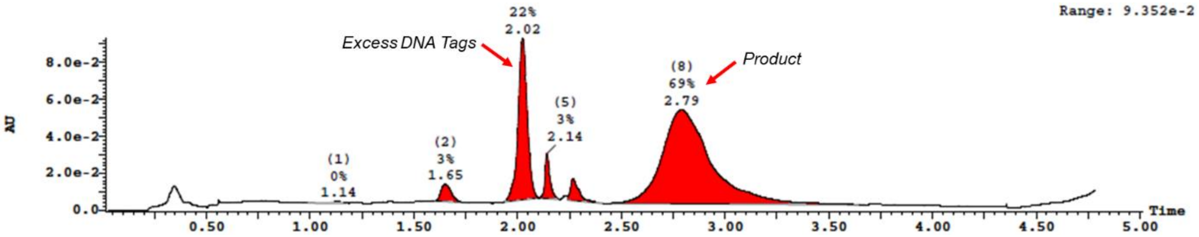

**Figure L4.** UPLC-MS analysis of pooled and buffer exchanged library after SNAr addition of triazines and cycle 2 amines.

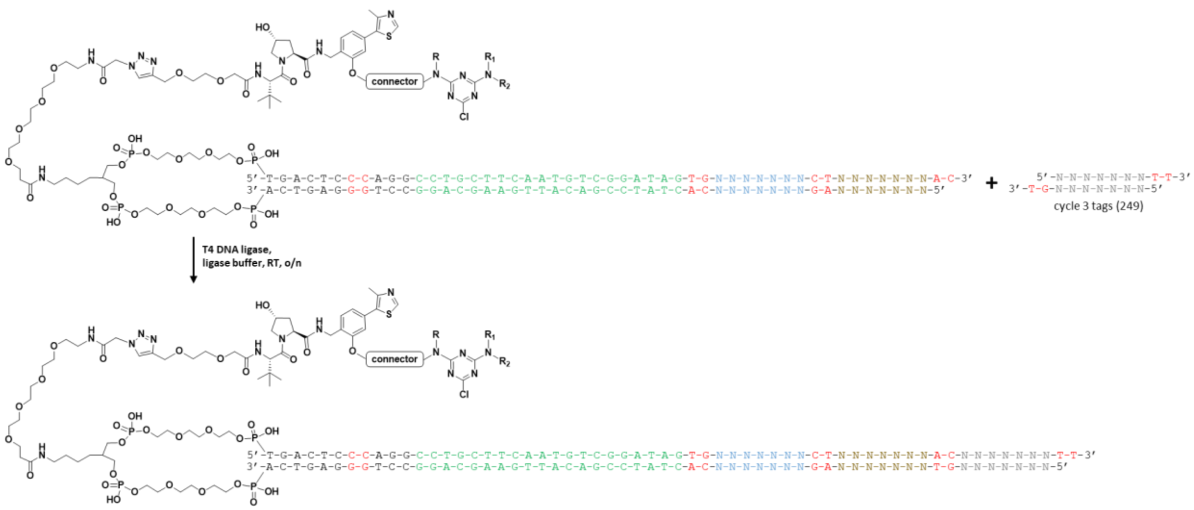

**Scheme L8.** *Library production step 5:* ligation of cycle 3 barcodes encoding for cycle 3 amine building blocks.

**Ligation of 249 cycle 3 barcodes for encoding of cycle 3 amines:** The pooled library from above was split out across 249 wells using 96-well plates (~1.0 nmol per well, 1 equiv). To each well was added a unique cycle 3 barcode (2.2 nmol, 2.2 equiv) and a freshly prepared solution of T4 DNA ligase in 5× ligase buffer (1.2 Weiss U / well) to give a 10 µL final reaction volume per well. After a brief centrifugation, the plates were incubated at room temperature overnight. Ligations were then analyzed by 4% agarose gel electrophoresis to confirm the reactions were complete (**Figure L5**) and DNA was EtOH precipitated in each well. The air-dried pellets were dissolved in 150 mM borate buffer pH 9.4 (18 µL) and used directly in the next step.

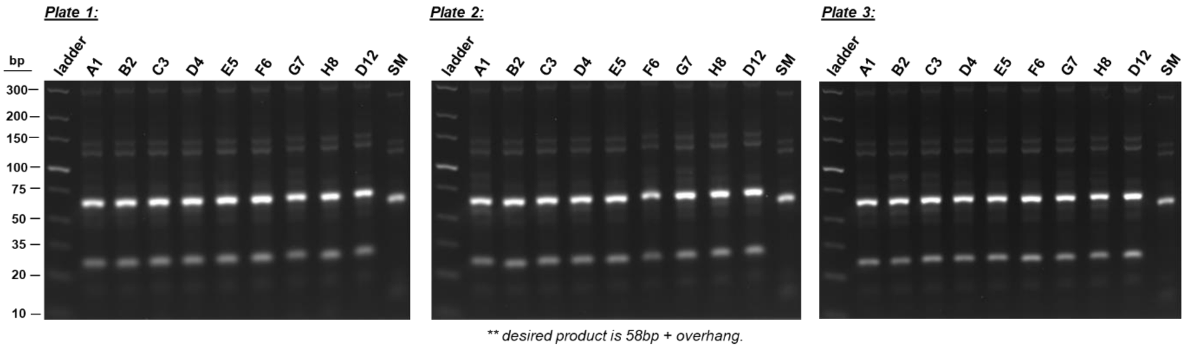

**Figure L5.** Agarose gel analysis of cycle 3 tag ligations across the diagonal wells of each 96-well plates.

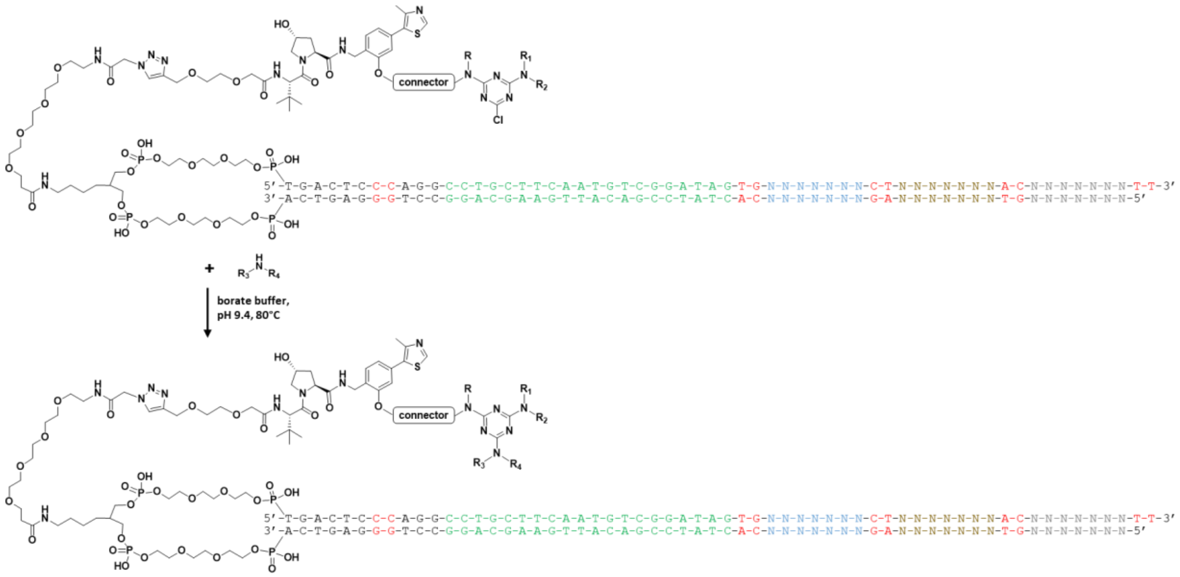

**Scheme L9.** *Library production step 6:* nucleophilic substitution of triazines with cycle 3 amine building blocks.

**Final substitution of triazines with 249 cycle 3 amines:** To each well from the above ligations was added a unique cycle 3 amine (200 mM solutions in 1:1 $CH_3CN$:$dH_2O$, 2 µL, 400 equiv) and the plates were sealed and heated to 80 °C for 16 hr. Reactions were then EtOH precipitated and the pellets were dissolved in $dH_2O$ and transferred to an Amicon Ultra 10 kDa centrifugal filter. The pooled library was buffer exchanged with ~10 volumes of $dH_2O$ and the final concentrated library was analyzed by UPLC-MS (**Figure L6**).

***UV:***

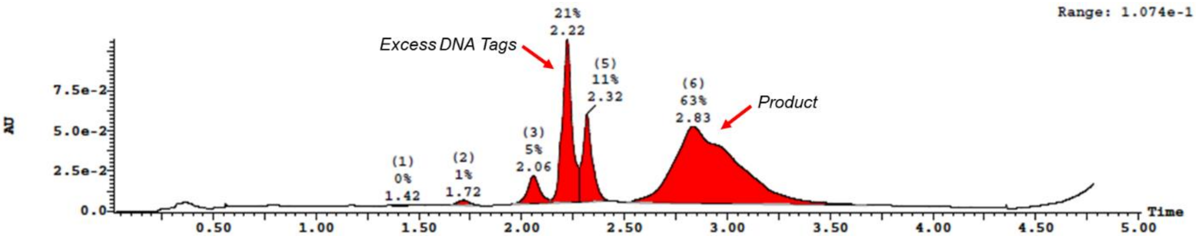

**Figure L6.** UPLC-MS analysis of pooled and buffer exchanged library after SNAr addition of triazines and cycle 2 amines.

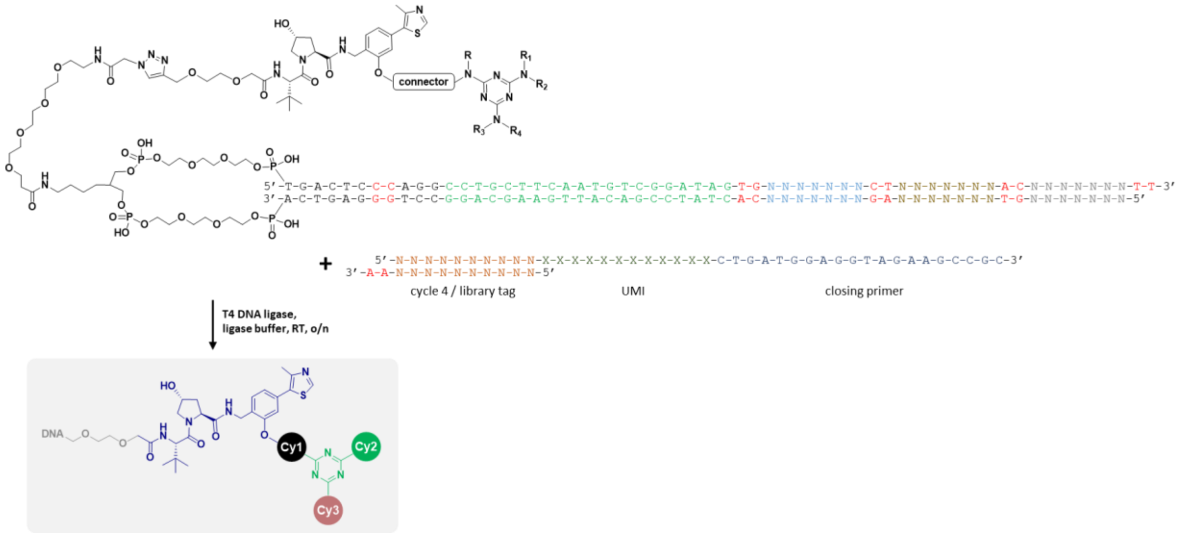

**Scheme L10.** *Library production step 7:* ligation of library closing primer containing a library tag, UMI, and reverse primer binding site to generate the final CIP-DEL.

**Closing tag ligation:** To the DNA barcodes was added a final closing DNA sequence consisting of a double-stranded library identification tag, a single-stranded random sequence providing a unique molecular identified (UMI), and a single-stranded reverse primer binding site. One quarter of the crude material was taken on for closing tag addition (260 nmol / 4 = 65 nmol of library). To the pooled library was added the closing tag (library tag: top 5′ – CAGAGTGCCA – 3′, 118 nmol, 1.8 equiv), 10× ligase buffer (Lucigen), and T4 DNA ligase (78 Weiss U) in a 400 µL total reaction volume. The ligation mixture was gently vortexed, briefly centrifuged, and incubated at room temperature overnight. Analysis of the reaction by agarose gel electrophoresis indicated near complete ligation (**Figure L7.a**) and the DNA was EtOH precipitated. The crude ligation mixture was purified by preparative agarose gel electrophoresis using a BioRad 491 Prep Cell per the manufacturer's instructions and as described previously[1]. The isolated fractions were analyzed by agarose gel (**Figure L7.b**) and product containing fractions were pooled in to a 10 kDa Amicon Ultra filter. The material was buffer exchanged with ~10 volumes of $dH_2O$ and concentrated. The final purified library was analyzed by UPLC-MS (**Figure L8**) and quantified by NanoDrop (25.1 nmol, ~39% yield) and QuBit dsDNA BR assay (Invitrogen) per the manufacturer's instructions (14.1 nmol, ~22% yield).

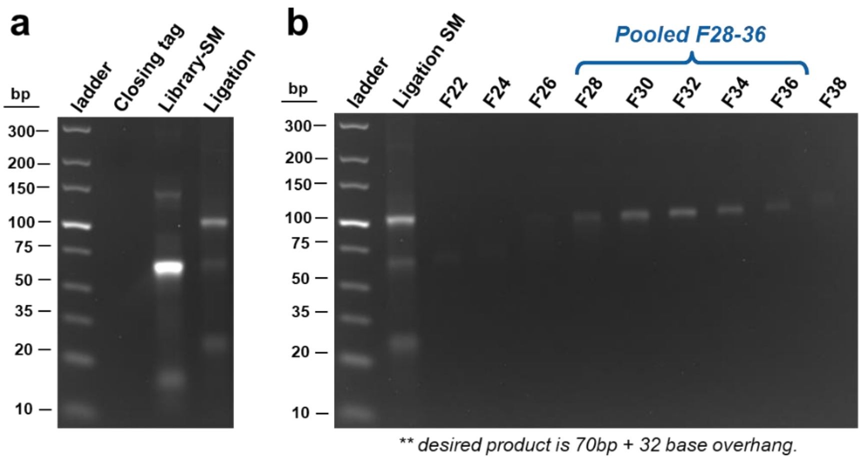

**Figure L7. (a)** Agarose gel analysis of closing tag ligation. **(b)** Agarose gel analysis of fractions obtained from BioRad 491 Prep Cell purification of final closed library.

***UV:***

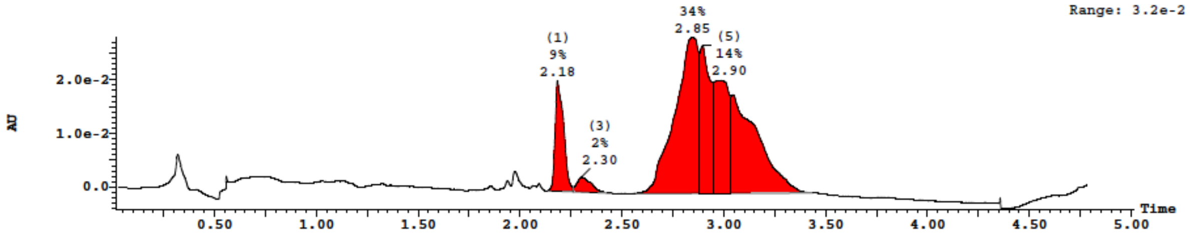

**Figure L8.** UPLC-MS analysis of final pooled, purified, and buffer exchanged library.

### NMR spectra of compounds

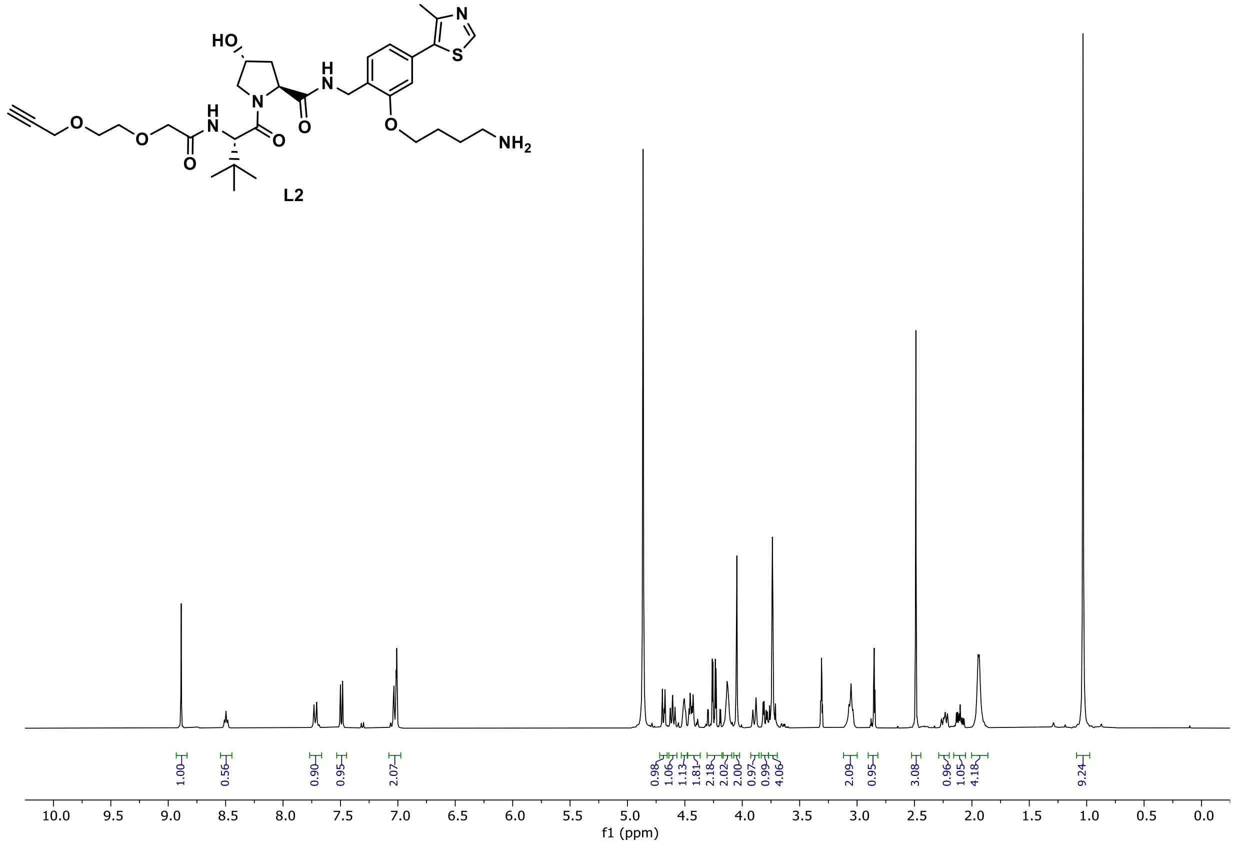

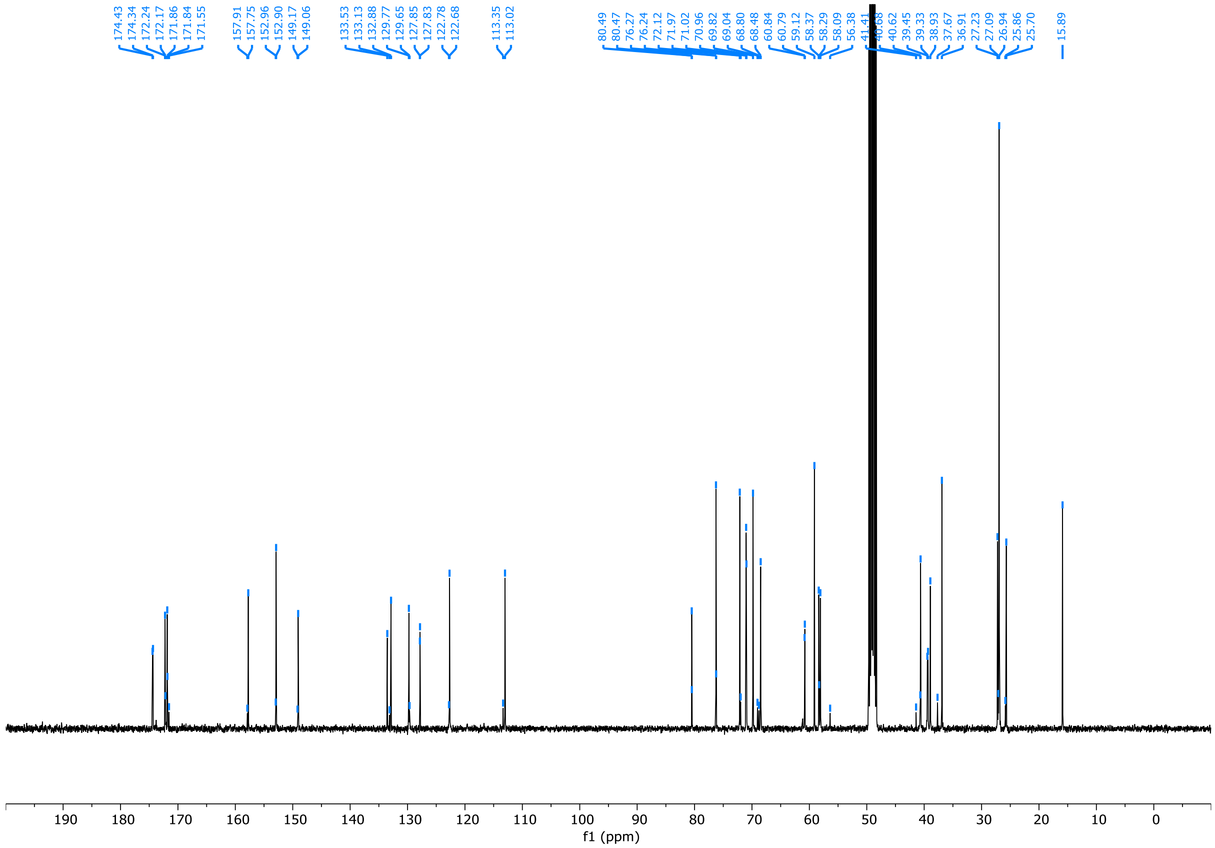
174.43
174.34
172.24
172.17
171.86
171.84
171.55
157.91
157.75
152.96
152.90
149.17
149.06
133.53
133.13
132.88
129.77
129.65
127.85
127.83
122.78
122.68
113.35
113.02
80.49
80.47
76.27
76.24
72.12
71.97
71.02
70.96
69.82
69.04
68.80
68.48
60.84
60.79
59.12
58.37
58.29
58.09
56.38
41.41
40.68
40.62
39.45
39.33
38.93
37.67
36.91
27.23
27.09
26.94
25.86
25.70
15.89
190 180 170 160 150 140 130 120 110 100 90 80 70 60 50 40 30 20 10 0
f1 (ppm)

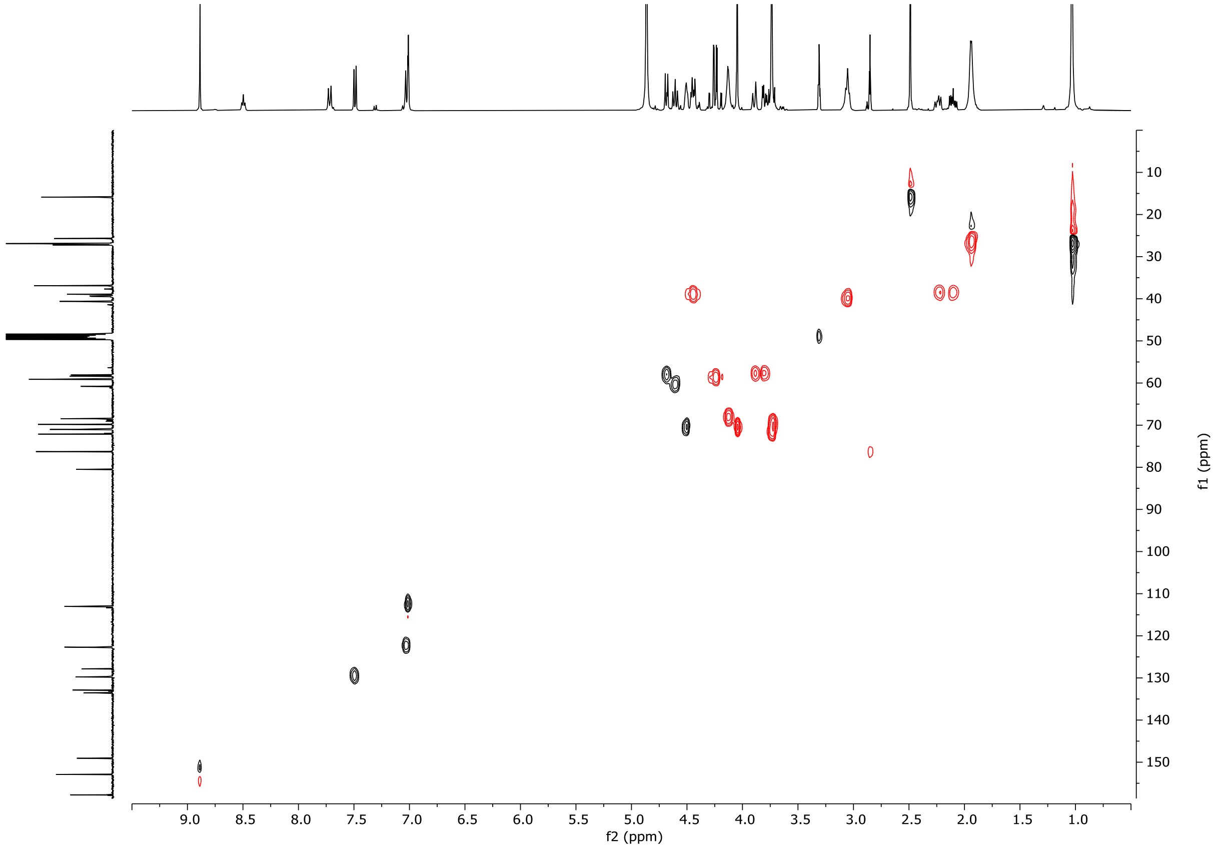
9.0 8.5 8.0 7.5 7.0 6.5 6.0 5.5 5.0 4.5 4.0 3.5 3.0 2.5 2.0 1.5 1.0
f2 (ppm)
10 20 30 40 50 60 70 80 90 100 110 120 130 140 150
f1 (ppm)

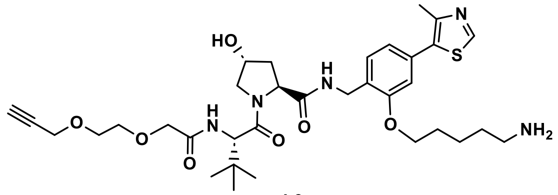
HO
N
H
N
O
O
O
H
N
O
O
N
S
O
NH2

L3

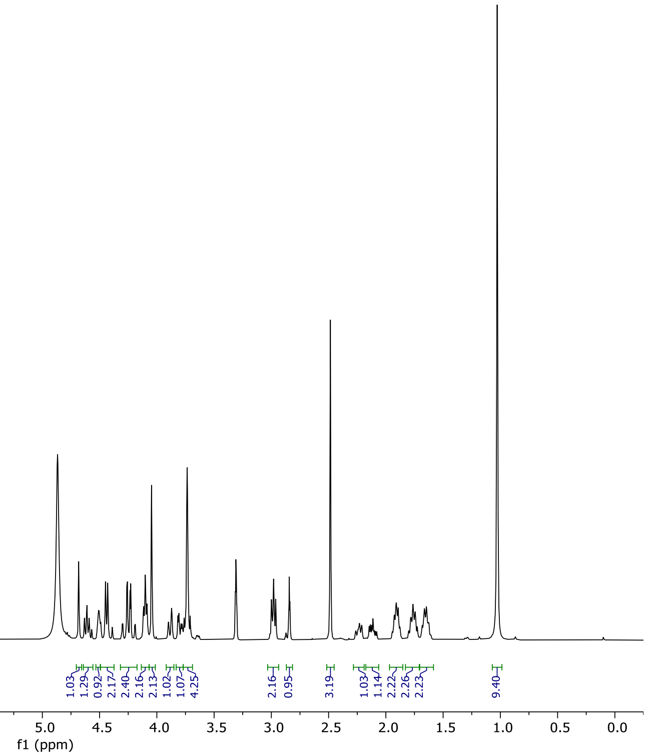
1.00
0.35
0.99
2.06
1.03
1.29
0.92
2.17
2.40
2.16
2.13
1.02
1.07
4.25
2.16
0.95
3.19
1.03
1.14
2.22
2.26
2.23
9.40
10.0 9.5 9.0 8.5 8.0 7.5 7.0 6.5 6.0 5.5 5.0 4.5 4.0 3.5 3.0 2.5 2.0 1.5 1.0 0.5 0.0
f1 (ppm)

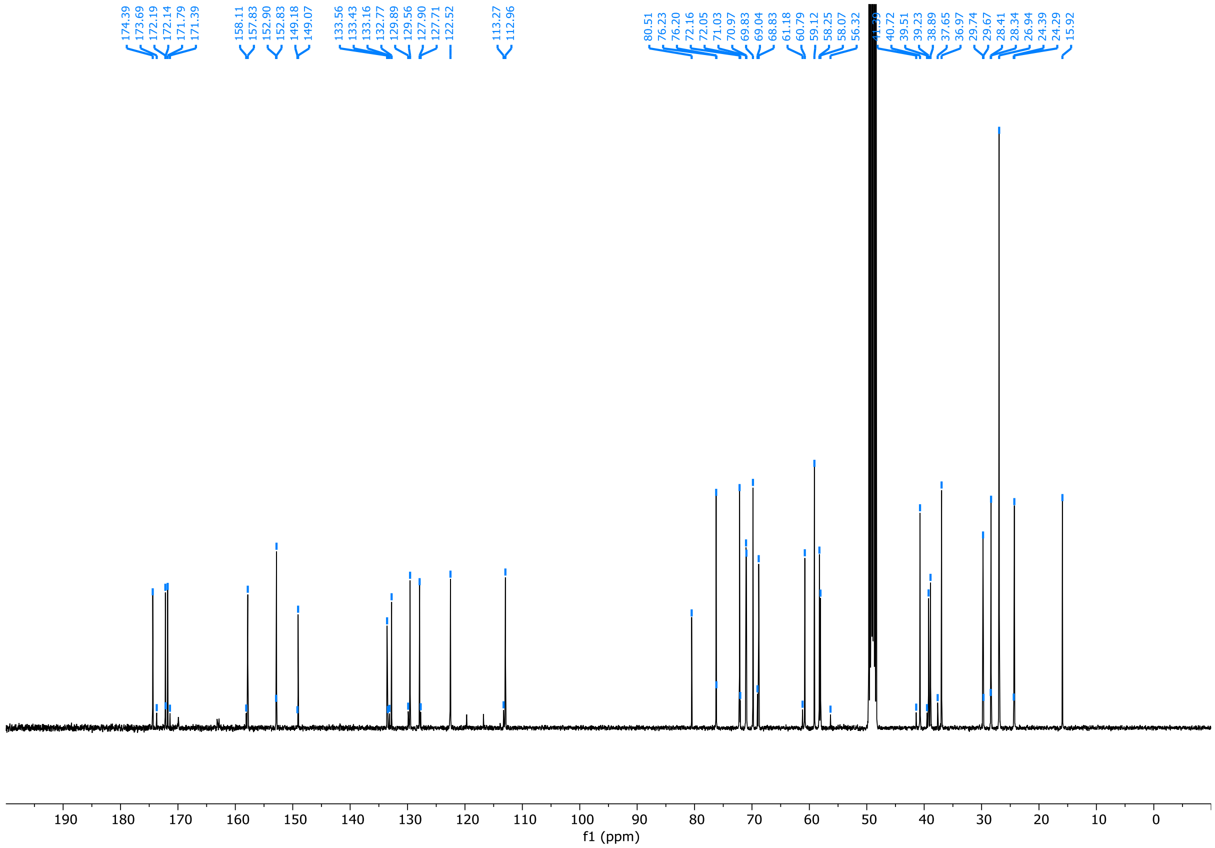
174.39
173.69
172.19
172.14
171.79
171.39
158.11
157.83
152.90
152.83
149.18
149.07
133.56
133.43
133.16
132.77
129.89
129.56
127.90
127.71
122.52
113.27
112.96
80.51
76.23
76.20
72.16
72.05
71.03
70.97
69.83
69.04
68.83
61.18
60.79
59.12
58.25
58.07
56.32
40.72
39.51
39.23
38.89
37.65
36.97
29.74
29.67
28.41
28.34
26.94
24.39
24.29
15.92
190 180 170 160 150 140 130 120 110 100 90 80 70 60 50 40 30 20 10 0
f1 (ppm)

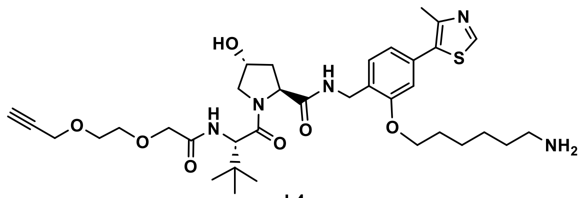
HO
N
H
N
O
O
N
H
O
O
O
N
S
O
NH2

**L4**

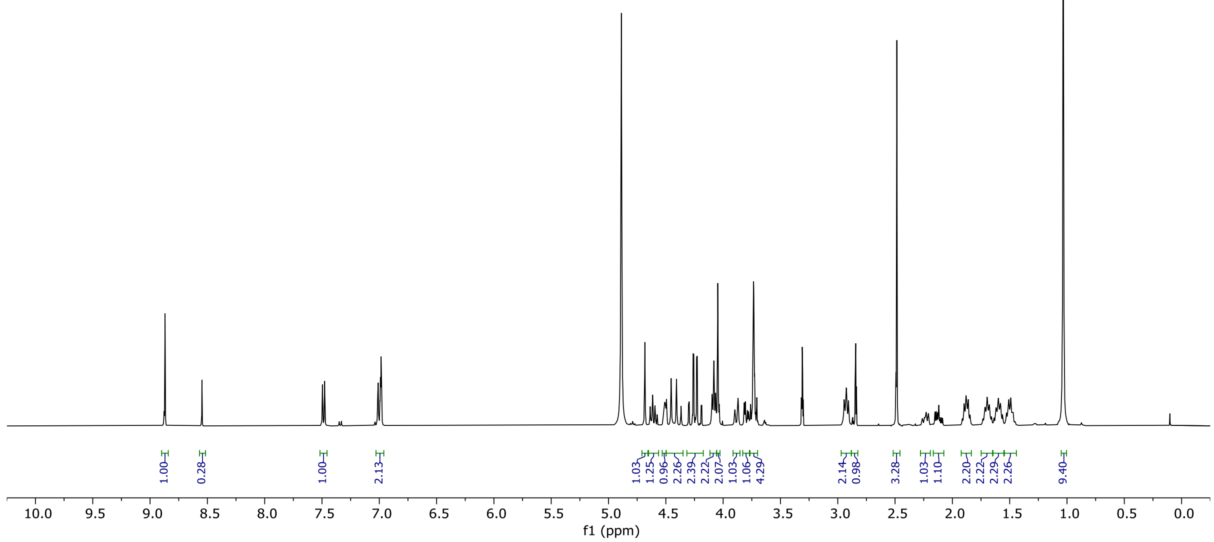
1.00
0.28
1.00
2.13
1.03
1.25
0.96
2.26
2.39
2.22
2.07
1.03
1.06
4.29
2.14
0.98
3.28
1.03
1.10
2.20
2.22
2.29
2.26
9.40
10.0 9.5 9.0 8.5 8.0 7.5 7.0 6.5 6.0 5.5 5.0 4.5 4.0 3.5 3.0 2.5 2.0 1.5 1.0 0.5 0.0
f1 (ppm)

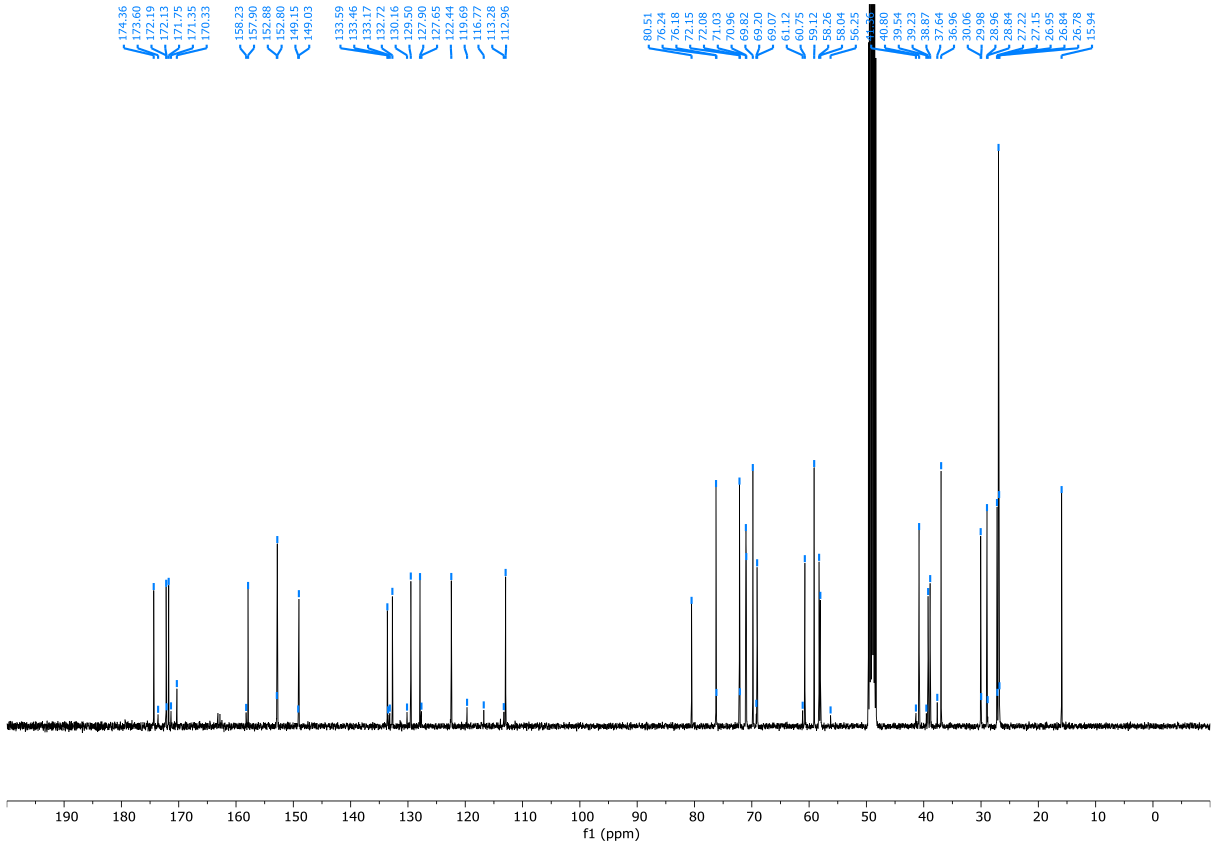
174.36
173.60
172.19
172.13
171.75
171.35
170.33
158.23
157.90
152.88
152.80
149.15
149.03
133.59
133.46
133.17
132.72
130.16
129.50
127.90
127.65
122.44
119.69
116.77
113.28
112.96
80.51
76.24
76.18
72.15
72.08
71.03
70.96
69.82
69.20
69.07
61.12
60.75
59.12
58.26
58.04
56.25
40.80
39.54
39.23
38.87
37.64
36.96
30.06
29.98
28.96
28.84
27.22
27.15
26.95
26.84
26.78
15.94
190 180 170 160 150 140 130 120 110 100 90 80 70 60 50 40 30 20 10 0
f1 (ppm)

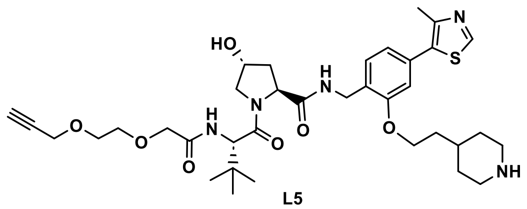
HO
N
H
N
O
O
O
N
H
O
O
N
S
O
NH

L5

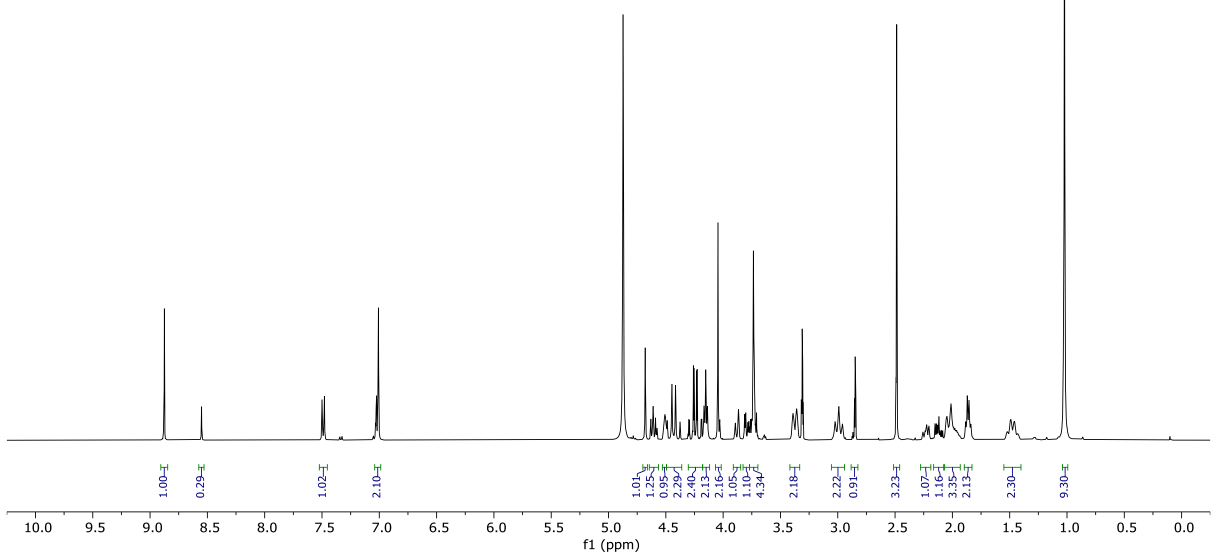
1.00
0.29
1.02
2.10
1.01
1.25
0.95
2.29
2.40
2.13
2.16
1.05
1.10
4.34
2.18
2.22
0.91
3.23
1.07
1.16
3.35
2.13
2.30
9.30
10.0 9.5 9.0 8.5 8.0 7.5 7.0 6.5 6.0 5.5 5.0 4.5 4.0 3.5 3.0 2.5 2.0 1.5 1.0 0.5 0.0
f1 (ppm)

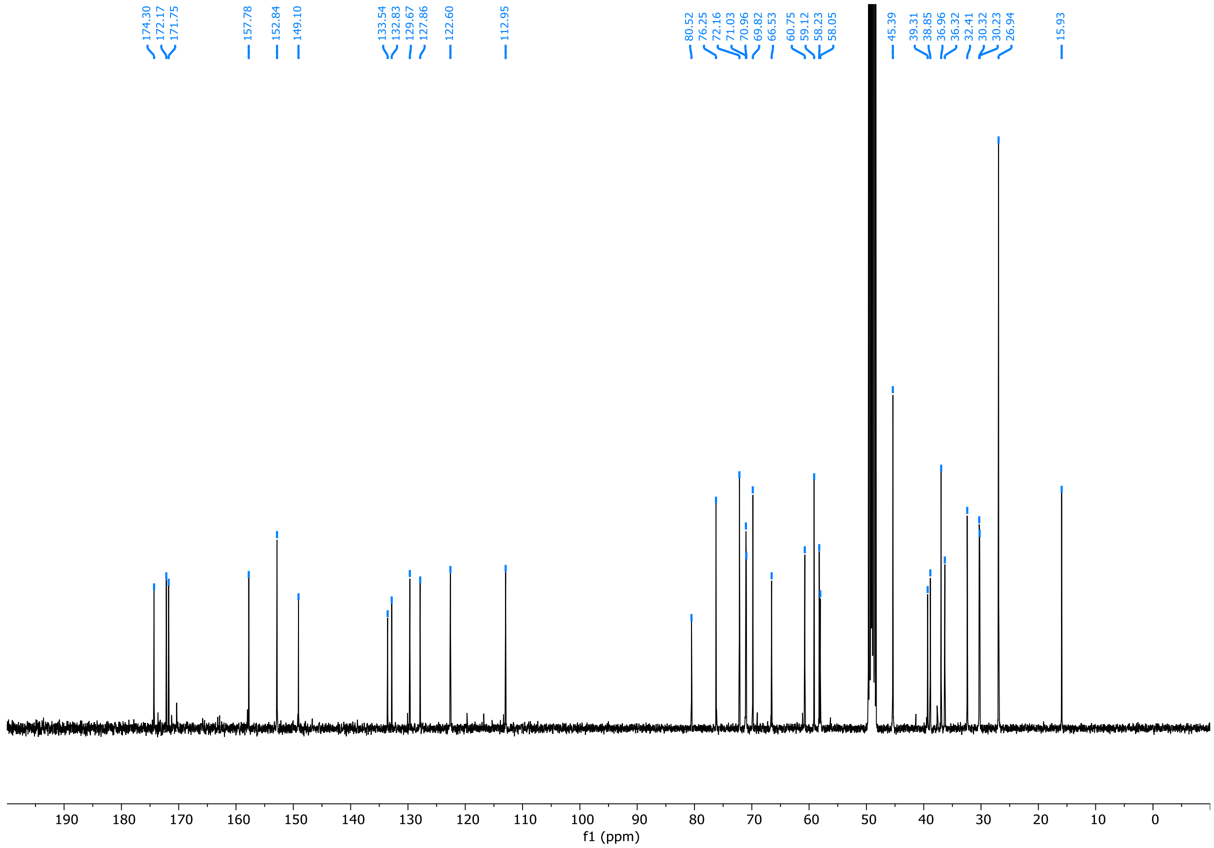
174.30
172.17
171.75
157.78
152.84
149.10
133.54
132.83
129.67
127.86
122.60
112.95
80.52
76.25
72.16
71.03
70.96
69.82
66.53
60.75
59.12
58.23
58.05
45.39
39.31
38.85
36.96
36.32
32.41
30.32
30.23
26.94
15.93
190 180 170 160 150 140 130 120 110 100 90 80 70 60 50 40 30 20 10 0
f1 (ppm)

**L6**

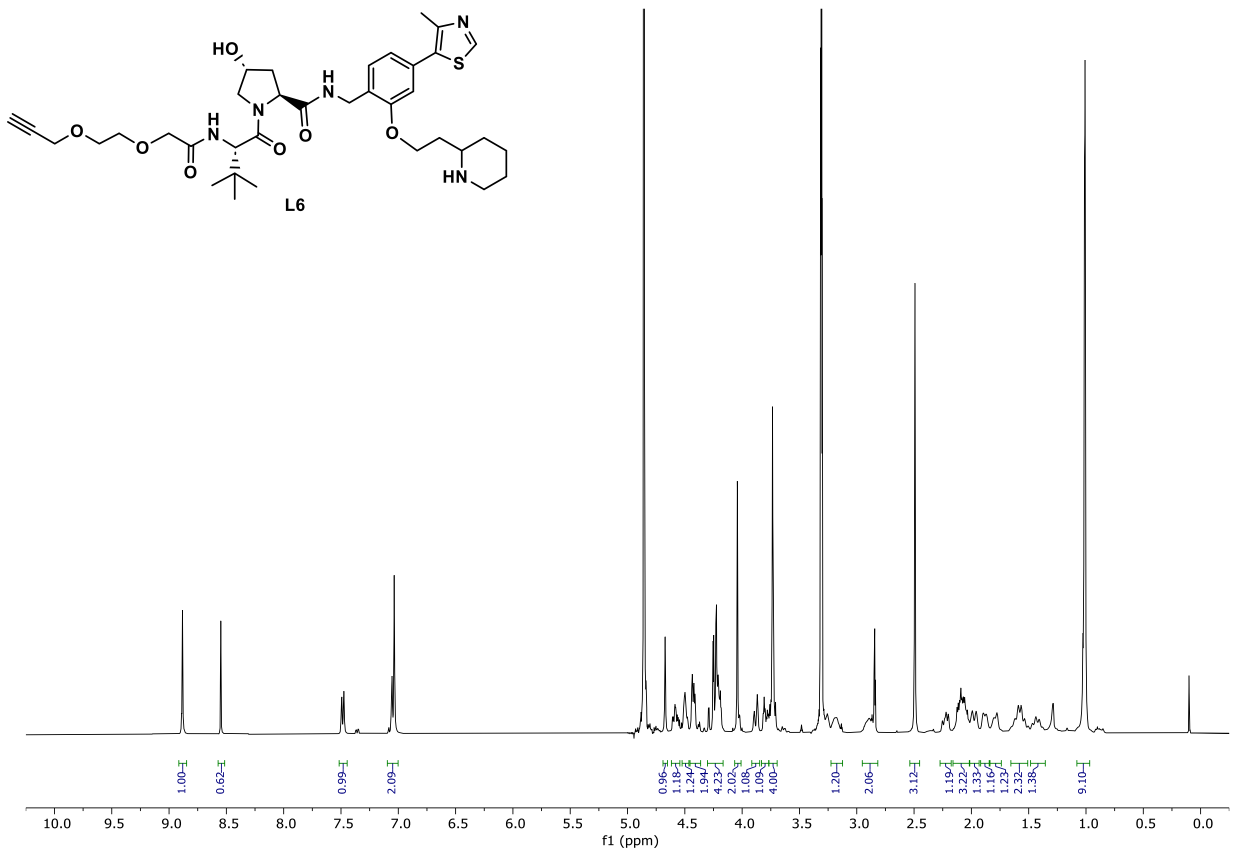

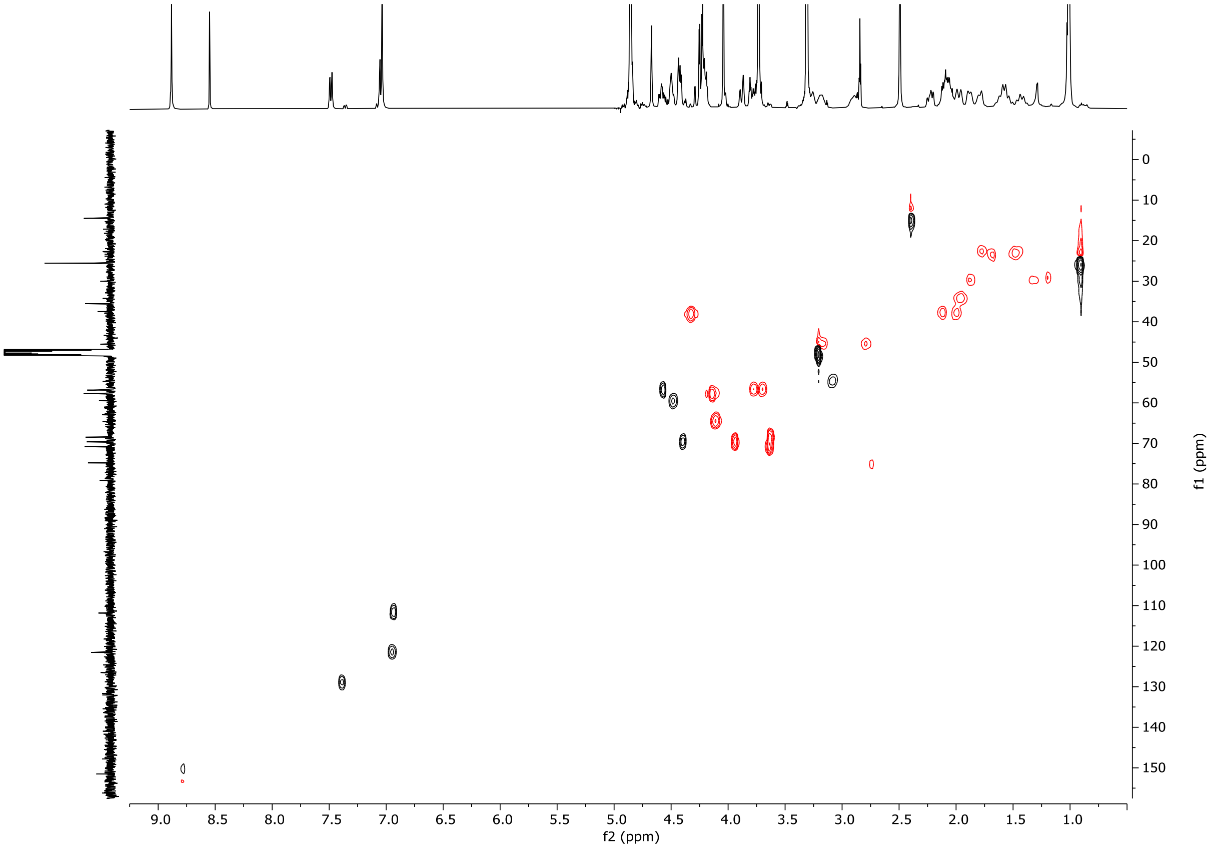

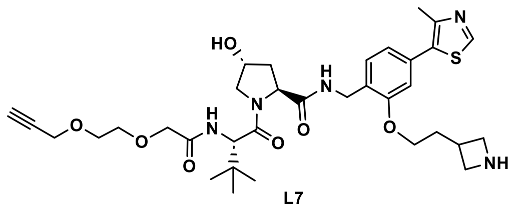
HO
N
H
O
O
O
N
H
N
O
S
N
O
NH

L7

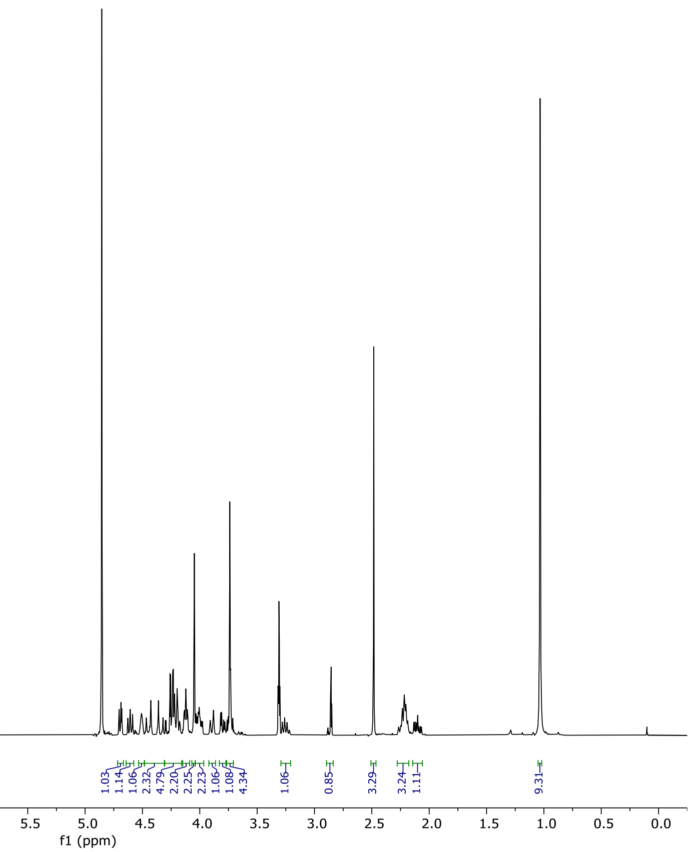
1.00
0.74
1.02
1.12
1.06
1.03
1.14
1.06
2.32
4.79
2.20
2.25
2.23
1.06
1.08
4.34
1.06
0.85
3.29
3.24
1.11
9.31
10.0 9.5 9.0 8.5 8.0 7.5 7.0 6.5 6.0 5.5 5.0 4.5 4.0 3.5 3.0 2.5 2.0 1.5 1.0 0.5 0.0
f1 (ppm)

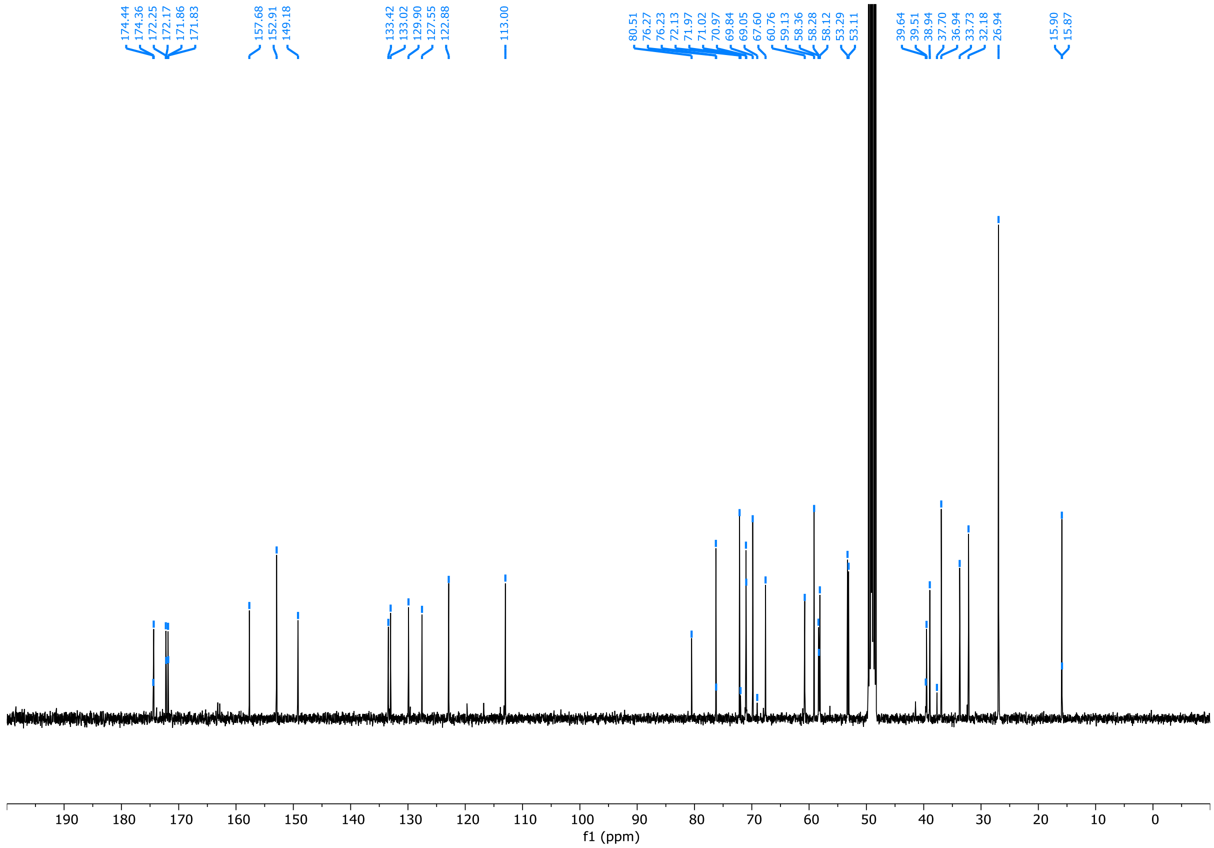
174.44
174.36
172.25
172.17
171.86
171.83
157.68
152.91
149.18
133.42
133.02
129.90
127.55
122.88
113.00
80.51
76.27
76.23
72.13
71.97
71.02
70.97
69.84
69.05
67.60
60.76
59.13
58.36
58.28
58.12
53.29
53.11
39.64
39.51
38.94
37.70
36.94
33.73
32.18
26.94
15.90
15.87
190 180 170 160 150 140 130 120 110 100 90 80 70 60 50 40 30 20 10 0
f1 (ppm)

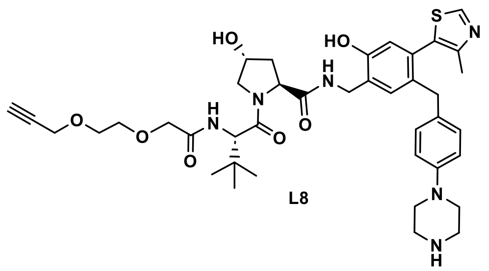
HO
HO
S
N
H
N
O
N
O
O
H
N
O
O
N
N
H

L8

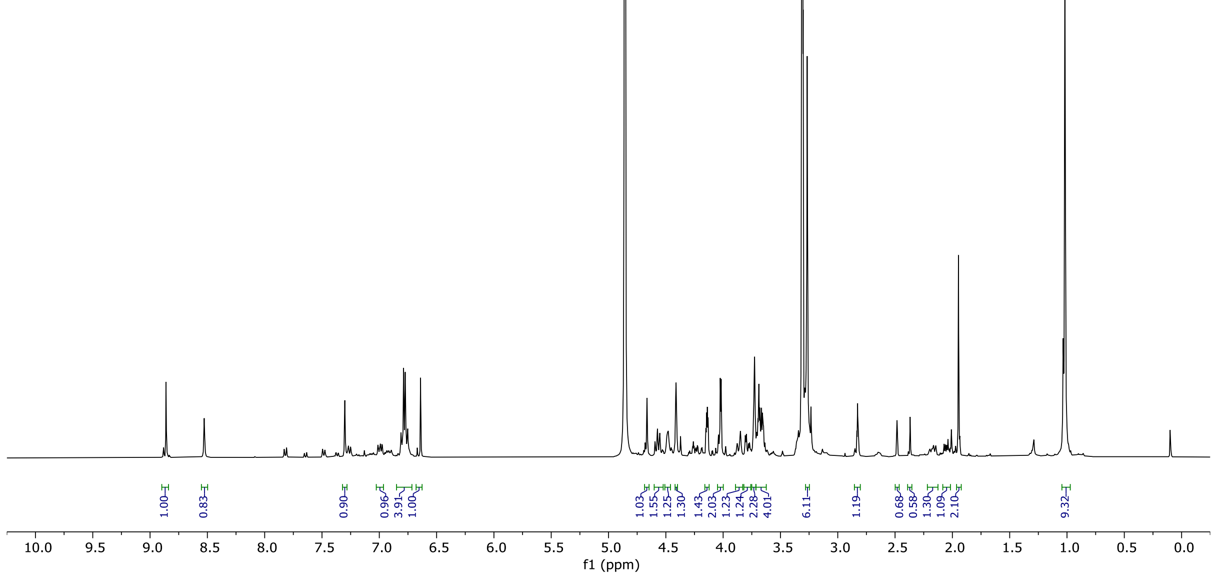
1.00
0.83
0.90
0.96
3.91
1.00
1.03
1.55
1.25
1.30
1.43
2.03
1.23
1.24
2.28
4.01
6.11
1.19
0.68
0.58
1.30
1.09
2.10
9.32
10.0 9.5 9.0 8.5 8.0 7.5 7.0 6.5 6.0 5.5 5.0 4.5 4.0 3.5 3.0 2.5 2.0 1.5 1.0 0.5 0.0
f1 (ppm)

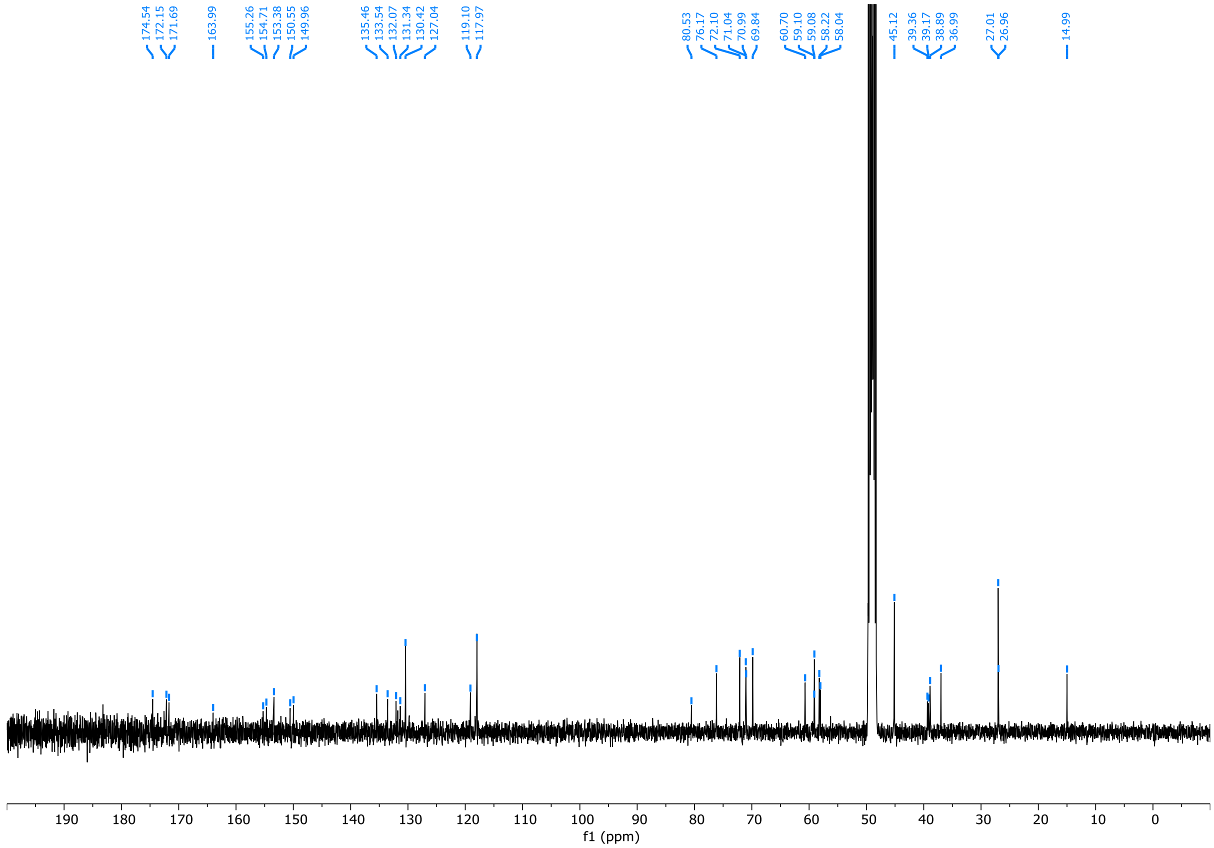
174.54
172.15
171.69
163.99
155.26
154.71
153.38
150.55
149.96
135.46
133.54
132.07
131.34
130.42
127.04
119.10
117.97
80.53
76.17
72.10
71.04
70.99
69.84
60.70
59.10
59.08
58.22
58.04
45.12
39.36
39.17
38.89
36.99
27.01
26.96
14.99
190 180 170 160 150 140 130 120 110 100 90 80 70 60 50 40 30 20 10 0
f1 (ppm)

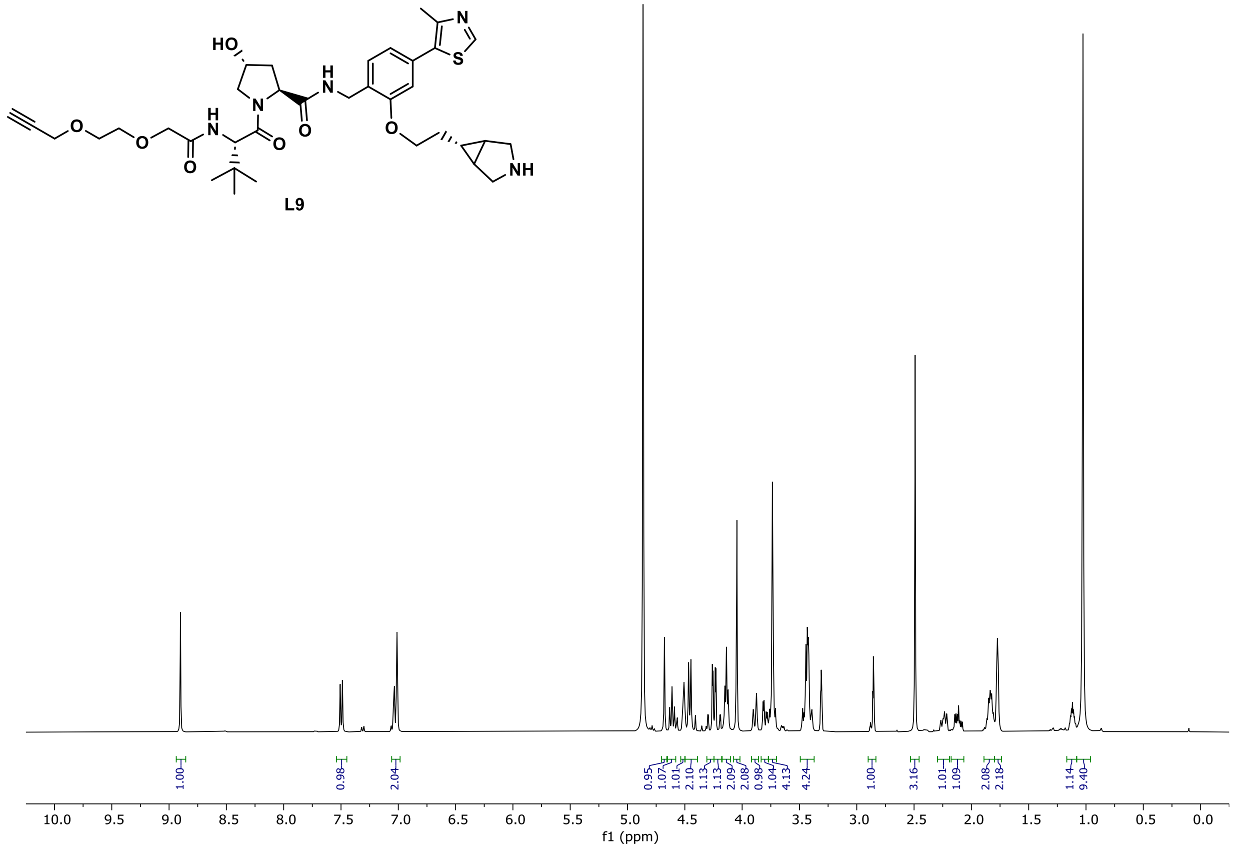
HO
N
H
O
S
NH
L9
1.00
0.98
2.04
0.95
1.07
1.01
2.10
1.13
1.13
2.09
2.08
0.98
1.04
4.13
4.24
1.00
3.16
1.01
1.09
2.08
2.18
1.14
9.40
10.0 9.5 9.0 8.5 8.0 7.5 7.0 6.5 6.0 5.5 5.0 4.5 4.0 3.5 3.0 2.5 2.0 1.5 1.0 0.5 0.0
f1 (ppm)

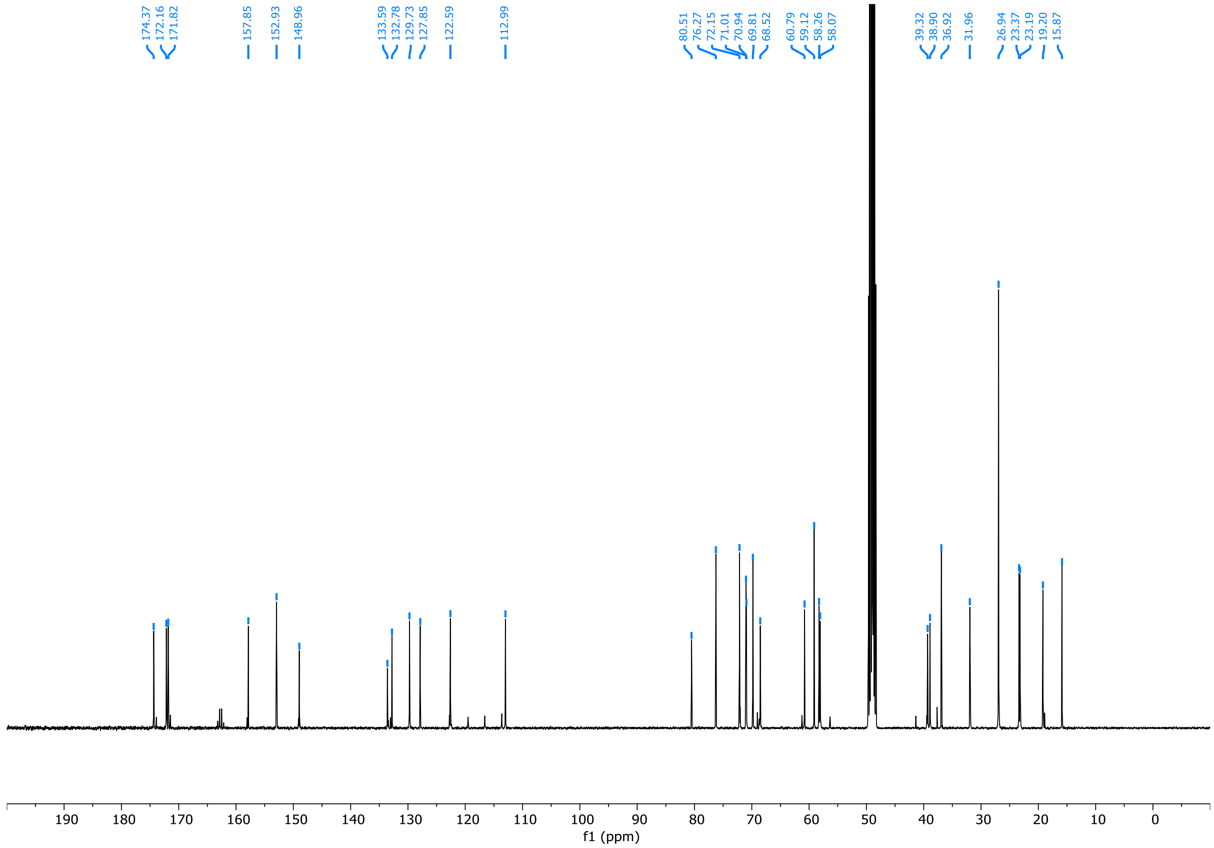
174.37
172.16
171.82
157.85
152.93
148.96
133.59
132.78
129.73
127.85
122.59
112.99
80.51
76.27
72.15
71.01
70.94
69.81
68.52
60.79
59.12
58.26
58.07
39.32
38.90
36.92
31.96
26.94
23.37
23.19
19.20
15.87
190 180 170 160 150 140 130 120 110 100 90 80 70 60 50 40 30 20 10 0
f1 (ppm)

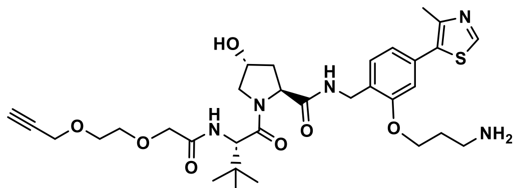

L10

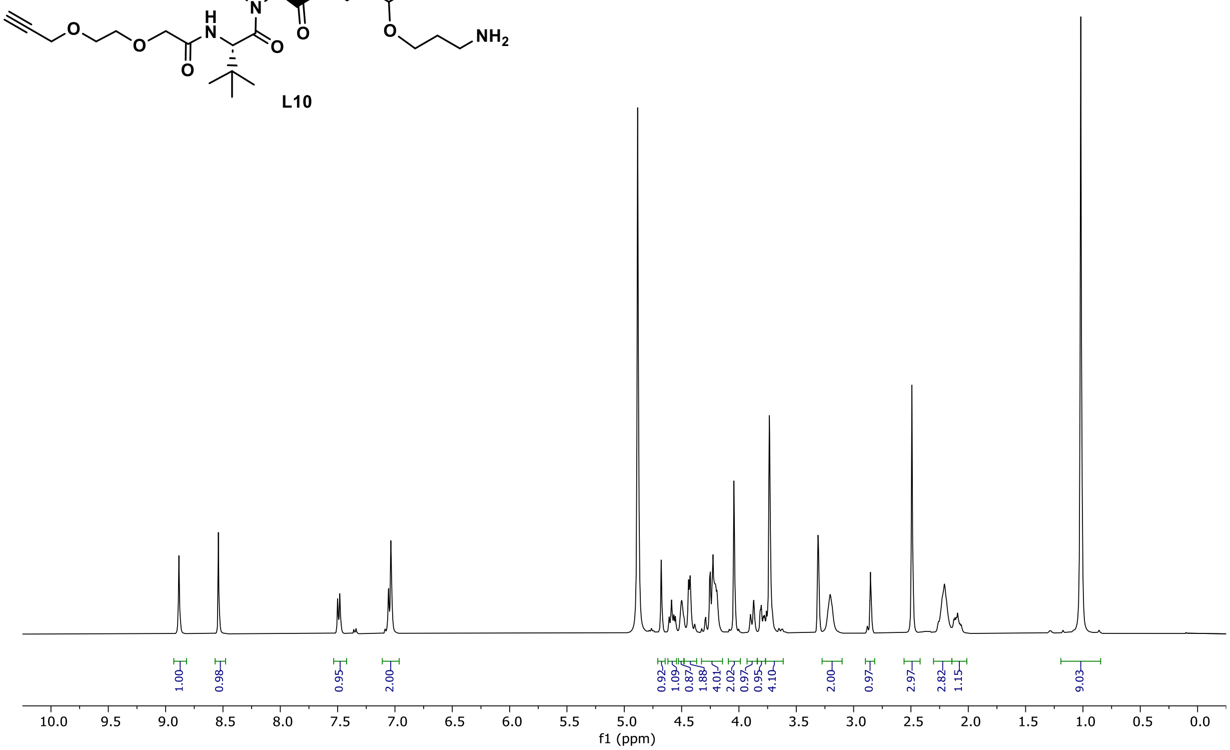

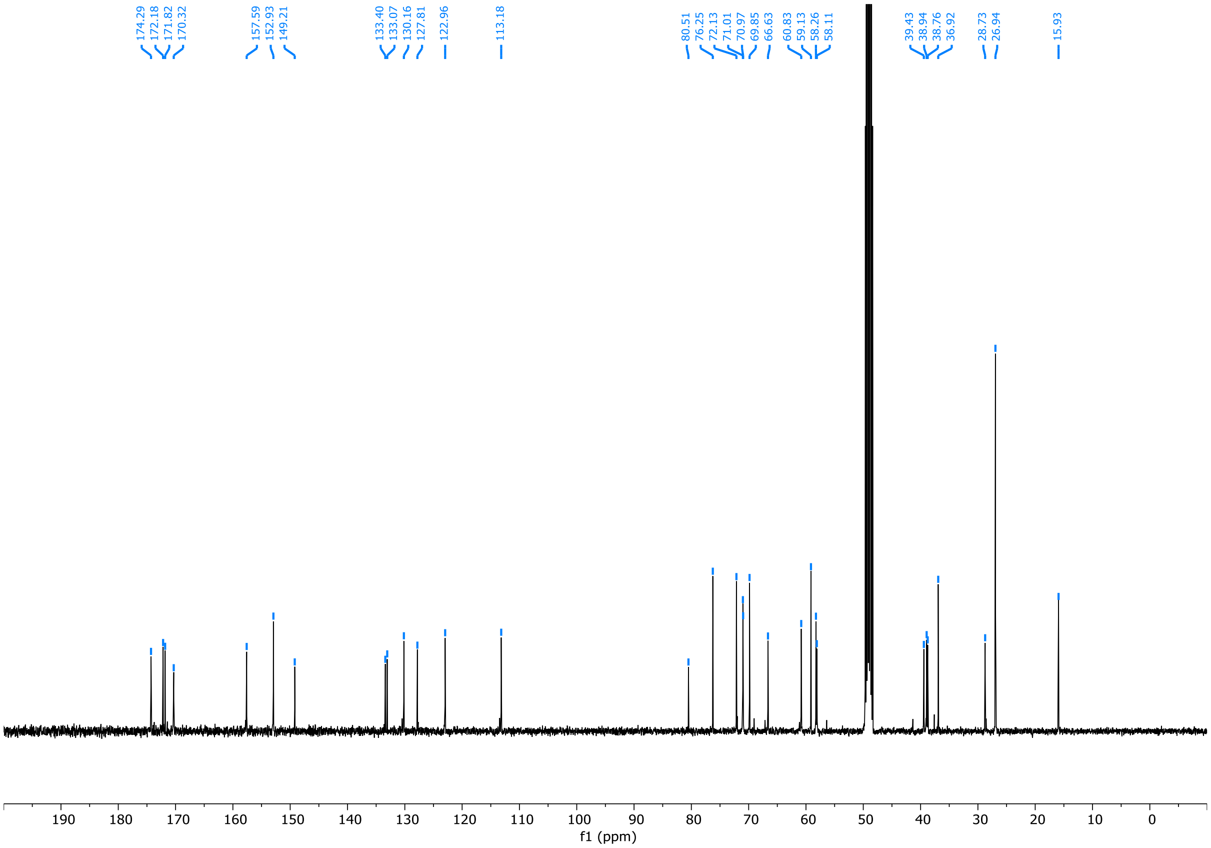

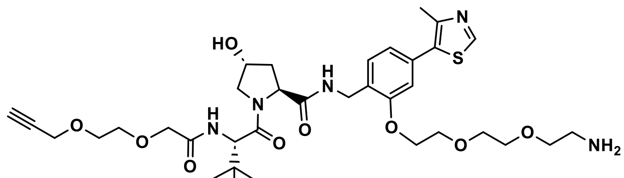
HO
N
H
N
S
O
O
O
O
NH2

L11

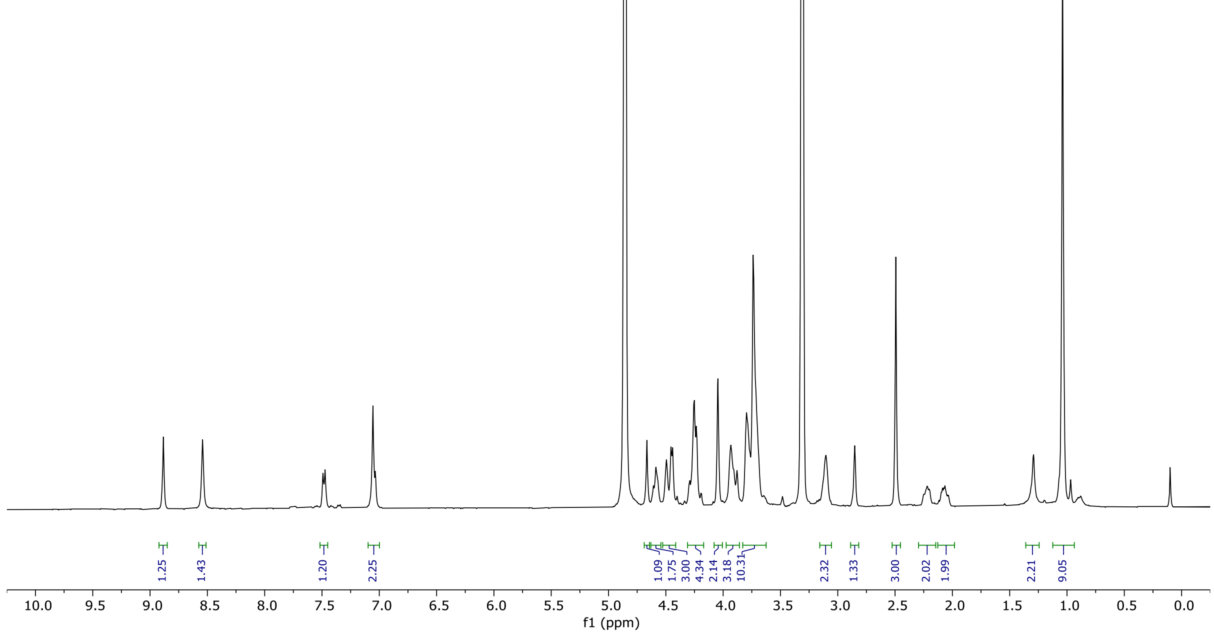
1.25
1.43
1.20
2.25
1.09
1.75
3.00
4.34
2.14
3.18
10.31
2.32
1.33
3.00
2.02
1.99
2.21
9.05
f1 (ppm)

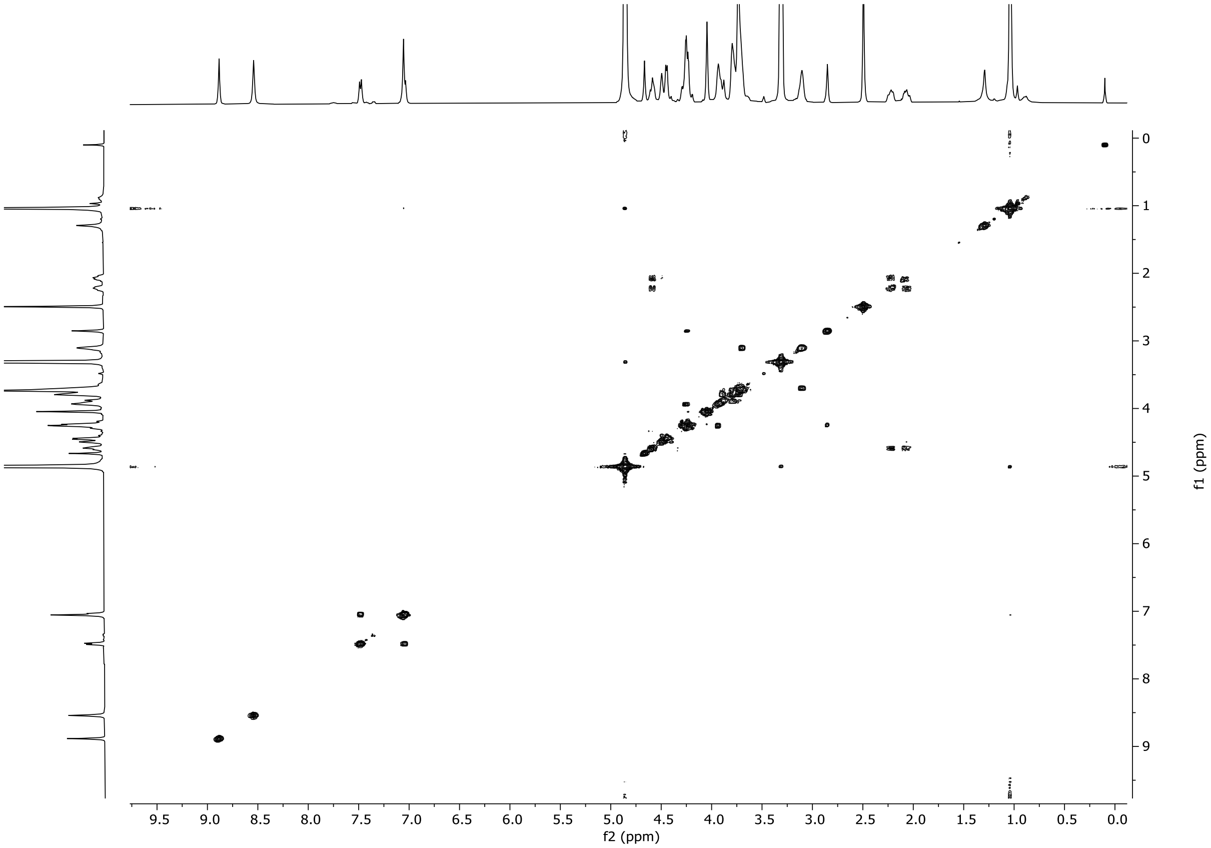
f2 (ppm)
f1 (ppm)

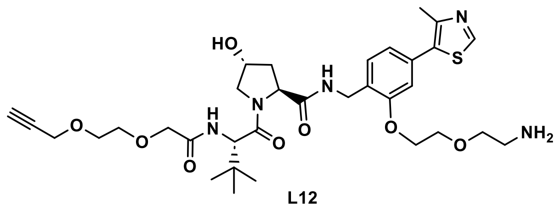

**L12**

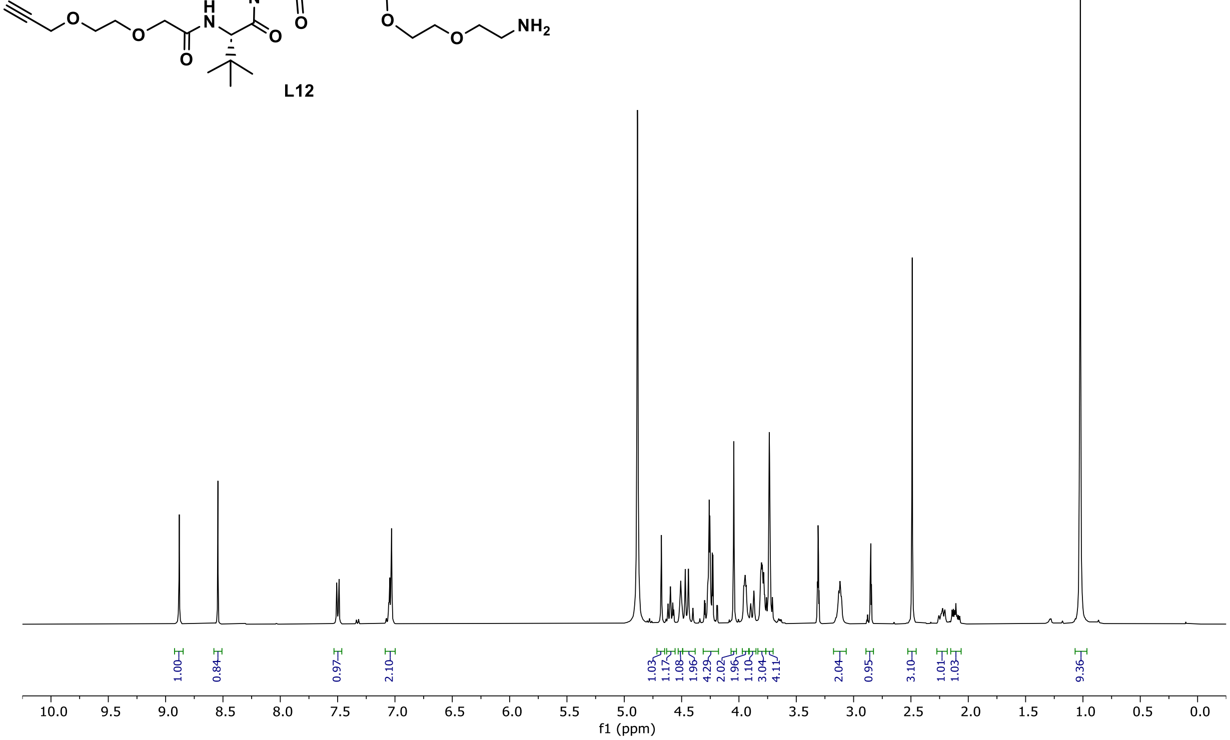
1.00
0.84
0.97
2.10
1.03
1.17
1.08
1.96
4.29
2.02
1.96
1.10
3.04
4.11
2.04
0.95
3.10
1.01
1.03
9.36
10.0 9.5 9.0 8.5 8.0 7.5 7.0 6.5 6.0 5.5 5.0 4.5 4.0 3.5 3.0 2.5 2.0 1.5 1.0 0.5 0.0
f1 (ppm)

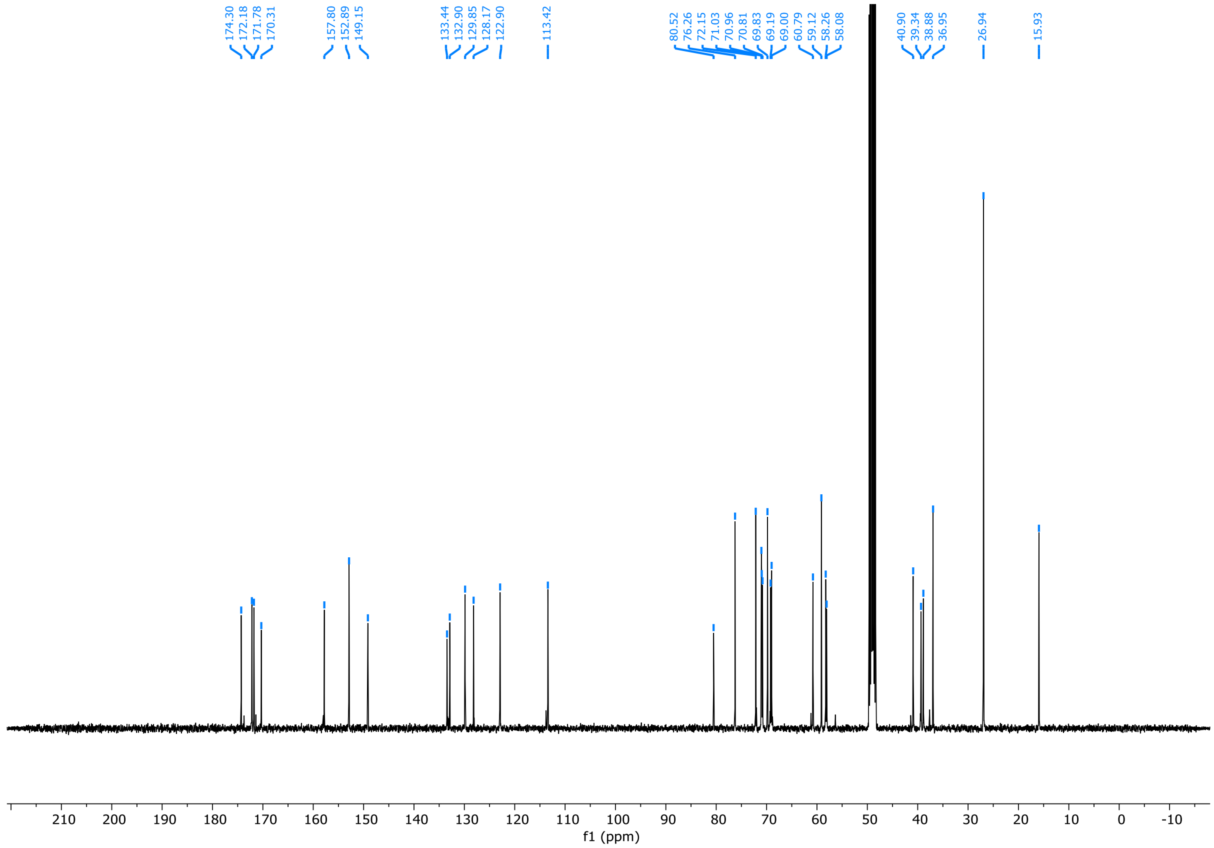
174.30
172.18
171.78
170.31
157.80
152.89
149.15
133.44
132.90
129.85
128.17
122.90
113.42
80.52
76.26
72.15
71.03
70.96
70.81
69.83
69.19
69.00
60.79
59.12
58.26
58.08
40.90
39.34
38.88
36.95
26.94
15.93
210 200 190 180 170 160 150 140 130 120 110 100 90 80 70 60 50 40 30 20 10 0 -10
f1 (ppm)

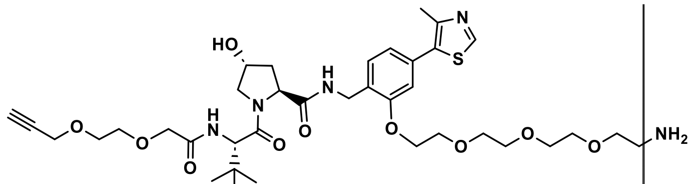
HO
N
H
N
O
O
O
N
H
O
O
N
S
O
O
O
O
NH2

**L13**

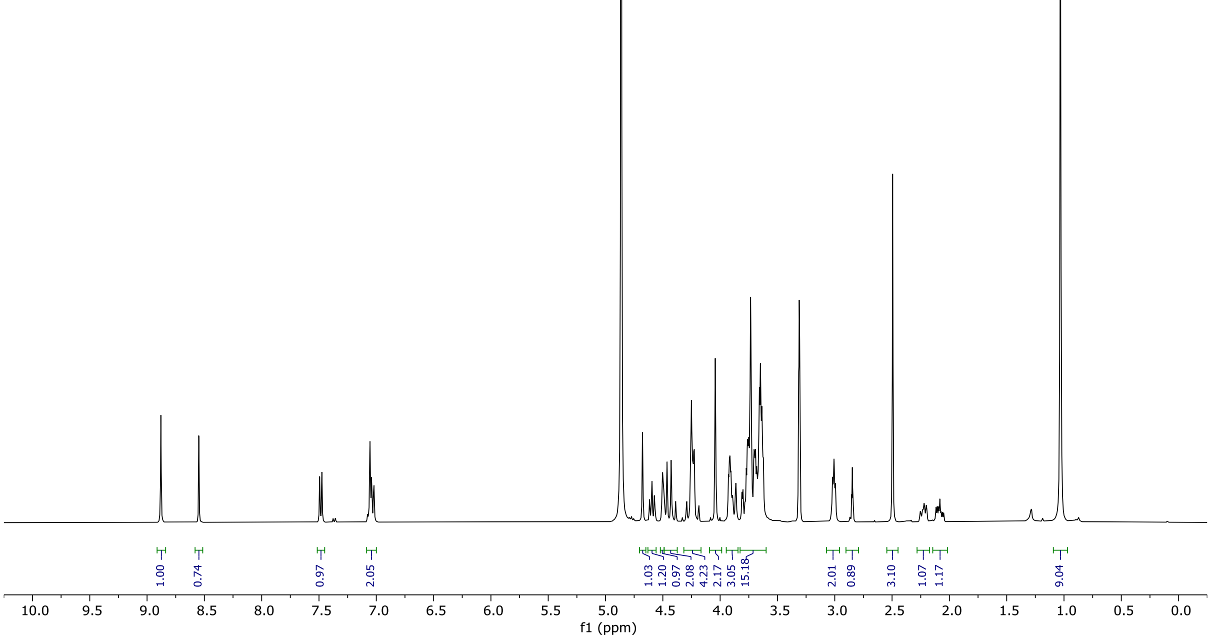
1.00
0.74
0.97
2.05
1.03
1.20
0.97
2.08
4.23
2.17
3.05
15.18
2.01
0.89
3.10
1.07
1.17
9.04
10.0 9.5 9.0 8.5 8.0 7.5 7.0 6.5 6.0 5.5 5.0 4.5 4.0 3.5 3.0 2.5 2.0 1.5 1.0 0.5 0.0
f1 (ppm)

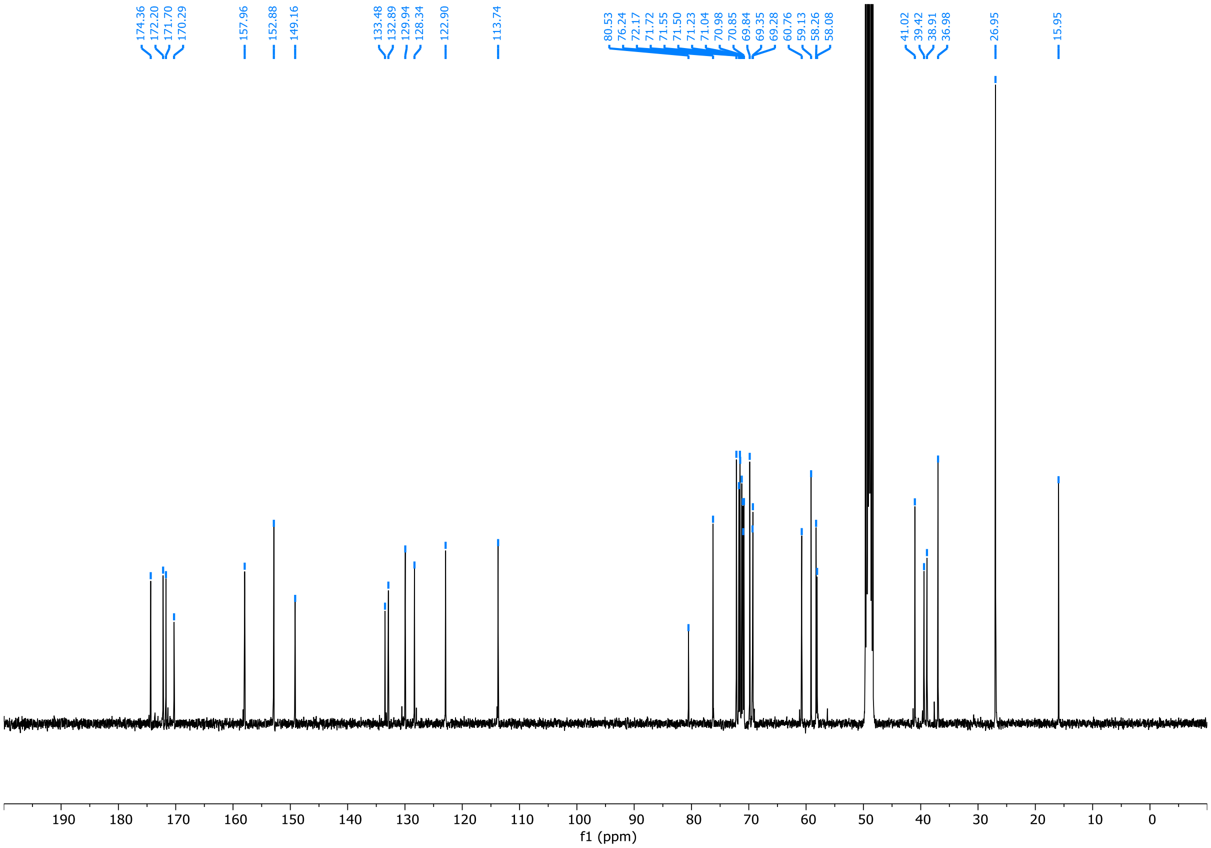
174.36
172.20
171.70
170.29
157.96
152.88
149.16
133.48
132.89
129.94
128.34
122.90
113.74
80.53
76.24
72.17
71.72
71.55
71.50
71.23
71.04
70.98
70.85
69.84
69.35
69.28
60.76
59.13
58.26
58.08
41.02
39.42
38.91
36.98
26.95
15.95
190 180 170 160 150 140 130 120 110 100 90 80 70 60 50 40 30 20 10 0
f1 (ppm)

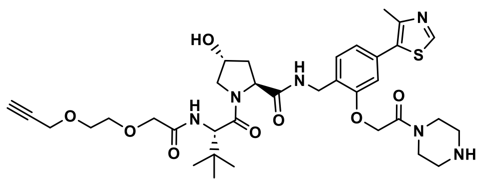

**L15**

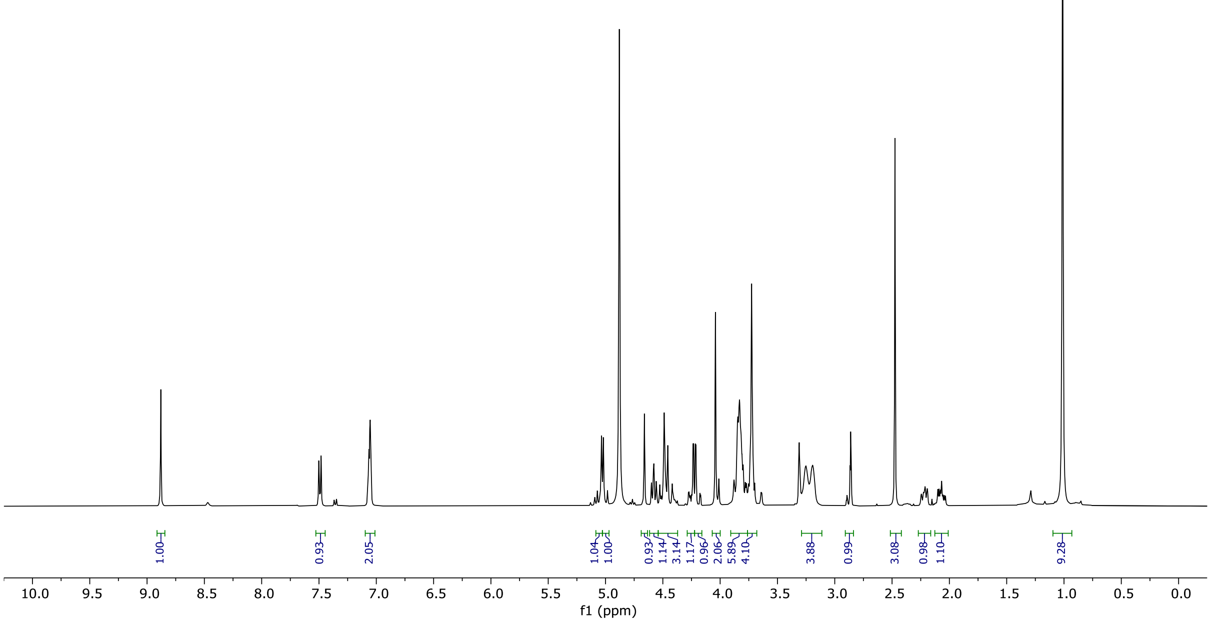
1.00
0.93
2.05
1.04
1.00
0.93
1.14
3.14
1.17
0.96
2.06
5.89
4.10
3.88
0.99
3.08
0.98
1.10
9.28
10.0 9.5 9.0 8.5 8.0 7.5 7.0 6.5 6.0 5.5 5.0 4.5 4.0 3.5 3.0 2.5 2.0 1.5 1.0 0.5 0.0
f1 (ppm)

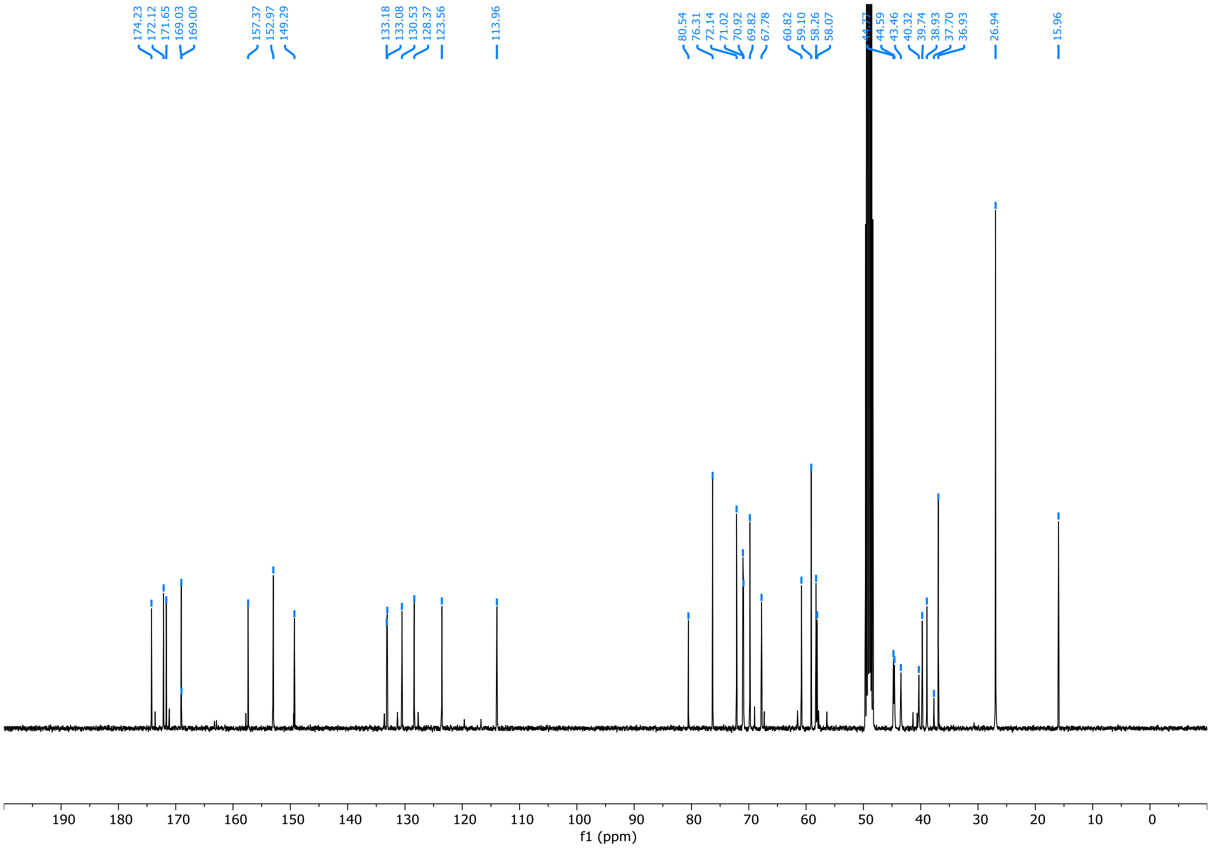
174.23
172.12
171.65
169.03
169.00
157.37
152.97
149.29
133.18
133.08
130.53
128.37
123.56
113.96
80.54
76.31
72.14
71.02
70.92
69.82
67.78
60.82
59.10
58.26
58.07
44.59
43.46
40.32
39.74
38.93
37.70
36.93
26.94
15.96
190 180 170 160 150 140 130 120 110 100 90 80 70 60 50 40 30 20 10 0
f1 (ppm)

### OFF-DNA COMPOUND SYNTHESIS

#### General information for compound synthesis

Unless otherwise noted, all reactions were performed in glass vials capped with silicon septa. Anhydrous dichloromethane (DCM), 1,2-dichloroethane (DCE), *N,N*-dimethylformamide (DMF), triethylamine ($Et_3N$), and *N,N*-Diisopropylethylamine (DIPEA) were purchased from Sigma Aldrich (with Sure/Seal™) and used as received. Other chemicals were purchased from various suppliers and used as received. Compound **INT-1** was synthesized according to the literature[2].

Thin layer chromatography (TLC) was performed using silica gel 60 $F_{254}$ glass plates (EMD Millipore) and visualized with UV light and/or staining with a solution of $KMnO_4$. Column chromatography was performed using Teledyne ISCO CombiFlash systems with RediSep Rf columns.

Ultra-performance liquid chromatography – mass spectrometry (UPLC-MS) was performed using Waters ACQUITY systems equipped with ACQUITY UPLC BEH C18 1.7 μM, 2.1 × 30 mm columns kept at 50°C. UV signals were monitored with ACQUITY UPLC PDA detectors (210 – 400 nm) and light scattering signals were recorded with ACQUITY ELSD detectors. Acidic gradients consisted of water (solvent A) and acetonitrile (solvent B) mobile phases both modified with 0.1% formic acid and basic gradients consisted of the same solvents modified with 5 mM ammonium hydroxide. Linear mobile phase gradients ran from 2% to 98% solvent B over a runtime of 2 mins and at a flowrate of 1.0 mL/min. High-resolution mass spectra (HRMS) were obtained with a Waters Xevo G2 QToF detector using positive mode electron spray ionization (ESI) and scan times of 0.2 sec.

Preparative high-performance liquid chromatography (prep-HPLC) was performed using a Waters system equipped with Waters 515 pumps, a Waters 2545 binary gradient module, a Waters ACQUITY QDa detector, a Waters 2998 PDA detector, and a Waters 2767 sample manager. Separations were performed with a Waters XBridge C18 OBD, 5 μM, 30 mm × 50 mm column using focused gradients of water (solvent A) and acetonitrile (solvent B) modified with 0.1% formic acid or 5 mM ammonium hydroxide (75 mL/min flowrate). Fraction collection was triggered using the UV signal and a user-defined target mass.

NMR Spectra were recorded on a Bruker AV-III spectrometer with $^{1}H$-NMR recorded at 400 MHz and $^{13}C$-NMR at 100 MHz or 151 MHz. Chemical shifts are expressed in ppm and referenced to residual solvent peaks with multiplicities reported as: s = singlet, d = doublet, t = triplet, q = quartet, m = multiplet, and br = broad signal. Coupling constants are reported in Hertz (Hz).

#### General synthetic scheme

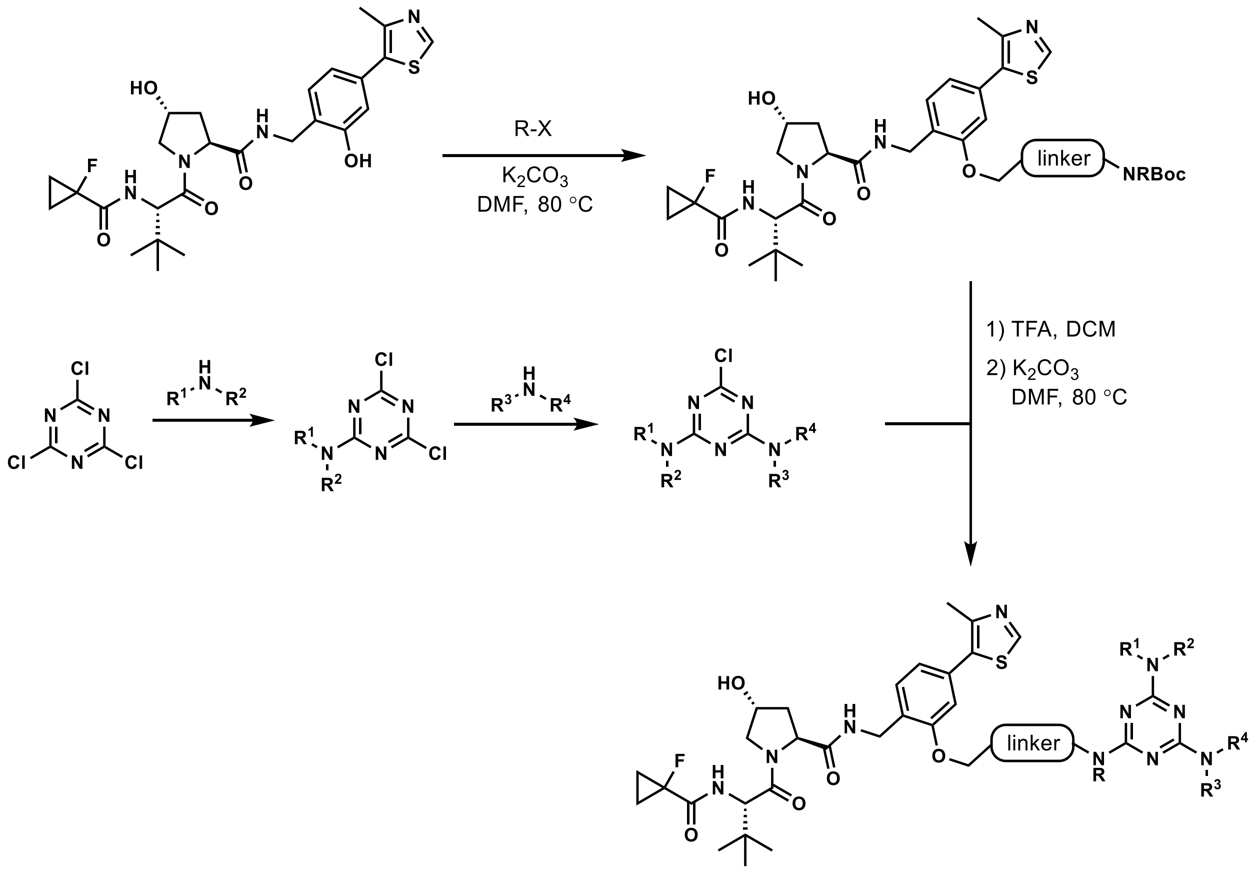

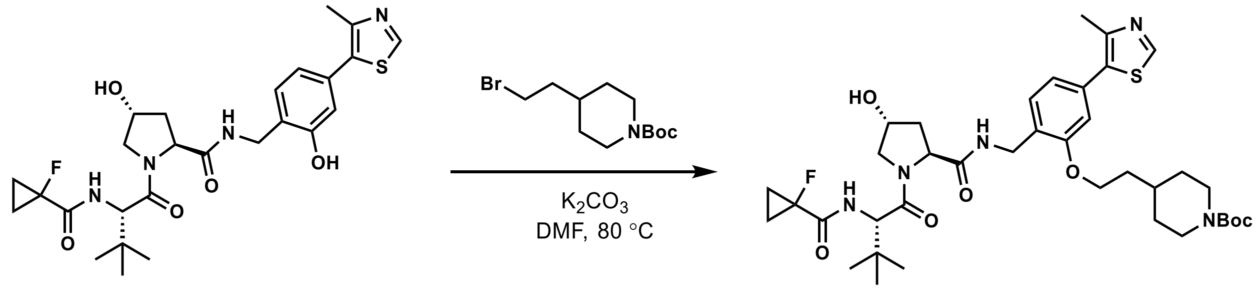

INT-1 INT-2

Compound **INT-2**: To a 1-dram glass vial was added compound **INT-1** (34.9 mg, 0.0655 mmol, 1 equiv.), *tert*-butyl 4-(2-bromoethyl)piperidine-1-carboxylate (48 mg, 0.16 mmol, 2.5 equiv.), $K_2CO_3$ (27 mg, 0.20 mmol, 3.0 equiv.) and DMF (0.8 mL). The reaction was heated at 80 °C for 12h. After cooling down to room temperature, the reaction was quenched with saturated aq. $NH_4Cl$ and extracted 3 times with DCM. The combined organic layer was dried over $Na_2SO_4$ and concentrated *in vacuo*. The residue was purified by flash column chromatography (0-10% MeOH/DCM) to afford compound **INT-2** (39.0 mg, 80% yield) as a pale yellow solid. $^1$H NMR (400 MHz, $CDCl_3$) δ 8.73 (s, 1H), 7.33 (d, *J* = 7.6 Hz, 1H), 7.27 (t, *J* = 5.8 Hz, 1H), 7.01 (dd, *J* = 8.7, 3.7 Hz, 1H), 6.94 (dd, *J* = 7.8, 1.7 Hz, 1H), 6.85 (d, *J* = 1.7 Hz, 1H), 4.72 (t, *J* = 7.6 Hz, 1H), 4.57 – 4.46 (m, 3H), 4.40 (dd, *J* = 14.9, 5.2 Hz, 1H), 4.17 – 4.02 (m, 4H), 3.98 (dt, *J* = 11.2, 2.0 Hz, 1H), 3.61 (dd, *J* = 11.2, 3.9 Hz, 1H), 2.71 (t, *J* = 12.4 Hz, 2H), 2.60 – 2.53 (m, 1H), 2.53 (s, 3H), 2.07 (ddt, *J* = 12.8, 8.2, 2.1 Hz, 1H), 1.81 (q, *J* = 6.3 Hz, 2H), 1.78 – 1.67 (m, 3H), 1.45 (s, 9H), 1.36 – 1.23 (m, 6H), 1.23 – 1.15 (m, 2H), 0.92 (s, 9H); $^{13}$C NMR (101 MHz, $CDCl_3$) δ 171.2, 170.5, 170.3, 157.0, 155.0, 150.6, 148.1, 132.2, 132.1, 129.6,

126.5, 121.7, 112.1, 79.5, 70.3, 65.8, 58.5, 57.6, 56.6, 55.2, 44.0, 43.2, 38.9, 35.9, 35.9, 35.4, 33.3, 32.2, 28.6, 26.4, 18.7, 17.4, 16.0, 13.9, 13.8; HRMS (ESI) *calcd* for $[C_{38}H_{55}FN_5O_7S]^+$ ($[M+H]^+$): *m/z* 744.3801, found: 744.3831.

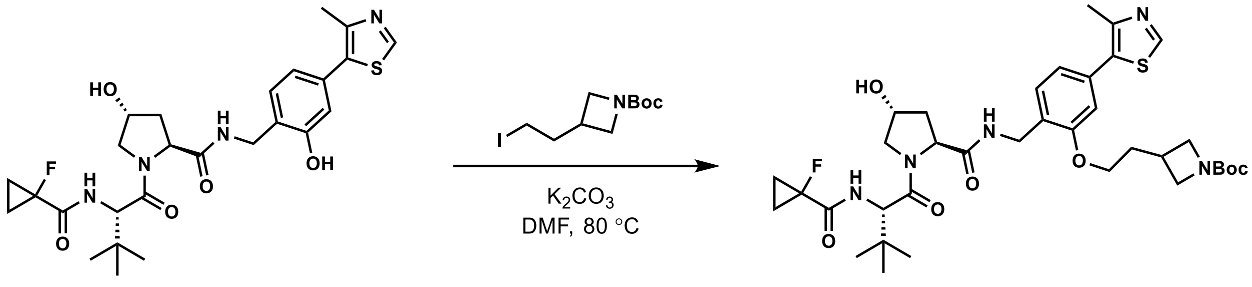

**INT-1** **INT-3**

Compound **INT-3**: To a 1-dram glass vial was added compound **INT-1** (25.3 mg, 0.0475 mmol, 1 equiv.), *tert*-butyl 3-(2-iodoethyl)azetidine-1-carboxylate (44 mg, 0.14 mmol, 3.0 equiv.), $K_2CO_3$ (26 mg, 0.19 mmol, 4.0 equiv.) and DMF (0.7 mL). The reaction was heated at 80 °C for 24h. After cooling down to room temperature, the reaction mixture was diluted with methanol (0.8 mL), filtered through 0.2 µm PTFE filter and purified by preparative HPLC (35-60% Acetonitrile/$H_2O$ with 0.1% formic acid, 3.5 min). The fractions containing pure desired product were combined and lyophilized to afford compound **INT-3** (12.8 mg, 37% yield) as a white solid. $^1H$ NMR (400 MHz, $CDCl_3$) δ 8.68 (s, 1H), 7.34 (d, *J* = 7.7 Hz, 1H), 7.29 – 7.24 (m, 1H), 7.08 – 7.00 (m, 1H), 6.96 (d, *J* = 7.7 Hz, 1H), 6.83 (d, *J* = 2.4 Hz, 1H), 4.77 – 4.65 (m, 1H), 4.61 – 4.46 (m, 3H), 4.45 – 4.32 (m, 1H), 4.04 – 3.86 (m, 3H), 3.74 – 3.16 (m, 6H), 2.77 (hept, *J* = 7.0 Hz, 1H), 2.62 – 2.51 (m, 1H), 2.51 (s, 3H), 2.21 – 2.03 (m, 2H), 1.88 – 1.75 (m, 1H), 1.49 – 1.43 (m, 3H), 1.36 – 1.23 (m, 4H), 0.97 – 0.87 (m, 9H); $^{13}C$ NMR (101 MHz, $CDCl_3$) δ 171.0, 170.4, 170.2, 156.5, 154.6, 150.4, 150.3, 148.6, 132.4, 132.3, 131.7, 129.9, 129.4, 126.2, 121.9, 112.0, 79.4, 70.2, 69.6, 66.1, 58.7, 58.5, 57.4, 56.5, 56.3, 48.9, 45.4, 38.7, 37.8, 35.8, 35.3, 33.9, 28.5, 26.3, 16.1, 13.8, 13.7; HRMS (ESI) *calcd* for $[C_{36}H_{51}FN_5O_7S]^+$ ($[M+H]^+$): *m/z* 716.3488, found: 716.3517.

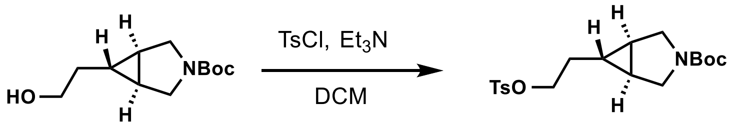

**INT-4** **INT-5**

Compound **INT-5**: To a 1-dram glass vial was added sequentially *tert*-butyl (1R,5S,6s)-6-(2-hydroxyethyl)-3-azabicyclo[3.1.0]hexane-3-carboxylate (42.8 mg, 0.188 mmol, 1 equiv.), 4-toluenesulfonyl chloride (72 mg, 0.38 mmol, 2.0 Eq), DCM (2 mL) and $Et_3N$ (79 µL, 0.56 mmol, 3.0 equiv). The reaction was allowed to stir at room temperature for 40h. The mixture was concentrated *in vacuo* and directly loaded on silica gel and purified by flash column chromatography (20-40% EtOAc/Heptane) to afford compound **INT-5** (67 mg, 93% yield) as a colorless oil. $^1H$ NMR (400 MHz, $CDCl_3$) δ 7.82 – 7.76 (m, 2H), 7.39 – 7.33 (m, 2H), 4.05 (t, *J* = 6.5 Hz, 2H), 3.46 (d, *J* = 10.8 Hz, 2H), 3.28 (dt, *J* = 10.8, 2.0 Hz, 2H), 2.46 (s, 3H), 1.60 (q, *J* = 6.8 Hz, 2H), 1.43 (s, 9H), 1.29 – 1.22 (m, 2H), 0.53 (tt, *J* = 6.9, 3.3 Hz, 1H); $^{13}C$ NMR (101 MHz, $CDCl_3$) δ 155.0, 145.0, 133.2, 130.0, 128.0, 79.4, 70.2, 48.3, 48.1, 31.1, 28.6, 22.8, 22.1, 21.8, 19.2; HRMS (ESI) *calcd* for $[C_{19}H_{27}NNaO_5S]^+$ ($[M+Na]^+$): *m/z* 404.1502, found: 404.1523.

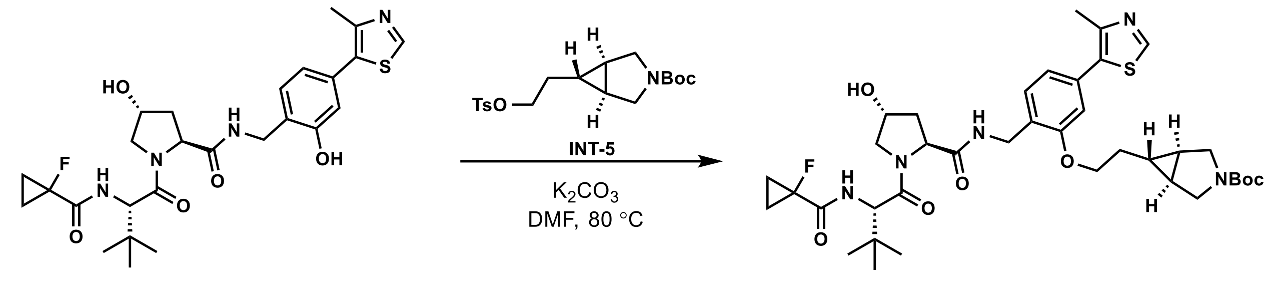

INT-1 INT-6

Compound **INT-6**: To a 1-dram glass vial was added compound **INT-1** (39.3 mg, 0.0738 mmol, 1 equiv.), *tert*-butyl (1R,5S,6s)-6-(2-(tosyloxy)ethyl)-3-azabicyclo[3.1.0]hexane-3-carboxylate (67 mg, 0.18 mmol, 2.4 equiv.), $K_2CO_3$ (31 mg, 0.22 mmol, 3.0 equiv.) and DMF (0.9 mL). The reaction was heated at 80 °C for 12h. After cooling down to room temperature, the reaction was quenched with saturated aq. $NH_4Cl$ and extracted 3 times with DCM. The combined organic layer was dried over $Na_2SO_4$ and concentrated *in vacuo*. The residue was purified by flash column chromatography (4-10% MeOH/DCM) to afford compound **INT-6** (42 mg, 77% yield) as a colorless oil. $^1H$ NMR (400 MHz, $CDCl_3$) δ 8.69 (s, 1H), 7.33 (d, *J* = 7.7 Hz, 1H), 7.25 – 7.16 (m, 1H), 7.03 (dd, *J* = 8.8, 3.5 Hz, 1H), 6.95 (d, *J* = 7.5 Hz, 1H), 6.86 (s, 1H), 4.69 (t, *J* = 7.8 Hz, 1H), 4.60 – 4.35 (m, 4H), 4.06 (t, *J* = 6.6 Hz, 2H), 3.95 (d, *J* = 11.1 Hz, 1H), 3.60 (td, *J* = 11.1, 4.2 Hz, 2H), 3.52 (d, *J* = 10.8 Hz, 1H), 3.34 (t, *J* = 10.1 Hz, 2H), 2.58 – 2.49 (m, 1H), 2.53 (s, 3H), 2.15 – 1.99 (m, 1H), 1.90 – 1.73 m, 2H), 1.42 (s, 9H), 1.37 (t, *J* = 3.5 Hz, 2H), 1.34 – 1.28 (m, 4H), 0.92 (d, *J* = 2.2 Hz, 9H), 0.73 – 0.69 (m, 1H); $^{13}C$ NMR (101 MHz, $CDCl_3$) δ 171.1, 170.5, 170.3, 170.1, 156.9, 155.2, 150.5, 148.4, 132.4, 132.0, 129.77, 129.73, 126.3, 121.7, 112.1, 79.6, 70.3, 67.9, 58.6, 57.5, 56.6, 55.8, 48.5, 48.3, 43.8, 36.0, 35.6, 28.6, 26.4, 20.0, 18.8, 17.4, 16.2; HRMS (ESI) *calcd* for $[C_{38}H_{53}FN_5O_7S]^+$ ($[M+H]^+$): *m/z* 742.3644, found: 742.3658.

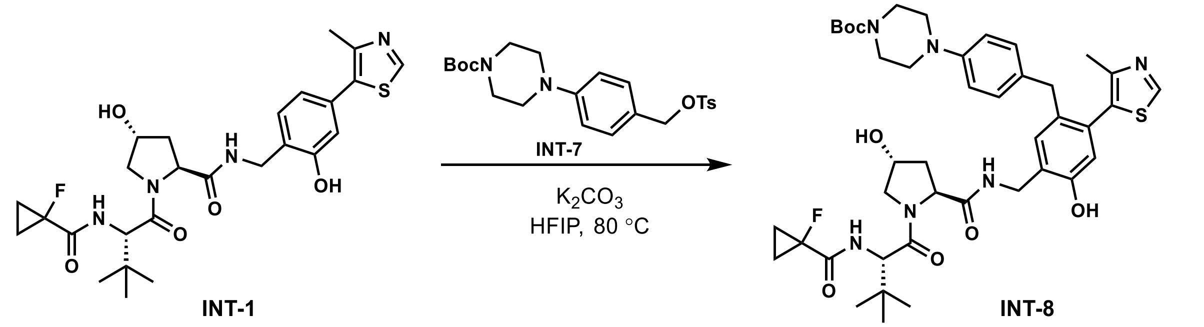

INT-1 INT-8

Compound **INT-8**: To a 1-dram glass vial was added *tert*-butyl 4-(4-(hydroxymethyl)phenyl)piperazine-1-carboxylate (38.9 mg, 0.133 mmol, 1 equiv.), TsCl (20.3 mg, 0.159 mmol, 0.80 equiv.), DCE (0.8 mL) and $Et_3N$ (55 µL, 0.40 mmol, 3.0 equiv.). The mixture was stirred at room temperature for 48h, when TLC showed full consumption of TsCl. The reaction was concentrated *in vacuo* and the crude compound **INT-7** was directly used in the next step.

To a 1-dram glass vial was added compound **INT-1** (24.0 mg, 0.0451 mmol, 1 equiv.), the crude tosylate **INT-7** prepared above (3.0 equiv.), $K_2CO_3$ (25 mg, 0.18 mmol, 4.0 equiv.) and HFIP (0.9 mL). The reaction was heated at 80 °C for 4h. After cooling down to room temperature, the reaction was quenched with saturated aq. $NH_4Cl$ and extracted 3 times with DCM. The combined organic layer was dried over $Na_2SO_4$ and concentrated *in vacuo*. The residue was purified by reverse-phase preparative HPLC (35%-60% acetonitrile/water with 0.1% formic acid) to afford the product **INT-8** (13.7 mg, 36% yield) as white powder. $^1H$ NMR (400 MHz, $CDCl_3$) δ 8.97 (s, 1H), 8.68 (s, 1H), 7.98 – 7.85 (t, *J* = 6.2 Hz, 1H), 6.96 (dd, *J* = 8.6, 3.6 Hz, 1H), 6.92 (s, 1H), 6.85 – 6.79 (m, 4H), 6.78 (s, 1H), 4.75 (t, *J* = 7.8 Hz, 1H), 4.57 – 4.51 (m, 1H), 4.45 (q, *J* = 7.0 Hz, 2H), 4.13 (dd, *J* = 14.5, 5.5 Hz, 1H), 4.01 (d, *J* = 11.4 Hz, 1H), 3.72 – 3.61 (m, 2H), 3.61 – 3.51 (m, 5H), 3.07 (brs, 4H), 2.57 (ddd, *J* = 12.7, 7.4, 4.6 Hz, 1H), 2.13 (s, 3H), 2.06 (dd, *J* = 13.7, 8.3 Hz, 1H), 1.47 (s, 9H), 1.36 – 1.27 (m, 4H), 0.88 (s, 9H); $^{13}C$ NMR (101 MHz, $CDCl_3$) δ 172.7, 171.6, 170.7, 170.5, 154.8, 153.8, 151.2, 150.0, 132.8, 132.6, 132.5, 129.7, 129.6, 124.8, 121.0, 117.2, 80.1, 79.6, 70.3, 57.9, 57.8, 56.6, 50.1, 40.3, 37.8, 35.3, 35.2, 29.8, 28.6, 26.4, 15.5, 14.0, 13.9; HRMS (ESI) *calcd* for $[C_{42}H_{56}FN_6O_7S]^+$ ($[M+H]^+$): *m/z* 807.3910, found: 807.3949.

Note: while using acetone as solvent resulted in a mixture of O/C-alkylation and mono/di-alkylation products (figure S4), we found using HFIP as solvent led to the selective formation of desired C-alkylation product.

**General procedure for the double $S_NAr$ reactions of cyanuric chloride**

Cyanuric chloride (1.0 equiv.) was suspended in anhydrous DCM (0.1 M) in a 2-dram glass vial under positive pressure of nitrogen and was cooled to 0 °C. A mixture of the amine building block (0.9 equiv.) and DIPEA (3.0 – 6.0 equiv., depending on salt state of the amine) dissolved in a small amount of DCM was added dropwise. The reaction was allowed to stir at 0 °C for 1 h. After completion of the reaction, the mixture was directly loaded onto silica gel and eluted with 0 – 10% MeOH/DCM to afford the crude mono-$S_NAr$ intermediate.

The mono-$S_NAr$ intermediate (1.0 equiv.) and the amine building block for the $2^{nd}$ $S_NAr$ reaction (1.2 equiv.) was mixed and dissolved in anhydrous DCM (0.1 M) in a 1-dram glass vial at room temperature. DIPEA (3.0 – 6.0 equiv., depending on salt state of the amine) was added dropwise and the resulting mixture was stirred at room temperature. Upon the completion of the reaction as indicated by UPLC-MS (reaction time varies from 1 h to 15 h), the reaction was concentrated *in vacuo* and then re-suspended in ethyl acetate (0.5 mL). The insoluble material was removed, and the resulting solution was concentrated and purified by flash column chromatography (typically 5 – 20% MeOH/DCM) to provide the double-$S_NAr$ product.

**General procedure for the final $S_NAr$ reactions of triazine**

To the Boc-protected linker-attached VHL ligand **INT-2, INT-3, INT-6** or **INT-8** (1.0 equiv) in a 1-dram glass vial was added DCM and TFA (v/v = 4:1, 0.02 M). After stirring at room temperature for 2 h, the reaction mixture was concentrated by rotary evaporation and further dried in high vacuum for 3 h. The resulting amine TFA salt was used directly in the next step without further purification.

To the same vial containing the crude amine was added the chlorotriazine (1.5 – 2.5 equiv.) and potassium carbonate (5.0 equiv), followed by anhydrous DMF (0.4 mL, ~ 0.03 M). The resulting mixture was heated at 80 °C for 12 h. The reaction was then cooled to room temperature, quenched with aq. $NH_4Cl$ (1 mL) and extracted with DCM (3 × 1.5 mL). The combined organic layers were concentrated *in vacuo* and the residue was purified by reverse-phase preparative HPLC (water-acetonitrile with 0.1% formic acid) to afford the final product as white powder.

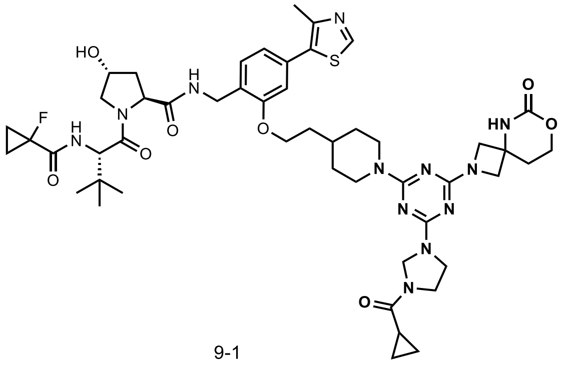

9-1

**Compound 9-1**: $^1$H NMR (400 MHz, $CDCl_3$) δ 8.69 (s, 1H), 7.32 (d, *J* = 7.8 Hz, 1H), 7.30 – 7.25 (m, 1H), 7.02 (dt, *J* = 8.2, 3.6 Hz, 1H), 6.95 (dd, *J* = 7.5, 1.6 Hz, 1H), 6.86 (s, 1H), 6.35 – 6.05 (m, 1H), 5.14 (s, 1H), 4.99 (s, 1H), 4.82 – 4.66 (m, 3H), 4.58 – 4.45 (m, 3H), 4.40 (dd, *J* = 14.9, 5.2 Hz, 1H), 4.30 (t, *J* = 5.3 Hz, 2H), 4.15 – 3.96 (m, 7H), 3.90 (s, 2H), 3.77 (s, 2H), 3.62 (dd, *J* = 11.2, 4.0 Hz, 1H), 2.85 – 2.70 (m, 2H), 2.62 – 2.52 (m, 1H), 2.52 (s, 3H), 2.28 – 2.19 (m, 2H), 2.07 (ddt, *J* = 13.0, 8.5, 2.0 Hz, 1H), 1.91 – 1.50 (m, 6H), 1.39 – 1.17 (m, 6H), 1.10 – 1.01 (m, 2H), 0.93 (s, 9H), 0.87 – 0.79 (m, 2H); $^{13}$C NMR (101 MHz, $CDCl_3$) δ 171.7, 171.4, 171.2, 170.5, 164.3, 156.8, 153.0, 150.5, 148.6, 132.4, 131.9, 129.7, 129.6, 126.3, 121.7, 112.1, 79.6, 70.3, 66.0, 65.8, 64.0, 62.6, 61.0, 60.6, 58.6, 57.6, 56.6, 51.4, 44.8, 44.2, 43.8, 43.5, 43.2, 38.9, 35.9, 35.4, 35.2, 33.9, 33.7, 32.2, 31.6, 26.5, 16.3, 13.9, 13.8, 12.8, 12.4, 8.1, 8.0, 1.3; HRMS (ESI) *calcd* for $[C_{49}H_{66}FN_{12}O_8S]^+$ ($[M+H]^+$): *m/z* 1001.4826, found: 1001.4841.

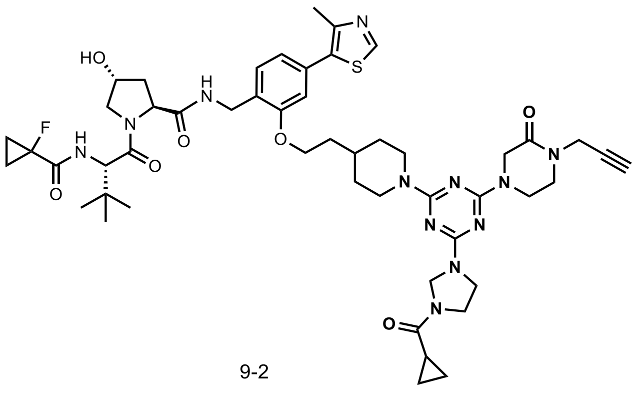

9-2

**Compound 9-2**: $^1$H NMR (400 MHz, $CDCl_3$) δ 8.69 (s, 1H), 7.33 (d, *J* = 7.7 Hz, 1H), 7.30 – 7.26 (m, 1H), 7.02 – 6.93 (m, 2H), 6.88 (s, 1H), 5.16 (s, 1H), 5.01 (s, 1H), 4.82 – 4.69 (m, 3H), 4.59 – 4.53 (m, 1H), 4.53 – 4.46 (m, 2H), 4.46 – 4.38 (m, 3H), 4.30 (s, 2H), 4.14 – 3.88 (m, 7H), 3.79 (s, 2H), 3.61 (dd, *J* = 10.8, 4.0 Hz, 1H), 3.52 (q, *J* = 5.9 Hz, 2H), 2.87 – 2.72 (m, 2H), 2.65 – 2.56 (m, 1H), 2.53 (s, 3H), 2.24 (t, *J* = 2.8 Hz, 1H), 2.13 – 2.04 (m, 1H), 1.92 – 1.76 (m, 5H), 1.69 – 1.55 (m, 1H), 1.38 – 1.14 (m, 6H), 1.11 – 1.00 (m, 2H), 0.92 (s, 9H), 0.89 – 0.78 (m, 2H); $^{13}$C NMR (101 MHz, $CDCl_3$) δ 171.7, 171.4, 171.3, 171.2, 170.4, 166.4, 164.3, 163.2, 156.94, 156.89, 150.4, 148.6, 132.5, 132.0, 129.9, 129.7, 126.4, 121.7, 112.21, 112.15, 79.6, 77.9, 77.7, 72.7, 70.3, 66.1, 65.9, 61.1, 60.7, 58.5, 58.4, 57.7, 57.6, 56.5, 47.7, 45.7, 44.8, 44.4, 43.8, 43.6, 43.2, 40.0, 37.0, 36.0, 35.81, 35.75, 35.6, 35.3, 35.2, 34.0, 33.7, 32.3, 29.8, 26.4, 16.3, 13.9, 13.8, 12.9, 12.4, 8.1, 8.0, 1.3; HRMS (ESI) *calcd* for $[C_{50}H_{65}FN_{12}O_7SNa]^+$ ($[M+Na]^+$): *m/z* 1019.4696, found: 1019.4724.

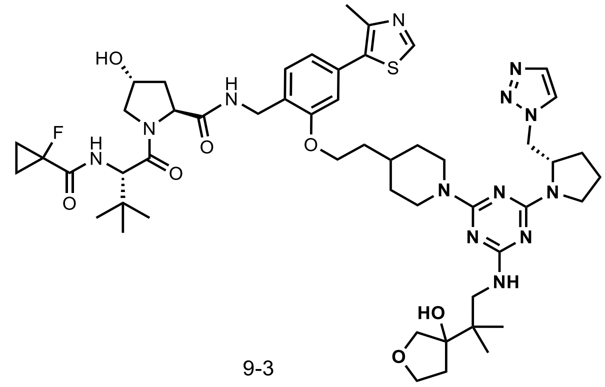

9-3

**Compound 9-3**: $^1$H NMR (400 MHz, $CDCl_3$) δ 8.68 (s, 1H), 7.68 (s, 1H), 7.42 (s, 1H), 7.33 (d, *J* = 7.7 Hz, 1H), 7.26 – 7.20 (m, 1H), 7.01 (dd, *J* = 9.0, 4.0 Hz, 1H), 6.96 (dd, *J* = 7.8, 1.6 Hz, 1H), 6.87 (d, *J* = 1.6 Hz, 1H), 5.18 (br, 1H), 4.89 – 4.59 (m, 5H), 4.58 – 4.47 (m, 4H), 4.42 (dd, *J* = 14.8, 5.3 Hz, 1H), 4.09 (dt, *J* = 7.3, 3.6 Hz, 2H), 4.04 – 3.77 (m, 4H), 3.62 (dd, *J* = 11.3, 3.9 Hz, 1H), 3.59 – 3.08 (m, 5H), 2.83 (br, 2H), 2.62 – 2.52 (m, 1H), 2.53 (s, 3H), 2.15 – 2.05 (m, 1H), 2.05 – 1.78 (m, 12H), 1.35 – 1.24 (m, 5H), 1.04 (s, 3H), 1.00 (s, 3H), 0.93 (s, 9H); $^{13}$C NMR (101 MHz, DMSO) δ 171.9, 169.0, 168.2, 168.0, 166.2, 163.8, 155.9, 151.5, 147.9, 133.3, 131.4, 130.9, 127.7, 126.9, 125.4, 120.8, 111.7, 84.4, 84.1, 79.3, 77.0, 75.1, 69.0, 66.8, 65.5, 58.8, 56.7, 56.6, 56.4, 50.8, 50.6, 47.3, 46.4, 42.9, 40.2, 39.9, 39.7, 39.5, 39.3, 39.1, 38.9, 37.9, 37.3, 36.1, 35.4, 32.9, 31.7, 27.7, 26.2, 22.1, 21.9, 16.0, 13.1, 13.0, 12.8, 12.7; HRMS (ESI) *calcd* for $[C_{51}H_{72}FN_{13}O_7SNa]^+$ ($[M+Na]^+$): *m/z* 1052.5275, found: 1052.5316.

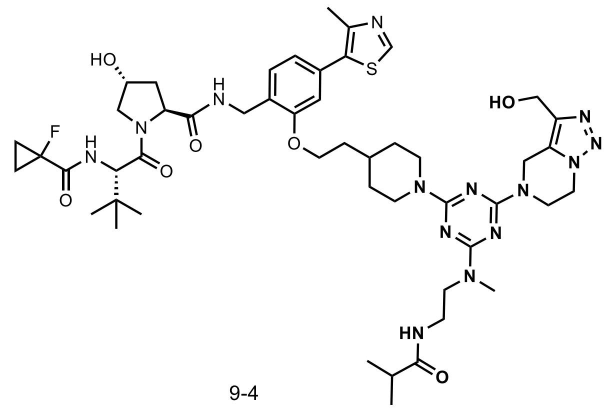

9-4

**Compound 9-4**: $^1$H NMR (400 MHz, $CDCl_3$) δ 8.69 (s, 1H), 7.33 (d, *J* = 7.7 Hz, 1H), 7.30 – 7.26 (m, 1H), 7.00 (dd, *J* = 9.0, 3.4 Hz, 1H), 6.96 (dd, *J* = 7.7, 1.5 Hz, 1H), 6.87 (d, *J* = 1.6 Hz, 1H), 5.39 – 5.18 (br, 1H), 5.18 – 5.01 (br, 1H), 4.95 – 4.66 (m, 5H), 4.61 – 4.48 (m, 3H), 4.48 – 4.27 (m, 3H), 4.34 – 4.21 (m, 2H), 4.10 (tt, *J* = 5.9, 2.9 Hz, 2H), 3.99 (d, *J* = 11.2 Hz, 1H), 3.72 (br, 2H), 3.62 (dd, *J* = 11.2, 3.9 Hz, 1H), 3.46 (br, 2H), 3.15 (s, 3H), 2.87 (t, *J* = 12.7 Hz, 2H), 2.63 – 2.53 (m, 1H), 2.53 (s, 3H), 2.25 (sept, *J* = 6.7 Hz, 1H), 2.15 – 2.04 (m, 1H), 1.98 – 1.77 (m, 5H), 1.37 – 1.26 (m, 6H), 1.03 (br, 6H), 0.93 (s, 9H); $^{13}$C NMR (101 MHz, $CDCl_3$) δ 171.2, 170.6, 170.5, 170.3, 156.8, 150.5, 148.6, 132.4, 131.9, 129.6, 129.1, 126.3, 121.8, 112.1, 79.6, 70.3, 65.9, 58.6, 57.6, 56.6, 48.0, 45.9, 44.1, 40.2, 39.9, 39.0, 37.2, 35.90, 35.86, 35.7, 35.3, 35.1, 33.7, 32.3, 32.2, 29.8, 26.4, 19.6, 16.3, 13.9, 13.8; HRMS (ESI) *calcd* for $[C_{49}H_{70}FN_{14}O_7S]^+$ ($[M+H]^+$): *m/z* 1017.5251, found: 1017.5261.

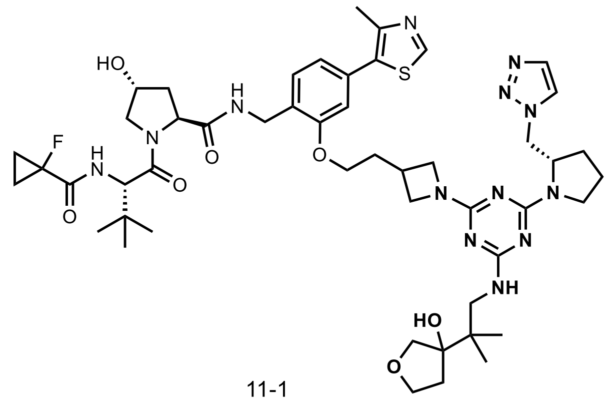

11-1

**Compound 11-1**: $^1$H NMR (400 MHz, $CDCl_3$) δ 8.67 (s, 1H), 7.68 (s, 1H), 7.43 (s, 1H), 7.35 (d, *J* = 6.4 Hz, 1H), 7.26 – 7.15 (m, 1H), 7.06 – 6.99 (m, 1H), 6.97 (d, *J* = 7.7 Hz, 1H), 6.85 (s, 1H), 5.22 – 4.93 (m, 1H), 4.84 – 4.35 (m, 8H), 4.12 – 3.37 (m, 16H), 2.71 – 2.53 (m, 1H), 2.52 (s, 3H), 2.26 – 1.68 (m, 10H), 1.37 – 1.26 (m, 3H), 1.04 (s, 3H), 1.00 (s, 3H), 0.93 (br, 9H). $^{13}$C NMR (101 MHz, $CDCl_3$) δ 171.0, 170.7, 170.4, 170.1, 166.0, 163.1, 156.6, 154.8, 150.5, 148.7, 134.0, 132.5, 131.8, 129.7, 126.3, 124.4, 122.1, 112.1, 107.4, 84.2, 75.8, 70.3, 69.6, 67.4, 58.6, 57.5, 56.6, 54.4, 51.4, 49.3, 47.0, 45.8, 40.7, 38.9, 38.2, 37.9, 36.2, 35.6, 31.5, 28.2, 26.4, 23.1, 22.6, 16.3, 13.9, 13.8. HRMS (ESI) *calcd* for $[C_{49}H_{69}FN_{13}O_7S]^+$ ($[M+H]^+$): *m/z* 1002.5142, found: 1002.5151.

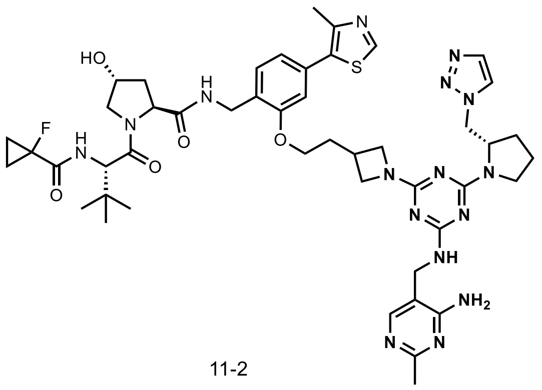

11-2

**Compound 11-2**: $^{1}$H NMR (400 MHz, DMSO) δ 8.97 (s, 1H), 8.63 – 8.45 (m, 1H), 8.10 – 7.84 (m, 2H), 7.71 (s, 1H), 7.42 (d, *J* = 7.8 Hz, 1H), 7.30 (dd, *J* = 9.3, 2.8 Hz, 1H), 7.23 – 7.07 (m, 1H), 7.07 – 7.00 (m, 1H), 7.00 – 6.67 (m, 3H), 5.20 (s, 1H), 4.81 – 4.48 (m, 4H), 4.43 – 4.26 (m, 4H), 4.26 – 4.00 (m, 6H), 3.84 – 3.57 (m, 6H), 2.83 – 2.64 (m, 1H), 2.45 (s, 3H), 2.27 (s, 3H), 2.25 – 2.20 (m, 1H), 2.19 – 2.00 (m, 2H), 1.96 – 1.64 (m, 5H), 1.41 – 1.31 (m, 2H), 1.21 (dd, *J* = 8.4, 3.4 Hz, 2H), 0.94 (s, 9H); $^{13}$C NMR (101 MHz, DMSO) δ 171.9, 169.0, 168.3, 168.1, 165.4, 165.2, 163.4, 163.2, 163.0, 162.8, 161.8, 155.8, 155.2, 151.5, 148.0, 133.4, 131.3, 131.0, 127.8, 127.1, 126.9, 125.4, 121.0, 111.8, 111.7, 79.3, 77.0, 69.5, 69.0, 58.9, 56.8, 56.6, 50.8, 48.4, 46.4, 45.1, 38.0, 37.9, 37.7, 37.4, 36.1, 27.8, 26.2, 25.2, 22.3, 16.0, 13.1, 13.0, 12.8, 12.7; HRMS (ESI) *calcd* for $[C_{47}H_{62}FN_{16}O_5S]^+$ ($[M+H]^+$): *m/z* 981.4788, found: 981.4816.

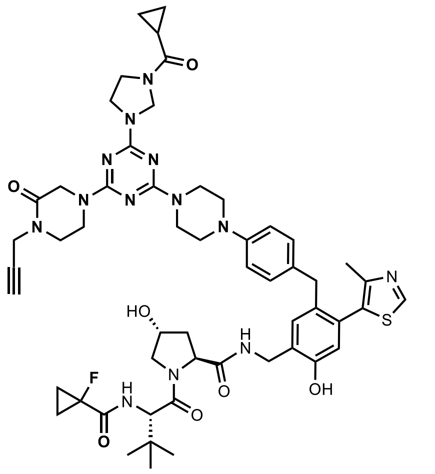

12-1

**Compound 12-1**: $^{1}$H NMR (400 MHz, DMSO) δ 9.67 (brs, 1H), 8.97 (s, 1H), 8.48 (t, *J* = 5.9 Hz, 1H), 7.23 (dd, *J* = 9.0, 2.8 Hz, 1H), 7.20 (s, 1H), 6.79 – 6.69 (m, 4H), 6.63 (s, 1H), 5.17 (s, 1H), 5.13 (s, 1H), 4.81 (s, 1H), 4.57 (d, *J* = 9.5 Hz, 1H), 4.47 (t, *J* = 8.3 Hz, 1H), 4.36 – 4.24 (m, 3H), 4.23 – 4.16 (m, 4H), 4.02 – 3.89 (m, 4H), 3.88 – 3.75 (m, 5H), 3.65 – 3.58 (m, 4H), 3.45 – 3.40 (m, 2H), 3.22 (t, *J* = 2.5 Hz, 1H), 3.04 (brs, 4H), 2.06 – 1.97 (m, 1H), 2.03 (s, 3H), 1.87 – 1.73 (m, 2H), 1.40 – 1.30 (m, 2H), 1.24 – 1.17 (m, 2H), 0.94 (s, 9H), 0.81 – 0.74 (m, 4H); $^{13}$C NMR (101 MHz, DMSO) δ 171.8, 170.7, 170.2, 168.8, 168.2, 168.0, 165.1, 164.2, 163.8, 162.8, 152.7, 152.1, 149.1, 132.1, 131.4, 129.8, 129.7, 129.2, 128.8, 126.3, 117.3, 115.9, 79.4, 79.0, 77.1, 74.6, 69.0, 60.4, 60.2, 58.8, 56.8, 56.6, 48.7, 47.0, 45.0, 44.1, 44.0, 43.3, 42.8, 42.6, 37.9, 37.1, 36.2, 34.7, 26.2, 15.1, 13.1, 13.0, 12.8, 12.7, 12.2, 11.7, 7.4, 7.3; HRMS (ESI) *calcd* for $[C_{54}H_{67}FN_{13}O_7S]^+$ ($[M+H]^+$): *m/z* 1060.4986, found: 1060.4965.

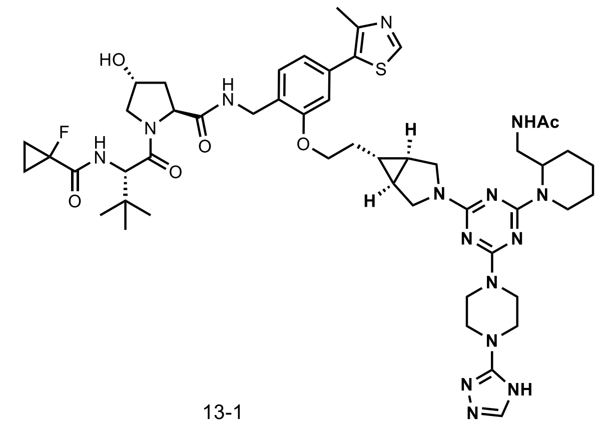

13-1

**Compound 13-1**: $^{1}$H NMR (400 MHz, $CDCl_3$) δ 8.68 (s, 1H), 7.78 (s, 1H), 7.50 – 7.30 (m, 1H), 7.33 (d, *J* = 7.7 Hz, 1H), 7.06 (dd, *J* = 8.9, 3.5 Hz, 1H), 6.95 (dd, *J* = 7.7, 1.6 Hz, 1H), 6.86 (d, *J* = 1.7 Hz, 1H), 6.82 (br, 1H), 5.06 (br, 1H), 4.83 – 4.61 (m, 2H), 4.61 – 4.47 (m, 3H), 4.39 (ddd, *J* = 14.9, 5.1, 2.3 Hz, 1H), 4.15 – 4.01 (m, 2H), 4.00 – 3.57 (m, 10H), 3.56 – 3.35 (m, 6H), 3.26 (br, 1H), 2.85 (br, 1H), 2.85 – 2.46 (m, 1H), 2.52 (s, 3H), 2.16 – 1.99 (m, 1H), 1.96 – 1.76 (m, 1H), 1.79 (s, 3H), 1.73 – 1.56 (m, 5H), 1.51 – 1.39 (m, 3H), 1.36 – 1.22 (m, 5H), 0.94 (s, 9H), 0.76 – 0.67 (m, 1H); $^{13}$C NMR (101 MHz, $CDCl_3$) δ 171.2, 170.7, 170.4, 170.2, 162.9, 156.8, 150.5, 148.6, 146.0, 132.4, 131.9, 129.5, 126.3, 121.8, 112.2, 79.5, 70.2, 68.0, 58.7, 57.6, 56.6, 48.6, 46.8, 42.7, 41.3, 38.8, 38.4, 36.1, 35.6, 31.7, 29.8, 27.6, 26.5, 25.5, 23.4, 22.7, 20.7, 19.9, 16.3, 13.97, 13.90, 13.87, 13.80; HRMS (ESI) *calcd* for $[C_{50}H_{69}FN_{15}O_6S]^+$ ($[M+H]^+$): *m/z* 1026.5255, found: 1026.5302.

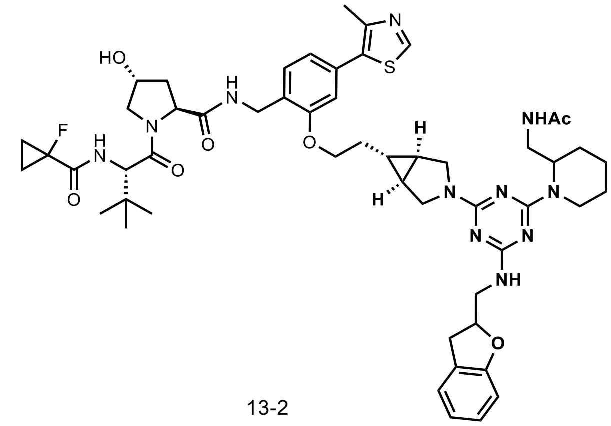

13-2

**Compound 13-2**: $^{1}$H NMR (400 MHz, DMSO) δ 8.97 (s, 1H), 8.50 (t, *J* = 5.4 Hz, 1H), 7.82 – 7.62 (br, 1H), 7.41 (d, *J* = 7.8 Hz, 1H), 7.30 (dd, *J* = 9.2, 2.5 Hz, 1H), 7.16 (d, *J* = 7.0 Hz, 1H), 7.05 (t, *J* = 7.6 Hz, 1H), 7.00 (s, 1H), 6.94 (d, *J* = 7.7 Hz, 1H), 6.78 (t, *J* = 7.2 Hz, 1H), 6.71 (d, *J* = 7.9 Hz, 1H), 6.74 – 6.60 (br, 1H), 5.19 (d, *J* = 3.4 Hz, 1H), 4.99 – 4.78 (m, 2H), 4.64 – 4.46 (m, 3H), 4.38 – 4.24 (m, 2H), 4.17 (dd, *J* = 16.4, 5.0 Hz, 1H), 4.10 (br, 2H), 3.77 – 3.67 (m, 1H), 3.66 – 3.56 (m, 2H), 3.55 – 3.25 (m, 6H), 3.18 (dd, *J* = 15.9, 9.0 Hz, 1H), 3.13 – 3.02 (m, 1H), 3.02 – 2.90 (m, 1H), 2.76 (t, *J* = 12.6 Hz, 1H), 2.45 (s, 3H), 2.12 – 2.03 (m, 1H), 1.91 (ddd, *J* = 12.8, 9.7, 4.3 Hz, 1H), 1.83 – 1.45 (m, 13H), 1.42 – 1.31 (m, 2H), 1.24 – 1.17 (m, 2H), 0.94 (s, 9H), 0.67 – 0.54 (m, 1H); $^{13}$C NMR (101 MHz, DMSO) δ 171.9, 169.6, 169.3, 169.0, 168.3, 168.1, 165.6, 164.6, 164.0, 163.2, 159.1, 156.0, 151.5, 147.9, 131.4, 131.0, 127.8, 126.9, 126.8, 125.2, 120.8, 120.2, 111.6, 109.1, 81.1, 79.4, 77.0, 69.0, 67.7, 58.9, 56.8, 56.6, 48.0, 43.9, 37.9, 37.4, 36.1, 32.3, 31.0, 26.2, 25.6, 25.2, 22.6, 22.0, 20.1, 19.2, 16.0, 13.1, 13.0, 12.9, 12.8. HRMS (ESI) *calcd* for $[C_{53}H_{69}FN_{11}O_7S]^+$ ($[M+H]^+$): *m/z* 1022.5081, found: 1022.5093.

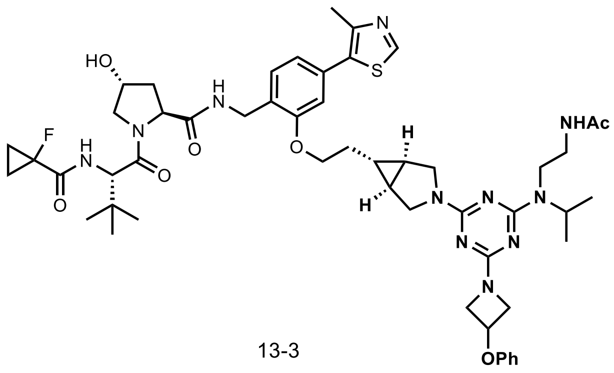

13-3

**Compound 13-3**: $^1$H NMR (400 MHz, DMSO) δ 8.96 (s, 1H), 8.50 (s, 1H), 7.97 – 7.88 (m, 1H), 7.40 (d, *J* = 7.8 Hz, 1H), 7.35 – 7.25 (m, 3H), 7.03 – 6.91 (m, 3H), 6.84 (d, *J* = 7.8 Hz, 2H), 5.21 (br, 1H), 5.04 (br, 1H), 4.88 – 4.72 (m, 1H), 4.58 (d, *J* = 9.2 Hz, 1H), 4.50 (t, *J* = 8.2 Hz, 1H), 4.42 – 4.24 (m, 4H), 4.17 (dd, *J* = 16.5, 5.2 Hz, 1H), 4.09 (t, *J* = 6.1 Hz, 2H), 3.90 – 3.77 (m, 2H), 3.72 (d, *J* = 11.0 Hz, 1H), 3.66 – 3.57 (m, 2H), 3.39 – 3.14 (m, 7H), 2.44 (s, 3H), 2.07 (dd, *J* = 12.4, 7.8 Hz, 1H), 1.91 (ddd, *J* = 13.0, 8.9, 4.5 Hz, 1H), 1.77 (s, 3H), 1.76 – 1.61 (m, 2H), 1.48 (br, 2H), 1.41 – 1.30 (m, 2H), 1.21 (dd, *J* = 8.1, 2.9 Hz, 2H), 1.06 (d, *J* = 6.1 Hz, 6H), 0.93 (s, 9H), 0.66 – 0.55 (m, 1H); $^{13}$C NMR (101 MHz, DMSO) δ 172.0, 169.6, 169.1, 168.4, 168.2, 166.4, 164.2, 163.8, 163.7, 156.5, 156.0, 151.6, 148.0, 131.5, 131.0, 129.9, 127.8, 127.0, 121.4, 120.8, 114.7, 111.7, 79.4, 77.1, 69.0, 67.7, 66.6, 59.0, 56.8, 56.7, 56.3, 48.1, 48.0, 45.1, 44.9, 40.8, 38.4, 38.3, 37.9, 37.5, 36.2, 31.1, 26.2, 22.7, 22.0, 20.2, 16.1, 13.2, 13.1, 12.9, 12.8. HRMS (ESI) *calcd* for $[C_{52}H_{68}FN_{11}O_7SNa]^+$ ($[M+Na]^+$): *m/z* 1032.4900, found: 1032.4932.

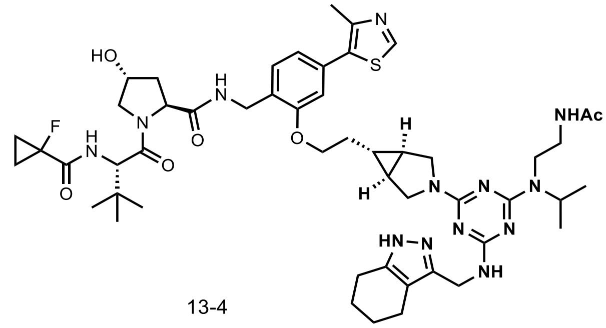

13-4

**Compound 13-4**: $^1$H NMR (400 MHz, DMSO) δ 8.97 (s, 1H), 8.56 – 8.45 (m, 1H), 8.02 – 7.86 (m, 1H), 7.42 (d, *J* = 7.8 Hz, 1H), 7.29 (dd, *J* = 9.2, 2.6 Hz, 1H), 7.01 (s, 1H), 6.94 (dd, *J* = 7.8, 1.2 Hz, 1H), 6.62 (br, 1H), 5.18 (br, 1H), 4.97 – 4.74 (m, 1H), 4.59 (d, *J* = 9.2 Hz, 1H), 4.51 (t, *J* = 8.2 Hz, 1H), 4.41 – 4.24 (m, 4H), 4.17 (dd, *J* = 16.5, 5.3 Hz, 1H), 4.10 (t, *J* = 5.8 Hz, 2H), 3.89 – 3.77 (m, 1H), 3.77 – 3.66 (m, 1H), 3.66 – 3.57 (m, 2H), 3.35 – 3.15 (m, 6H), 2.50 – 2.45 (m, 2H), 2.45 (s, 3H), 2.36 (t, *J* = 5.2 Hz, 1H), 2.12 – 2.03 (m, 1H), 1.91 (ddd, *J* = 12.7, 9.1, 4.2 Hz, 1H), 1.77 (s, 3H), 1.77 – 1.69 (m, 2H), 1.69 – 1.55 (m, 4H), 1.49 (br, 2H), 1.36 (ddd, *J* = 18.2, 5.9, 3.5 Hz, 2H), 1.21 (dd, *J* = 8.5, 2.7 Hz, 2H), 1.06 (br, 6H), 0.94 (s, 9H), 0.67 – 0.56 (m, 1H); $^{13}$C NMR (101 MHz, DMSO) δ 171.9, 169.3, 169.0, 168.2, 168.0, 165.2, 165.1, 164.3, 163.9, 155.9, 151.5, 147.9, 131.4, 130.9, 127.8, 126.9, 120.7, 111.8, 111.6, 79.3, 77.0, 69.0, 67.6, 58.8, 56.7, 56.6, 47.9, 38.4, 37.9, 37.4, 36.1, 31.0, 26.21, 26.17, 23.0, 22.6, 21.8, 20.2, 19.8, 16.0, 13.1, 13.0, 12.8, 12.7; HRMS (ESI) *calcd* for $[C_{51}H_{71}FN_{13}O_7S]^+$ ($[M+H]^+$): *m/z* 1012.5350, found: 1012.5353.

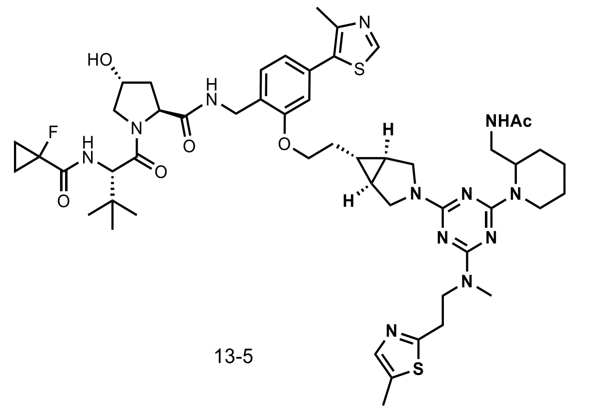

13-5

**Compound 13-5**: $^{1}$H NMR (400 MHz, $CDCl_3$) δ 8.68 (s, 1H), 7.36 – 7.28 (m, 2H), 7.26 – 7.18 (m, 1H), 7.01 (dd, *J* = 9.1, 3.7 Hz, 1H), 6.96 (d, *J* = 7.5 Hz, 1H), 6.87 (s, 1H), 4.98 (br, 1H), 4.80 – 4.60 (m, 2H), 4.60 – 4.46 (m, 3H), 4.41 (dd, *J* = 14.7, 5.0 Hz, 1H), 4.07 (t, *J* = 6.4 Hz, 2H), 3.92 (br, 4H), 3.70 – 3.55 (m, 2H), 3.49 (br, 2H), 3.33 – 3.15 (m, 3H), 3.08 (s, 3H), 2.89 (br, 1H), 2.53 (s, 3H), 2.53 – 2.46 (m, 1H), 2.43 (s, 3H), 2.08 – 2.04 (m, 1H), 1.97 – 1.75 (m, 6H), 1.75 – 1.56 (m, 6H), 1.56 – 1.40 (m, 3H), 1.36 – 1.26 (m, 4H), 0.92 (s, 9H), 0.82 – 0.65 (m, 1H); $^{13}$C NMR (101 MHz, $CDCl_3$) δ 171.1, 170.7, 170.5, 170.3, 170.1, 156.9, 150.4, 148.6, 140.0, 133.8, 132.5, 131.9, 129.8, 126.3, 121.8, 112.2, 79.6, 74.6, 70.2, 68.0, 58.6, 57.5, 56.6, 49.8, 48.8, 39.0, 36.0, 35.6, 31.8, 31.4, 26.4, 25.5, 25.4, 25.0, 23.4, 22.7, 20.8, 19.6, 16.3, 13.91, 13.86, 13.81, 13.76, 12.1; HRMS (ESI) *calcd* for $[C_{51}H_{70}FN_{12}O_6S_2]^+$ ($[M+H]^+$): *m/z* 1029.4961, found: 1029.4994.

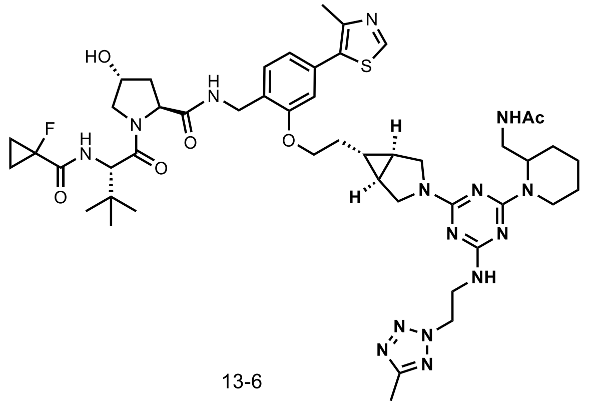

13-6

**Compound 13-6**: $^{1}$H NMR (400 MHz, $CDCl_3$) δ 8.68 (s, 1H), 7.39 – 7.27 (m, 2H), 6.99 (d, *J* = 8.9 Hz, 1H), 6.96 (dd, *J* = 7.6, 1.5 Hz, 1H), 6.87 (d, *J* = 1.7 Hz, 1H), 6.58 (br, 1H), 5.01 (br, 1H), 4.87 – 4.58 (m, 4H), 4.58 – 4.45 (m, 3H), 4.41 (d, *J* = 14.3 Hz, 1H), 4.08 (t, *J* = 6.0 Hz, 2H), 4.05 – 3.78 (m, 5H), 3.62 (dd, *J* = 11.0, 4.1 Hz, 1H), 3.62 – 3.33 (m, 3H), 3.33 – 2.98 (m, 1H), 2.83 (br, 1H), 2.53 (s, 4H), 2.52 (s, 3H), 2.10 – 2.02 (m, 1H), 2.00 – 1.76 (m, 5H), 1.76 – 1.57 (m, 5H), 1.57 – 1.37 (m, 3H), 1.36 – 1.26 (m, 4H), 0.93 (s, 9H), 0.81 – 0.67 (m, 1H); $^{13}$C NMR (101 MHz, $CDCl_3$) δ 171.2, 170.5, 170.2, 165.8, 163.3, 156.9, 150.4, 148.7, 132.5, 131.9, 129.7, 126.3, 121.8, 112.2, 79.6, 70.3, 67.9, 58.6, 57.6, 56.5, 52.4, 51.3, 49.6, 48.7, 40.1, 39.0, 38.5, 35.9, 35.5, 31.7, 29.8, 27.1, 26.4, 25.4, 23.4, 22.7, 22.5, 20.8, 19.7, 19.1, 16.3, 13.9, 13.8, 11.0; HRMS (ESI) *calcd* for $[C_{48}H_{67}FN_{15}O_6S]^+$ ($[M+H]^+$): *m/z* 1000.5098, found: 1000.5129.

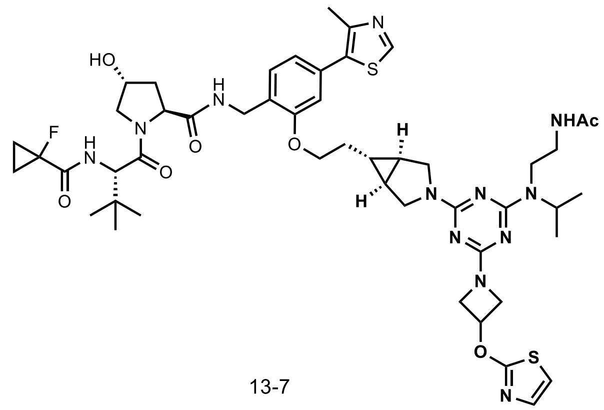

13-7

**Compound 13-7**: $^{1}$H NMR (400 MHz, $CDCl_3$) δ 8.68 (s, 1H), 7.68 (br, 1H), 7.32 (d, *J* = 7.8 Hz, 1H), 7.22 (t, *J* = 6.0 Hz, 1H), 7.11 (d, *J* = 4.0 Hz, 1H), 7.00 (dd, *J* = 8.6, 3.6 Hz, 1H), 6.96 (dd, *J* = 7.7, 1.2 Hz, 1H), 6.87 (d, *J* = 1.2 Hz, 1H), 6.72 (d, *J* = 3.9 Hz, 1H), 5.45 (tt, *J* = 6.6, 4.0 Hz, 1H), 4.89 (h, *J* = 6.8 Hz, 1H), 4.70 (t, *J* = 7.7 Hz, 1H), 4.61 – 4.33 (m, 6H), 4.23 – 4.09 (m, 2H), 4.07 (t, *J* = 6.4 Hz, 2H), 4.00 – 3.82 (m, 3H), 3.69 – 3.31 (m, 7H), 2.62 – 2.51 (m, 1H), 2.53 (s, 3H), 2.03 (t, *J* = 10.9 Hz, 1H), 1.87 (s, 3H), 1.84 – 1.60 (m, 2H), 1.47 (s, 2H), 1.36 – 1.24 (m, 4H), 1.14 (d, *J* = 6.7 Hz, 6H), 0.92 (s, 9H), 0.78 – 0.66 (m, 1H); $^{13}$C NMR (101 MHz, $CDCl_3$) δ 173.1, 171.2, 170.4, 170.2, 166.4, 165.8, 164.1, 156.9, 150.4, 148.6, 137.2, 132.5, 131.9, 129.7, 126.3, 121.8, 112.2, 79.6, 70.7, 70.2, 68.0, 58.6, 57.6, 56.7, 56.4, 48.5, 46.4, 42.8, 40.1, 39.0, 35.8, 35.4, 31.8, 29.8, 26.4, 23.4, 22.6, 20.5, 20.3, 16.3, 13.9, 13.8; HRMS (ESI) *calcd* for $[C_{49}H_{66}FN_{12}O_7S_2]^+$ ($[M+H]^+$): *m/z* 1017.4597, found: 1017.4617.

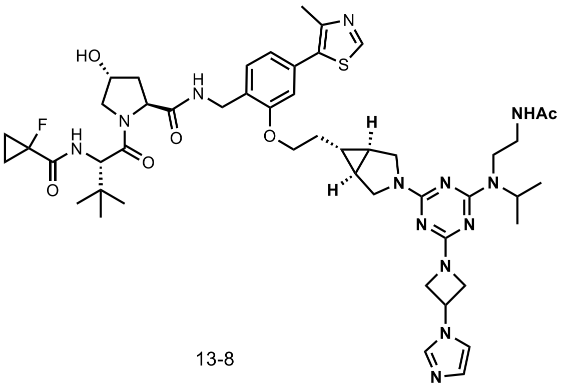

13-8

**Compound 13-8**: $^{1}$H NMR (400 MHz, $CDCl_3$) δ 8.68 (s, 1H), 8.18 (br, 1H), 7.40 – 7.26 (m, 3H), 7.02 (dd, *J* = 8.8, 3.4 Hz, 1H), 6.96 (dd, *J* = 7.5, 1.6 Hz, 1H), 6.86 (d, *J* = 1.6 Hz, 1H), 5.12 (br, 1H), 4.88 (h, *J* = 6.7 Hz, 1H), 4.70 (t, *J* = 7.4 Hz, 1H), 4.64 – 4.45 (m, 5H), 4.41 (dd, *J* = 14.9, 5.1 Hz, 1H), 4.22 (br, 2H), 4.07 (t, *J* = 6.1 Hz, 2H), 3.99 – 3.83 (m, 3H), 3.70 – 3.32 (m, 7H), 2.52 (s, 3H), 2.52 – 2.45 (m, 1H), 2.12 – 1.98 (m, 1H), 1.48 (s, 2H), 1.36 – 1.25 (m, 4H), 1.16 (d, *J* = 6.7 Hz, 6H), 0.92 (s, 9H), 0.80 – 0.69 (m, 1H); $^{13}$C NMR (151 MHz, $CDCl_3$) δ 171.0, 170.4, 170.3, 170.1, 170.0, 166.1, 165.4, 163.7, 156.7, 150.3, 148.5, 132.3, 131.8, 129.4, 126.1, 121.6, 112.0, 79.0, 70.0, 67.8, 58.53, 58.48, 57.4, 56.4, 48.5, 48.4, 46.2, 42.0, 40.2, 38.7, 35.8, 35.41, 35.36, 31.6, 29.7, 26.3, 23.2, 22.5, 20.5, 20.1, 16.1, 13.8, 13.7, 13.6; HRMS (ESI) *calcd* for $[C_{49}H_{66}FN_{13}O_6SNa]^+$ ($[M+Na]^+$): *m/z* 1006.4856, found: 1006.4977.

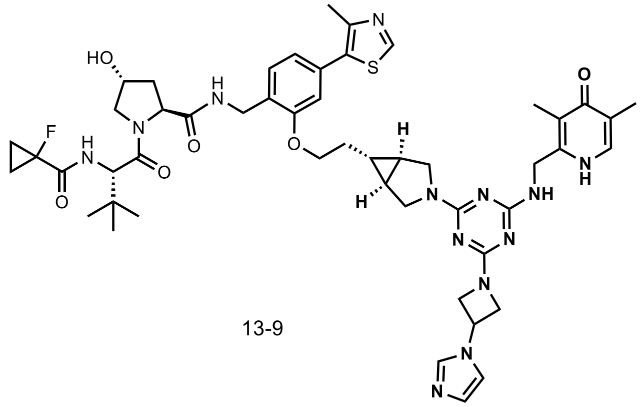

13-9

**Compound 13-9**: $^{1}$H NMR (400 MHz, $CDCl_3$) δ 8.67 (s, 1H), 7.64 (s, 1H), 7.44 – 7.26 (m, 2H), 7.19 – 7.08 (m, 2H), 7.08 – 7.01 (m, 1H), 7.00 – 6.09 (m, 1H), 6.86 (s, 1H), 6.10 (br, 1H), 5.11 – 4.90 (m, 1H), 4.82 – 4.28 (m, 9H), 4.25 – 3.99 (m, 4H), 3.99 – 3.72 (m, 3H), 3.72 – 3.57 (m, 1H), 3.51 – 3.29 (m, 2H), 2.51 (s, 3H), 2.50 – 2.33 (m, 1H), 2.08 (s, 4H), 1.97 (s, 3H), 1.94 – 1.83 (m, 1H), 1.78 – 1.61 (m, 1H), 1.57 – 1.40 (m, 2H), 1.40 – 1.25 (m, 4H), 0.93 (s, 9H), 0.78 – 0.66 (m, 1H); $^{13}$C NMR (101 MHz, $CDCl_3$) δ 178.9, 171.0, 170.9, 170.2, 170.0, 166.1, 165.9, 163.8, 163.7, 156.8, 150.5, 148.6, 144.1, 136.4, 132.8, 132.5, 131.9, 130.4, 129.8, 126.1, 123.2, 121.8, 121.7, 121.0, 117.0, 112.1, 112.0, 79.7, 70.1, 67.9, 67.7, 59.0, 58.9, 57.5, 56.7, 48.7, 46.9, 40.3, 40.2, 39.3, 39.0, 36.2, 35.8, 31.7, 29.8, 26.4, 22.8, 22.6, 22.4, 20.9, 16.3, 14.3, 13.9, 13.8, 13.7, 13.6, 10.4; HRMS (ESI) *calcd* for $[C_{50}H_{62}FN_{13}O_6SNa]^+$ ($[M+Na]^+$): *m/z* 1014.4543, found: 1014.4509.

### NMR spectra of compounds

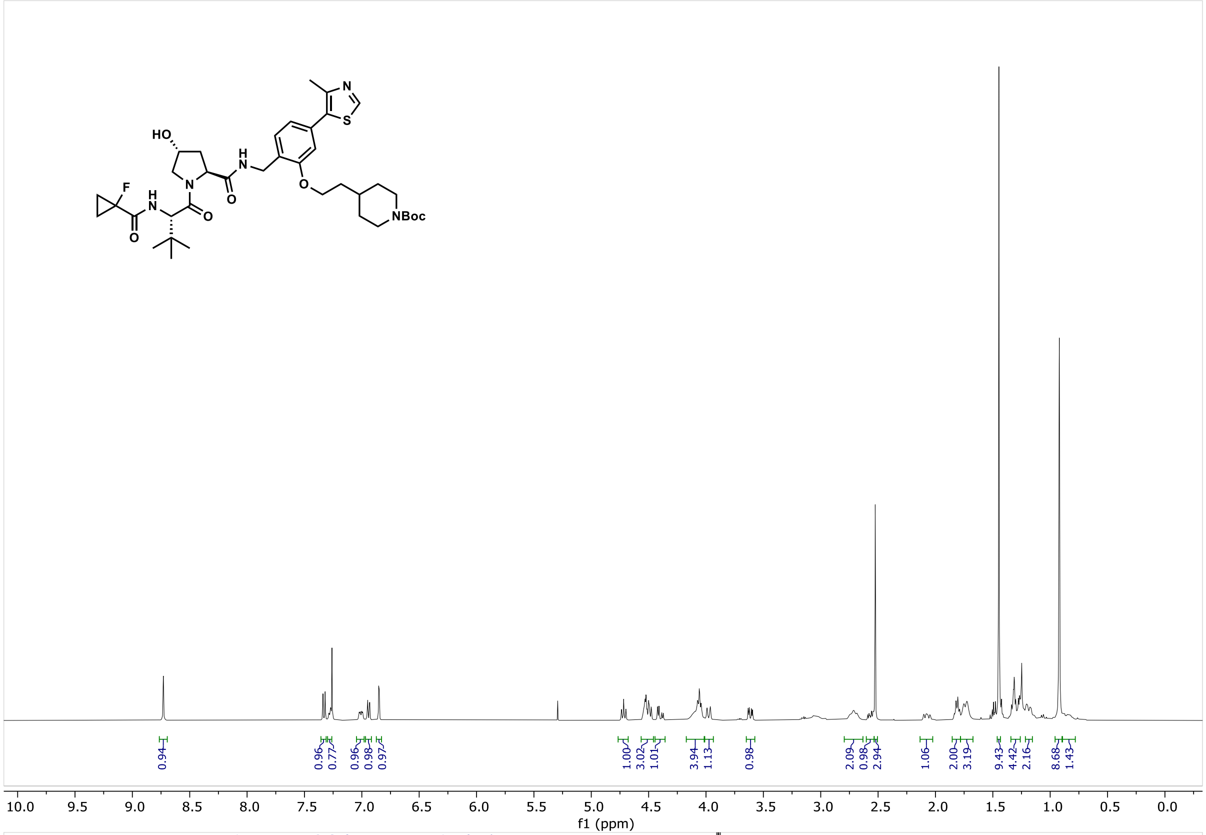

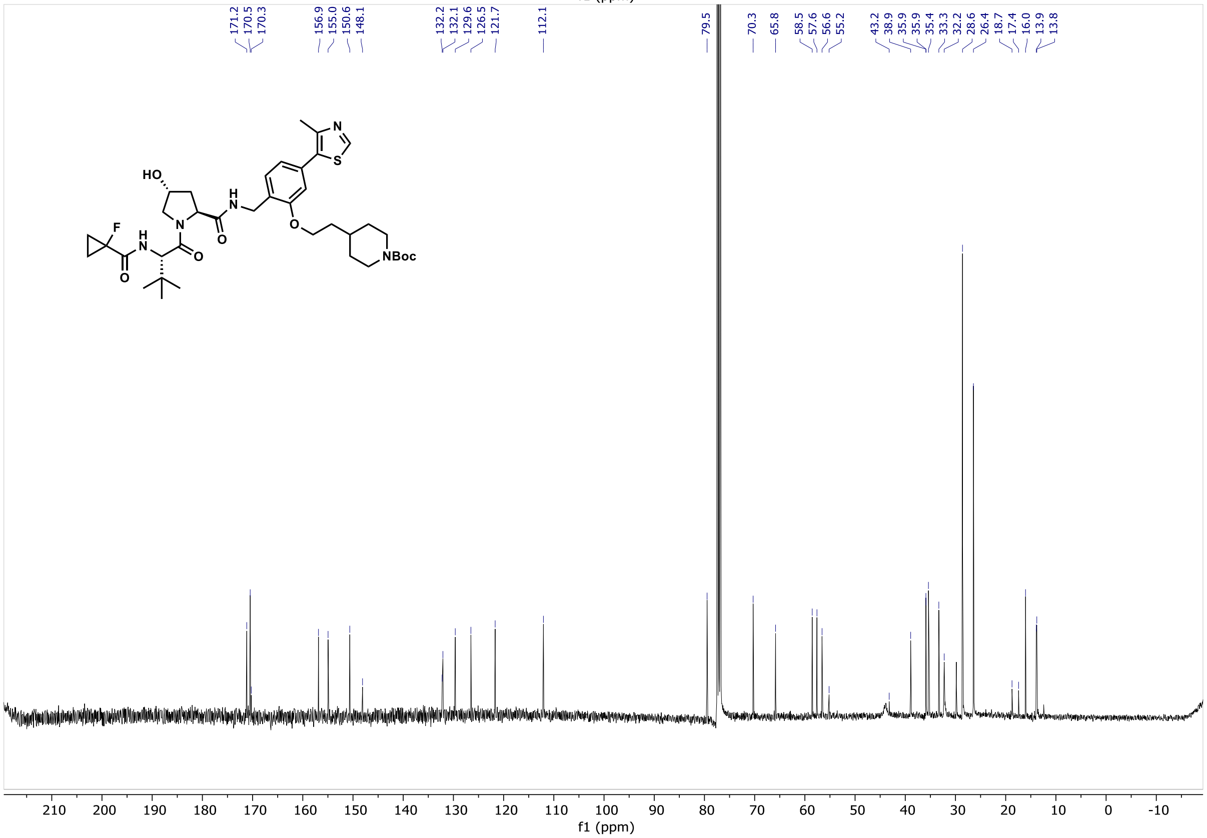

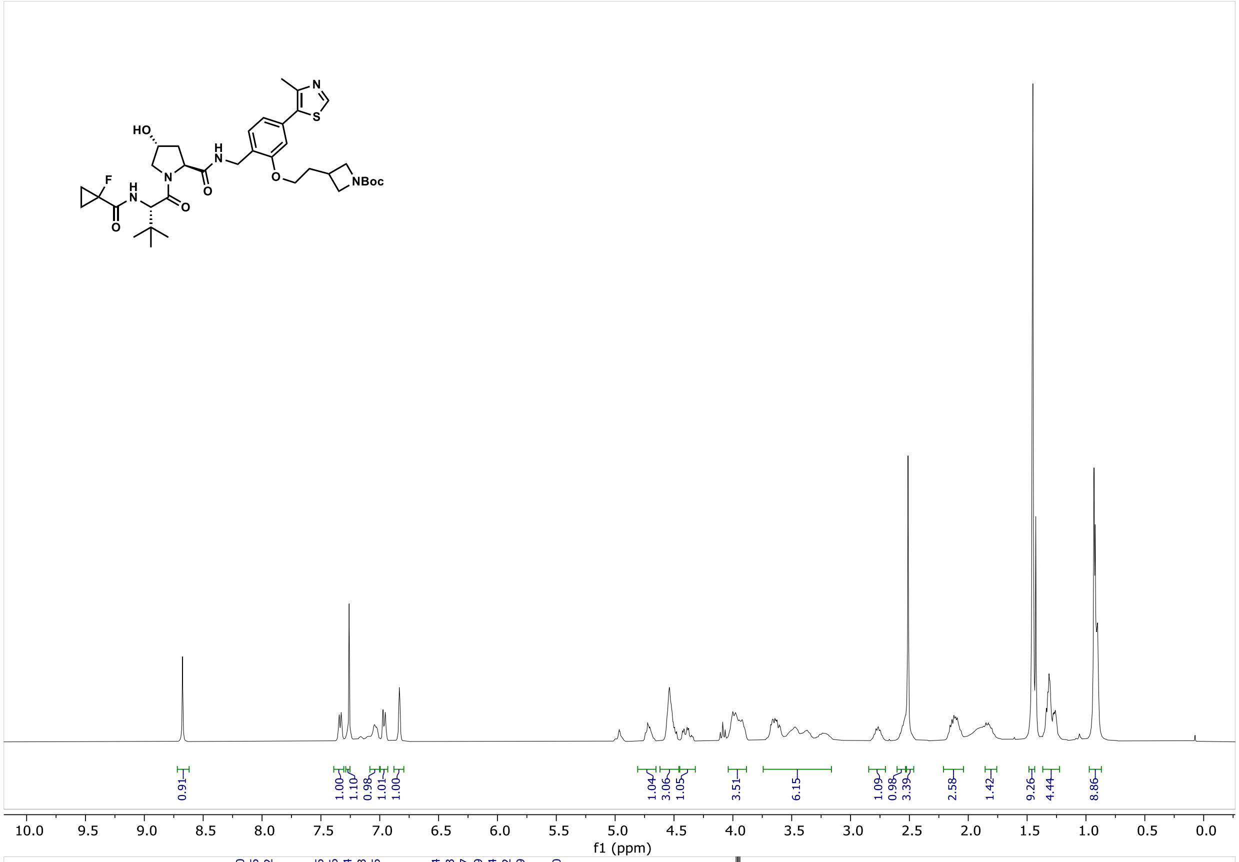
HO
N
H
N
O
O
F
H
N
O
O
N
S
O
NBoc
0.91
1.00
1.10
0.98
1.01
1.00
1.04
3.06
1.05
3.51
6.15
1.09
0.98
3.39
2.58
1.42
9.26
4.44
8.86
10.0 9.5 9.0 8.5 8.0 7.5 7.0 6.5 6.0 5.5 5.0 4.5 4.0 3.5 3.0 2.5 2.0 1.5 1.0 0.5 0.0
f1 (ppm)

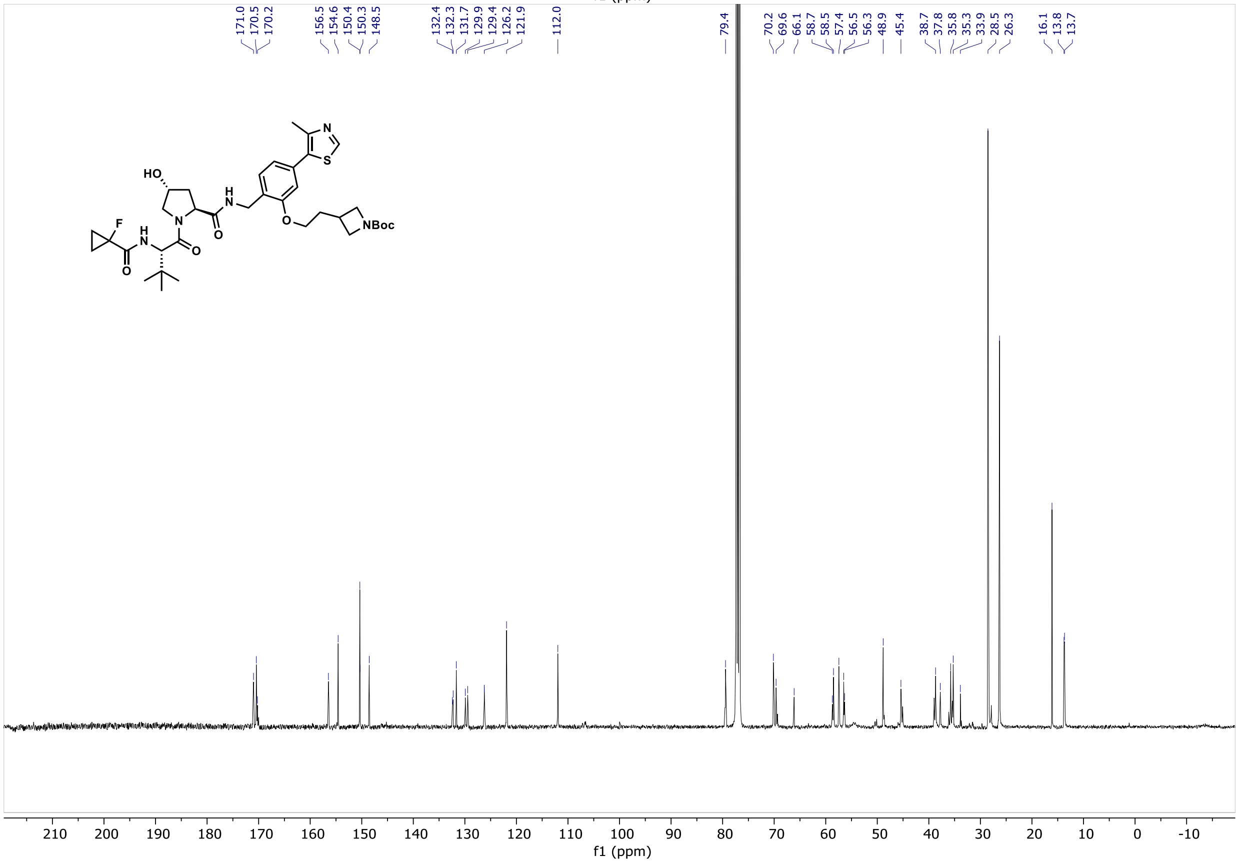
171.0
170.5
170.2
156.5
154.6
150.4
150.3
148.5
132.4
132.3
131.7
129.9
129.4
126.2
121.9
112.0
79.4
70.2
69.6
66.1
58.7
58.5
57.4
56.5
56.3
48.9
45.4
38.7
37.8
35.8
35.3
33.9
28.5
26.3
16.1
13.8
13.7
HO
N
H
N
O
O
F
H
N
O
O
N
S
O
NBoc
210 200 190 180 170 160 150 140 130 120 110 100 90 80 70 60 50 40 30 20 10 0 -10
f1 (ppm)

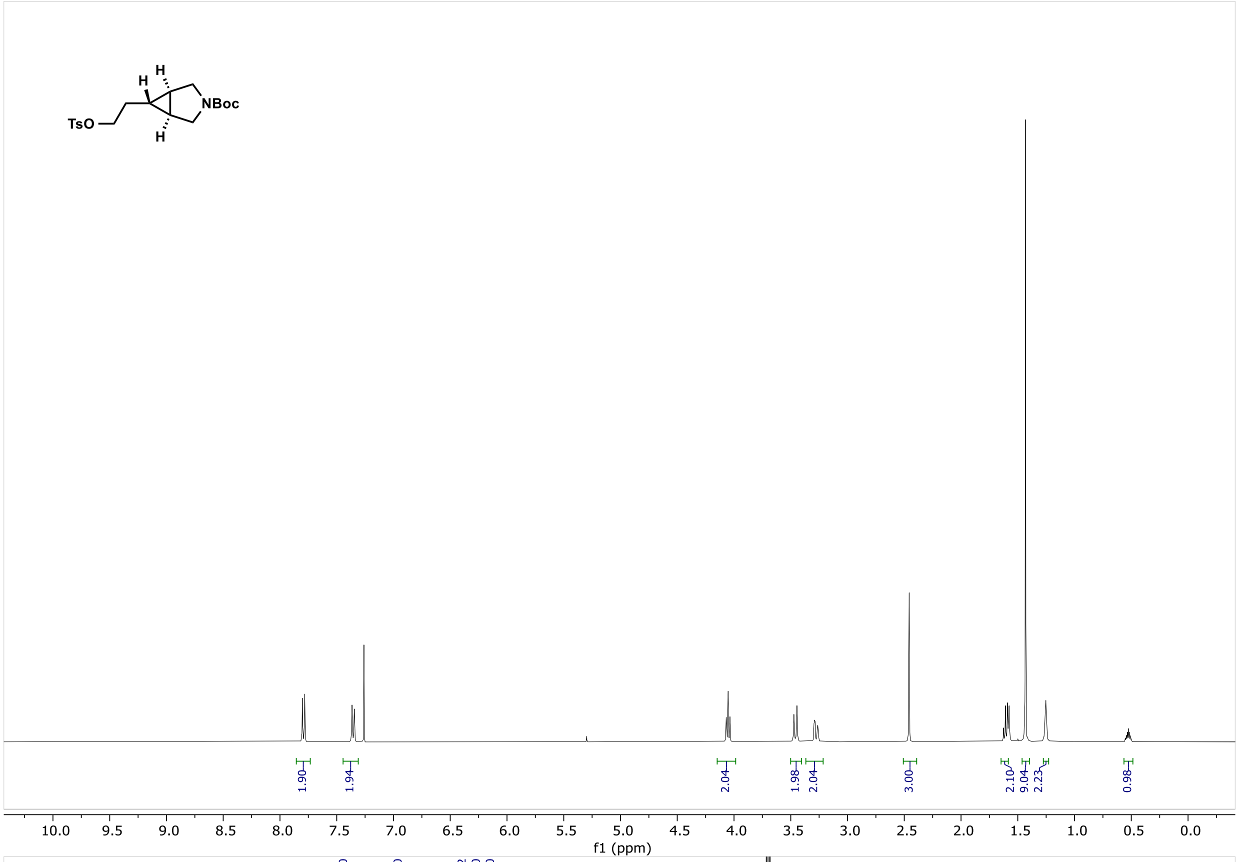
TsO
NBoc
H
H
H
1.90
1.94
2.04
1.98
2.04
3.00
2.10
9.04
2.23
0.98
10.0 9.5 9.0 8.5 8.0 7.5 7.0 6.5 6.0 5.5 5.0 4.5 4.0 3.5 3.0 2.5 2.0 1.5 1.0 0.5 0.0
f1 (ppm)

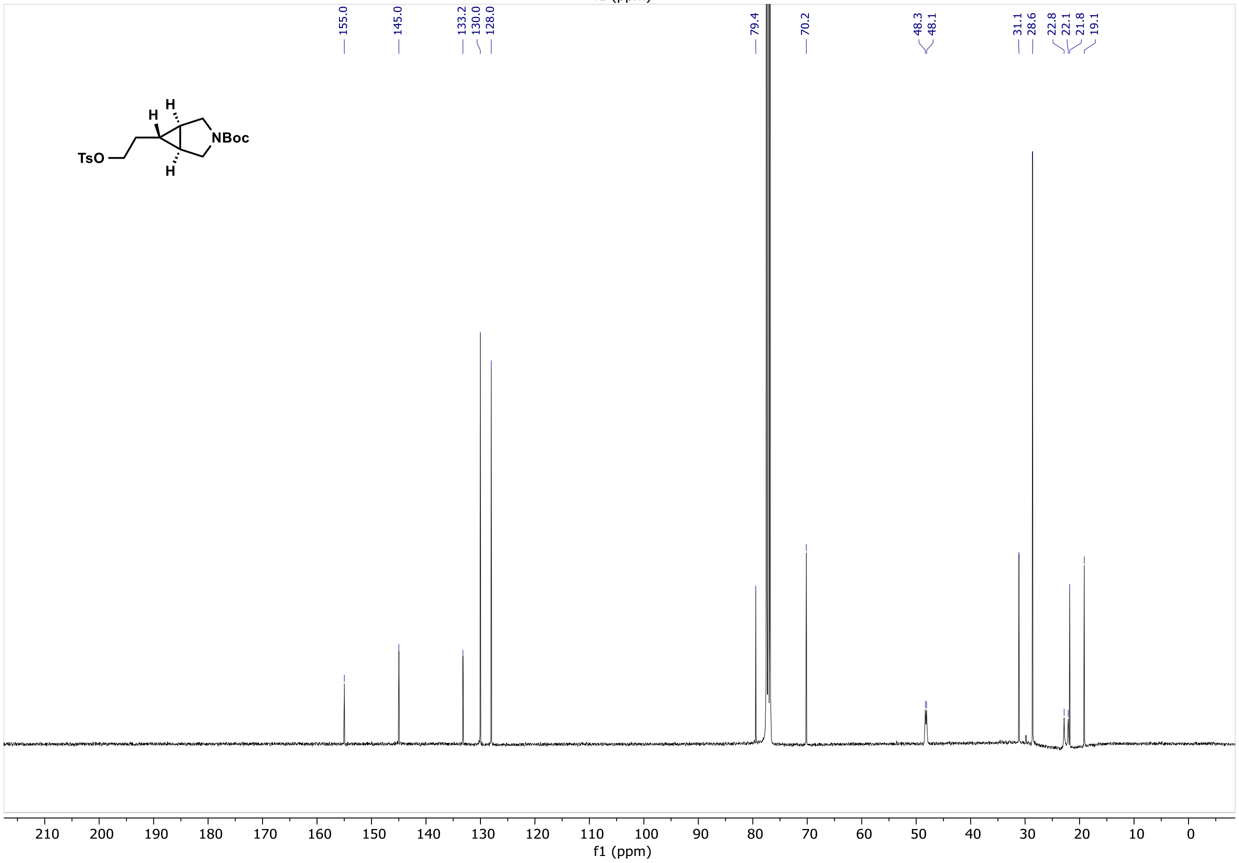
155.0
145.0
133.2
130.0
128.0
79.4
70.2
48.3
48.1
31.1
28.6
22.8
22.1
21.8
19.1
TsO
NBoc
H
H
H
210 200 190 180 170 160 150 140 130 120 110 100 90 80 70 60 50 40 30 20 10 0
f1 (ppm)

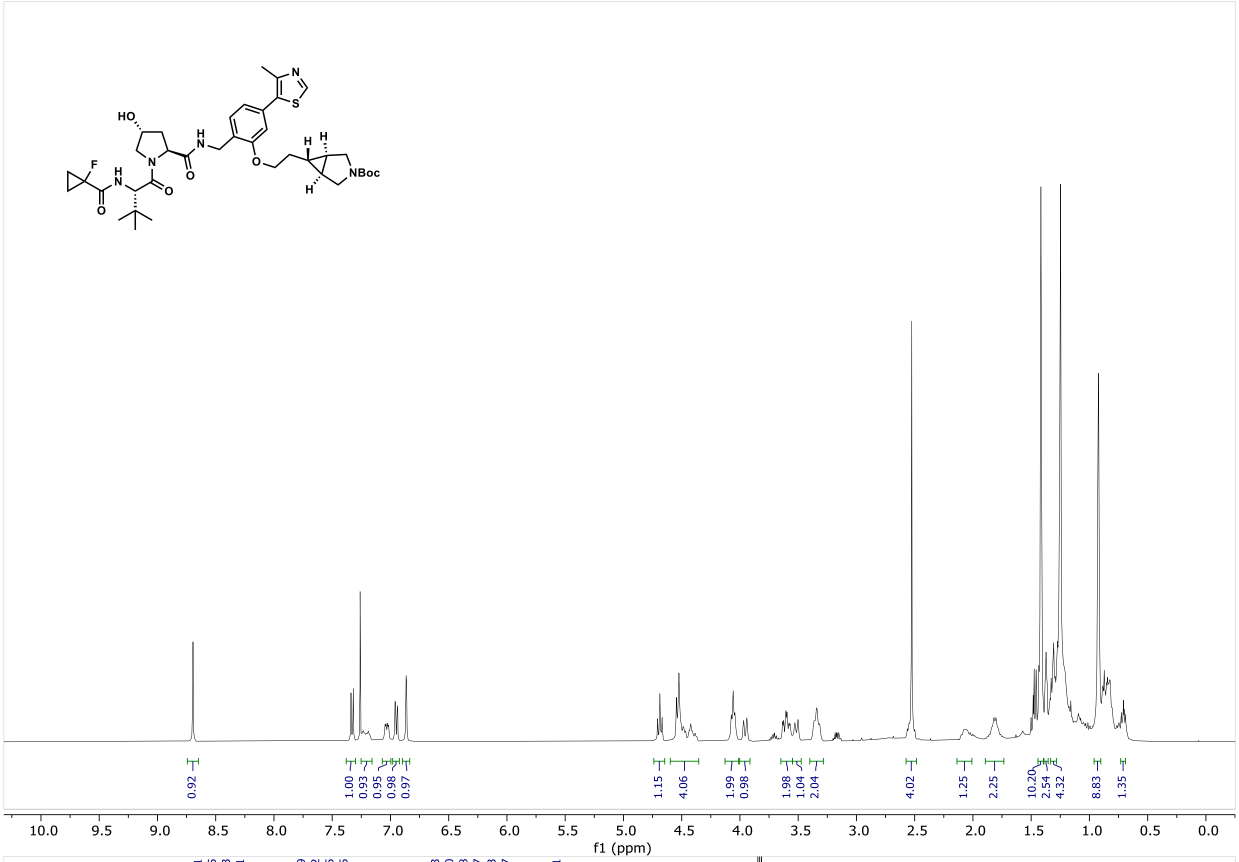
HO
N
H
NH
F
O
S
N
NBoc
0.92
1.00
0.93
0.95
0.98
0.97
1.15
4.06
1.99
0.98
1.98
1.04
2.04
4.02
1.25
2.25
10.20
2.54
4.32
8.83
1.35
10.0 9.5 9.0 8.5 8.0 7.5 7.0 6.5 6.0 5.5 5.0 4.5 4.0 3.5 3.0 2.5 2.0 1.5 1.0 0.5 0.0
f1 (ppm)

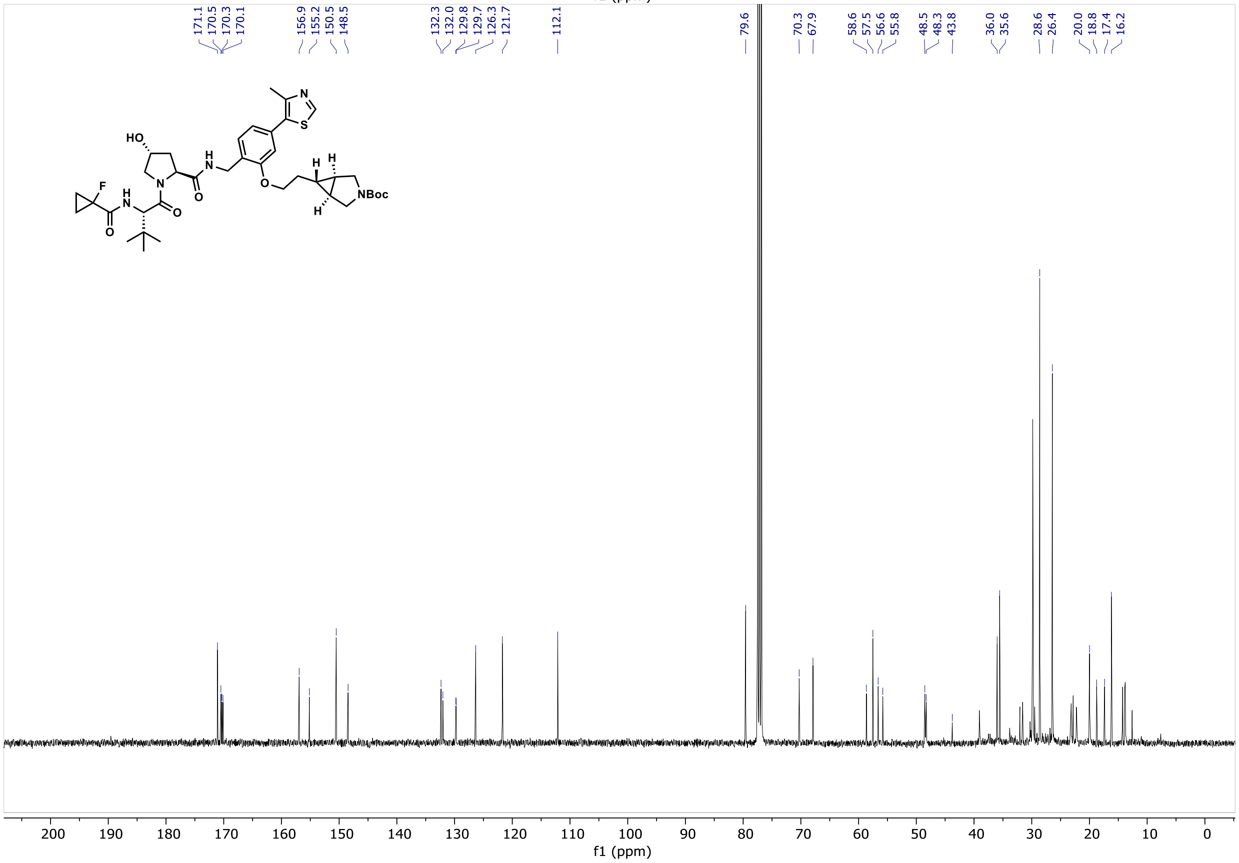
171.1
170.5
170.3
170.1
156.9
155.2
150.5
148.5
132.3
132.0
129.8
129.7
126.3
121.7
112.1
79.6
70.3
67.9
58.6
57.5
56.6
55.8
48.5
48.3
43.8
36.0
35.6
28.6
26.4
20.0
18.8
17.4
16.2
HO
N
H
NH
F
O
S
N
NBoc
200 190 180 170 160 150 140 130 120 110 100 90 80 70 60 50 40 30 20 10 0
f1 (ppm)

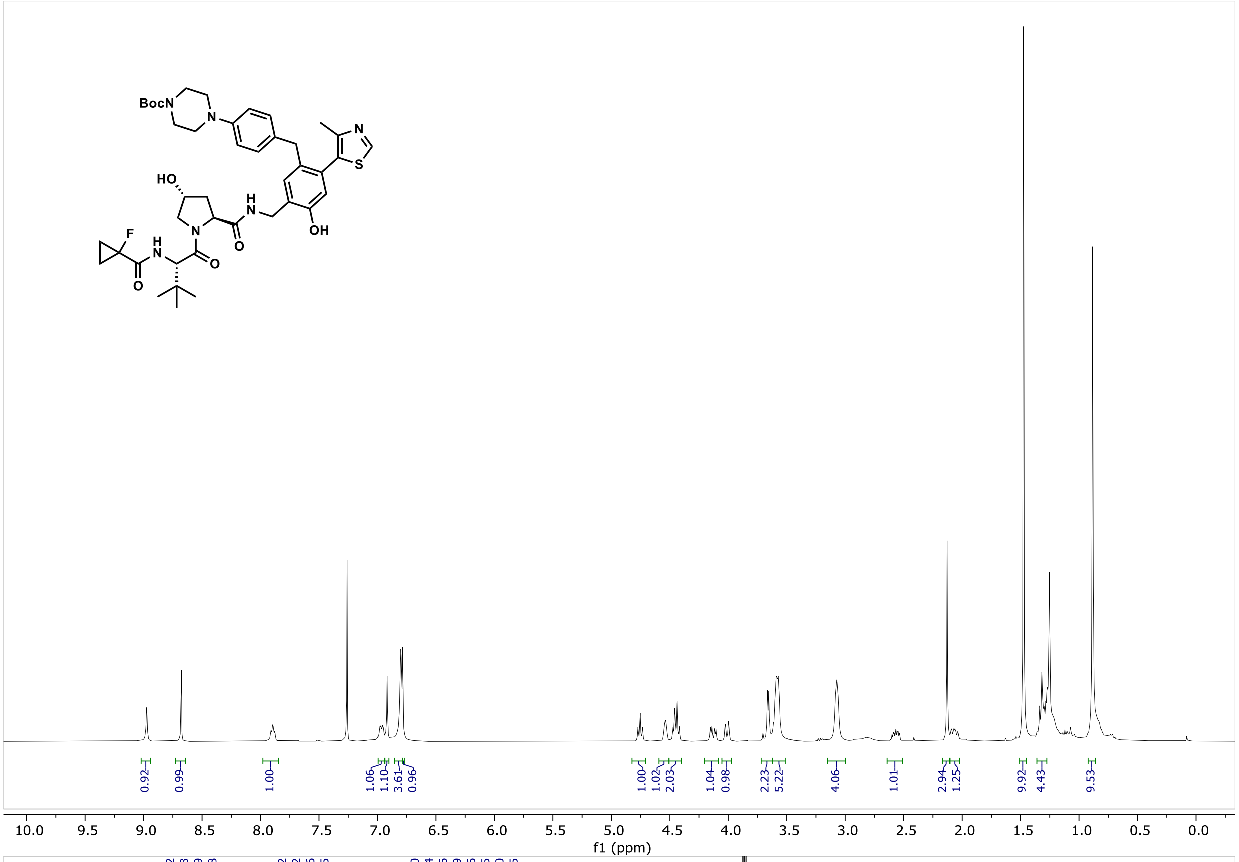
BocN
HO
OH
F
N
S
H
O
0.92
0.99
1.00
1.06
1.10
3.61
0.96
1.00
1.02
2.03
1.04
0.98
2.23
5.22
4.06
1.01
2.94
1.25
9.92
4.43
9.53
10.0 9.5 9.0 8.5 8.0 7.5 7.0 6.5 6.0 5.5 5.0 4.5 4.0 3.5 3.0 2.5 2.0 1.5 1.0 0.5 0.0
f1 (ppm)

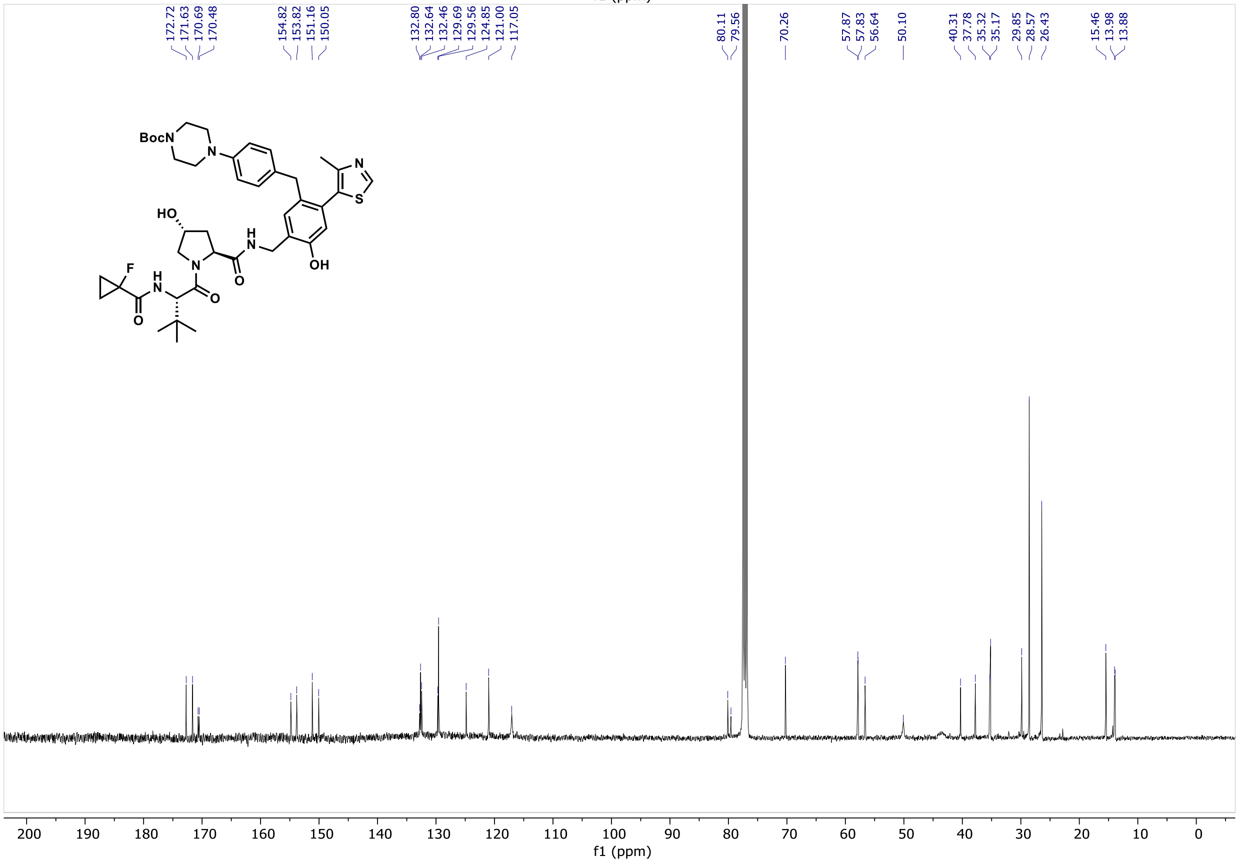
172.72
171.63
170.69
170.48
154.82
153.82
151.16
150.05
132.80
132.64
132.46
129.69
129.56
124.85
121.00
117.05
80.11
79.56
70.26
57.87
57.83
56.64
50.10
40.31
37.78
35.32
35.17
29.85
28.57
26.43
15.46
13.98
13.88
BocN
HO
OH
F
N
S
H
O
200 190 180 170 160 150 140 130 120 110 100 90 80 70 60 50 40 30 20 10 0
f1 (ppm)

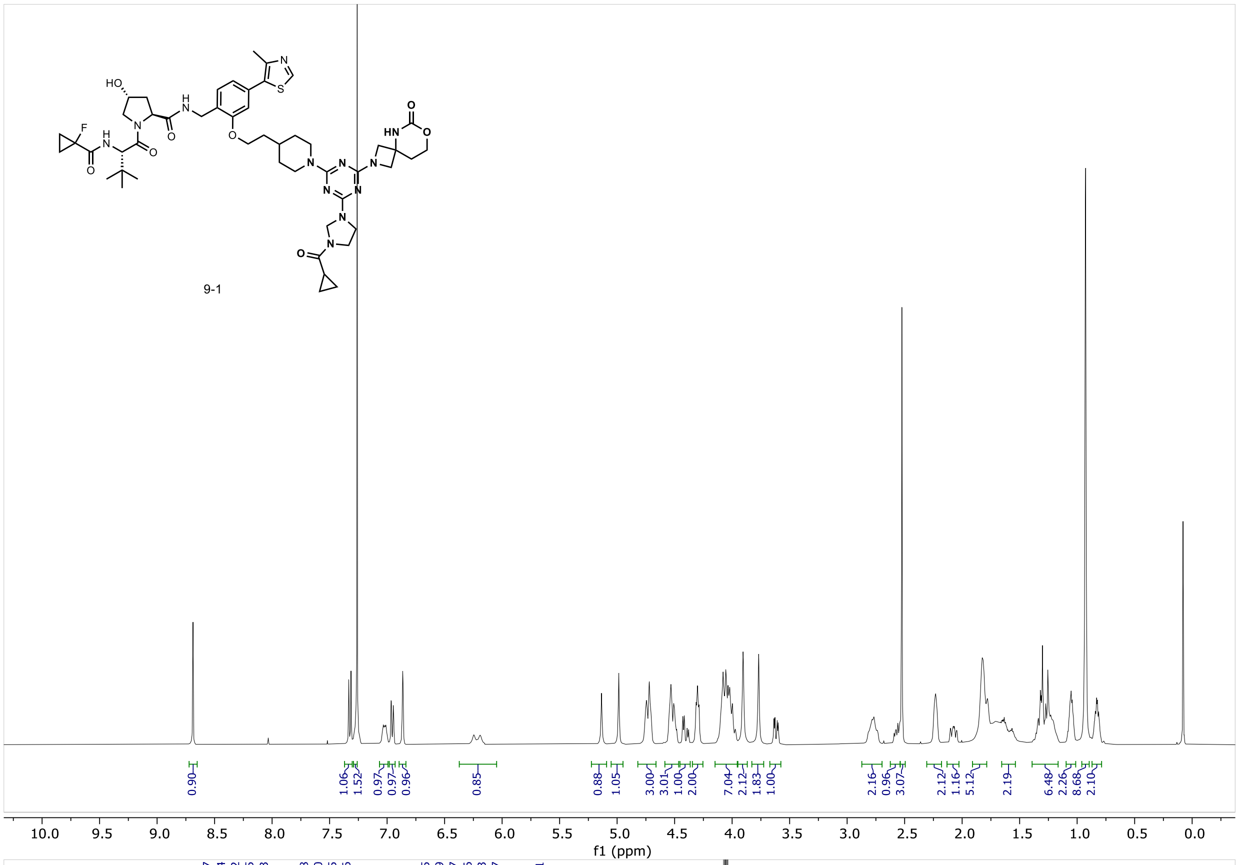
9-1
0.90
1.06
1.52
0.97
0.97
0.96
0.85
0.88
1.05
3.00
3.01
1.00
2.00
7.04
2.12
1.83
1.00
2.16
0.96
3.07
2.12
1.16
5.12
2.19
6.48
2.26
8.68
2.10
f1 (ppm)

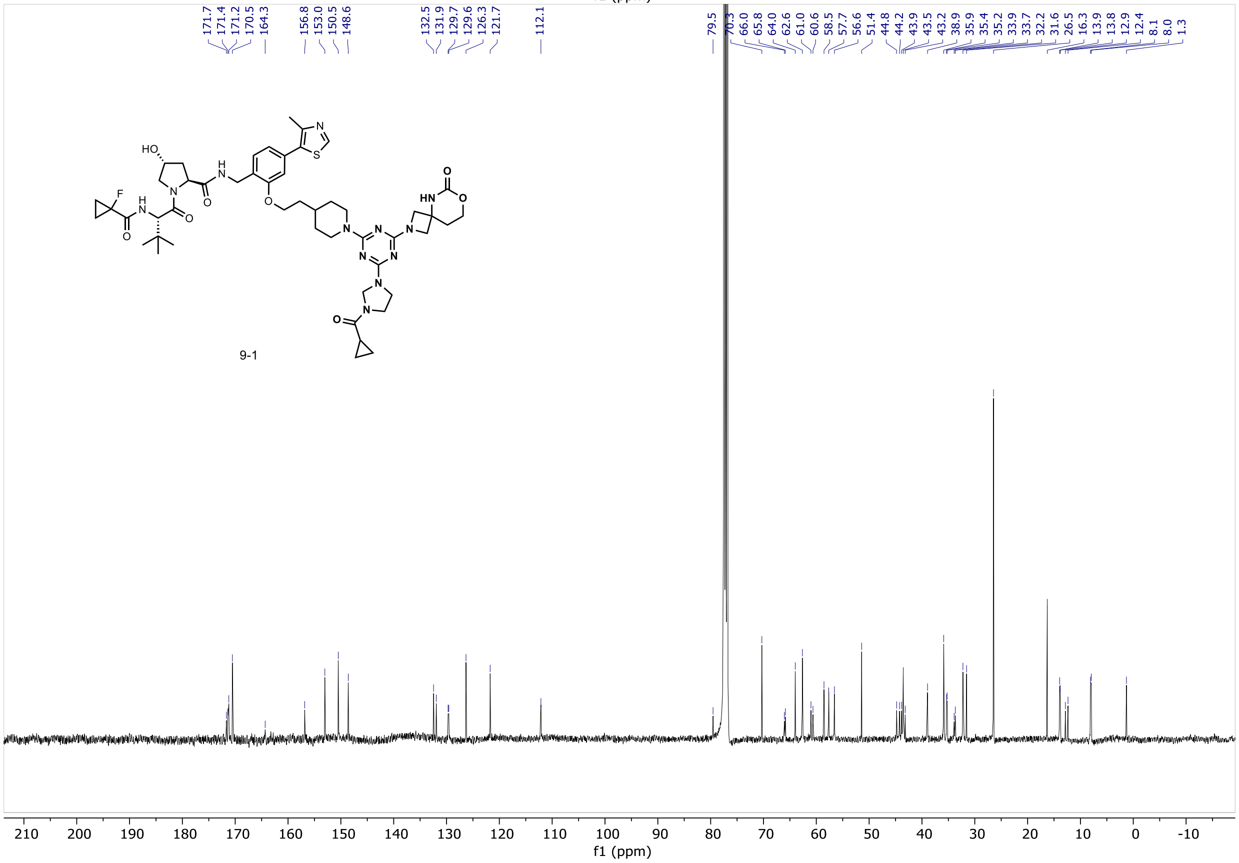
171.7
171.4
171.2
170.5
164.3
156.8
153.0
150.5
148.6
132.5
131.9
129.7
129.6
126.3
121.7
112.1
79.5
70.3
66.0
65.8
64.0
62.6
61.0
60.6
58.5
57.7
56.6
51.4
44.8
44.2
43.9
43.5
43.2
38.9
35.9
35.4
35.2
33.9
33.7
32.2
31.6
26.5
16.3
13.9
13.8
12.9
12.4
8.1
8.0
1.3
9-1
f1 (ppm)

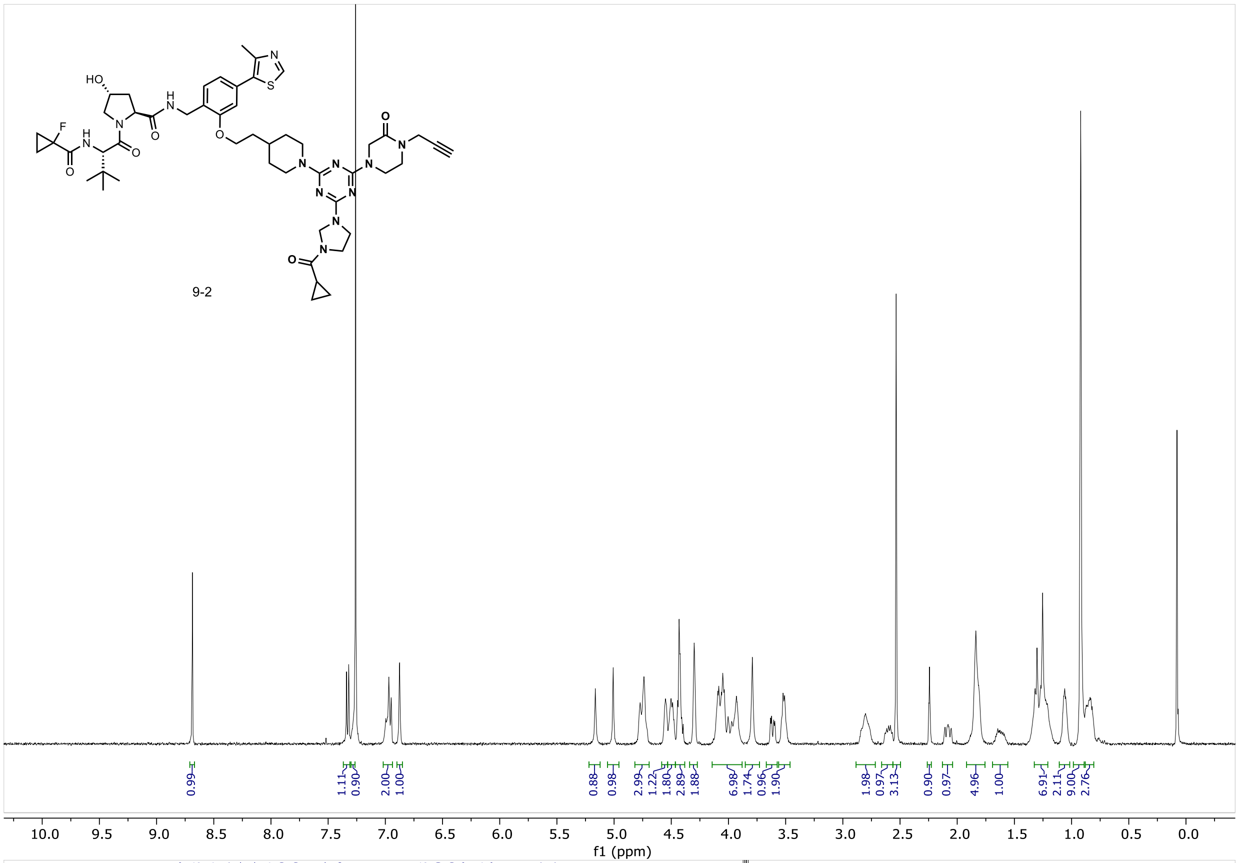
9-2
0.99 1.11 0.90 2.00 1.00 0.88 0.98 2.99 1.22 1.80 2.89 1.88 6.98 1.74 0.96 1.90 1.98 0.97 3.13 0.90 0.97 4.96 1.00 6.91 2.11 9.00 2.76
10.0 9.5 9.0 8.5 8.0 7.5 7.0 6.5 6.0 5.5 5.0 4.5 4.0 3.5 3.0 2.5 2.0 1.5 1.0 0.5 0.0
f1 (ppm)

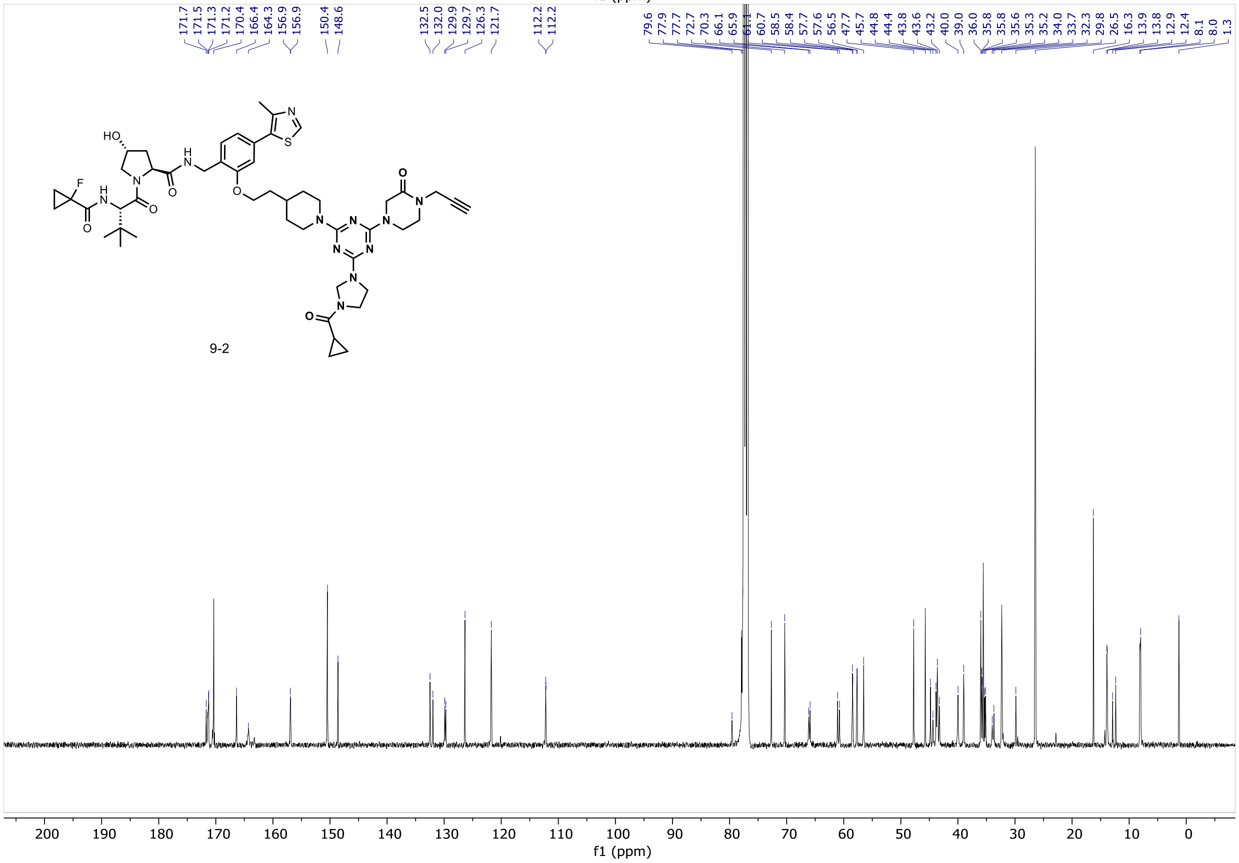
171.7 171.5 171.3 171.2 170.4 166.4 164.3 156.9 156.9 150.4 148.6 132.5 132.0 129.9 129.7 126.3 121.7 112.2 112.2 79.6 77.9 77.7 72.7 70.3 66.1 65.9 61.1 60.7 58.5 58.4 57.7 57.6 56.5 47.7 45.7 44.8 44.4 43.8 43.6 43.2 40.0 39.0 36.0 35.8 35.8 35.6 35.3 35.2 34.0 33.7 32.3 29.8 26.5 16.3 13.9 13.8 12.9 12.4 8.1 8.0 1.3
9-2
200 190 180 170 160 150 140 130 120 110 100 90 80 70 60 50 40 30 20 10 0
f1 (ppm)

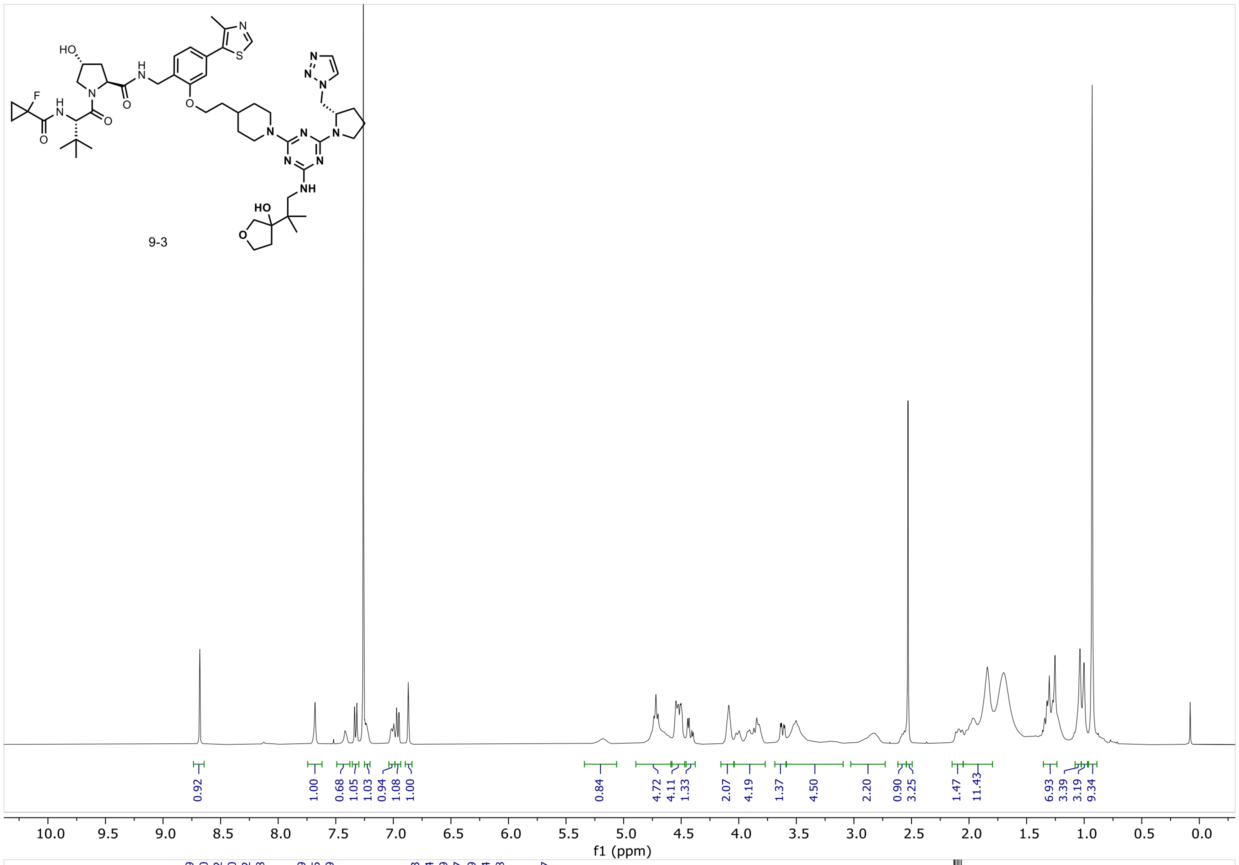
9-3
0.92
1.00
0.68
1.05
1.03
0.94
1.08
1.00
0.84
4.72
4.11
1.33
2.07
4.19
1.37
4.50
2.20
0.90
3.25
1.47
11.43
6.93
3.39
3.19
9.34
10.0 9.5 9.0 8.5 8.0 7.5 7.0 6.5 6.0 5.5 5.0 4.5 4.0 3.5 3.0 2.5 2.0 1.5 1.0 0.5 0.0
f1 (ppm)

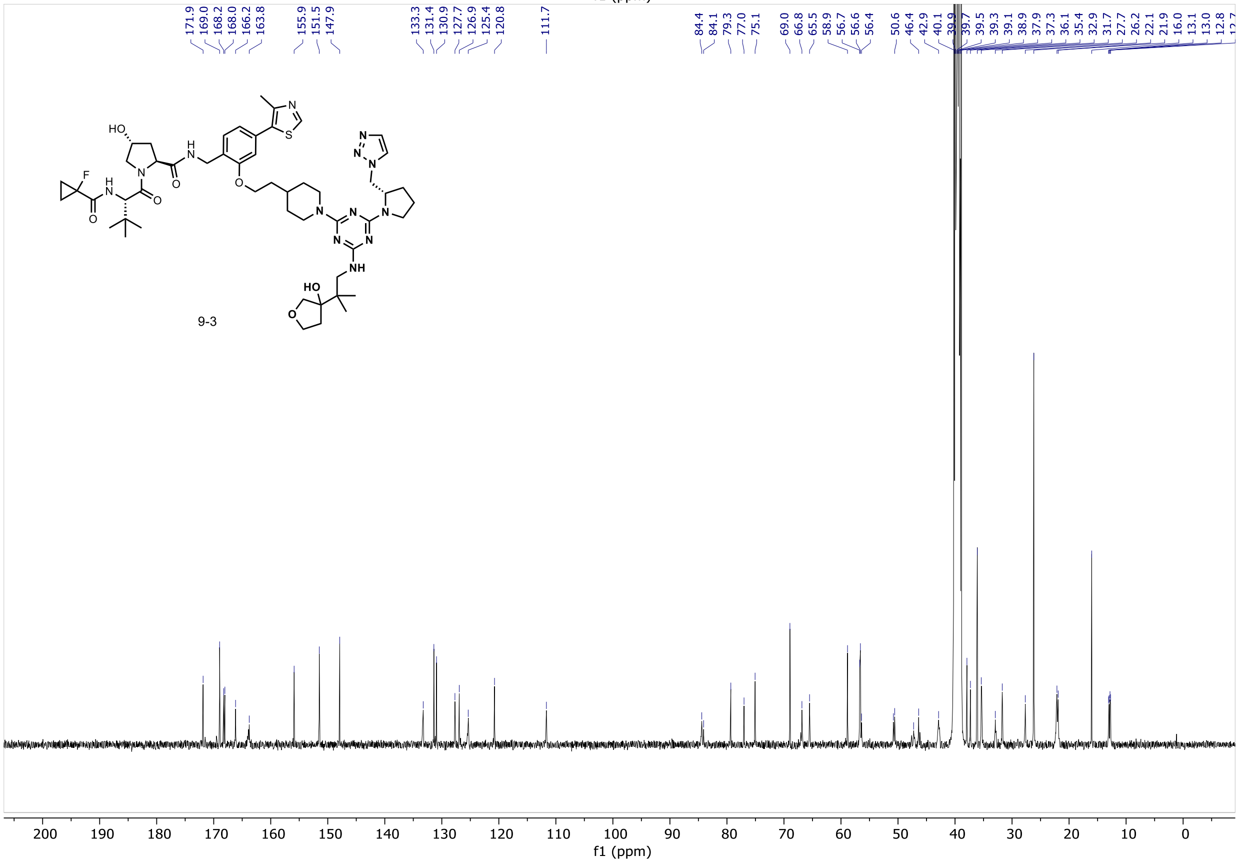
171.9
169.0
168.2
168.0
166.2
163.8
155.9
151.5
147.9
133.3
131.4
130.9
127.7
126.9
125.4
120.8
111.7
84.4
84.1
79.3
77.0
75.1
69.0
66.8
65.5
58.9
56.7
56.6
56.4
50.6
46.4
42.9
40.1
39.9
39.7
39.5
39.3
39.1
38.9
37.9
37.3
36.1
35.4
32.9
31.7
27.7
26.2
22.1
21.9
16.0
13.1
13.0
12.8
9-3
200 190 180 170 160 150 140 130 120 110 100 90 80 70 60 50 40 30 20 10 0
f1 (ppm)

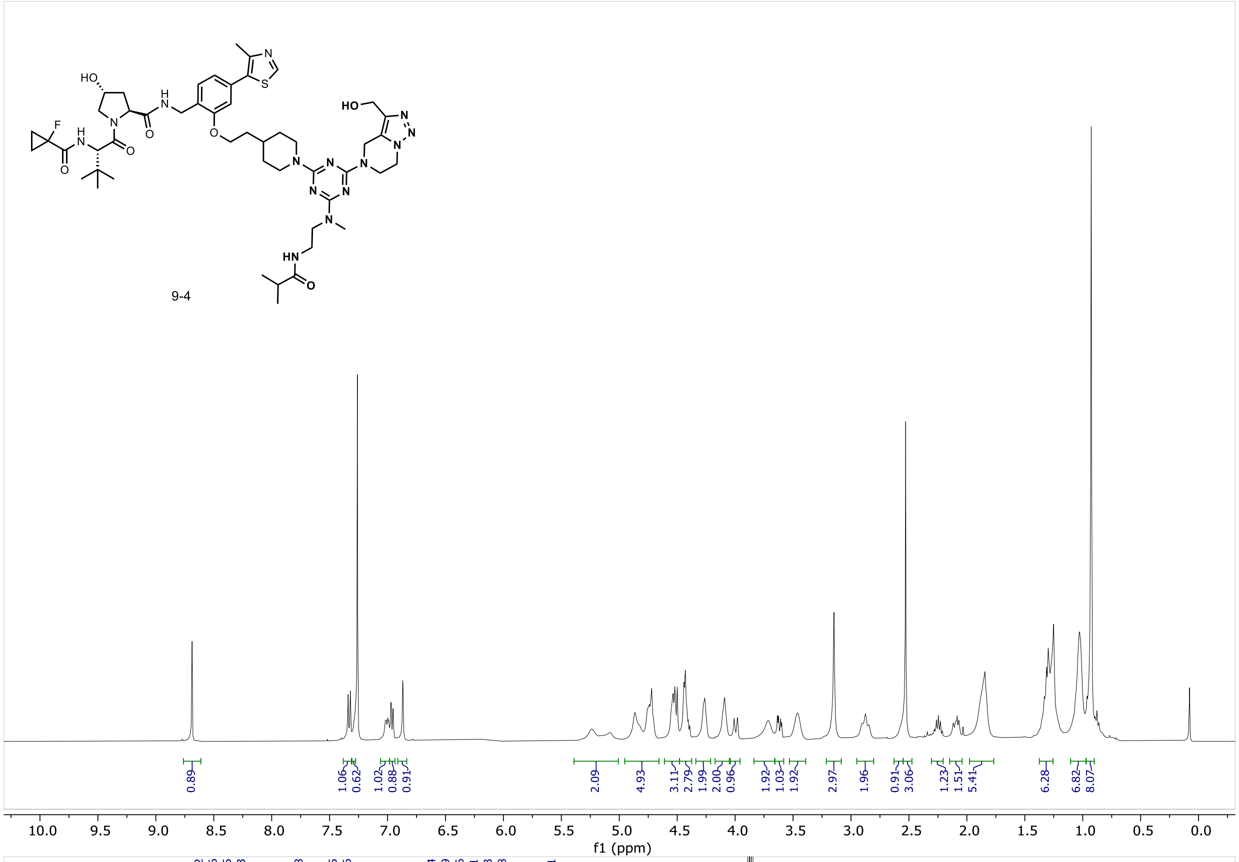
9-4
f1 (ppm)

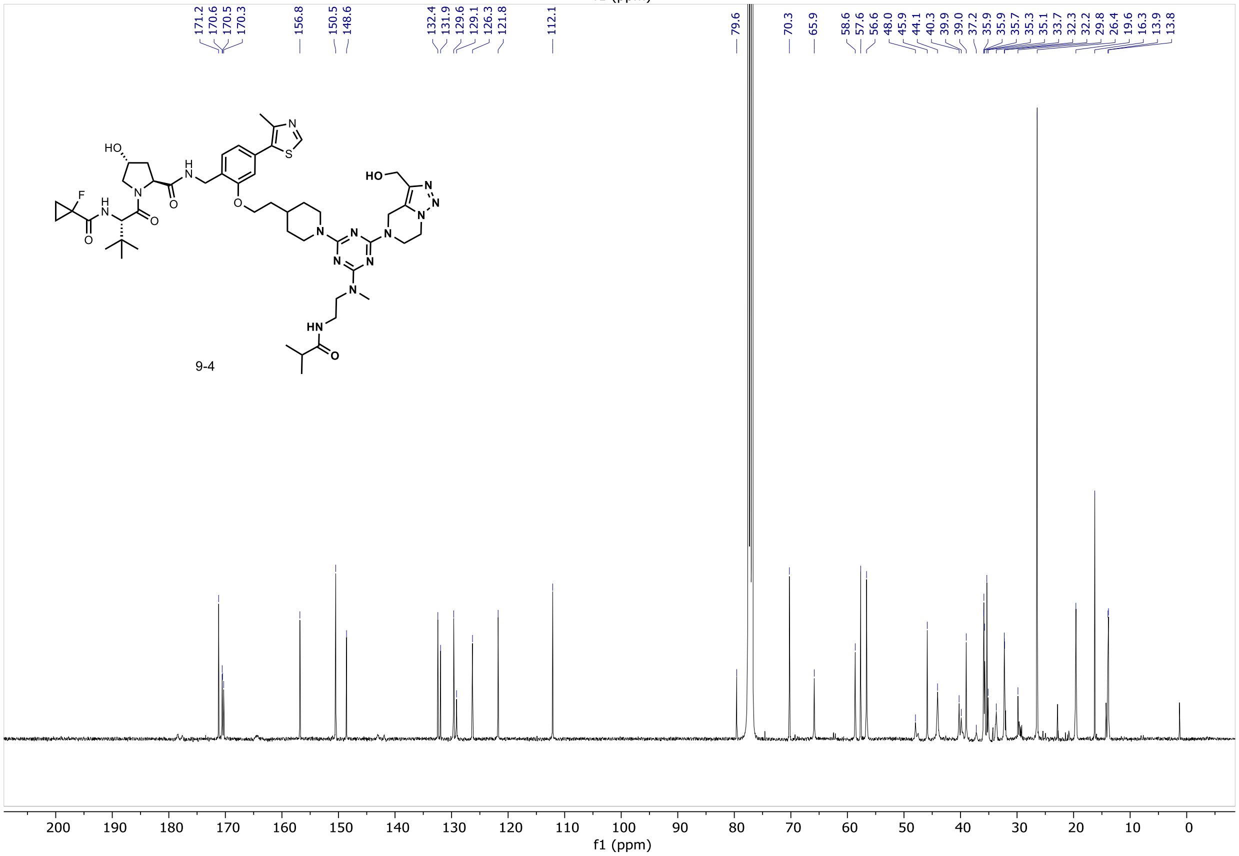
9-4
f1 (ppm)

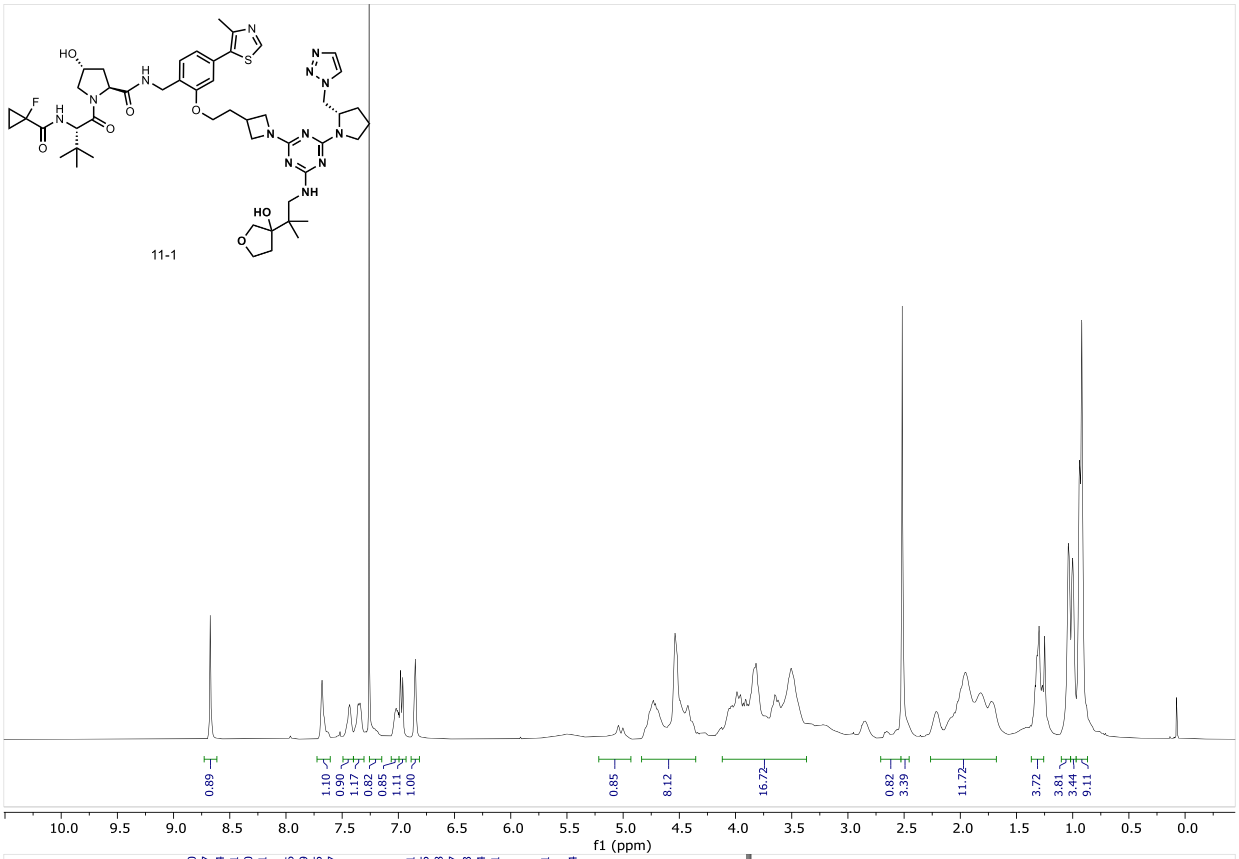
11-1
0.89
1.10
0.90
1.17
0.82
0.85
1.11
1.00
0.85
8.12
16.72
0.82
3.39
11.72
3.72
3.81
3.44
9.11
f1 (ppm)

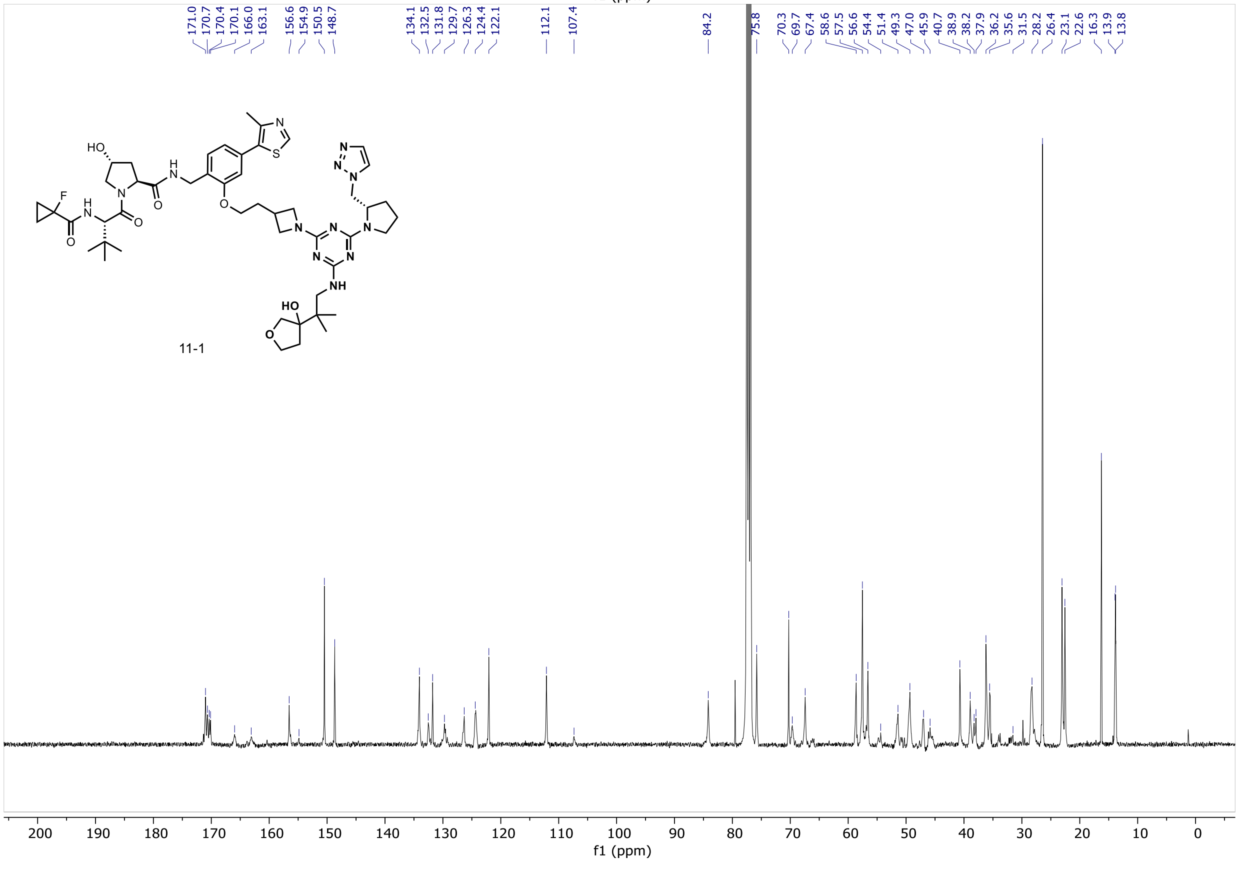
171.0
170.7
170.4
170.1
166.0
163.1
156.6
154.9
150.5
148.7
134.1
132.5
131.8
129.7
126.3
124.4
122.1
112.1
107.4
84.2
75.8
70.3
69.7
67.4
58.6
57.5
56.6
54.4
51.4
49.3
47.0
45.9
40.7
38.9
38.2
37.9
36.2
35.6
31.5
28.2
26.4
23.1
22.6
16.3
13.9
13.8
11-1
f1 (ppm)

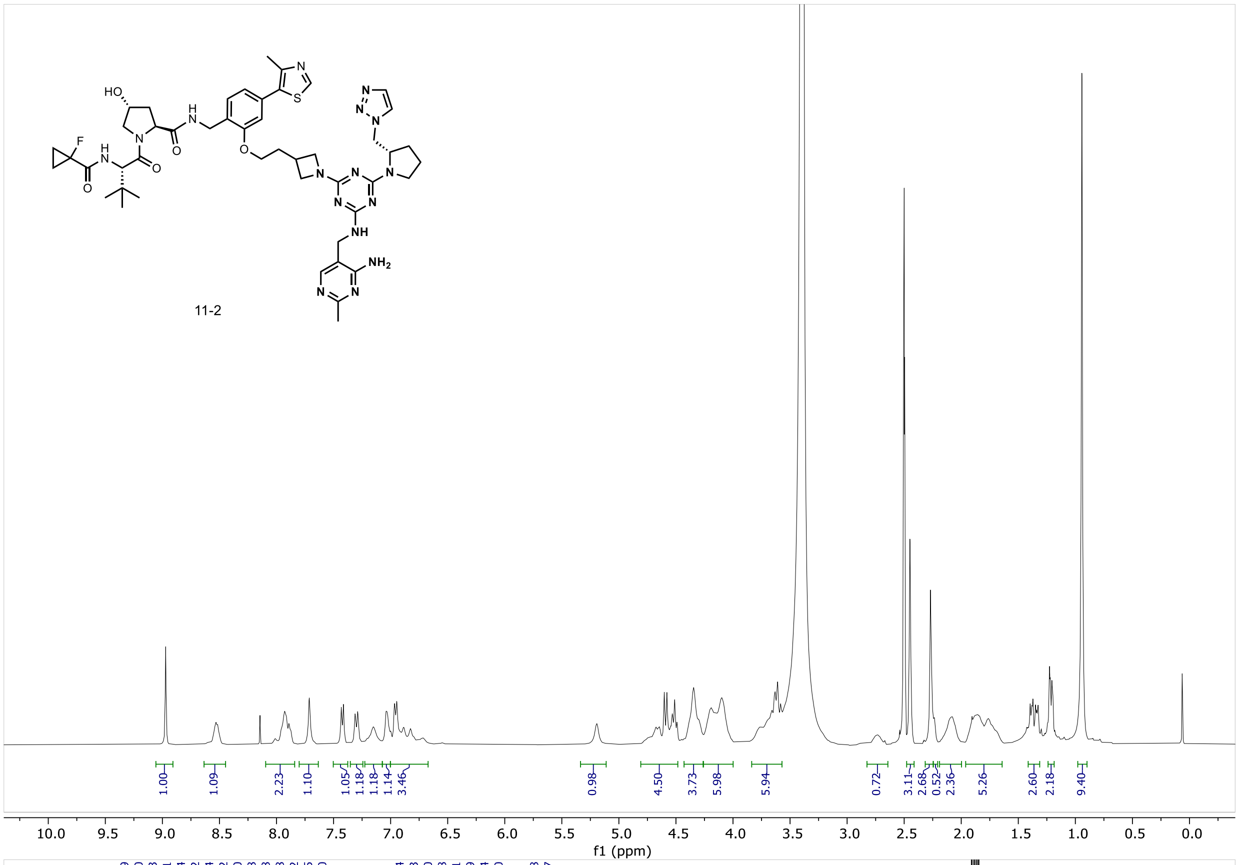
11-2
1.00 1.09 2.23 1.10 1.05 1.18 1.18 1.14 3.46 0.98 4.50 3.73 5.98 5.94 0.72 3.11 2.68 0.52 2.36 5.26 2.60 2.18 9.40
f1 (ppm)

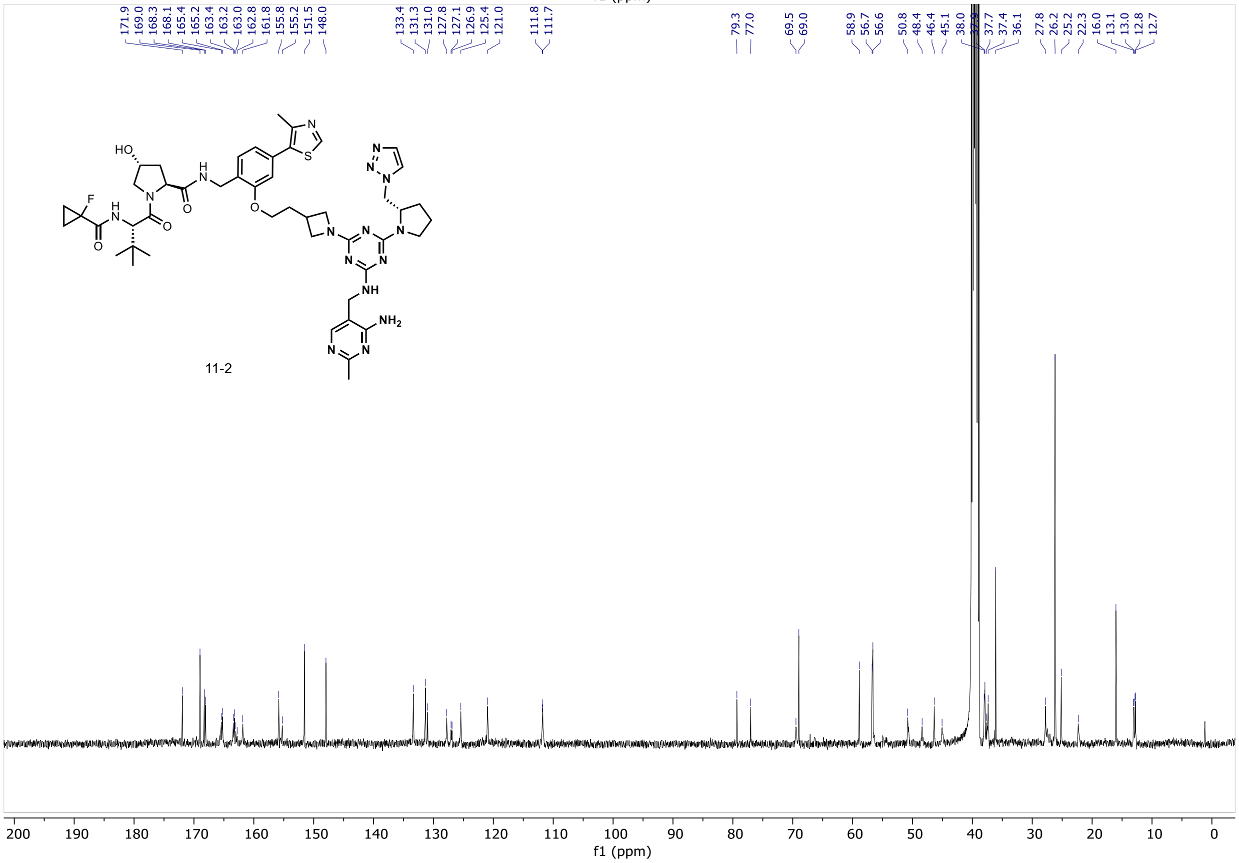
171.9 169.0 168.3 168.1 165.4 165.2 163.4 163.2 163.0 162.8 161.8 155.8 155.2 151.5 148.0 133.4 131.3 131.0 127.8 127.1 126.9 125.4 121.0 111.8 111.7 79.3 77.0 69.5 69.0 58.9 56.7 56.6 50.8 48.4 46.4 45.1 38.0 37.7 37.4 36.1 27.8 26.2 25.2 22.3 16.0 13.1 13.0 12.8 12.7
11-2
f1 (ppm)

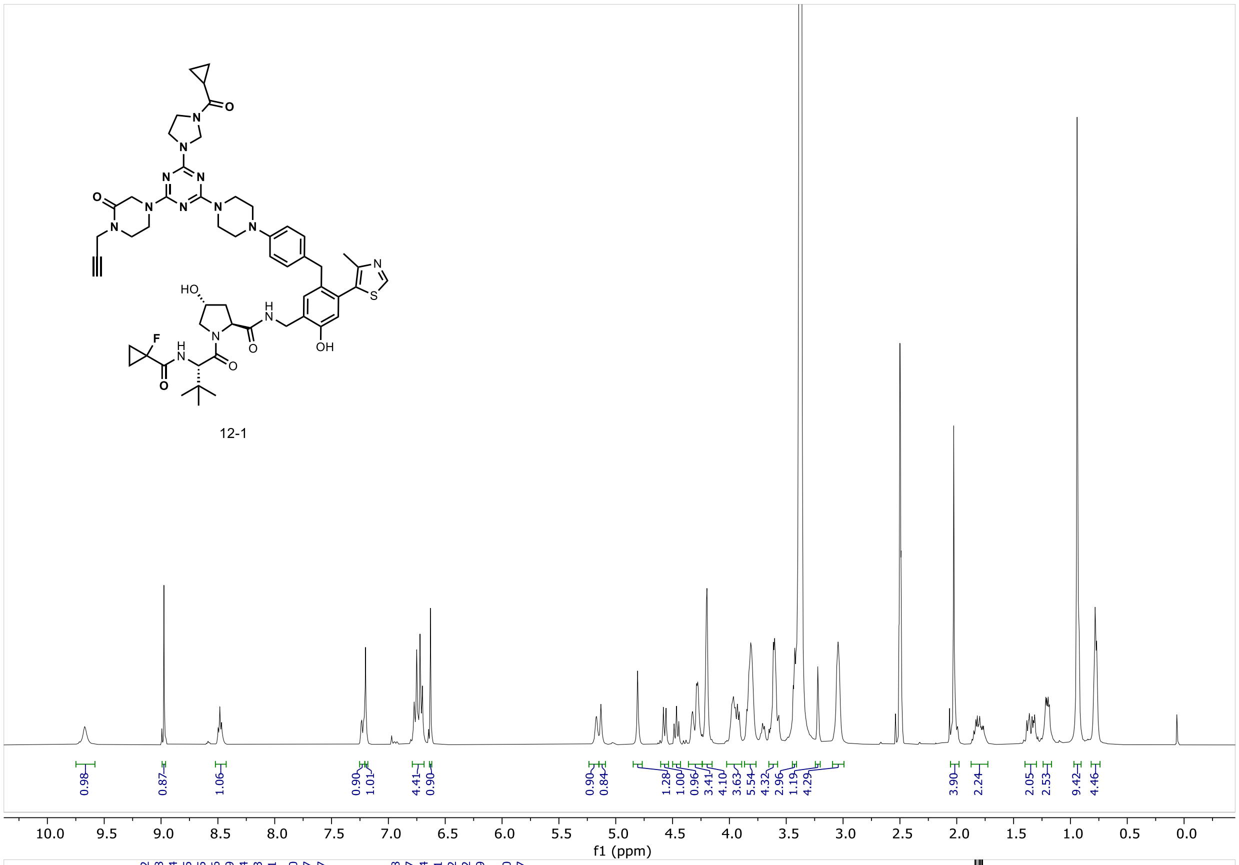
12-1
0.98
0.87
1.06
0.90
1.01
4.41
0.90
0.90
0.84
1.28
1.00
0.96
3.41
4.10
3.63
5.54
4.32
2.96
1.19
4.29
3.90
2.24
2.05
2.53
9.42
4.46
10.0 9.5 9.0 8.5 8.0 7.5 7.0 6.5 6.0 5.5 5.0 4.5 4.0 3.5 3.0 2.5 2.0 1.5 1.0 0.5 0.0
f1 (ppm)

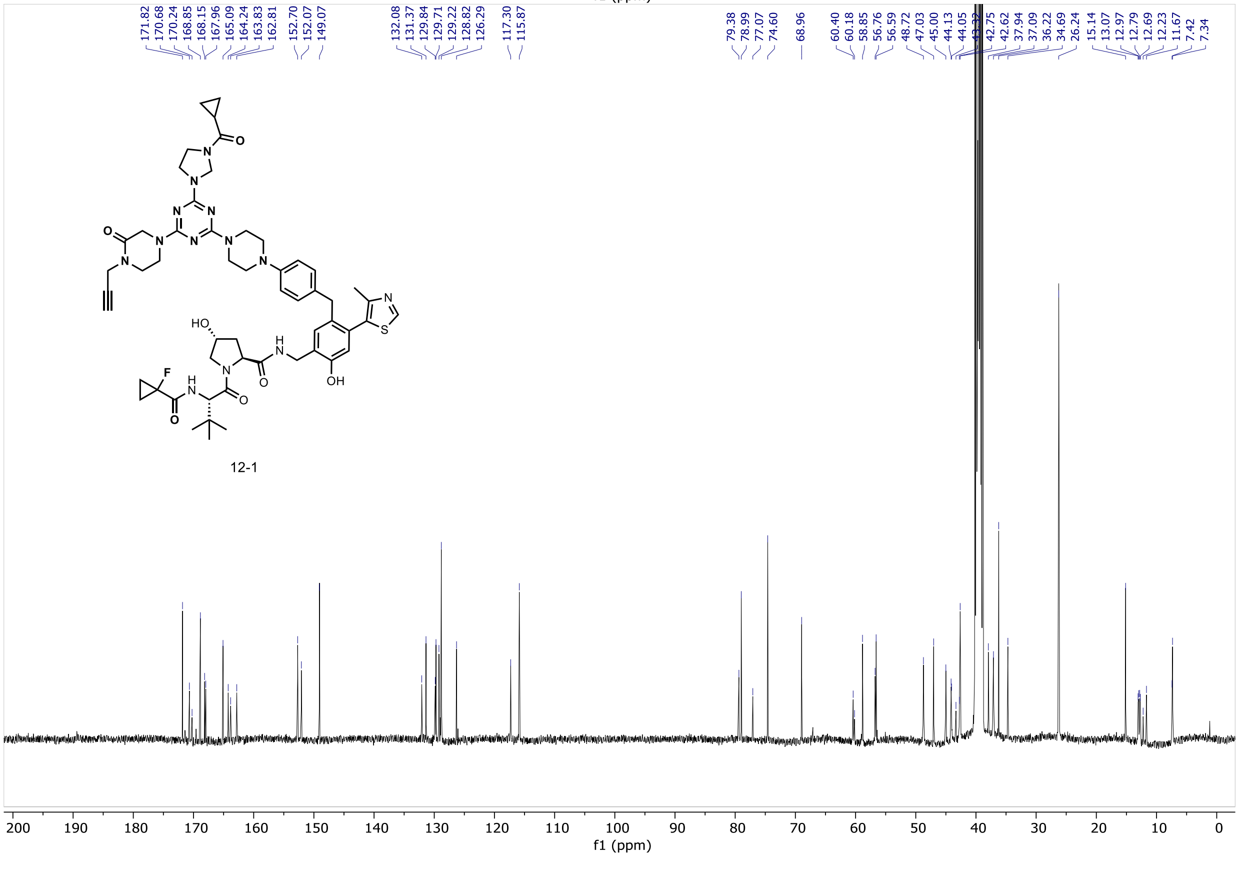
171.82
170.68
170.24
168.85
168.15
167.96
165.09
164.24
163.83
162.81
152.70
152.07
149.07
132.08
131.37
129.84
129.71
129.22
128.82
126.29
117.30
115.87
79.38
78.99
77.07
74.60
68.96
60.40
60.18
58.85
56.76
56.59
48.72
47.03
45.00
44.13
44.05
42.75
42.62
37.94
37.09
36.22
34.69
26.24
15.14
13.07
12.97
12.79
12.69
12.23
11.67
7.42
7.34
12-1
200 190 180 170 160 150 140 130 120 110 100 90 80 70 60 50 40 30 20 10 0
f1 (ppm)

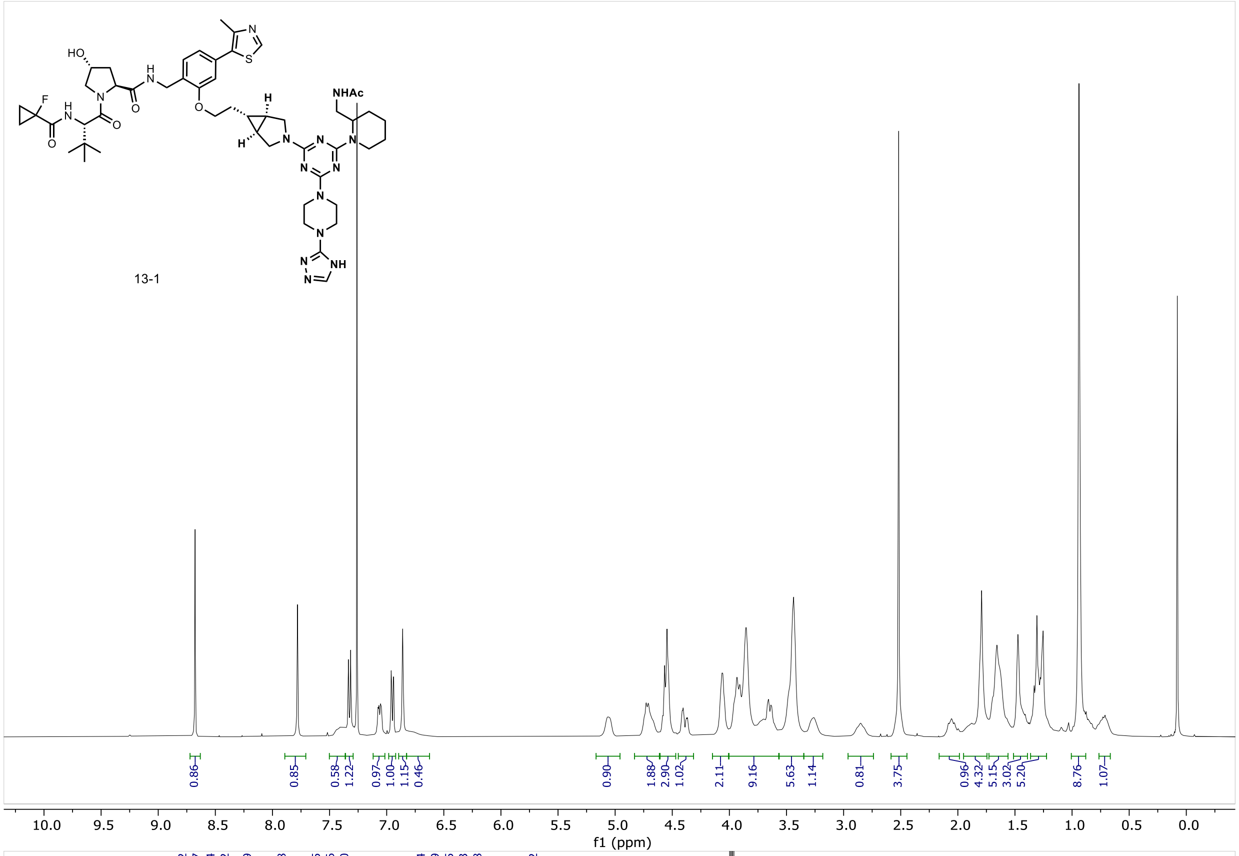
13-1
NHAc
HO
F
0.86 0.85 0.58 1.22 0.97 1.00 1.15 0.46 0.90 1.88 2.90 1.02 2.11 9.16 5.63 1.14 0.81 3.75 0.96 4.32 5.15 3.02 5.20 8.76 1.07
10.0 9.5 9.0 8.5 8.0 7.5 7.0 6.5 6.0 5.5 5.0 4.5 4.0 3.5 3.0 2.5 2.0 1.5 1.0 0.5 0.0
f1 (ppm)

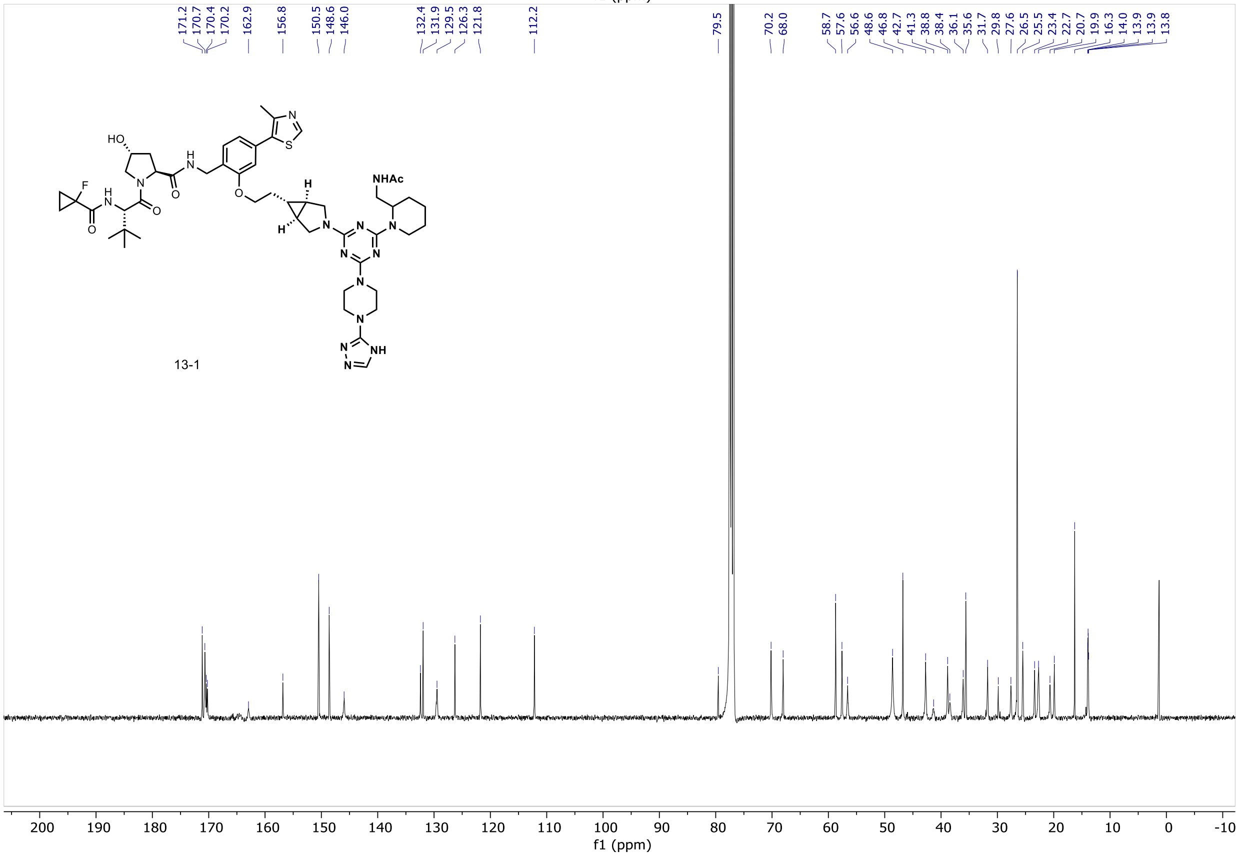
171.2 170.7 170.4 170.2 162.9 156.8 150.5 148.6 146.0 132.4 131.9 129.5 126.3 121.8 112.2 79.5 70.2 68.0 58.7 57.6 56.6 48.6 46.8 42.7 41.3 38.8 38.4 36.1 35.6 31.7 29.8 27.6 26.5 25.5 23.4 22.7 20.7 19.9 16.3 14.0 13.9 13.9 13.8
13-1
NHAc
HO
F
200 190 180 170 160 150 140 130 120 110 100 90 80 70 60 50 40 30 20 10 0 -10
f1 (ppm)

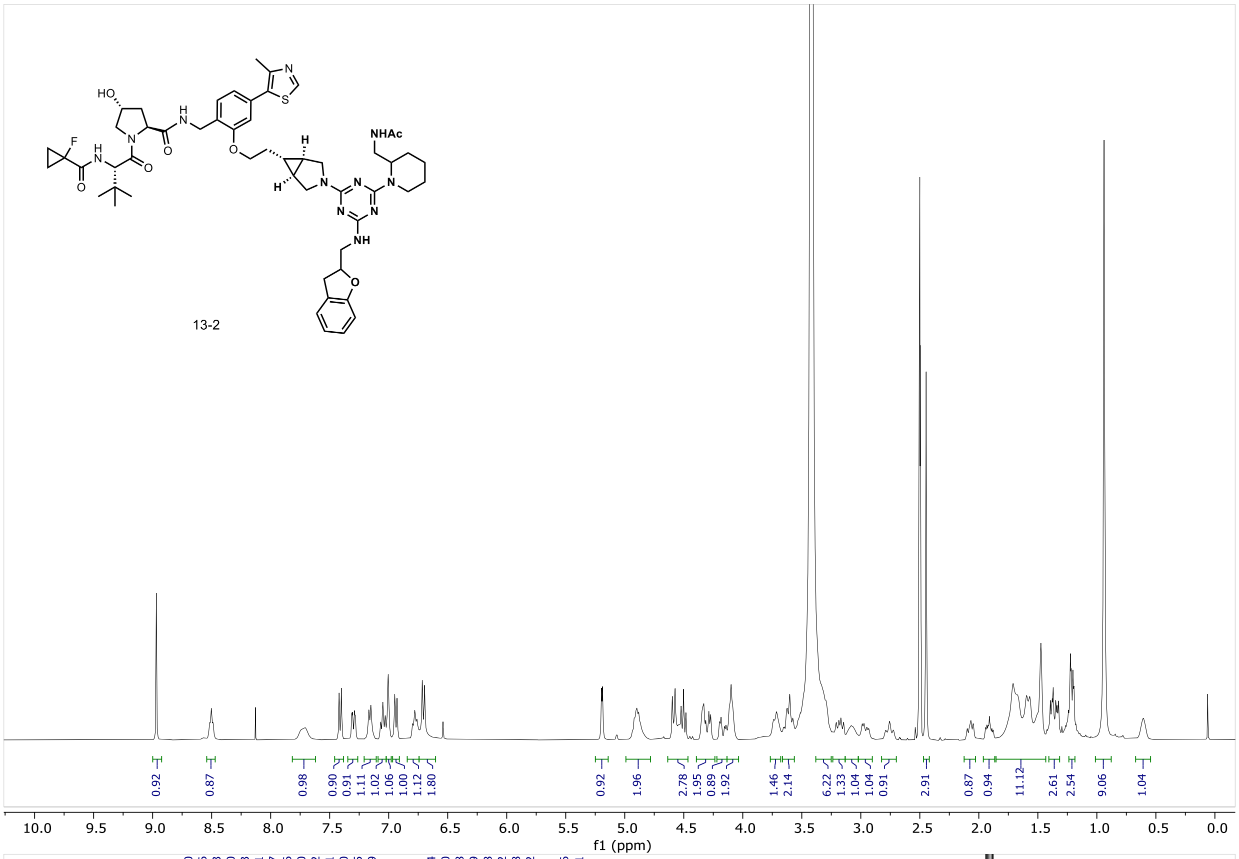
HO
NHAc
13-2
0.92
0.87
0.98
0.90
0.91
1.11
1.02
1.06
1.00
1.12
1.80
0.92
1.96
2.78
1.95
0.89
1.92
1.46
2.14
6.22
1.33
1.04
1.04
0.91
2.91
0.87
0.94
11.12
2.61
2.54
9.06
1.04
10.0 9.5 9.0 8.5 8.0 7.5 7.0 6.5 6.0 5.5 5.0 4.5 4.0 3.5 3.0 2.5 2.0 1.5 1.0 0.5 0.0
f1 (ppm)

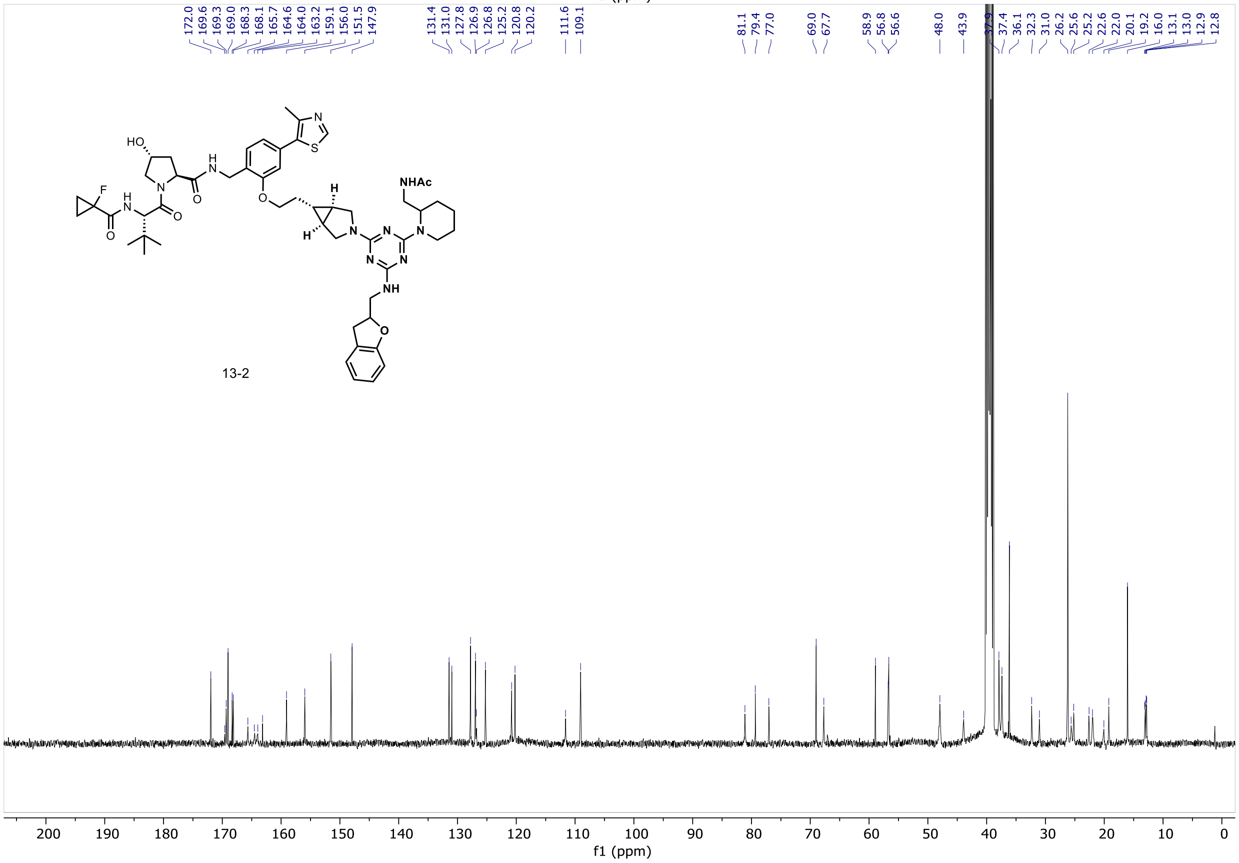
172.0
169.6
169.3
169.0
168.3
168.1
165.7
164.6
164.0
163.2
159.1
156.0
151.5
147.9
131.4
131.0
127.8
126.9
126.8
125.2
120.8
120.2
111.6
109.1
81.1
79.4
77.0
69.0
67.7
58.9
56.8
56.6
48.0
43.9
37.4
36.1
32.3
31.0
26.2
25.6
25.2
22.6
22.0
20.1
19.2
16.0
13.1
13.0
12.9
12.8
HO
NHAc
13-2
200 190 180 170 160 150 140 130 120 110 100 90 80 70 60 50 40 30 20 10 0
f1 (ppm)

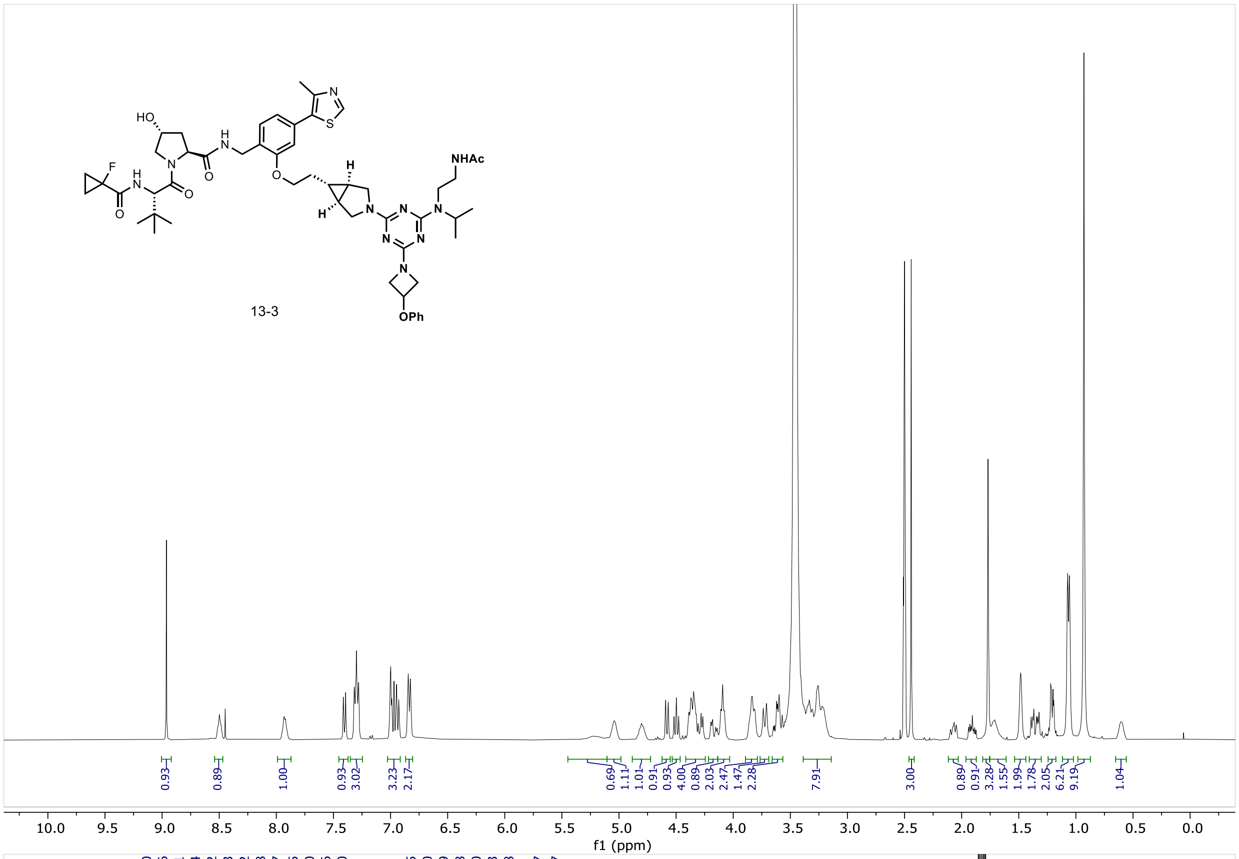

NHAc
OPh
13-3
0.93
0.89
1.00
0.93
3.02
3.23
2.17
0.69
1.11
1.01
0.91
0.93
4.00
0.89
2.03
2.47
1.47
2.28
7.91
3.00
0.89
0.91
3.28
1.55
1.99
1.78
2.05
6.21
9.19
1.04
f1 (ppm)

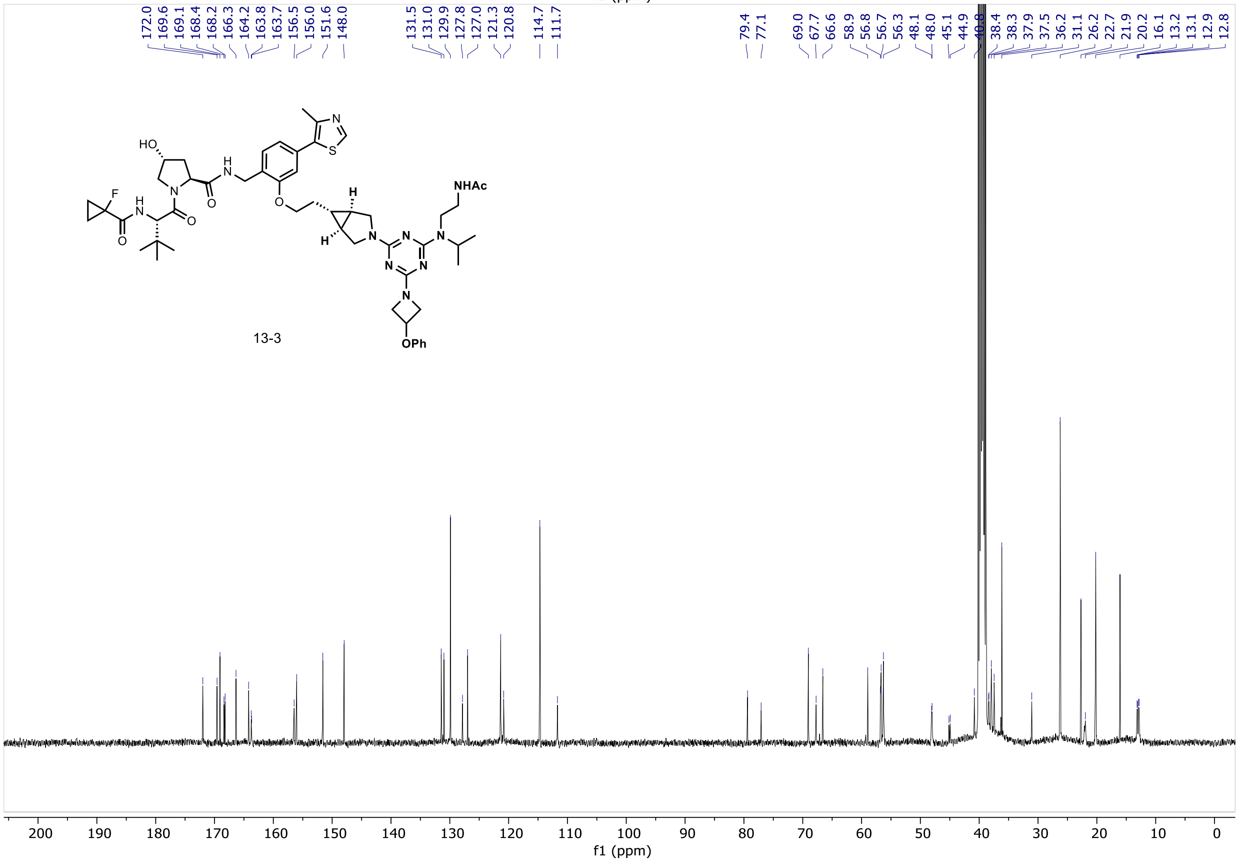

172.0
169.6
169.1
168.4
168.2
166.3
164.2
163.8
163.7
156.5
156.0
151.6
148.0
131.5
131.0
129.9
127.8
127.0
121.3
120.8
114.7
111.7
79.4
77.1
69.0
67.7
66.6
58.9
56.8
56.7
56.3
48.1
48.0
45.1
44.9
38.4
38.3
37.9
37.5
36.2
31.1
26.2
22.7
21.9
20.2
16.1
13.2
13.1
12.9
12.8
NHAc
OPh
13-3
f1 (ppm)

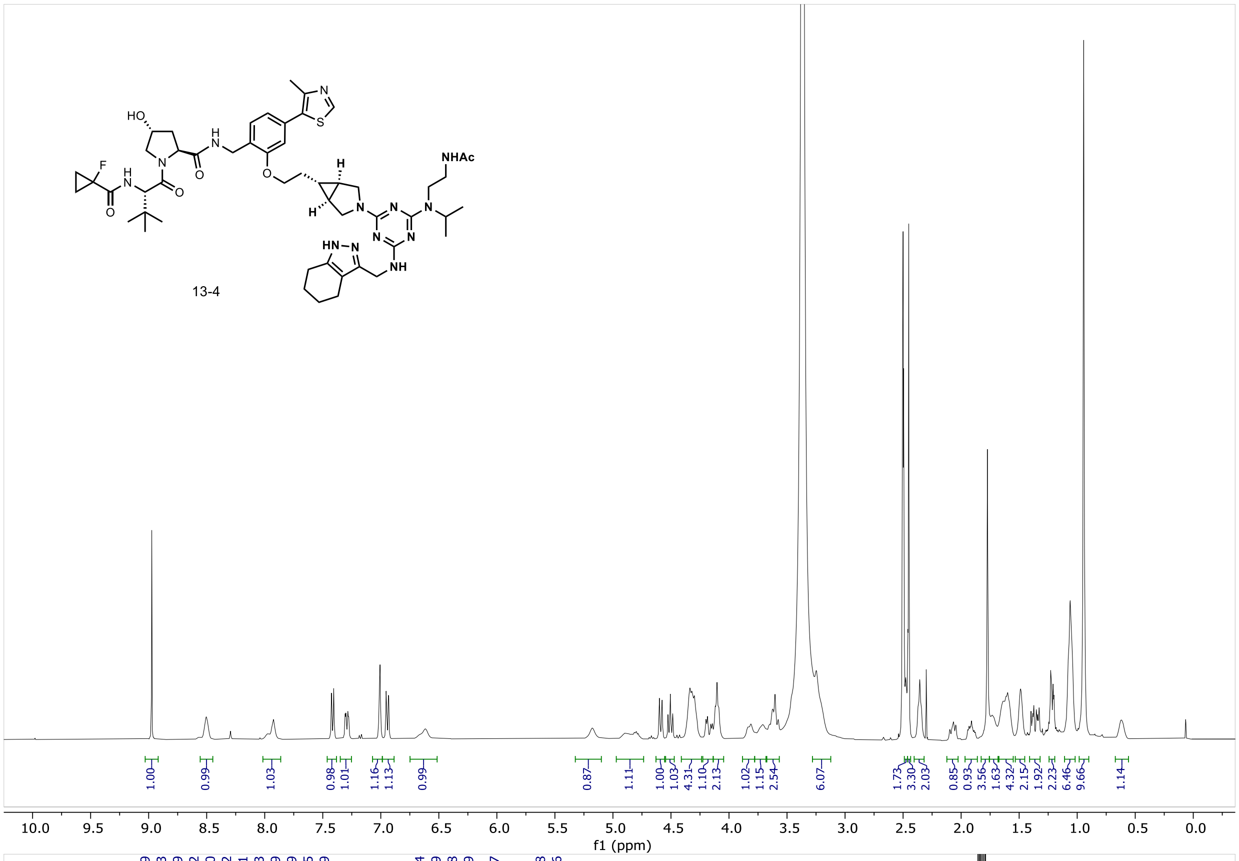
NHAc
HO
HN
HN
HN–N
NH
13-4
1.00
0.99
1.03
0.98
1.01
1.16
1.13
0.99
0.87
1.11
1.00
1.03
4.31
1.10
2.13
1.02
1.15
2.54
6.07
1.73
3.30
2.03
0.85
0.93
3.56
1.63
4.32
2.15
1.92
2.23
6.46
9.66
1.14
10.0 9.5 9.0 8.5 8.0 7.5 7.0 6.5 6.0 5.5 5.0 4.5 4.0 3.5 3.0 2.5 2.0 1.5 1.0 0.5 0.0
f1 (ppm)

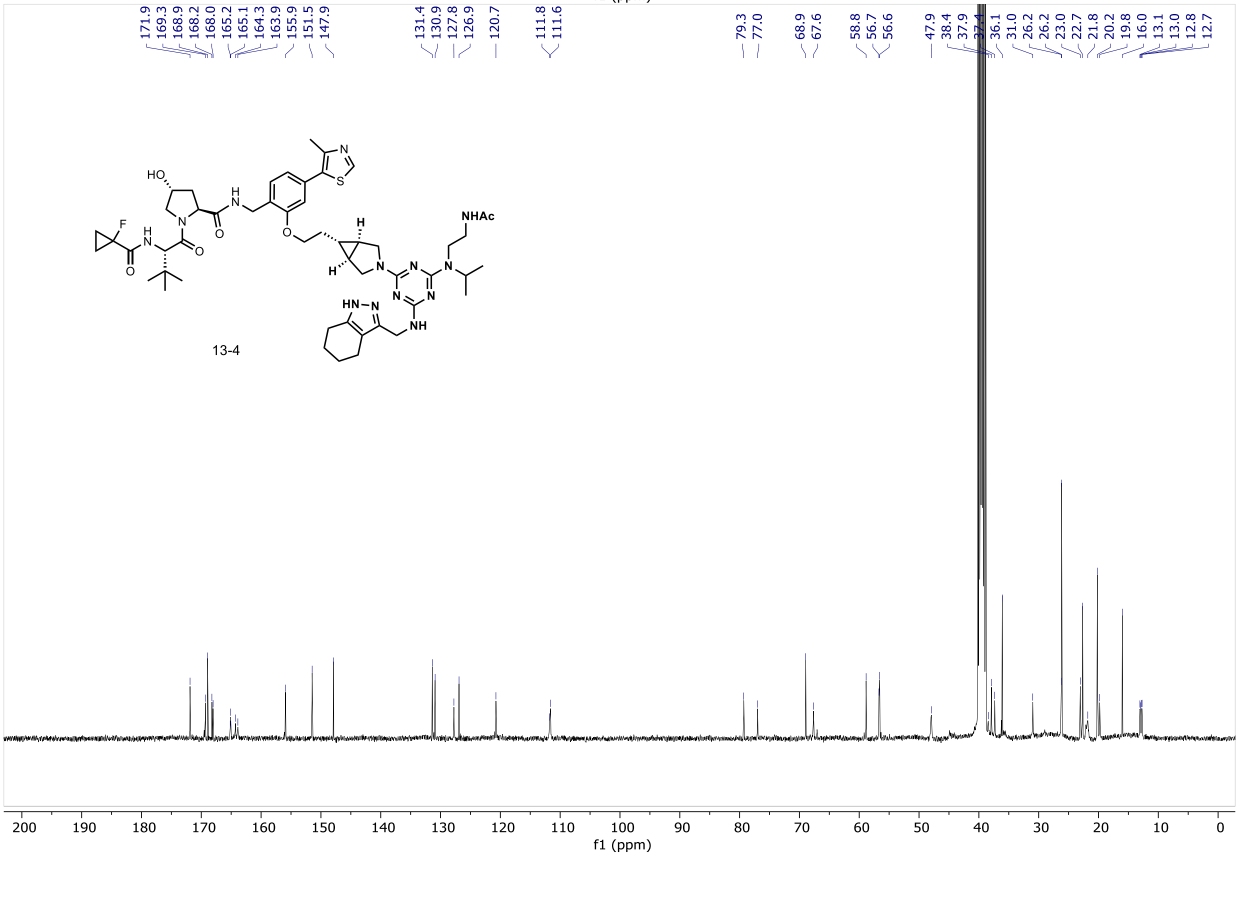
171.9
169.3
168.9
168.2
168.0
165.2
165.1
164.3
163.9
155.9
151.5
147.9
131.4
130.9
127.8
126.9
120.7
111.8
111.6
79.3
77.0
68.9
67.6
58.8
56.7
56.6
47.9
38.4
37.9
37.4
36.1
31.0
26.2
26.2
23.0
22.7
21.8
20.2
19.8
16.0
13.1
13.0
12.8
12.7
NHAc
HO
HN
HN–N
NH
13-4
200 190 180 170 160 150 140 130 120 110 100 90 80 70 60 50 40 30 20 10 0
f1 (ppm)

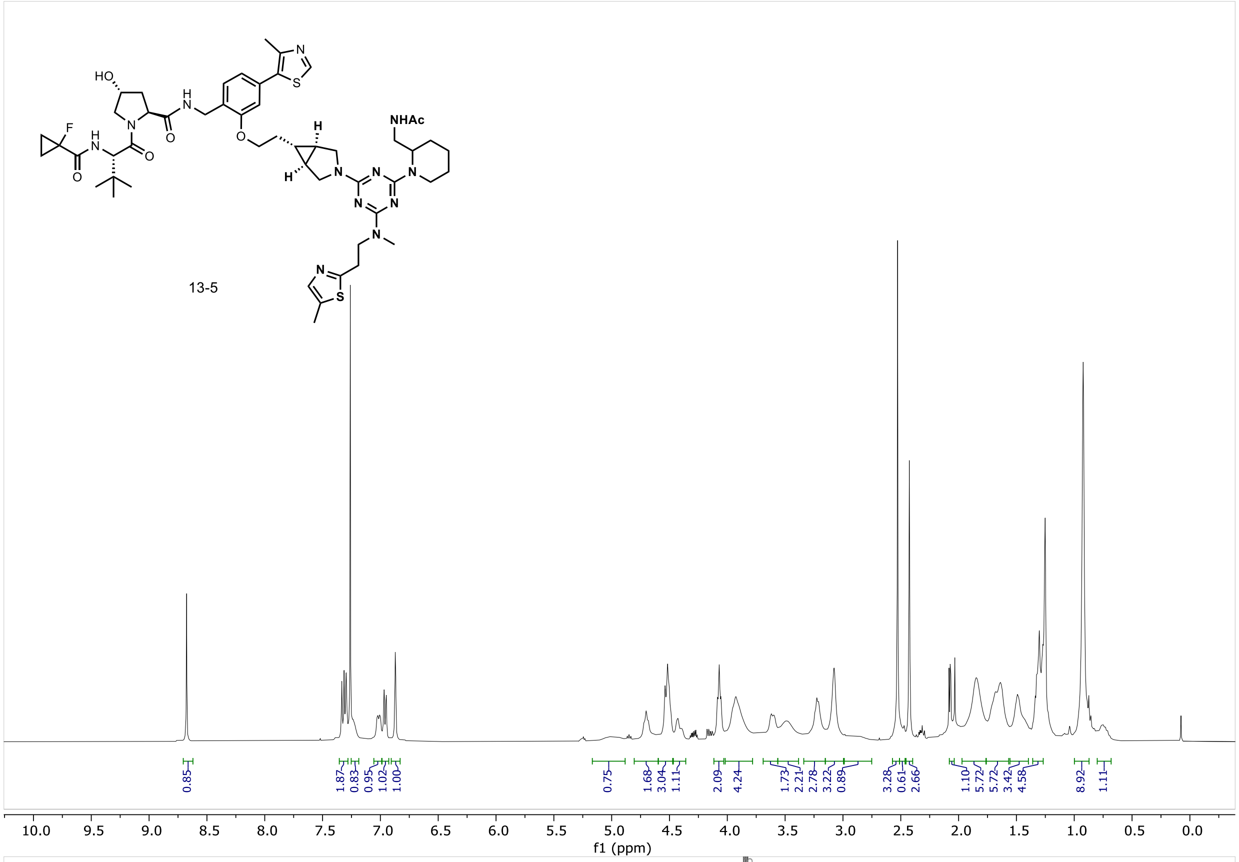
HO
F
NHAc
13-5
0.85
1.87
0.83
0.95
1.02
1.00
0.75
1.68
3.04
1.11
2.09
4.24
1.73
2.21
2.78
3.22
0.89
3.28
0.61
2.66
1.10
5.72
5.72
3.42
4.58
8.92
1.11
f1 (ppm)

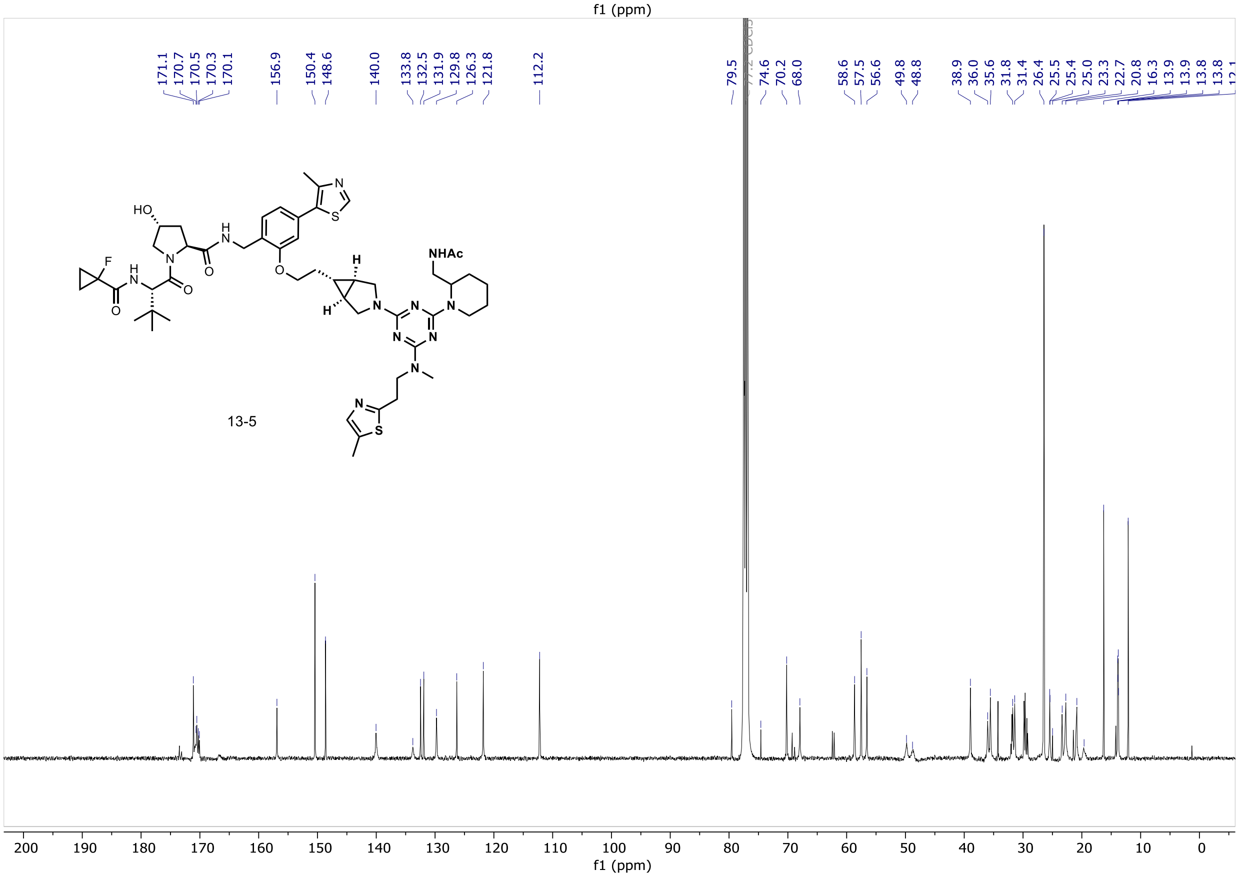
171.1
170.7
170.5
170.3
170.1
156.9
150.4
148.6
140.0
133.8
132.5
131.9
129.8
126.3
121.8
112.2
79.5
74.6
70.2
68.0
58.6
57.5
56.6
49.8
48.8
38.9
36.0
35.6
31.8
31.4
26.4
25.5
25.4
25.0
23.3
22.7
20.8
16.3
13.9
13.9
13.8
13.8
HO
F
NHAc
13-5
f1 (ppm)

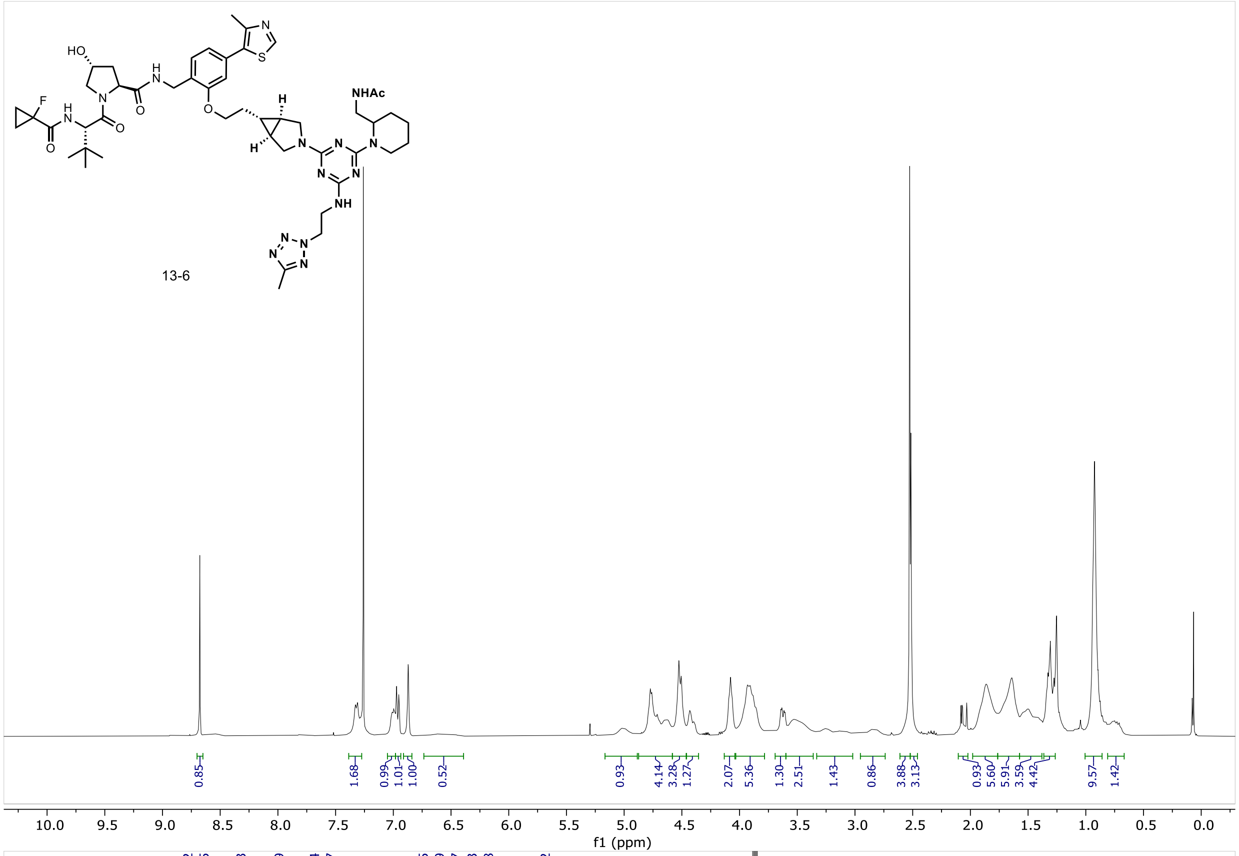
13-6
0.85
1.68
0.99
1.01
1.00
0.52
0.93
4.14
3.28
1.27
2.07
5.36
1.30
2.51
1.43
0.86
3.88
3.13
0.93
5.60
5.91
3.59
4.42
9.57
1.42
f1 (ppm)

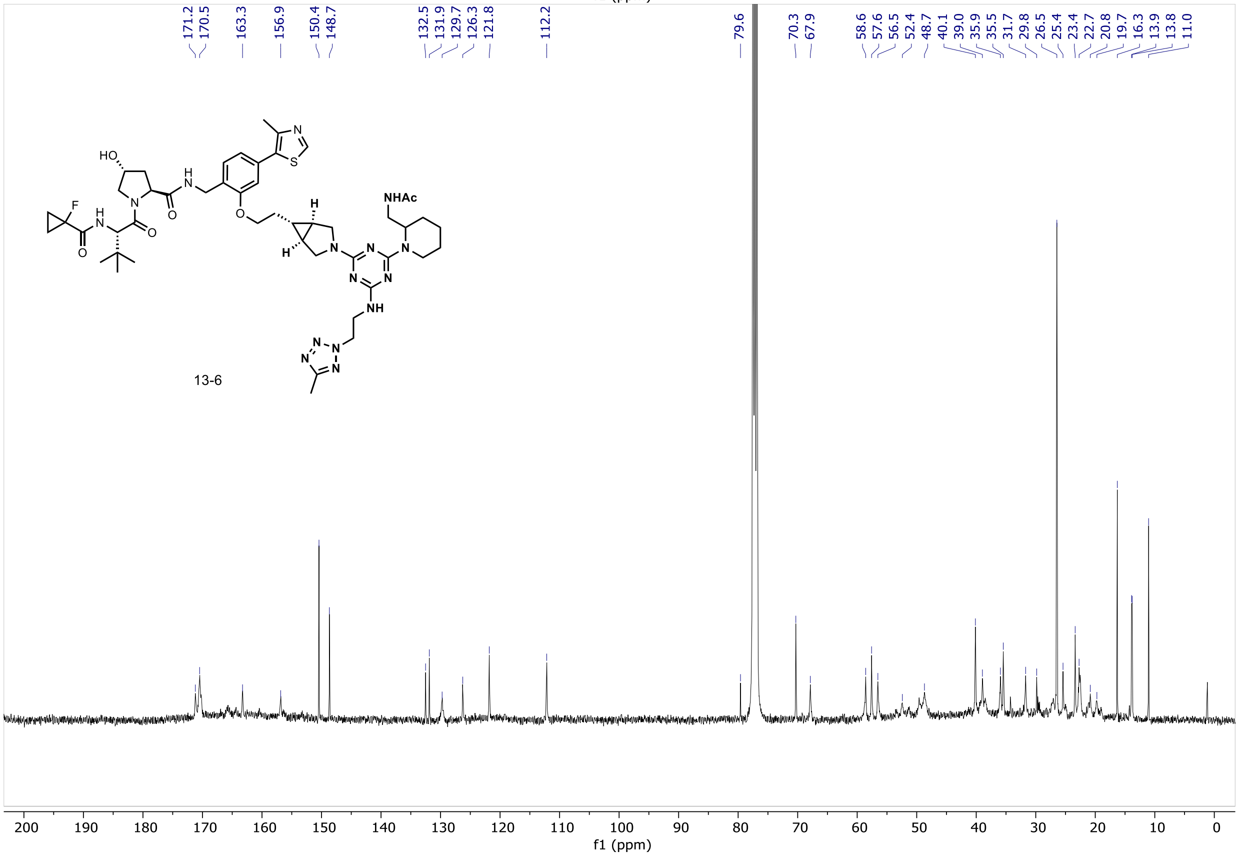
171.2
170.5
163.3
156.9
150.4
148.7
132.5
131.9
129.7
126.3
121.8
112.2
79.6
70.3
67.9
58.6
57.6
56.5
52.4
48.7
40.1
39.0
35.9
35.5
31.7
29.8
26.5
25.4
23.4
22.7
20.8
19.7
16.3
13.9
13.8
11.0
13-6
f1 (ppm)

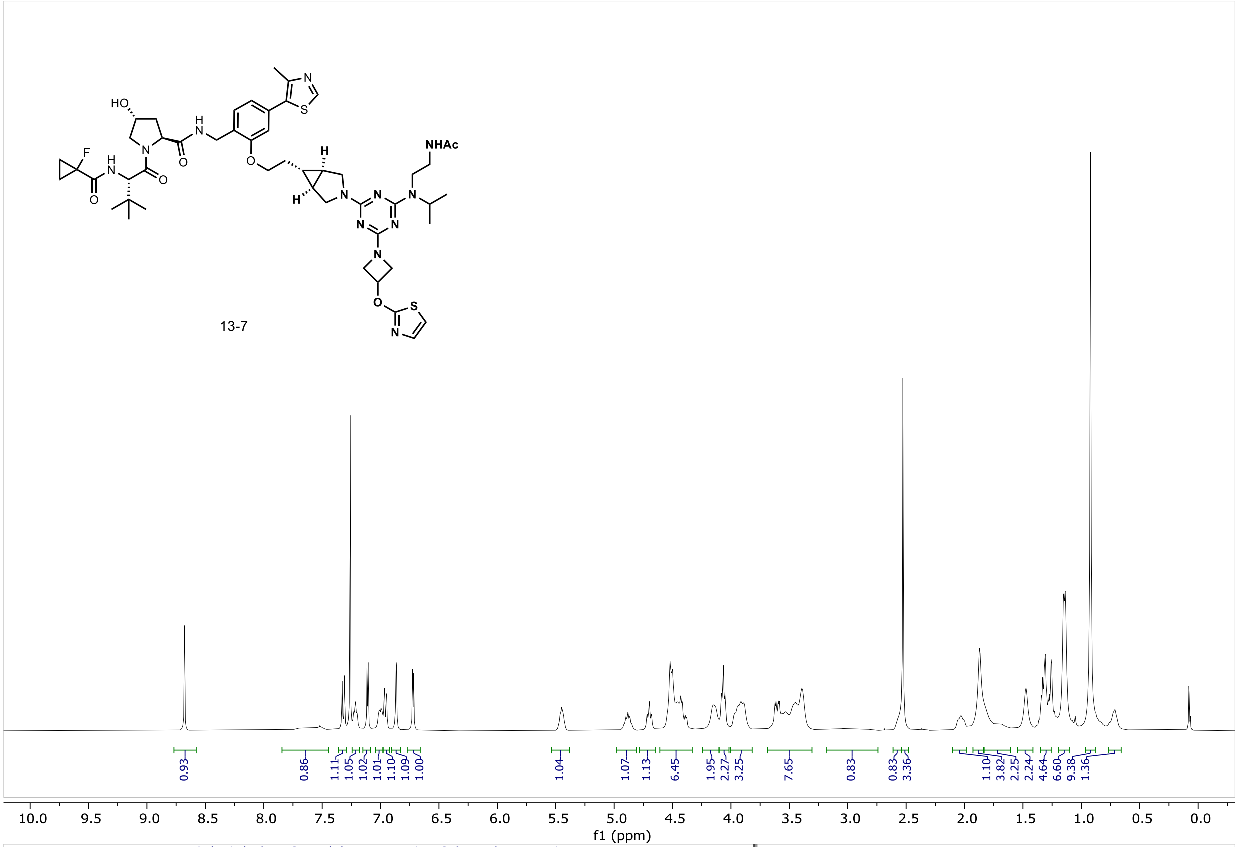
HO
NHAc
13-7
0.93
0.86
1.11
1.05
1.02
1.01
1.10
1.09
1.00
1.04
1.07
1.13
6.45
1.95
2.27
3.25
7.65
0.83
0.83
3.36
1.10
3.82
2.25
2.24
4.64
6.60
9.38
1.36
f1 (ppm)

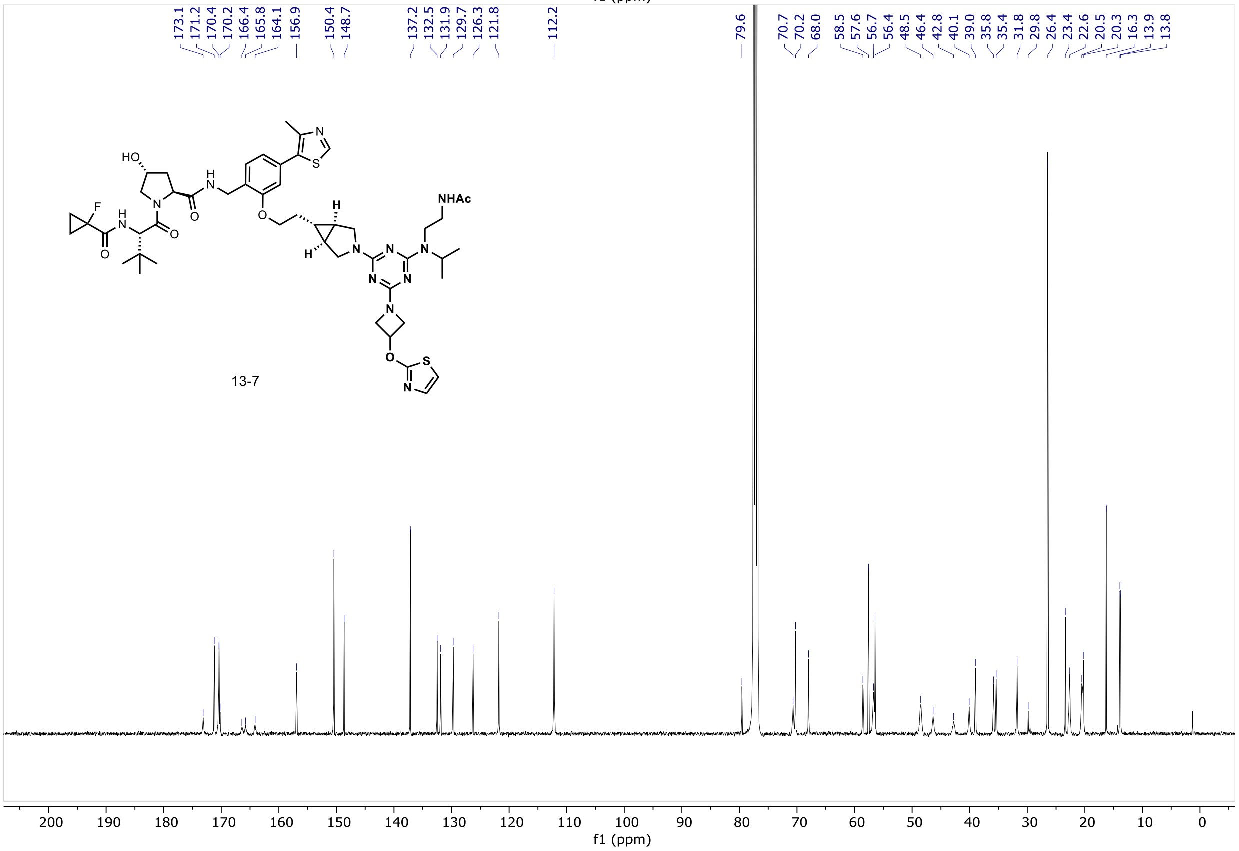
173.1
171.2
170.4
170.2
166.4
165.8
164.1
156.9
150.4
148.7
137.2
132.5
131.9
129.7
126.3
121.8
112.2
79.6
70.7
70.2
68.0
58.5
57.6
56.7
56.4
48.5
46.4
42.8
40.1
39.0
35.8
35.4
31.8
29.8
26.4
23.4
22.6
20.5
20.3
16.3
13.9
13.8
HO
NHAc
13-7
f1 (ppm)

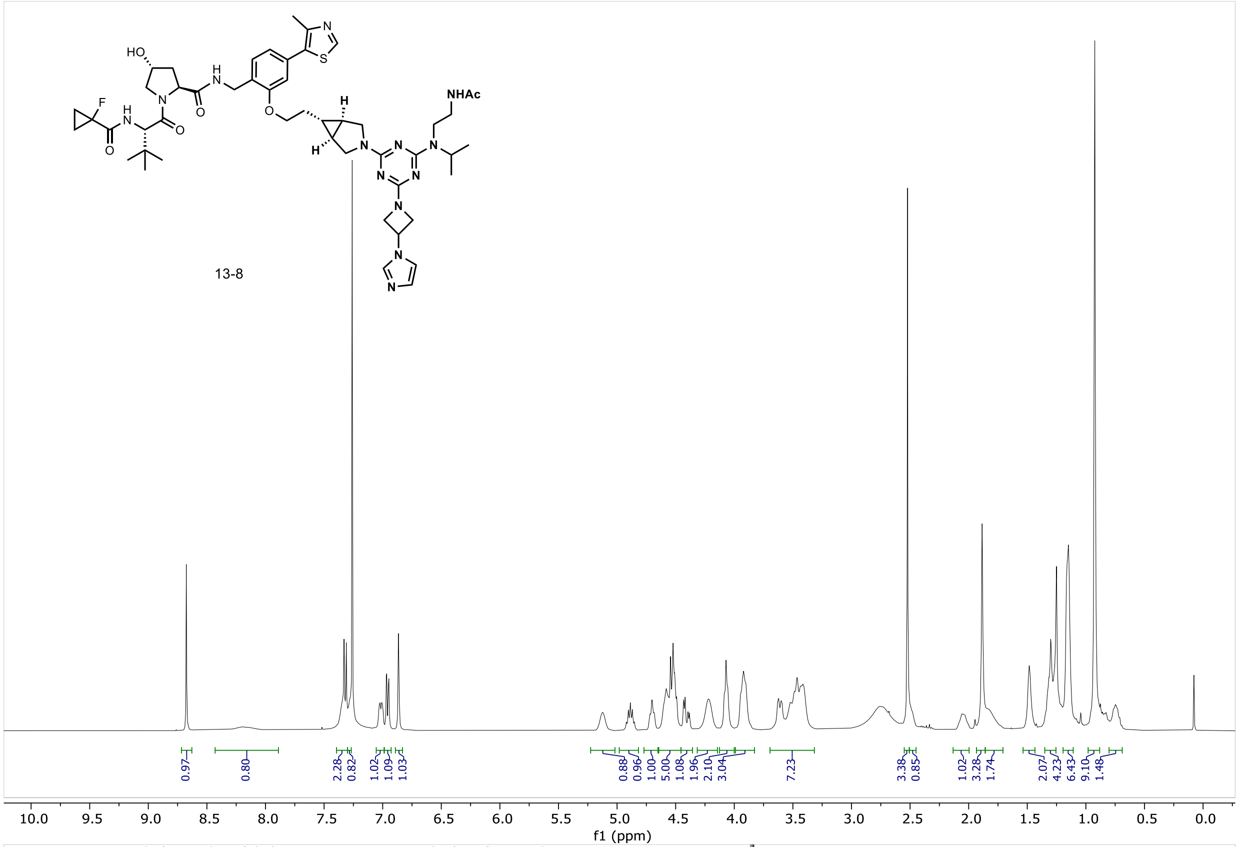

NHAc
HO
F
13-8
0.97
0.80
2.28
0.82
1.02
1.09
1.03
0.88
0.96
1.00
5.00
1.08
1.96
2.10
3.04
7.23
3.38
0.85
1.02
3.28
1.74
2.07
4.23
6.43
9.10
1.48
10.0 9.5 9.0 8.5 8.0 7.5 7.0 6.5 6.0 5.5 5.0 4.5 4.0 3.5 3.0 2.5 2.0 1.5 1.0 0.5 0.0
f1 (ppm)

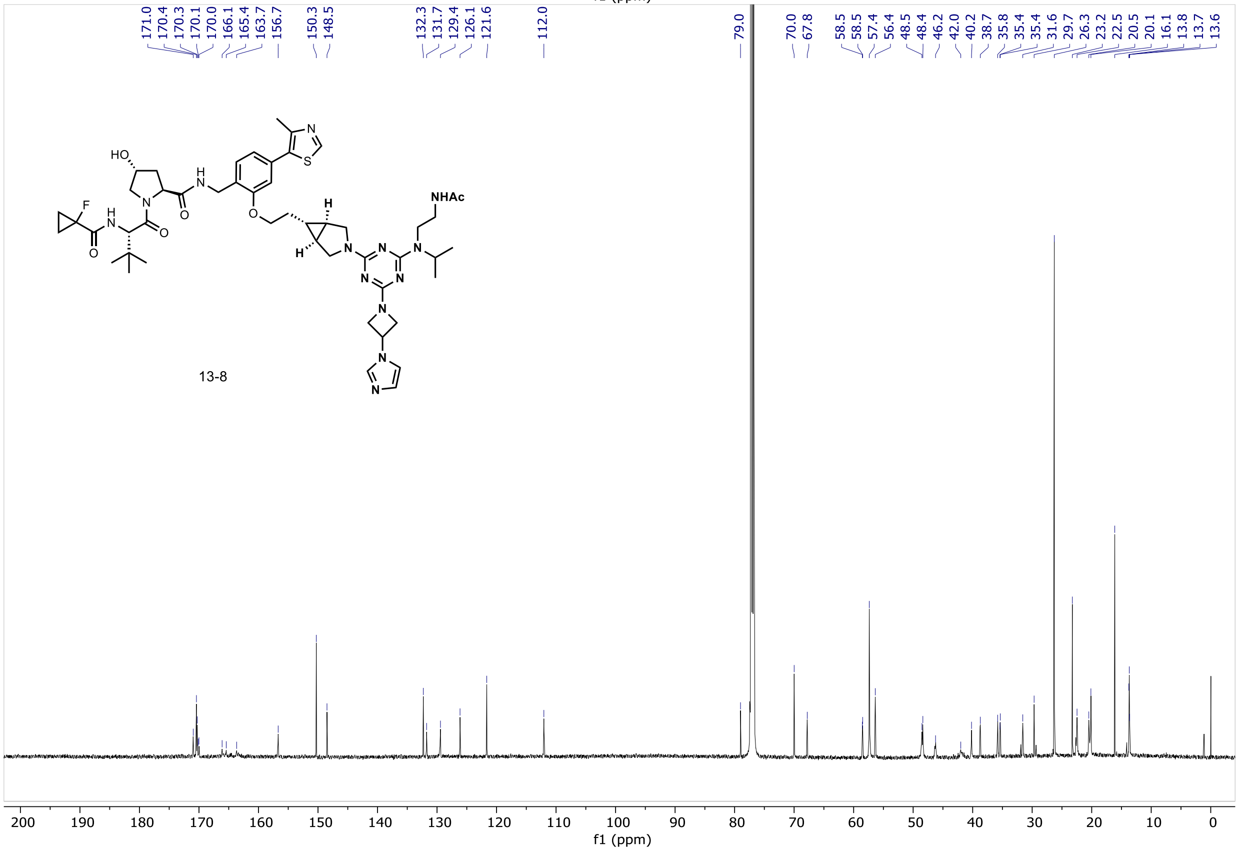

171.0
170.4
170.3
170.1
170.0
166.1
165.4
163.7
156.7
150.3
148.5
132.3
131.7
129.4
126.1
121.6
112.0
79.0
70.0
67.8
58.5
58.5
57.4
56.4
48.5
48.4
46.2
42.0
40.2
38.7
35.8
35.4
35.4
31.6
29.7
26.3
23.2
22.5
20.5
20.1
16.1
13.8
13.7
13.6
NHAc
HO
F
13-8
200 190 180 170 160 150 140 130 120 110 100 90 80 70 60 50 40 30 20 10 0
f1 (ppm)

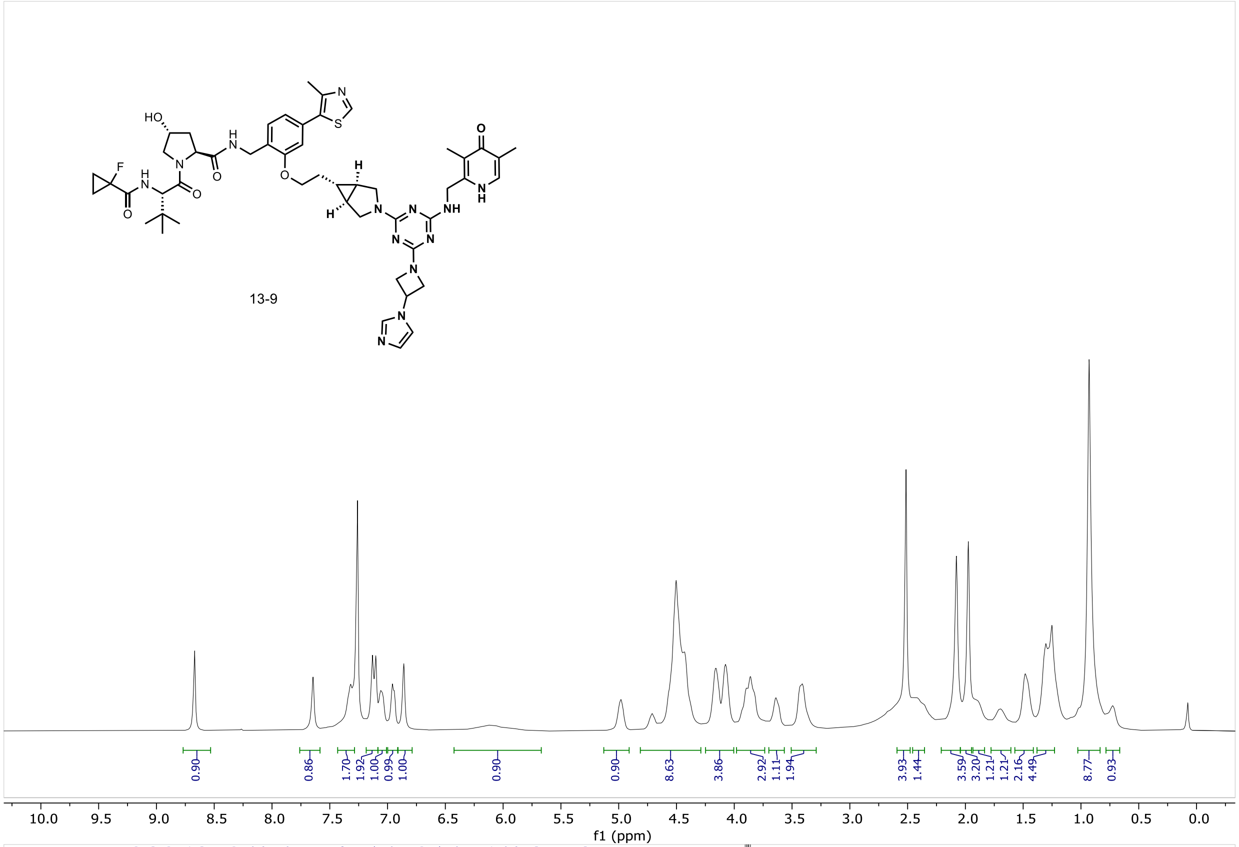

13-9
0.90
0.86
1.70
1.92
1.00
0.99
1.00
0.90
0.90
8.63
3.86
2.92
1.11
1.94
3.93
1.44
3.59
3.20
1.21
1.21
2.16
4.49
8.77
0.93
f1 (ppm)

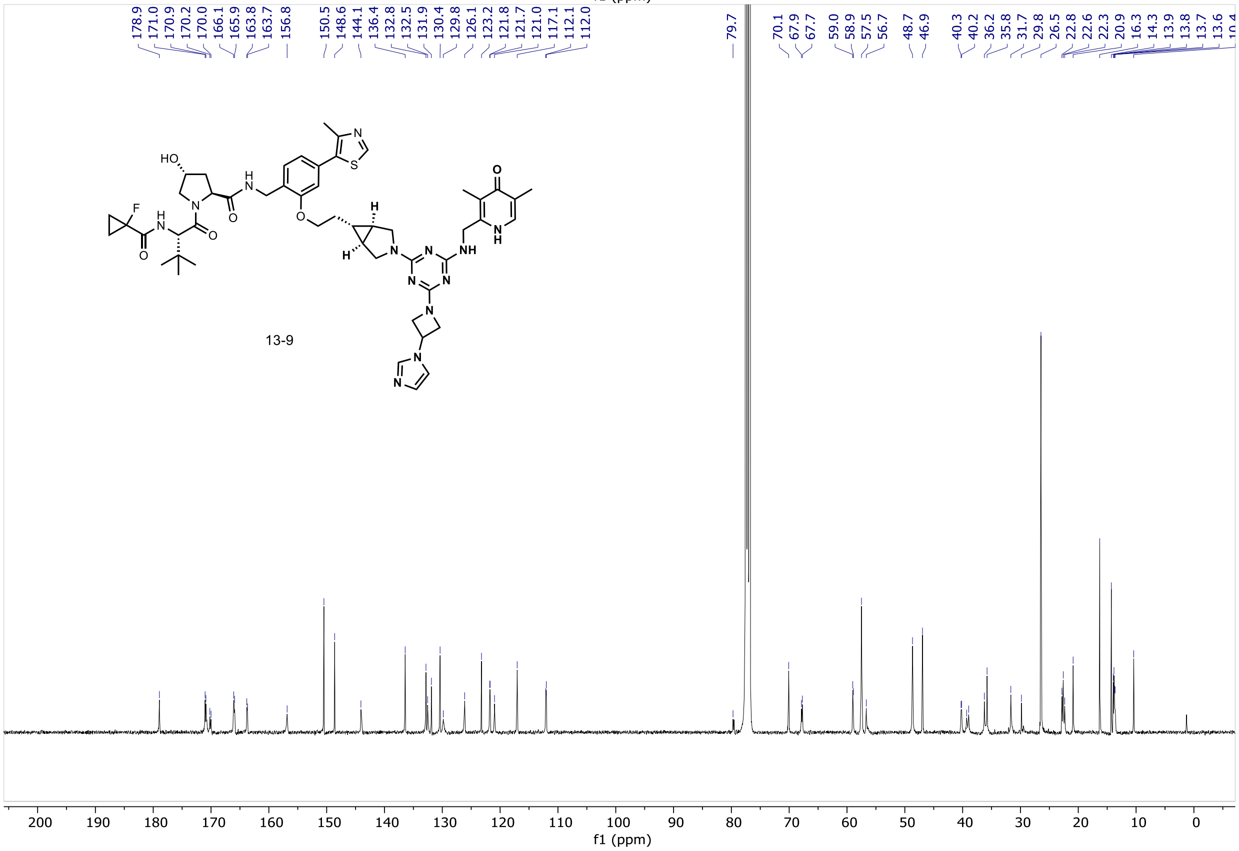

178.9
171.0
170.9
170.2
170.0
166.1
165.9
163.8
163.7
156.8
150.5
148.6
144.1
136.4
132.8
132.5
131.9
130.4
129.8
126.1
123.2
121.8
121.7
121.0
117.1
112.1
112.0
79.7
70.1
67.9
67.7
59.0
58.9
57.5
56.7
48.7
46.9
40.3
40.2
36.2
35.8
31.7
29.8
26.5
22.8
22.6
22.3
20.9
16.3
14.3
13.9
13.8
13.7
13.6
13-9
f1 (ppm)

#### SUPPLEMENTARY FIGURES

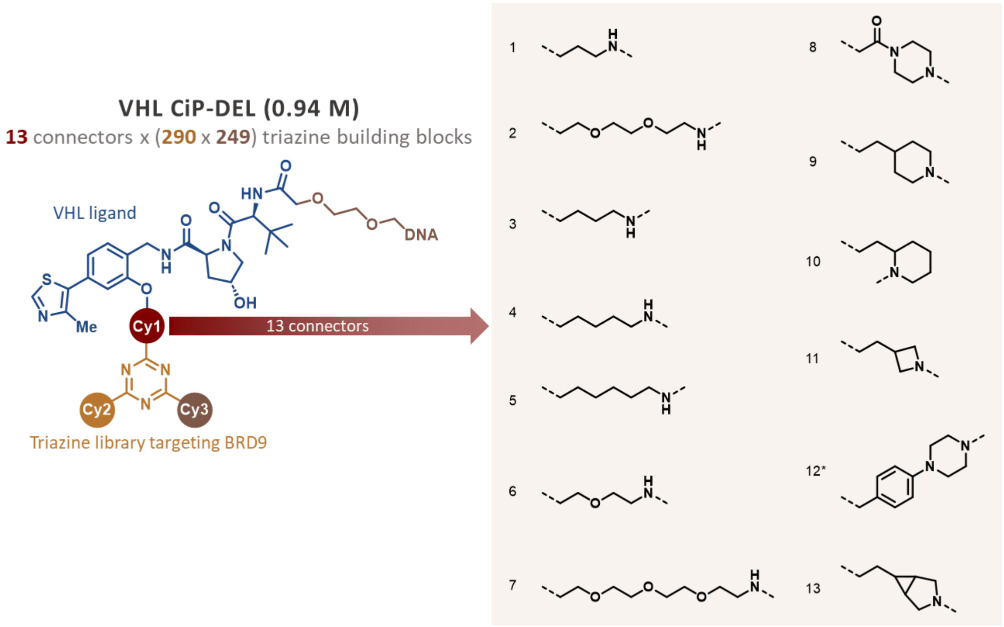

**Supplementary Figure S1. Structures of 13 connectors (cycle 1) in the VHL CIP-DEL.** Connectors 9, 11, 12, 13 were predominantly enriched in the CIP-DEL screen with BRD9 and VCB. *Note that connector 12 is attached to the VHL ligand via the carbon para to the phenolic oxygen. See **Supplementary Figure S4** for more details.

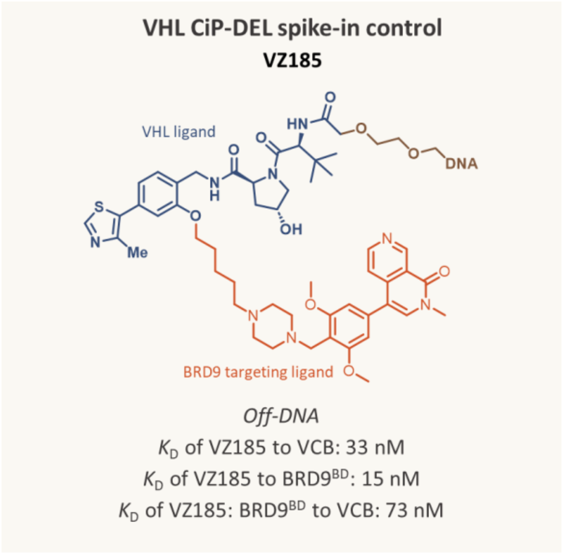

**Supplementary Figure S2. Structure of DNA-encoded VZ185 that was spiked into the VHL CIP-DEL.** Binary and ternary $K_D$s of VZ185 (off-DNA) are reported values[2].

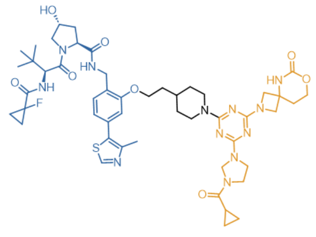

**9-1** (9, 275, 131)

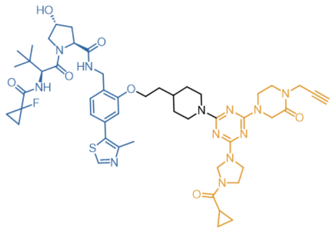

**9-2** (9, 275, 235)

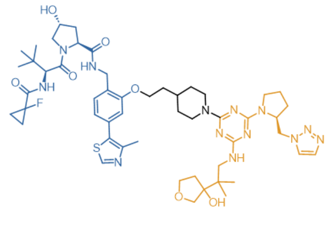

**9-3** (9, 87, 174)

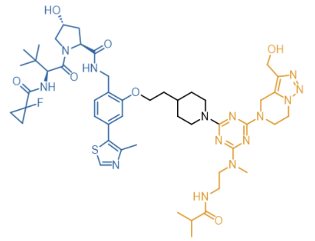

**9-4** (9, 126, 158)

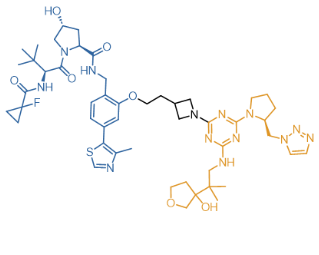

**11-1** (11, 87, 174)

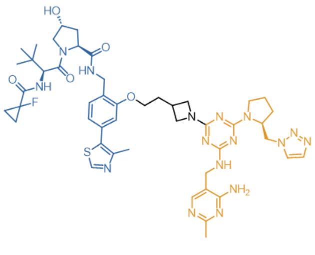

**11-2** (11, 197, 174)

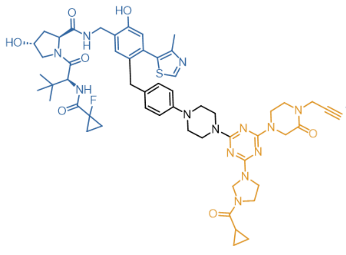

**12-1** (12, 275, 235)

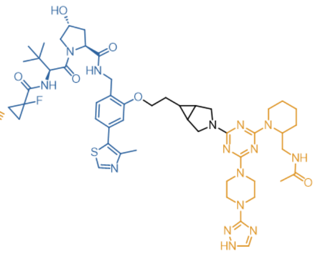

**13-1** (13, 163, 109)

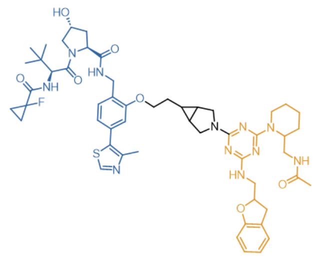

**13-2** (13, 253, 109)

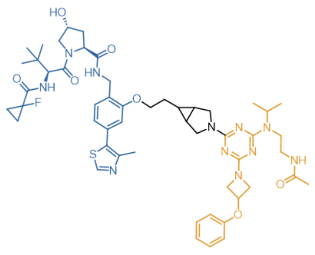

**13-3** (13, 117, 161)

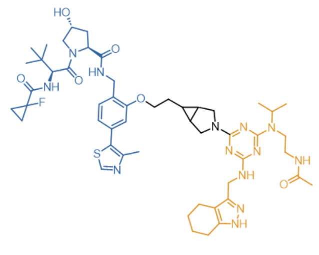

**13-4** (13, 117, 12)

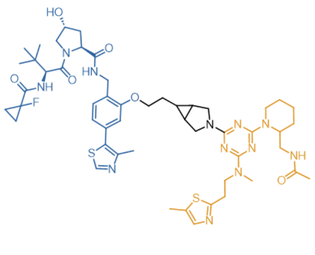

**13-5** (13, 130, 109)

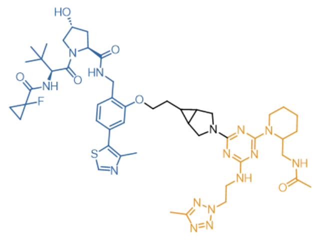

**13-6** (13, 18, 109)

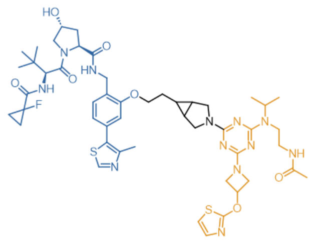

**13-7** (13, 117, 156)

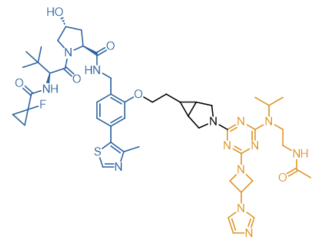

**13-8** (13, 117, 185)

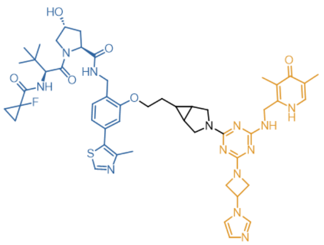

**13-9** (13, 52, 185)

**Supplementary Figure S3. Structures of off-DNA compounds that were enriched in the CIP-DEL screen with BRD9 and VCB.** All compounds have the VHL ligand (blue) connected to a substituted triazine (orange) via a connector (black). Compound short names were shown in bold. (x, y, z) identifies the compound by (connector, triazine's appendage building block 1, triazine's appendage building block 2), equivalent to (cycle 1, cycle 2, cycle 3) in the library.

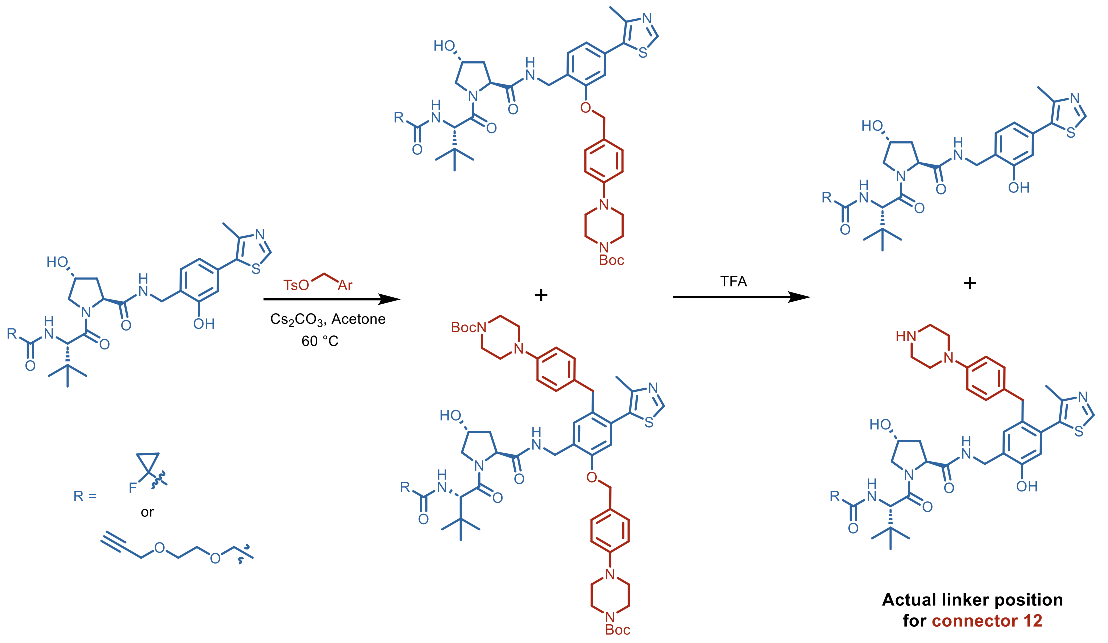

**Supplementary Figure S4. Connector 12 was connected to the VHL ligand via its carbon para to the anticipated phenolic oxygen.** Different from other connectors in the library, both C- and O- alkylations were observed when performing phenol alkylation reaction with connector 12, due to the high reactivity of the benzyl tosylate. However, the newly formed C-O bond was unstable and underwent cleavage upon treatment with TFA in the subsequent step; the resulting C-alkylation product was the only isolable product included in the library.

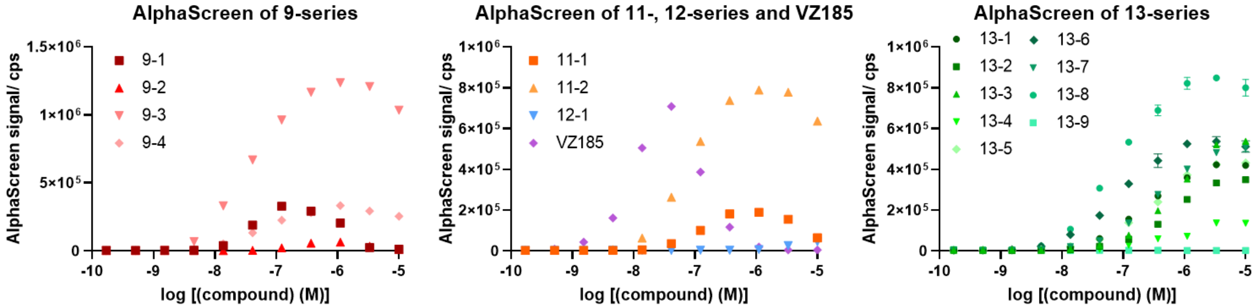

**Supplementary Figure S5. Ternary complex formation of off-DNA compounds or positive control VZ185 with purified recombinant BRD9 and VCB, as measured by AlphaScreen assays.** Data shown are 2 technical replicates ± SD, which have been independently reproduced. Each sample mixture contains 20 µL of 50 nM BRD9, 20 nM VCB, off-DNA compounds which were 3-fold serial diluted from 10 µM to 0.17 nM, 0.02 mg/mL acceptor beads, and 0.02 mg/mL donor beads, in 25 mM HEPES pH 7.4, 150 mM NaCl, 0.05% Tween-20 (w/v). Under the tested conditions, there was generally less hook effect for the 13-series compared to compounds such as 9-1, 11-1 and VZ185.

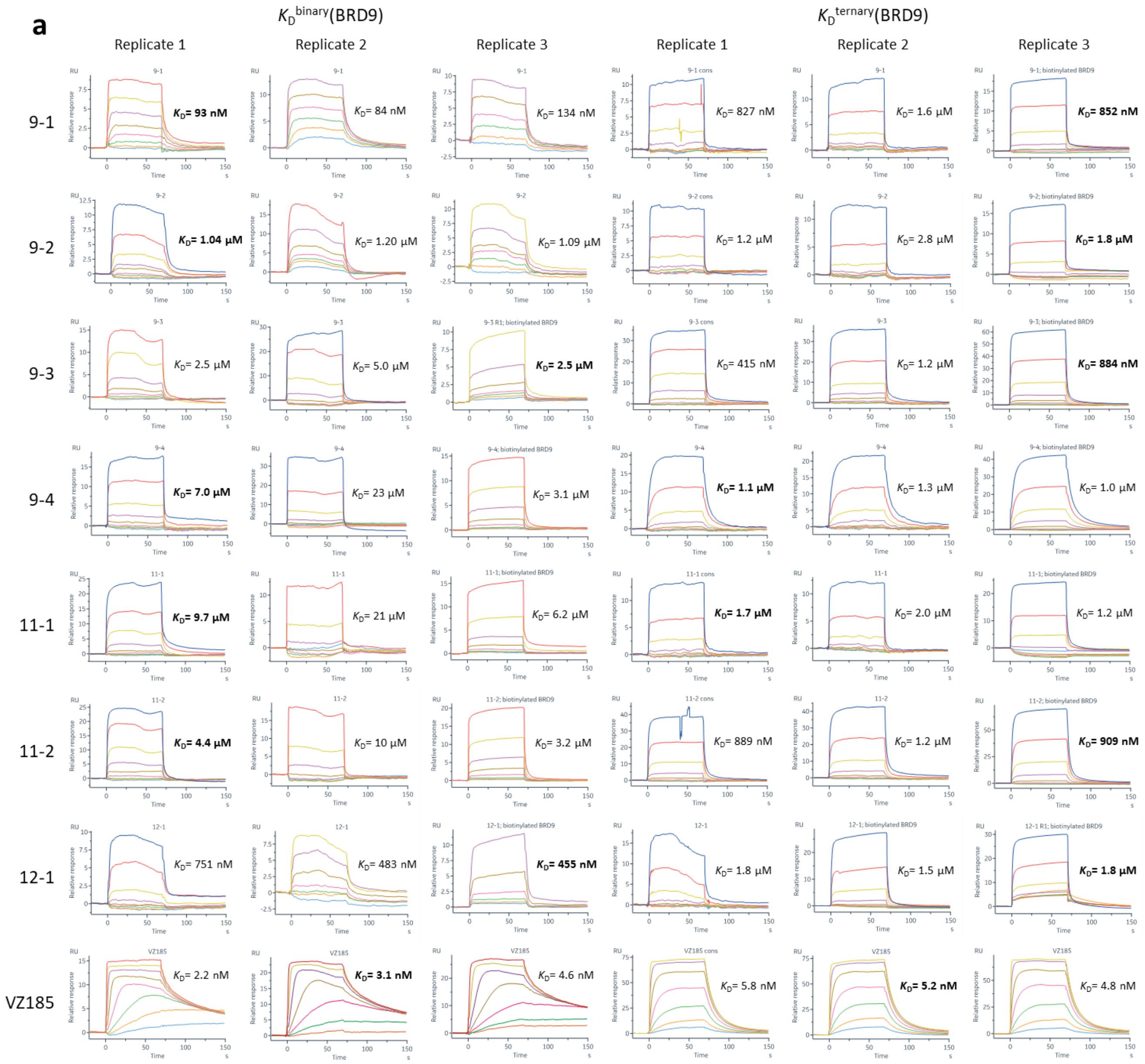

(continued on next page)

**b**

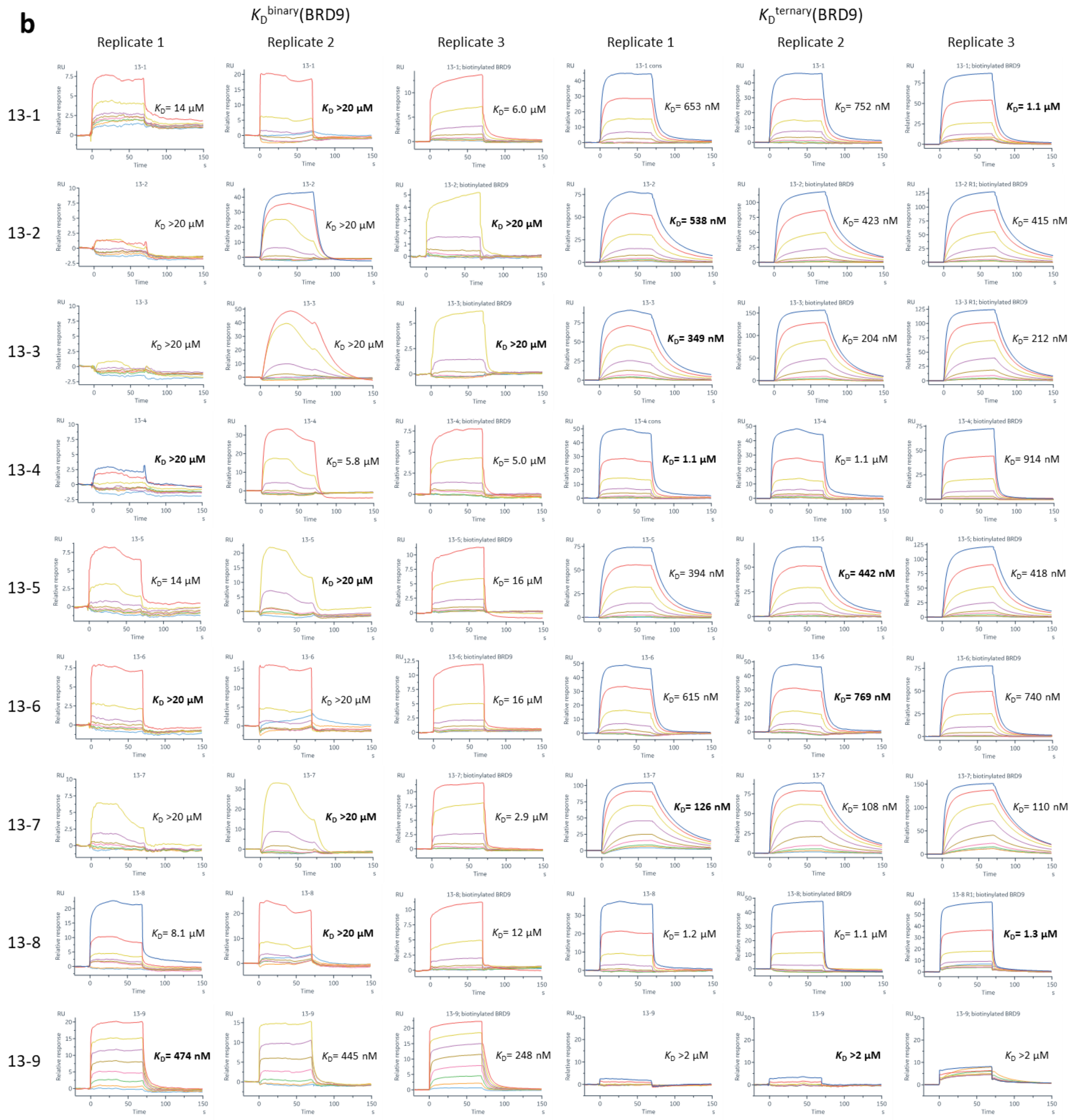

**Supplementary Figure S6a-b. Binary and ternary *K*Ds of off-DNA compounds or the positive control VZ185 with biotinylated BRD9 immobilized on streptavidin chip, as measured by SPR.** Compounds were 3-fold diluted from 20 μM and 2 μM for binary and ternary $K_D$ measurements with BRD9, respectively. For ternary $K_D$ measurements with BRD9, compounds were pre-incubated with 2 μM VCB. SPR running buffer: 20 mM HEPES, 100 mM NaCl, 0.005% tween-20 (w/v), 0.2 mM TCEP, 1% DMSO. Three independent replicates were shown. $K_D$s were obtained by steady state fit. $K_D$ values used for calculating the reported cooperativity are in bold. See ‘Methods’ for details.

**c**

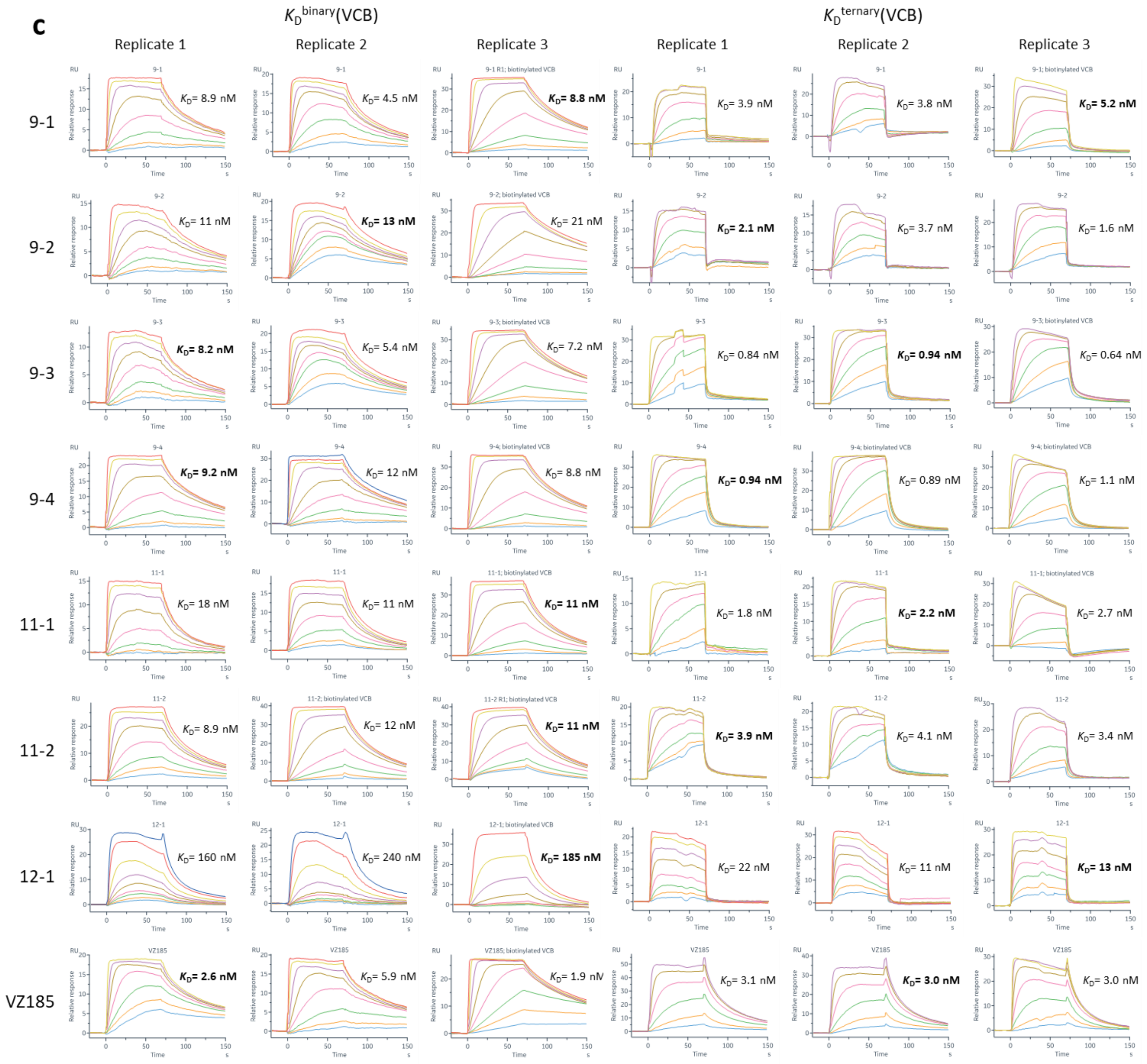

(continued on next page)

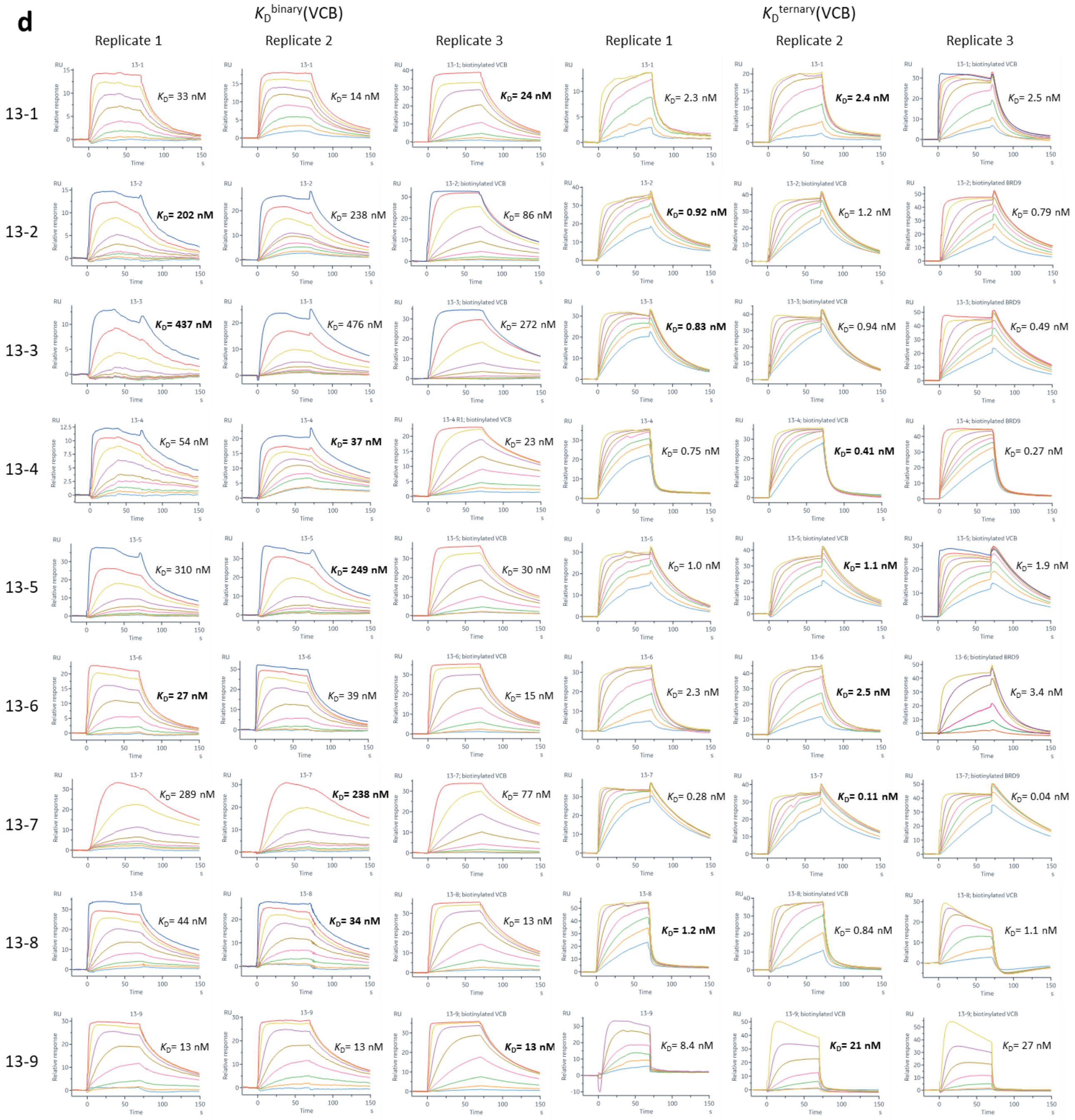

**Supplementary Figure S6c-d. Binary and ternary *K*Ds of off-DNA compounds and the positive control VZ185 with biotinylated VCB immobilized on streptavidin chip, as measured by SPR.** Compounds were 3-fold diluted from 2 μM for binary and ternary $K_D$ measurements with VCB. For ternary $K_D$ measurements with VCB, compounds were pre-incubated with 20 μM BRD9 and the sample compartment was cooled to 15 °C for BRD9 stability. SPR running buffer: 20 mM HEPES, 100 mM NaCl, 0.005% tween-20 (w/v), 0.2 mM TCEP, 1% DMSO. Three independent replicates were shown. $K_D$s were obtained by steady state fit. $K_D$ values used for calculating the reported cooperativity are in bold. See 'Methods' for details.

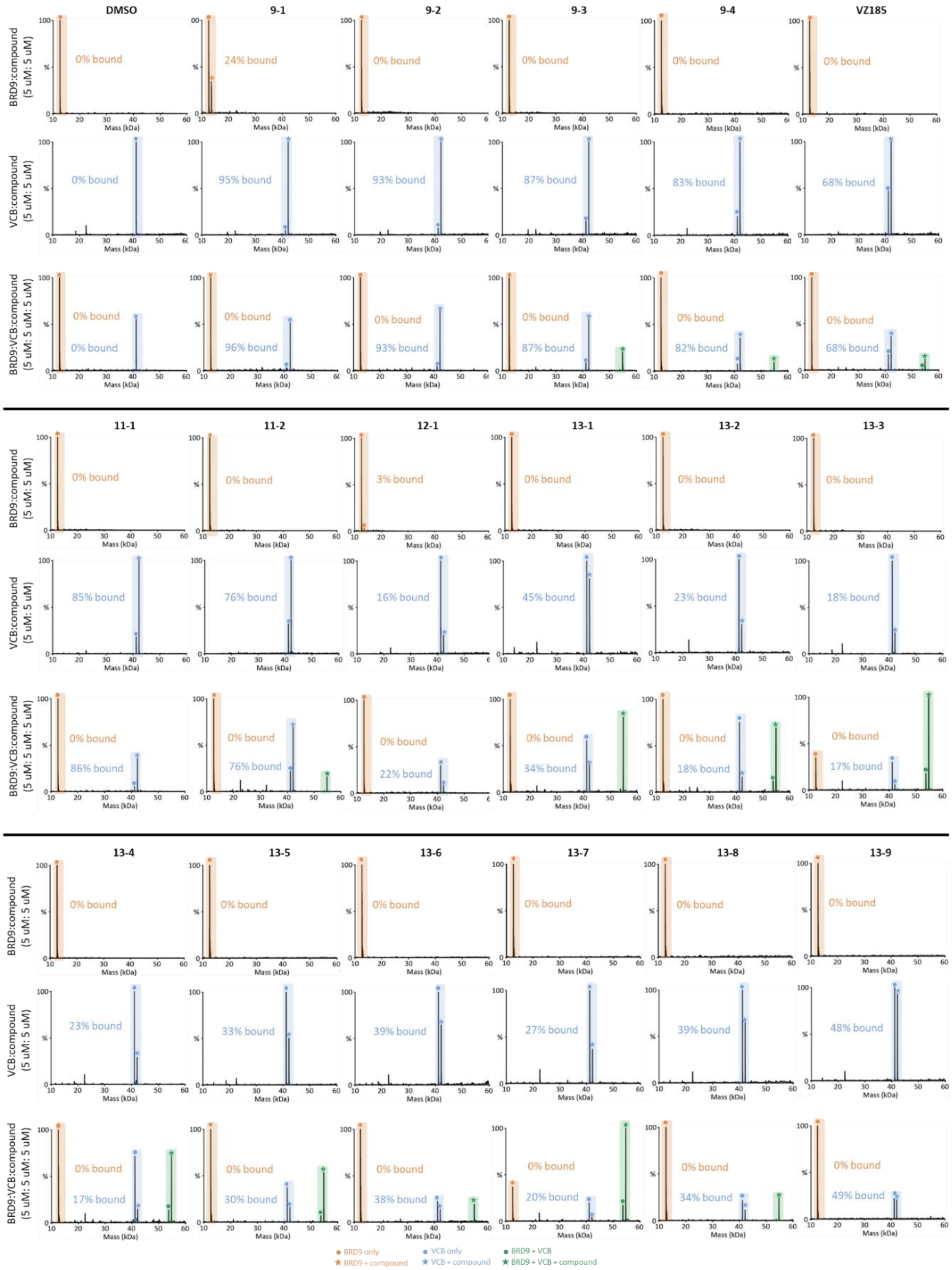

**Supplementary Figure S7. Non-denaturing MS analysis of off-DNA compounds or positive control VZ185 binding to BRD9 alone, VCB alone, or a mixture of BRD9 and VCB.** Data shown are deconvoluted to the actual mass of the species. Desolvation energy: -5V. Concentrations of compound, BRD9 and VCB: each at 5 µM. Buffer: 20 mM ammonium acetate, pH 7.5, 1% DMSO. See 'Methods' for details. Despite weaking binary binding to VCB (blue), more ternary complexes (green) were observed for the more cooperative 13-series in general.

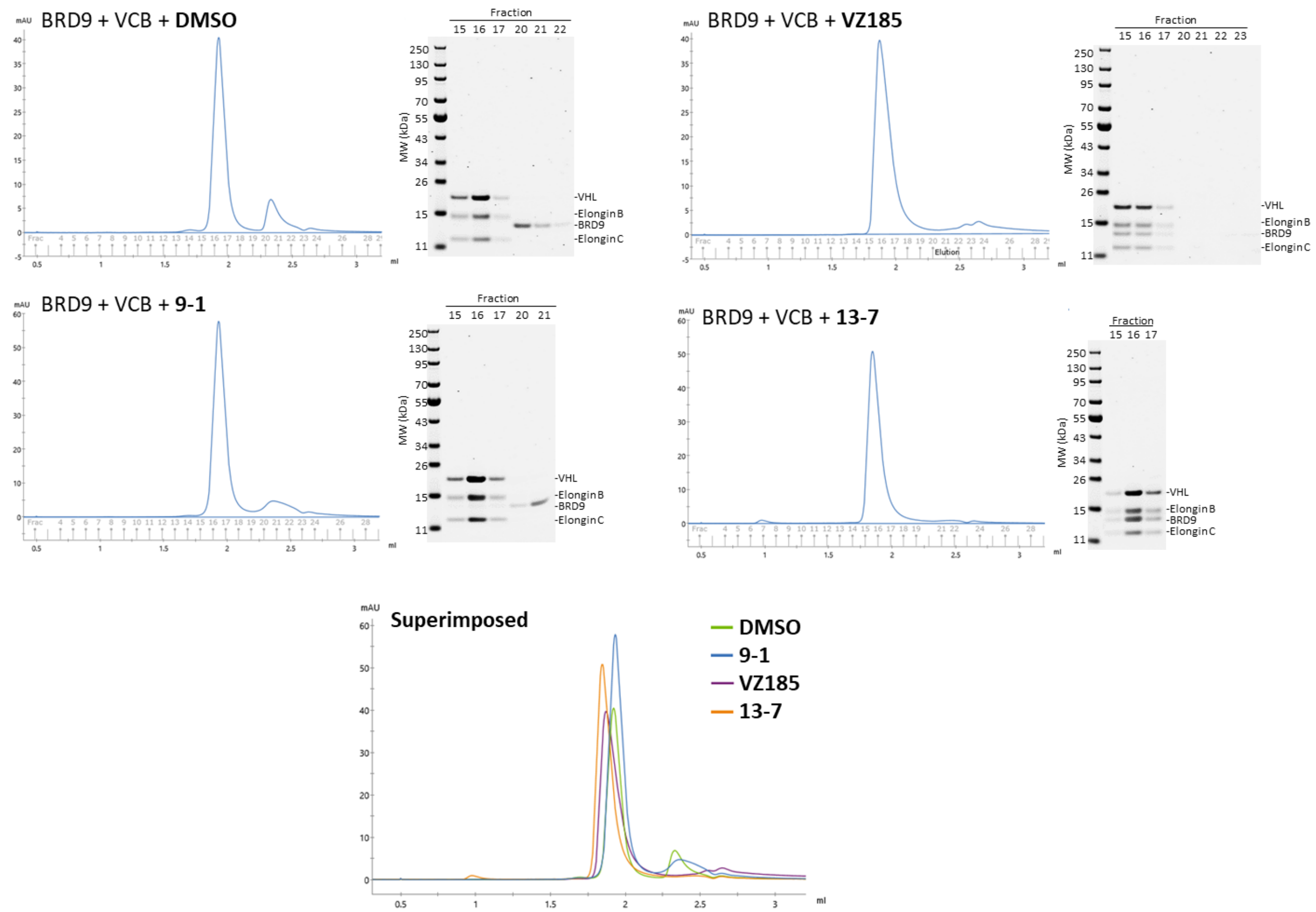

**Supplementary Figure S8. Size exclusion chromatography (SEC) UV traces of equimolar mixture of BRD9, VCB and compounds of interest.** The compound, namely 9-1, VZ185, 13-7 or DMSO only control, was mixed with BRD9 and VCB (each at 30 μM) in 20 mM HEPES, 100 mM NaCl, pH 7.4. See 'Methods' for details. The superimposed SEC traces showed a rightward shift of the elution peak towards lower molecular weight was observed from 13-7 (VCB and BRD9 co-elution), VZ185 to 9-1/DMSO (VCB only). SDS-PAGE gel analysis was performed on the fractions of the elution peaks. For 9-1/DMSO, VCB and BRD9 were eluted separately, whereas for 13-7, and to a lesser extent VZ185, co-elution of VCB and BRD9 was observed.

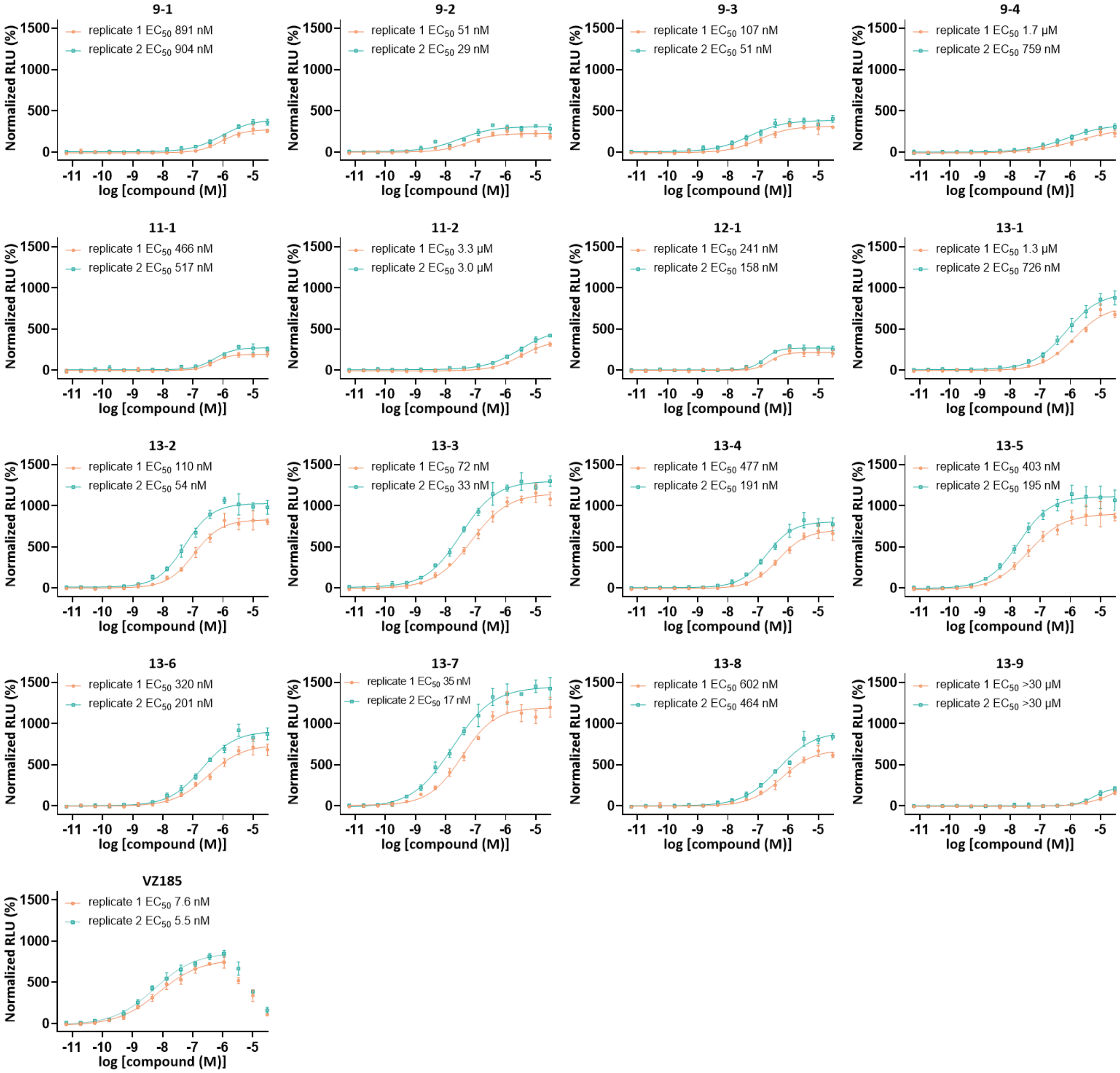

**Supplementary Figure S9. Ternary complex formation of off-DNA compounds or positive control VZ185 with SmBiT-BRD9 and VHL-LgBiT in HEK293T cells, as measured by NanoBiT assays.** Data shown are 2 independent replicates (orange and blue) of mean ± SD, n = 4 technical replicates. See 'Methods' for details.

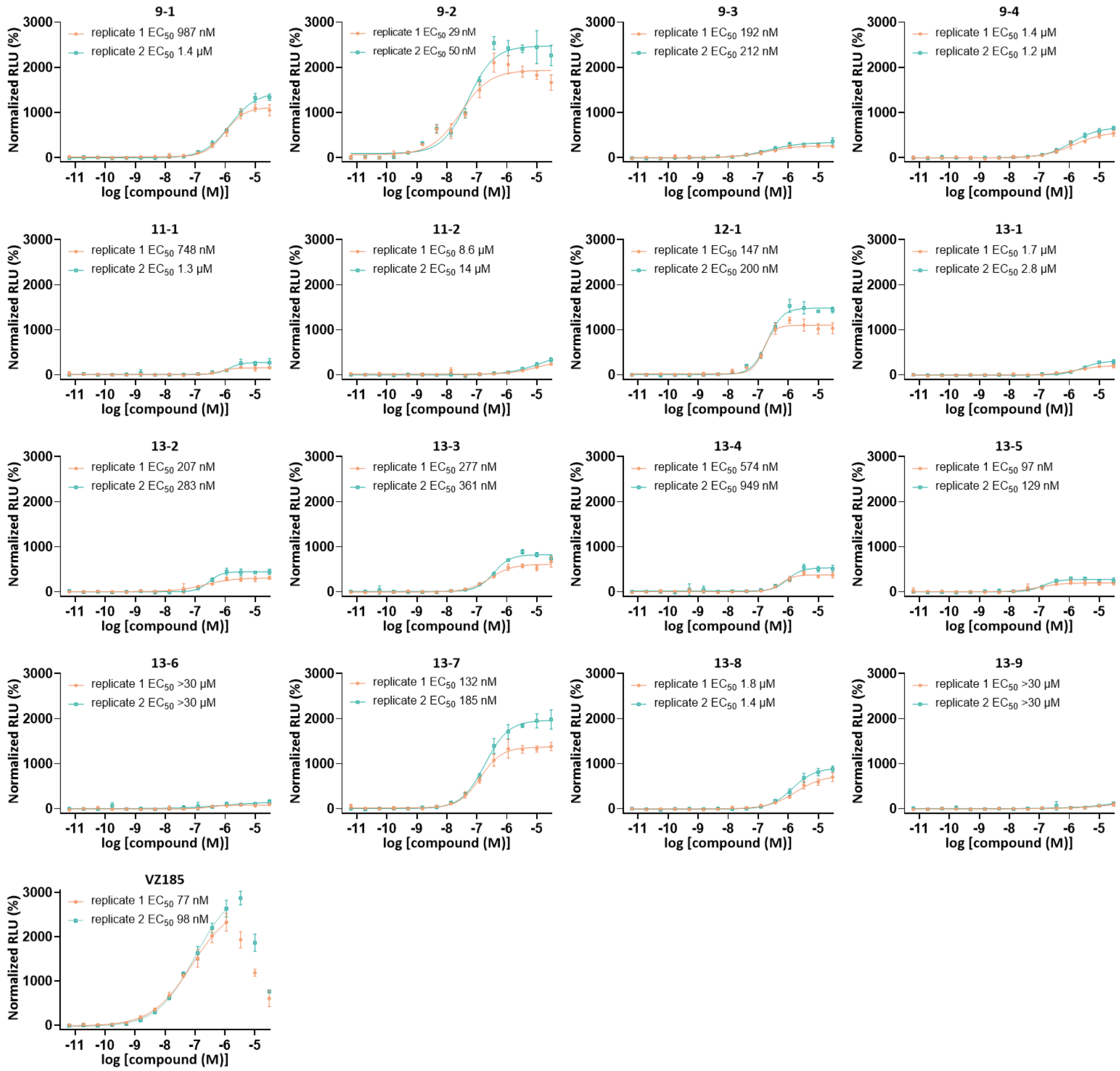

**Supplementary Figure S10. Ternary complex formation of off-DNA compounds or positive control VZ185 with SmBiT-BRD7 and VHL-LgBiT in HEK293T cells, as measured by NanoBiT assays.** Data shown are 2 independent replicates (orange and blue) of mean ± SD, n = 4 technical replicates. See 'Methods' for details.

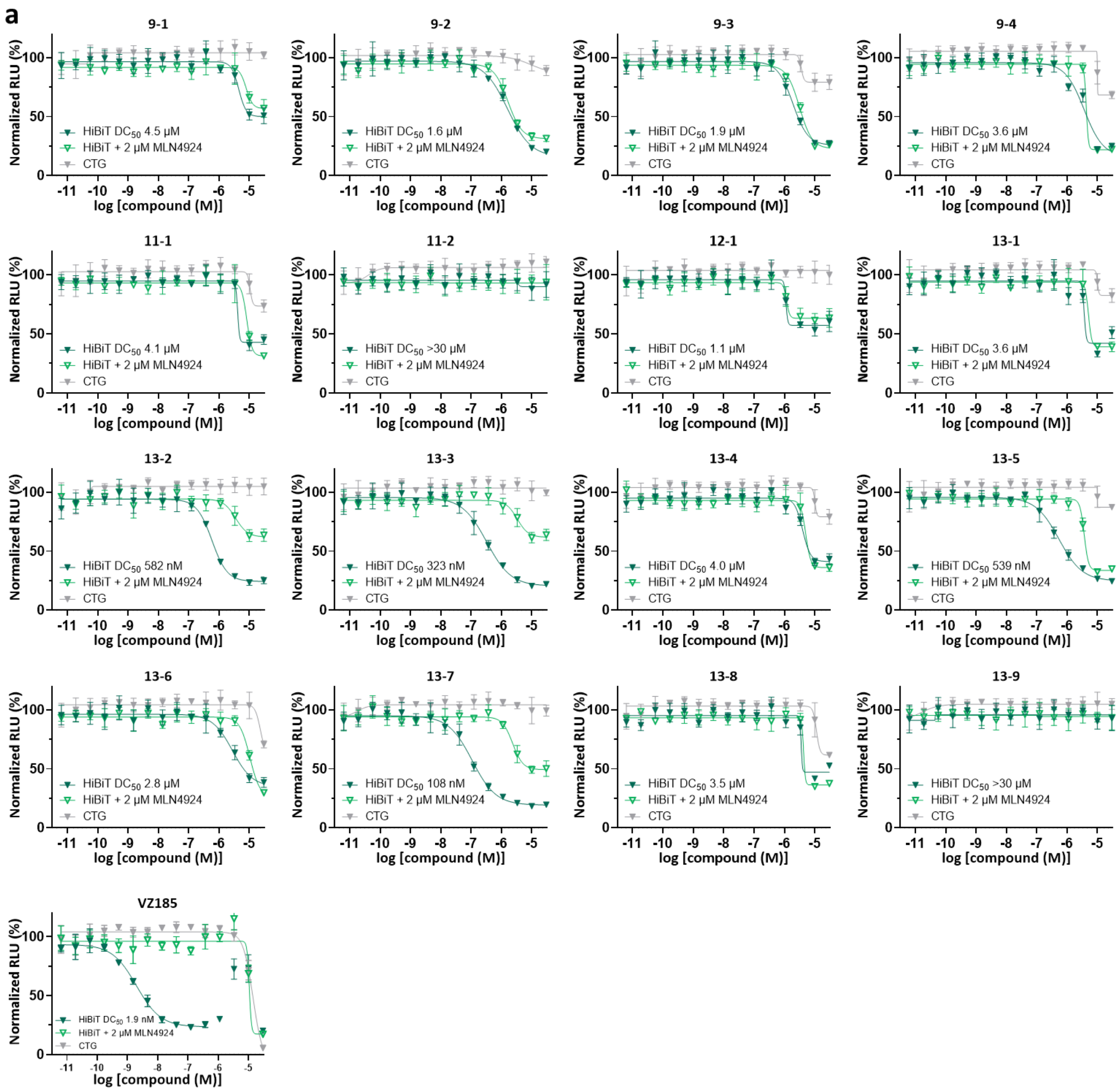

(continued on next page)

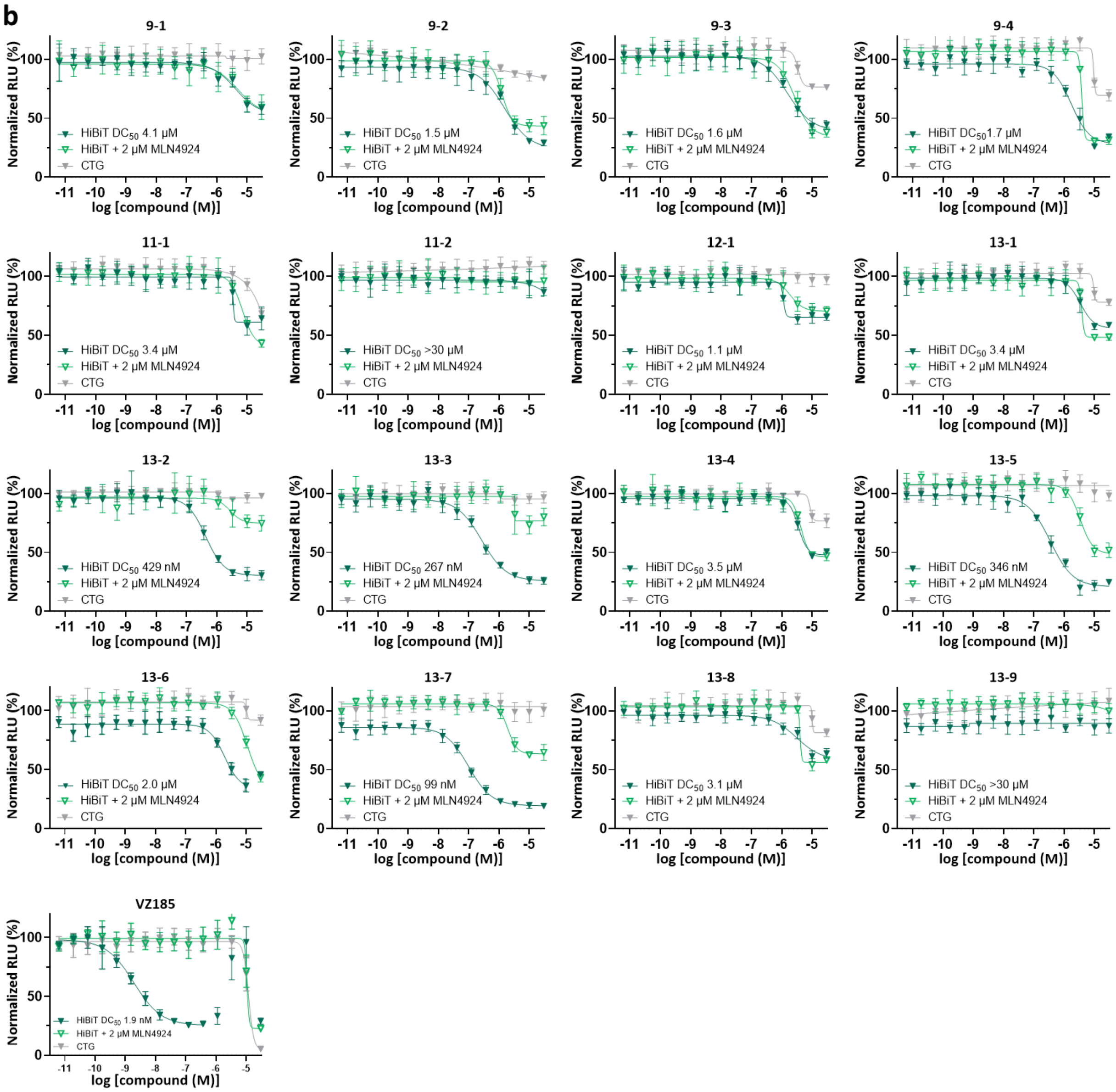

**Supplementary Figure S11. BRD9 degradation induced by off-DNA compounds or positive control VZ185 in HEK293T cells, as measured by HiBiT assays.** In addition to the standard HiBiT assays measuring BRD9-HiBiT degradation (dark green), neddylation inhibitor MLN4924[3] (2 µM) was added to access that the BRD9-HiBiT degradation is Cullin-RING E3 ligase, namely VHL, dependent (green). Degradation was not fully reversed by MLN4924 for some off-DNA compounds, for which we are not fully clear of the reasons after multiple investigations (see **Supplementary Figures S11** and **S12**). CellTiter-Glo (CTG) assays (gray) were performed in parallel to verify that the decrease in the luminescent signals of the HiBiT assays was due to BRD9-HiBiT degradation but not cell death. Data shown are 2 independent replicates (**a** and **b**) of mean ± SD, n = 4 technical replicates. See 'Methods' for details.

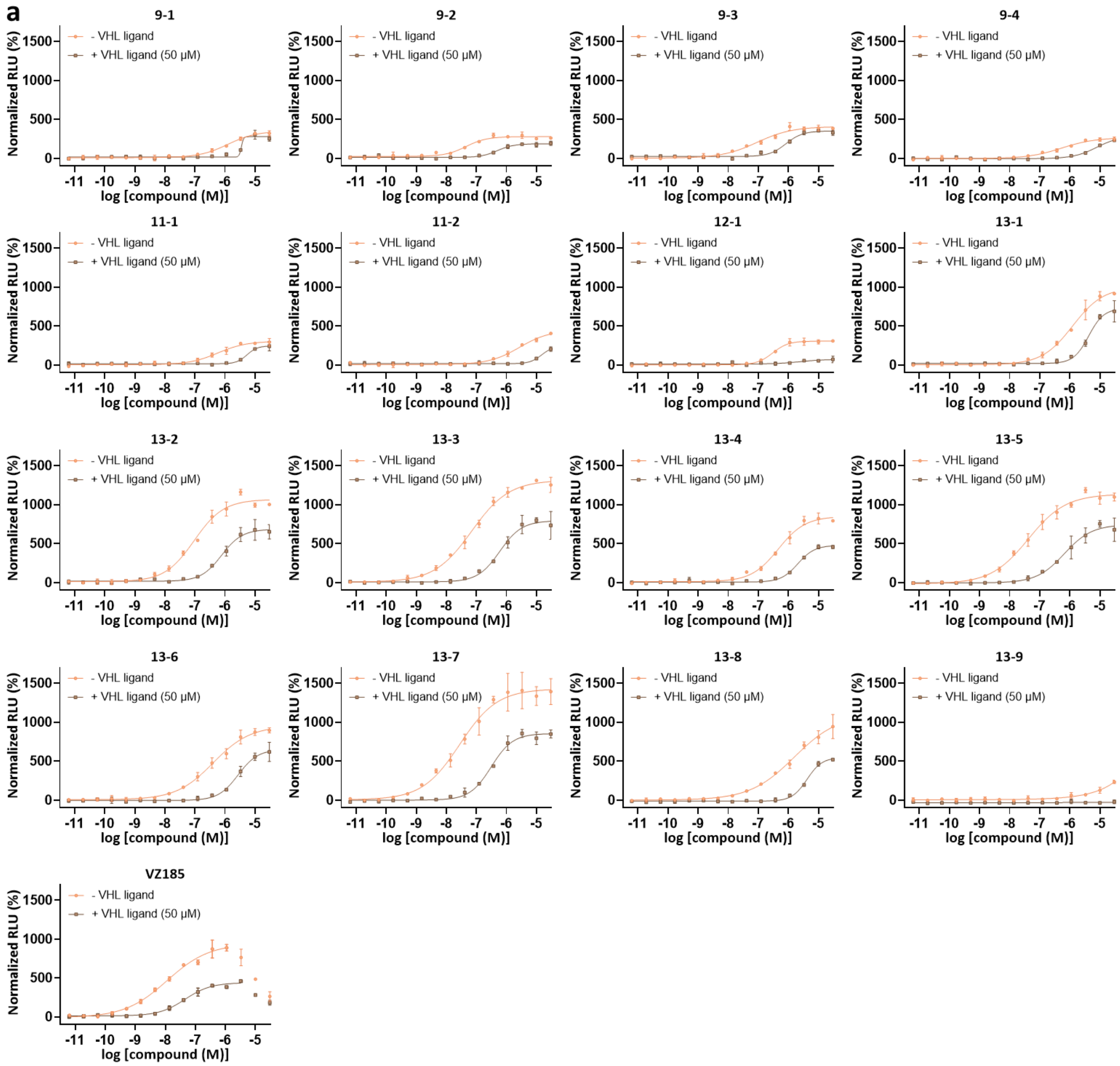

(continued on next page)

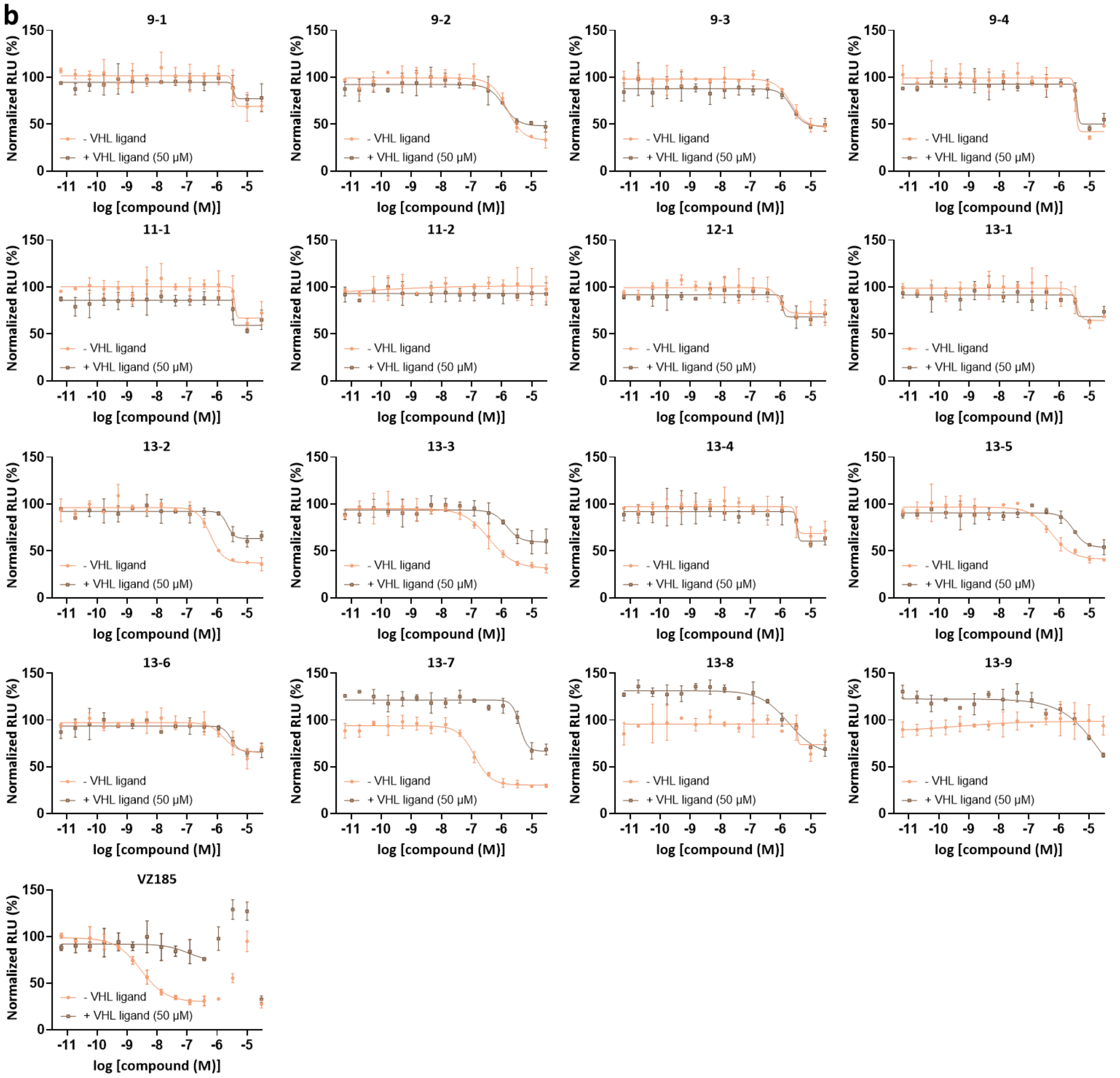

**Supplementary Figure S12. NanoBiT and HiBiT assays of off-DNA compounds or positive control VZ185 with VHL ligand (50 μM) competition.** The VHL ligand used to compete with the off-DNA compounds and VZ185 for binding to VHL is VH101[4], the VHL binding ligand of VZ185. **a.** Ternary complex formation of off-DNA compounds or positive control VZ185 with SmBiT-BRD9 and VHL-LgBiT in HEK293T cells, with and without VH101 (50 μM), as measured by NanoBiT assays. **b.** BRD9 degradation induced by off-DNA compounds or positive control VZ185 in HEK293T cells, with and without VH101 (50 μM), as measured by HiBiT assays. While $EC_{50}$s of the off-DNA compounds were shifted higher upon VH101 competition in NanoBiT assays, a similar extent of shift in the $DC_{50}$s was not observed for some compounds upon VH101 competition in HiBiT assays, suggesting that the BRD9-LgBiT degradation induced by those compounds was not fully VHL dependent. Data shown are mean ± SD, n = 2 technical replicates. See 'Methods' for details.

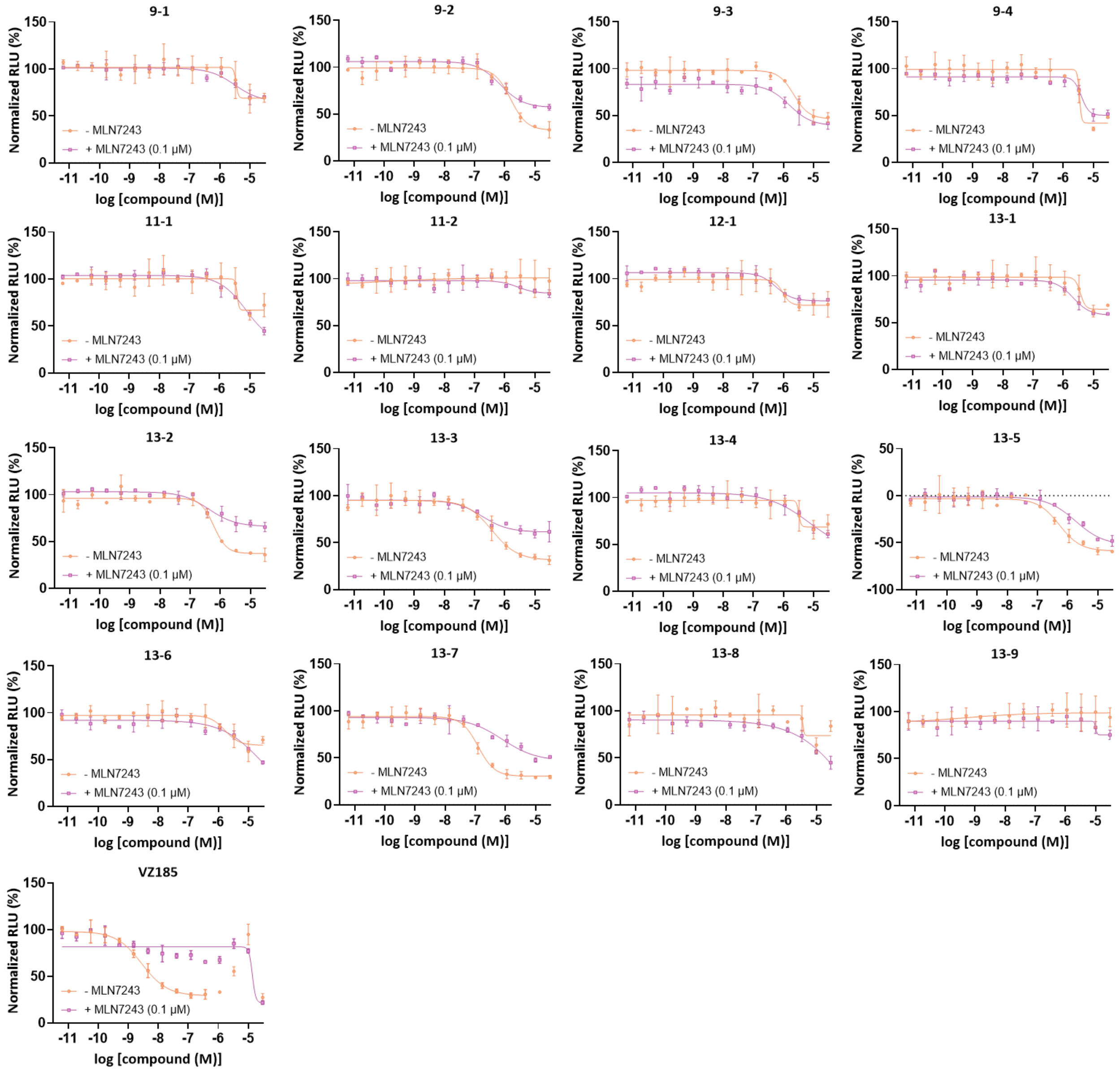

**Supplementary Figure S13. BRD9 degradation induced by off-DNA compounds or positive control VZ185 in HEK293T cells, with and without MLN7243 (0.1 µM), as measured by HiBiT assays.** MLN7243 is an inhibitor of ubiquitin-activating enzyme and its addition in HiBiT assays allows the assessment of whether BRD9-LgBiT degradation was ubiquitin-dependent. A shift of $DC_{50}$s upon the addition of MLN7243 in HiBiT assays was not observed for some compounds, suggesting that the BRD9-LgBiT degradation by those compounds was not fully ubiquitin dependent. Data shown are mean ± SD, n = 2 technical replicates. See 'Methods' for details.
